# Metagenome-based metabolic modelling predicts unique microbial interactions in deep-sea hydrothermal plume microbiomes

**DOI:** 10.1101/2022.11.06.515352

**Authors:** Dinesh Kumar Kuppa Baskaran, Shreyansh Umale, Zhichao Zhou, Karthik Raman, Karthik Anantharaman

## Abstract

Deep-sea hydrothermal vents are abundant on the ocean floor and play important roles in ocean biogeochemistry. In vent ecosystems such as hydrothermal plumes, microorganisms rely on reduced chemicals and gases in hydrothermal fluids to fuel primary production and form diverse and complex microbial communities. However, microbial interactions that drive these complex microbiomes remain poorly understood. Here, we use microbiomes from the Guaymas Basin hydrothermal system in the Pacific Ocean to shed more light on the key species in these communities and their interactions. We built metabolic models from metagenomically assembled genomes (MAGs) and infer possible metabolic exchanges and horizontal gene transfer (HGT) events within the community. We highlight possible archaea–archaea and archaea–bacteria interactions and their contributions to robustness of the community. Cellobiose, D-Mannose 1-phosphate, O_2_, CO_2_, and H_2_S were among the most exchanged metabolites. Ten microbes, including eight bacteria and two archaea, were identified as key contributors. These microorganisms uniquely enhanced the metabolic capabilities of the community by donating metabolites that cannot be produced by any other community member. Archaea from the DPANN group stood out as key microbes, benefiting significantly from accepting metabolites from other members of the microbiome. Amino acids were the key auxotrophy driving metabolic interactions in the community. Finally, over 200 horizontal gene transfer events were predicted in the community, the majority of which were between *Gammaproteobacteria* and *Alphaproteobacteria*. Overall, our study provides key insights into the microbial interactions that drive community structure and organisation in complex hydrothermal plumes and deep-sea microbiomes.

## Introduction

Deep-sea hydrothermal vents are abundant across mid-ocean ridges, back-arc basins, and volcanoes on the ocean floor. Hydrothermal vents emit hot fluids rich in reduced chemicals, gases, and metals. These hot fluids (up to 400 °C) mix with the cold seawater (2-4 °C) to form vent chimneys and hydrothermal plumes. While vent chimneys are formed by precipitation and solidification of minerals, hydrothermal plumes are turbulent environments that can rise hundreds of meters from the seafloor to achieve neutral buoyancy and spread across the ocean over hundreds to thousands of kilometers^1, 2^. Microbial activity in hydrothermal vents is driven by the presence of potential energy sources such as H_2_S, Fe, Mn, CH_4_ and H_2_^3, 4^. Hydrothermal plumes are associated with a strong redox gradient formed due to the presence of highly reduced electron donors from vents which mix with the cold seawater rich in electron acceptors such as oxygen and nitrate, which can provide microorganisms with sufficient energy to fix carbon into biomass^1, 2^. Microbial communities thrive in such harsh environments partly due to metabolic interactions associated with their ability for interdependent utilization of substrates^5–7^. Hydrothermal vent microbial communities form the base of the food chain in these environments and have been shown to play a significant role in mediating various elemental cycles in ocean ecosystems^8, 9^. Hydrothermal vent habitats also harbour the growth of a very specialized set of organisms like giant tubeworms (vestimentiferans), Pompeii worms (*Alvinella pompejana*), vesicomyid clams, vent mussels (*Bathymodiolus elongatus*), scaly-foot snails (*Chrysomallon squamiferum*), and crabs (*Kiwa* spp.). Flora and fauna in this ecosystem flourish as a result of close symbiosis with chemosynthetic microbes consisting primarily of bacteria and archaea.

Increasingly, omics-based approaches have focused on the study of uncultivated microorganisms and there is a growing recognition that microbial metabolic interactions are key in maintaining microbial community structure and function in diverse environments, including in the deep-sea. The problem of unculturability in microbes that pervades different ecosystems makes it a challenge to isolate and characterize metabolic interactions using conventional microbiological tools^10^. Metabolic interactions are the threads holding a community of microbes together^11–13^. Therefore, studying these interactions can enable us to gain mechanistic insights into community function^14, 15^. While metagenome-based interpretation of microbial genomes (as implemented in the software METABOLIC) can predict auxotrophies that can imply the presence of microbial interactions, metabolic modeling represents a more powerful approach in predicting metabolic interactions. To this end, *in silico* modelling approaches offer a promising alternative to study microbial metabolism in general^16^, and community metabolic interactions in particular^17–19^. Genome-scale metabolic models^20^ can be built using whole genomes or metagenomically assembled genomes (MAGs) of microbes^21, 22^. These models capture the metabolic capabilities of an organism. Metabolic models of all known members of a community allow us to study community interactions using various graph-based and constraint-based approaches^17–19^.

In hydrothermal vents and plumes, prior studies have focused on the genomic characterization of microbial and metabolic diversity, but little is known about the role of metabolic dependencies and interactions in these microbiomes. In this study, we use deep-sea hydrothermal vents in Guaymas Basin in the Pacific Ocean as a model system to study the functional underpinnings of microbial communities in hydrothermal vent plumes and the interactions that keep them together. In particular, this study focuses on: (i) the coexistence of archaea and bacteria and the cross-domain metabolic interactions between them, and (ii) evolutionary processes in hydrothermal plume microbial communities, including horizontal gene transfers (HGTs)^1^. Our study implicates the metabolite environment in which these microbes grow to play a major role in determining interactions.

Through a combination of computational methods, we illustrate the central role of metabolic interactions in the organisation of such complex microbial communities, and identify the key microbes mediating these interactions. We employed a graph-based approach for predicting the influence of every member on the metabolism of other microorganisms in the community. In order to study microbial evolution in these extreme conditions, we predicted all possible HGT events in the community from metagenomic data. Microbial communities are often involved in HGT to maintain uniform cooperation. HGTs provide advantages to individual microbes by reducing the susceptibility of a member to opportunistic invasion and extreme environmental conditions. Here, we identified several genes that were possibly horizontally transferred, and map the cellular processes impacted by them. Overall, our study illustrates the potential of computational approaches like metabolic modelling to unravel the complex web of metabolic and genetic interactions that drive the organisation of microbial communities.

## Materials and Methods

Figure 1 provides a pictorial representation of the approaches used in this study. This research work starts with building genome-scale metabolic networks of microbes of the communities from their respective metagenomically-assembled genomes.

**Figure 1.**
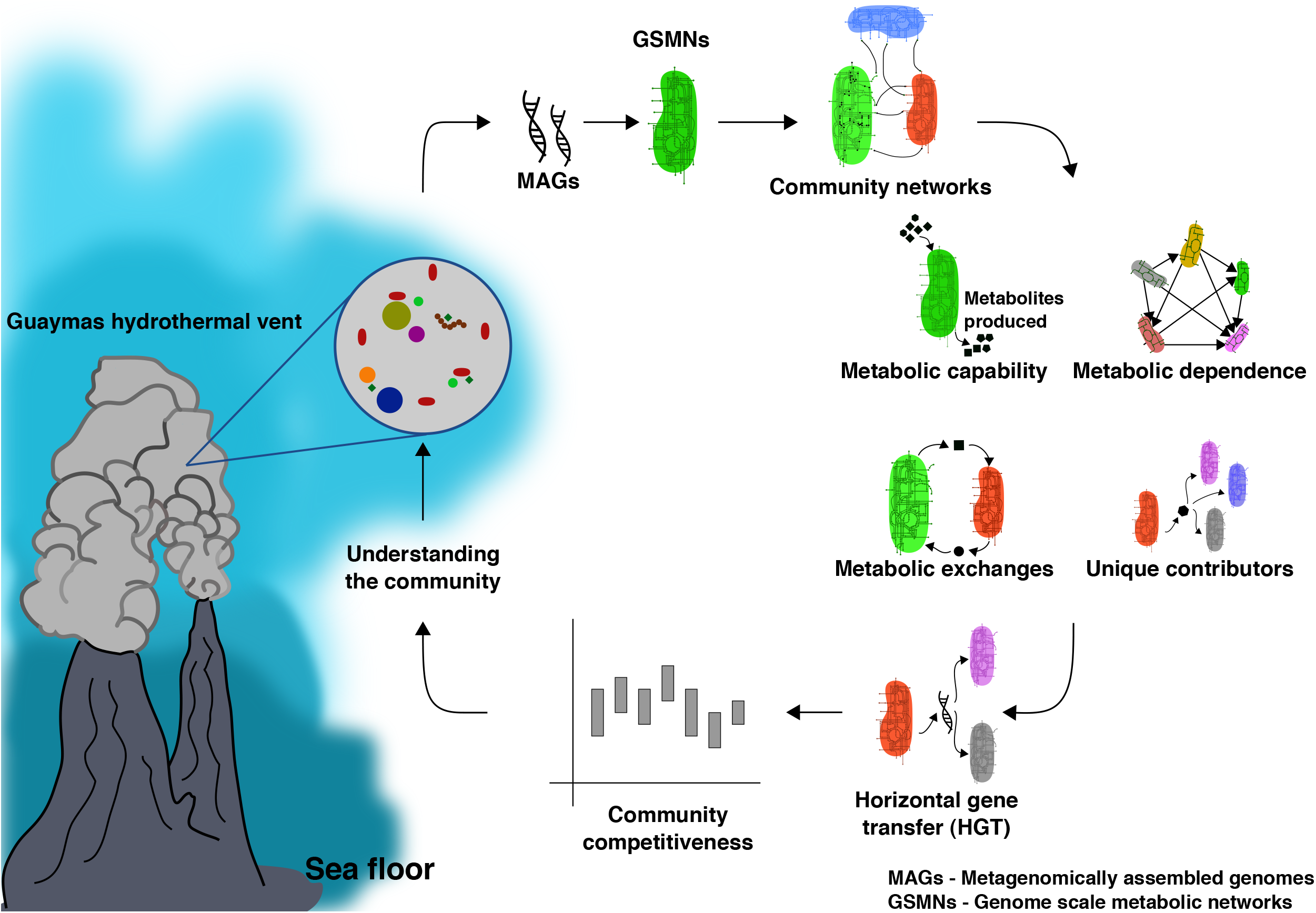
Summary of the process followed in studying the Guaymas microbiome. This study starts with the construction of genome-scale metabolic networks using tools like CarveMe and MetQuest from the metagenomically-assembled genomes (MAGs) of corresponding microbes. This allows us to further construct metabolic networks for two-member communities and higher-order communities. The next step involves predicting the characteristics of the community, like the metabolic capability of microbes in the community, metabolic dependence of the community, metabolic exchanges possible in the community and unique contributors in the community. Further, genetic interactions between microbes were predicted using MetaCHIP, a tool for predicting horizontal gene transfers (HGTs). As the last step, the competitiveness of the community is determined at a different community scale and compared against other microbial communities.

### Metagenomic datasets and model building

The Guaymas hydrothermal plume microbiome data^23^ consists of metagenomically assembled genomes (MAGs) of 98 microbes. These MAGs fulfil the MIMAG high-quality criteria^24^ on completeness and contamination which are available in our GitHub repository. Only these MAGs were used for further reconstructing the genome-scale metabolic models. Briefly, the samples were collected from plumes of Guaymas Basin, the Gulf of California and high-throughput shotgun sequencing was performed on the DNA. Metagenomic sequences were assembled into scaffolds and binned into corresponding metagenomically assembled genomes (MAGs). A detailed description of sampling, DNA extraction, and processing of MAGs is described in detail elsewhere.^23, 25^.

In this study, 98 MAGs corresponding to 98 OTUs were used to construct draft genome-scale metabolic models using CarveMe^21^. Along with this, we also used the data from a recent comparative study of the East Pacific Rise microbiome^26^ for studying the level of competition in the community (MRO analysis) discussed later in the article (see Studying the level of competition in the community). Given that these bacteria and archaea remain mostly uncultured and poorly characterized, the metabolic models were reconstructed without any gap-filling to avoid any biases. Hence, these draft metabolic models only represented the metabolism captured in the MAGs and which could be annotated.

### Determining the metabolite environment of Guaymas hydrothermal vent ecosystem

Four different metabolite conditions were used for performing all the analyses in this study: (1) Guaymas media (GM) which simulates conditions in the hydrothermal plumes of Guaymas Basin-MMJHS medium^27^ with methanol, (2) JW1 media^28^ with sulfite, thiosulfate, elemental sulfur, sodium sulfide, cysteine hydrochloride, methanol, (3) Marine Broth 2216^29^ with sulfite, thiosulfate, elemental sulfur, sodium sulfide, cysteine hydrochloride, methanol and (4) components of all three media combined (referred as “all-media” hereon).

### Predicting metabolic capabilities of microbes in the community

MetQuest^30^, a Python package built based on a graph-theoretic algorithm was employed to predict metabolic reactions that can be active and inactive in the given media conditions. This is achieved in two steps:

1. Constructing metabolic networks by assembling reactions into pathways using a dynamic programming-based approach.
2. Identifying all the reactions that are active (visited) and inactive (stuck) for a given set of starting/seed metabolites.

These seed metabolites are essentially the components of nutrient media on which the community needs to be grown or simulated. Since the algorithm requires only the topological information of metabolic networks, just the draft metabolic reconstructions of microbes are sufficient. The components in the media are important because the analyses performed in this study depends mainly on the environmental metabolome in which they are present (**Supplementary File S1**). This is due to the fact that metabolic support received or provided by a microbe to other members of the community varies with the media conditions. The metabolites that can be produced from the active reactions tells the metabolic capability of the microbes in the given media.

### Predicting metabolic dependencies of microbes in the community

Metabolic dependence is the dependence of one microbe on another microbe in the community for the activation of certain inactive (stuck) metabolic reactions. A reaction is active only when all the required substrates are available; this unavailability of substrates gives rise to dependencies. It was observed that the number of stuck reactions decreased when microbes were in a community versus when in individual state. This was due to the activation of previously inactive reactions led by availability of metabolites through the exchange of metabolites from other microbes. These reactions are referred as relieved reactions. A score called Metabolic support index (MSI)^31^ was used to determine this metabolic dependence of microbes. The formula for calculating MSI goes as follows:

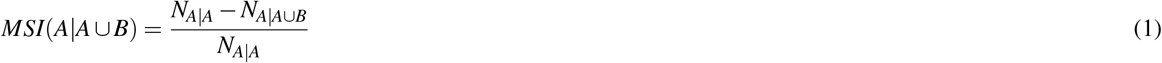

where N_A|A∪B_ represents the number of stuck reactions (reactions that are inactive in the given media condition) in *A* in the presence of *B* and N_A|A_ is number of stuck reactions in A when A is in isolation. Each reaction stuck/not executable is the loss of a metabolic capability of the metabolic network and MSI calculates the gain of metabolic capability. MSI gives distinct values for both the members of a pair, i.e., MSI of *A* in *A* ∪ *B* (or just *AB*, for brevity) community is different from MSI of *B* in *AB*,and hence it is a directional quantity. As an example, if MSI of A in AB is 0.041, this means that 4.1% of inactive reactions in microbe A can be activated by microbe B by exchange of required metabolites that were not available to microbe A in the absence of B. This value can be as high as one (*MSI* = 1) and as low as zero (*MSI* = 0). This step is called MSI analysis and was performed for all possible microbial pairs 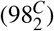 in the community.

### Visualising pairwise interaction networks

In order to visualise the results of pairwise MSI analysis, metabolic interaction networks were constructed. Different types of network visualisation were used viz., Cytoscape^32^ for visualising interactions between the microbes. In the “MSI network”, each node corresponds to the microbe and an edge between them indicates a potential interaction, i.e. a non-zero MSI value. Since MSI is directional, the interactions are captured via directed networks. The node on the arrowhead side is the “receiving” microbe while the node on the source side is the “supporting” microbe.

Another way of visualising the metabolic interactions was using chord diagrams. The chord diagrams were generated using the R package Chord diagrams, using home-grown scripts **(shared via GitHub)**. For this, initially the microbes were grouped into their corresponding microbial classes and then the interaction between each of the 98 microbes with microbial classes of Guaymas microbiome was represented using the chord diagrams. Again, the node on the arrowhead side is the “receiving” microbe/class while the node in the source side is the “supporting” microbe/class, and the chord thickness was mapped to the number of microbes in a class that interact with the target microbe. All the networks generated for the archaea and bacteria under study are available in **Supplementary File S2**.

### Predicting possible metabolic exchanges in all microbial pairs

Metabolic exchanges are the metabolites transferred from one microbe to another leading to the revival of stuck reactions. A list of stuck and relieved reactions was obtained for all the microbes in the respective communities. The reactants of relieved reactions that are transport reactions are the metabolites received during exchange.

### Identifying higher order interactions (CSI analysis)

#### Support offered by a group of microbes to the community

Here, the microbes in the community were pooled into different clusters based on the microbial classes they belong to such that each microbial class forms a cluster. There were 24 clusters formed corresponding to the 24 microbial classes present in the Guaymas community. The support offered by a cluster as a whole on the community can be determined by knocking out clusters and studying the reactions relieved in the presence of a particular cluster. Considering *X* as the microbial community and A the cluster to be removed, then *Ã* is the community without the cluster *A*. This can be represented as *Ã* = (X – A)^33^.

Then the formula for support index becomes

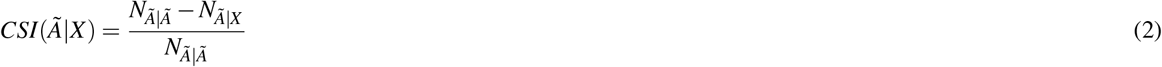

where *N_Ã|X_* is the number of stuck reactions in the community in the presence of cluster *A*, and *N_Ã|Ã_* captures the number of stuck reactions in the community when cluster *A* is removed from the community.

#### Unique contributors of the community

A unique contributor of a community is a microbe that has the potential to expand the metabolic niche of a community by contributing a unique metabolite to the community, thereby activating metabolic capabilities in the microbes. In order to determine the potential of a microbe A to support its community, the metabolic network of the community can be simulated with and without microbe A. Then, the support offered by A is the fraction of reactions relieved in the presence of A. Here, *X* as the microbial community and *A* the microbe to be removed, then *Ã* is the community without the microbe A. This can be represented as *Ã* = (*X* – *A*). The formula for support index is the same as Eq. 2. Every member of the community can be knocked out one by one to study the support offered by every microbe in the community. By this method any unique contributors in the community can be identified.

### Predicting HGT events

HGT events can be studied from metagenomic datasets of a community using MetaCHIP^34^. HGT analysis was performed using MetaCHIP v1.7.5 on all phylum, class, order, family and genus levels of taxonomic classification. Broadly, MetaCHIP first clusters query MAGs according to phylogenies and performs an all-versus-all blastn for all genes across genomes (parametric step). Next, the blastn matches for each gene is compared across taxa and is considered to be an HGT event if the best match comes from a non-self taxa. MetaCHIP then uses a phylogenetic approach to (i) reconcile differences between species and gene trees using RANGER-DTL^35^ and (ii) identify the direction of the putative transfer event. The enumerated HGT events can be visualised using the *circlize* package in R. Finally, egg-NOG mapper^36^ is used to map the HGT genes to corresponding functional categories.

### Studying the level of competition in the community

It is possible to predict the level of competition in a community by knowing the nutrient requirements of microbes in the community. We used SMETANA^37^ to calculate the metabolic resource overlap (MRO), which is the maximal overlap of minimal nutrient requirements of members of a community. SMETANA is formulated as a mixed linear integer problem (MILP) that enumerates the set of essential metabolic exchanges within a community of *N* species with non-zero growth of the *N* species subject to mass balance constraints. SMETANA does not use any biological objective functions which makes it unique. For every member *i* in a group of *N* distinct microbes, SMETANA enumerates the set of minimal nutritional components required for growth, **M_i_**. Nutritional requirement sets **M_i_** were used to compute MRO as described in the original paper. For the comparative analyses across different ecosystems, 1000 random communities were generated for community sizes ranging from 2–10 for four different ecosystems, *viz*. Guaymas^23^, East Pacific Rise^26^, anaerobic digestion^38^ and the gut^39^. This analysis is called MRO analysis.

## Results

In this study, we use 98 MAGs described previously from Guaymas Basin hydrothermal plumes to understand metabolic interactions and evolution in hydrothermal systems. Both bacteria and archaea are abundant members of hydrothermal plume microbiomes, yet play distinct roles in these environments. In this study we draw various insights about the uncultured bacteria and archaea, including bacteria depending on abundant hydrothermally-derived sulfur. Our observations were drawn from four major *in silico* analyses, MSI analysis, CSI analysis, HGT analyses, and MRO studies performed on these microbes. Overall, 26 (15 archaea and 11 bacteria) out of 98 MAGs were the main focus of this research, though these analyses were performed on all 98 microbes of the community. In comparison to bacteria, archaeal biology is still extremely under-explored, and their metabolic and functional potential is not well studied primarily due to the difficulty of culturing them^40–42^. Archaea are known to play important roles in hydrothermal vent ecosystems, and throughout the pelagic oceans such as in ammonia oxidation and transformation of organic compounds^2, 43–45^. Therefore, in order to understand and highlight the functional importance of ‘microbial dark matter’ in hydrothermal plumes, a significant focus of this study is on the archaeal members of this community and their interactions with other archaeal and bacterial species in Guaymas basin (**Refer Supplementary Table 1** for the list of archaea in the community). The Guaymas archaeome comprises three classes, *Poseidoniia, Nanoarchaeia*, and *Nitrososphaeria*.

In any microbial community, the ability of a microbe to produce or consume a metabolite is subject to the metabolite/media environment those microbes inhabit. In this study, four different media conditions (GM media, JW1 media, marine broth 2216 and an all-media) were used to study this community. To arrive at these four media conditions, three suitable media used to replicate sea water were selected from literature viz., MMJHS medium^27^, JW1 media^28^, and Marine broth 2216^29^. To these media components, sulfur and methane sources (sulfite, thiosulfate, elemental sulfur, sodium sulfide, cysteine hydrochloride, methanol) were added to specialize them towards Guaymas hydrothermal vent systems thus obtaining GM media, JW media and marine broth 2216 (see Determining the metabolite environment of Guaymas hydrothermal vent ecosystem in Materials and Methods). Components of all three media are possible constituents of hydrothermal vent environments, hence having a synthetic media like all-media by combining all the three media might provide a closer representation of the habitat.

Many observations were made about the metabolic capability of microbes in different media and the implicated metabolic exchanges. Oxygen and ornithine were some of the most exchanged metabolites in all-media and JW1 media, but the microbes in GM media and marine broth 2216 were unable to produce oxygen resulting in the absence of their exchanges in these environments. L-serine was the only metabolite exchanged irrespective of media conditions. This observation shows the capability of the community to compensate an absence of a metabolite through exchange. This helps in maintaining the robustness of the community.

### Archaea–bacteria pairs show high interaction potential in the hydrothermal plume microbiome

In order to determine the influence of bacteria present in the ecosystem on the metabolism of archaea, pairwise MSI analysis was performed under four different media conditions. (Described in Determining the metabolite environment of Guaymas hydrothermal vent ecosystem in Materials and Methods). Briefly, in this analysis, a score called Metabolic Support Index (MSI) (Predicting metabolic dependencies of microbes in the community in Materials and Methods) is calculated for every possible pair of microbes (98*C*_2_ pairs), which measures the increase in metabolic capabilities of a microbe while in a community versus as an individual organism. Microbes in the community gain different metabolic capabilities through the exchange of metabolites. MSI provides distinct values for both the members of a pair, i.e., MSI of A in AB community is different from MSI of B in AB, and hence is a directional quantity.

We identified the most interesting archaea–bacteria microbial pairs on all four media based on high MSI scores. The highest MSI score observed in the Guaymas microbiome was 0.052 between an archaeon and a bacterium: *Flavobacteriaceae* bacterium UWMA 0314 →*Candidatus* Pacearchaeota archaeon UWMA 0287 (the arrow goes from donor to acceptor) in JW1 media, which was primarily due to the exchange of metabolites cellobiose and D-Mannose 1-phosphate. These metabolites activated many metabolic reactions in *Candidatus* Pacearchaeota archaeon UWMA 0287. In this interaction, *Flavobacteriaceae* bacterium UWMA 0314 is not predicted to receive any metabolite from its partner (MSI =0) in all four media. *Flavobacteriaceae* bacterium UWMA 0314 →Nitrosopumilus sp UWMA 0263, *Gammaproteobacteria* bacterium UWMA 0261 →*Candidatus* Pacearchaeota archaeon UWMA 0287 were other archaea–bacteria microbial pairs with high interaction potential in the Guaymas microbiome (Figure 2 represents all the pairwise interactions between *Candidatus* Pacearchaeota archaeon UWMA 0287 and other microbial classes). Among the ^98^C_2_=4753 pairs possible in the community, the main emphasis was given to those where the receiver acquires at least a 1% increase in the metabolic capability (i.e., *MSI* >= 0.01). **Refer Supplementary File S3 for the entire list of MSIs.**

**Figure 2.**
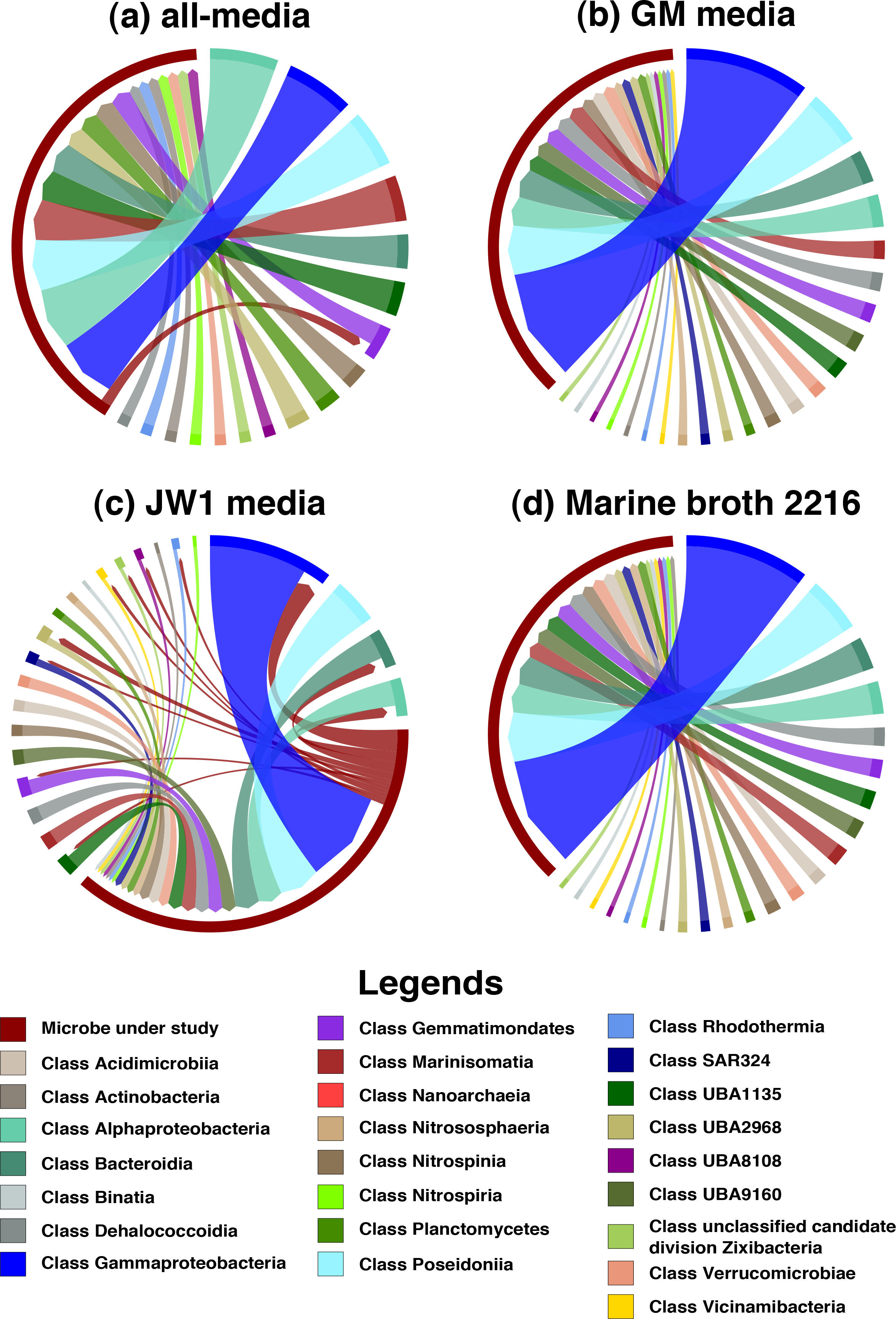
Pairwise MSI interactions of *Candidatus* Pacearchaeota archaeon UWMA 0287 with other microbes in the community. This chord diagram shows all possible metabolic interactions between *Candidatus* Pacearchaeota archaeon UWMA 0287 and other microbial classes present in the Guaymas microbiome in four different media conditions (a) all-media, (b) GM media, (c) JW1 media, (d) Marine Broth 2216. All the interacting microbes are grouped under their corresponding microbial class except *Candidatus* Pacearchaeota archaeon UWMA 0287. The chord starts from the donor microbe/class towards the recipient microbe/class. The thickness of the chord represents the number of microbes participating in the interaction from the same class. The colours are mapped to microbial classes.

In most of the archaea-bacteria interactions, archaea were always found to be on the “acceptor” side while bacteria “donate” metabolites. A possible explanation for this is that archaea have reduced metabolic capabilities than the bacteria in the Guaymas community. It is possible that the understudied nature of archaea manifests in a greater proportion of unannotated genes in their genomes leading to the impression of them having reduced metabolic capabilities. An MSI value (interaction) is always attributed to a set of exchanges leading to the gain of metabolic capabilities in the acceptor microbe. The metabolites frequently exchanged in the archaea–bacteria interactions mentioned above were cellobiose, D-Mannose 1-phosphate, O_2_, CO_2_, and H_2_S, among others, but the exchange of any one of these metabolites can lead to gain of comparatively greater metabolic capabilities in the acceptor microbe.

Though archaea–bacteria interactions were widely observed in GM media, JW1 media and all-media, they were lower in marine broth 2216. *Fuerstia* sp UWMA 0333 → *Candidatus* Pacearchaeota archaeon UWMA 0287, *Planctomycetaceae* bacterium UWMA 0346 → *Candidatus* Pacearchaeota archaeon UWMA 0287, *Sneathiellales* bacterium UWMA 0353 → *Candidatus* Pacearchaeota archaeon UWMA 0287, and *Gemmatimonadetes* bacterium UWMA 0339 → *Candidatus Nitrosopelagicus* sp UWMA 0359 were the only high potential archaea–bacteria interactions observed in marine broth 2216. Among these *Sneathiellales* bacterium UWMA 0353 → *Candidatus* Pacearchaeota archaeon UWMA 0287 was observed in all four media.

### Archaea-archaea interactions are dominated by DPANN archaea as acceptor microbes

The microbial interactions Marine Group II euryarchaeote UWMA 0266 → *Candidatus* Pacearchaeota archaeon UWMA 0287, Marine Group II euryarchaeote UWMA 0275 → *Candidatus* Pacearchaeota archaeon UWMA 0287, Marine Group II euryarchaeote UWMA 0279 → *Candidatus* Pacearchaeota archaeon UWMA 0287, Marine Group II euryarchaeote UWMA 0283 → *Candidatus* Pacearchaeota archaeon UWMA 0287, Marine Group II euryarchaeote UWMA 0350 → *Candidatus* Pacearchaeota archaeon UWMA 0287 were seen in GM media, JW1 media and all-media. Like in archaea–bacteria interactions, *Candidatus* Pacearchaeota archaeon UWMA 0287 were always the acceptors in these archaea–archaea interactions too. Cellobiose and CO_2_ exchanged from Marine Group II euryarchaeotes to Pacearchaeota has the potential to activate many metabolic capabilities in Pacearchaeota.

Unlike Pacearchaeota, archaea of class *Poseidoniia* can act as both acceptors and as donors in the Guaymas community. Interestingly, these archaea exhibited three distinct interaction patterns:

1. Marine Group II euryarchaeote UWMA 0266, Marine Group II euryarchaeote UWMA 0275, Marine Group II euryarchaeote UWMA 0279, Marine Group II euryarchaeote UWMA 0283, Marine Group II euryarchaeote UWMA 0350 and Marine Group II euryarchaeote UWMA 0352 showed similar interaction patterns.
2. Marine Group II euryarchaeote UWMA 0323, Marine Group II euryarchaeote UWMA 0328, Marine Group II euryarchaeote UWMA 0344, Marine Group II euryarchaeote UWMA 0357, Marine Group III euryarchaeote UWMA 0284 showed similar interaction patterns.
3. Marine Group III euryarchaeote UWMA 0340 was distinct from other members of *Poseidoniia*. The interaction pattern of this microbe was the sparsest in comparison to other members of this group.

Another significant archaea-archaea interaction involves *Candidatus* Nitrosopelagicus sp UWMA 0359 and Nitrosopumilus sp UWMA 0263 which belong to the class *Nitrososphaeria* (Figure 3 represents all the pairwise interactions between Nitrosopumilus sp UWMA 0263 and other microbial classes). These organisms show potential interactions among themselves in JW1 media and in all-media through the exchange of ornithine, putrescine, and H_2_S.

**Figure 3.**
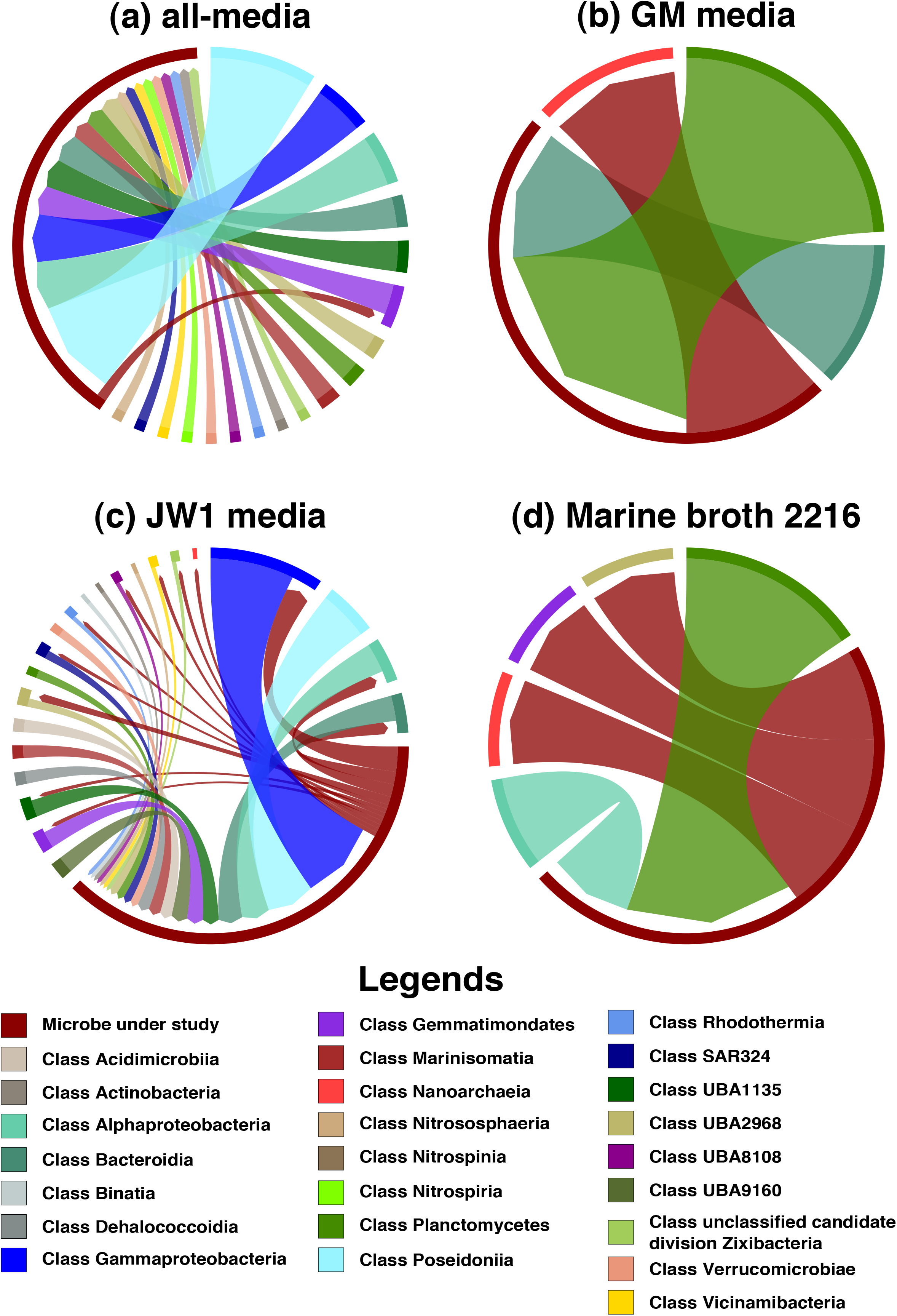
Pairwise MSI interactions of Nitrosopumilus sp UWMA 0263 with other microbes in the community. This chord diagram shows all possible metabolic interactions between Nitrosopumilus sp UWMA 0263 and other microbial classes present in the Guaymas microbiome in four different media conditions (a) all-media, (b) GM media, (c) JW1 media, (d) Marine Broth 2216. All the interacting microbes are grouped under their corresponding microbial class except Nitrosopumilus sp UWMA 0263. The chord starts from the donor microbe/class towards recipient microbe/class. The thickness of the chord represents the number of microbes participating in the interaction from the same class. The colours are mapped to microbial classes.

### Interactions of archaea in the Guaymas microbiome

*Candidatus* Pacearchaeota archaeon UWMA 0287 is an archaeon belonging to class Nanoarchaeota from the superphylum DPANN. Members of DPANN (including this class) are characterised by small genomes, and limited metabolic capabilities due to which they are predicted to rely on other microbes for most of their biosynthetic needs^41, 46–48^. It was also evident from the pairwise MSI analyses that Pacearchaeota are the largest beneficiary archaeon of Guaymas microbiome in GM media, JW1 media and all-media, while in marine broth 2216 Gemmatimonadetes bacterium UWMA 0339 benefited more. Though Pacearchaeota showed potential interactions with members of every other microbial class present in the Guaymas microbiome, most of the interactions were dominated by members of *Gammaproteobacteria, Poseidoniia, Alphaproteobacteria* and *Bacteroidia* (Figure 2). As the microbes receiving the greatest benefits from interactions in the community, Pacearchaeota receive cellobiose, O_2_, CO_2_, and H_2_S from its partners (Figure 5a). These exchanges were not seen in all four media, for example, the exchange of CO_2_ was restricted to GM media and marine broth 2216 alone as CO2 was already present in JW1 media and all-media. Among these, cellobiose can be seen in all interactions of Pacearchaeota except in marine broth 2216. Cellobiose is a disaccharide molecule and is a known carbon source for hyperthermophilic archaea^49^. Our models indicate that Pacearchaeota are able to accept cellobiose and hydrolyse it to use as a carbon source, thus leading to gain of many metabolic capabilities and high MSI in media except marine broth 2216. Pacearchaeota had the capability to donate metabolites like ornithine, putrescine, 4-aminobutanal (obtained during the metabolism of arginine) to other microbes only in all-media and JW1 media (Figure 5b). The metabolites exchanged in all other microbes are documented in **Supplementary File S4 and S5**.

**Figure 4.**
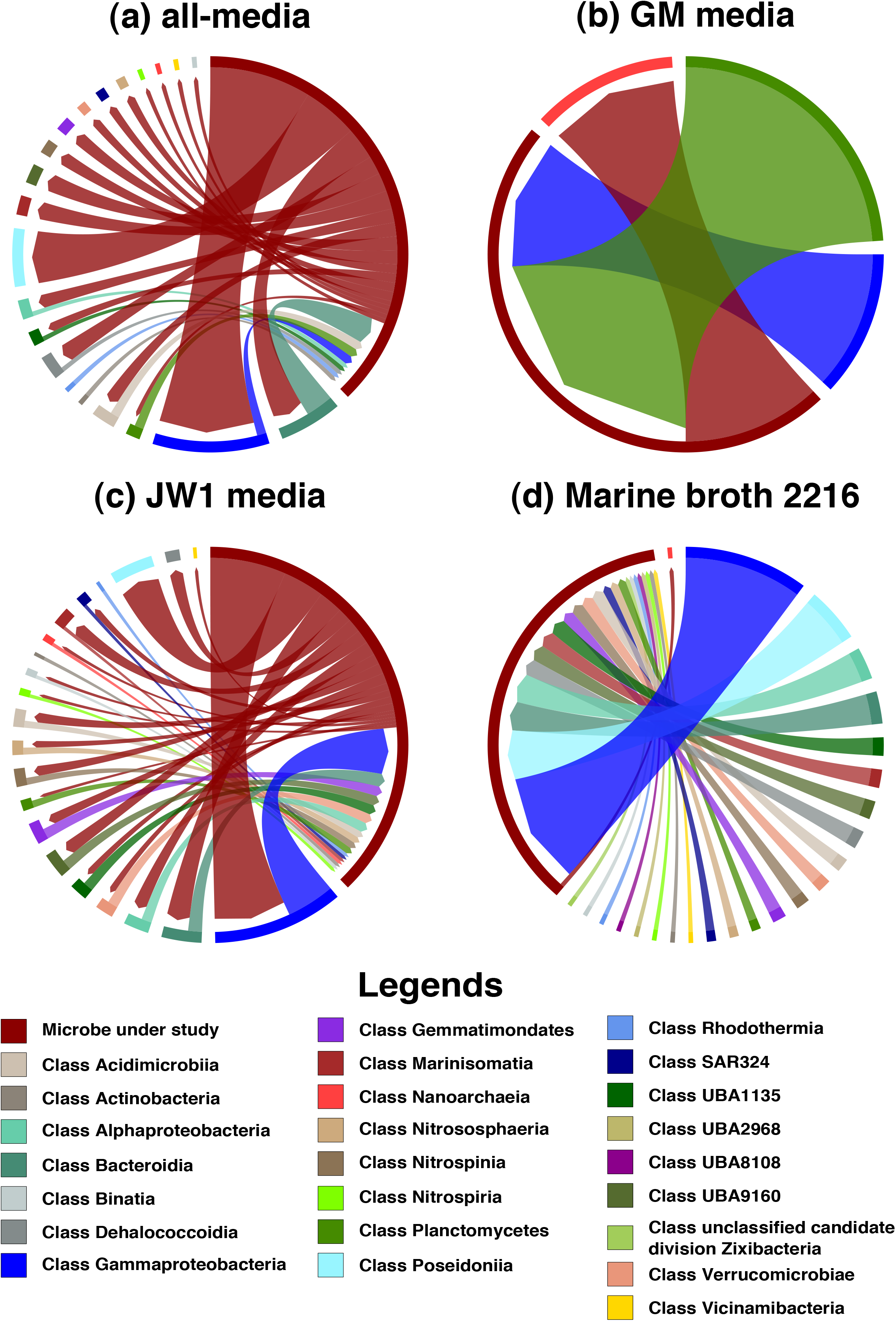
Pairwise MSI interactions of *Candidatus* Handelsmanbacteria bacterium UWMA 0286 with other microbes in the community. This chord diagram shows all possible metabolic interactions between *Candidatus* Handelsmanbacteria bacterium UWMA 0286 and other microbial classes present in the Guaymas microbiome in four different media conditions (a) all-media, (b) GM media, (c) JW1 media, (d) Marine Broth 2216. All the interacting microbes are grouped under their corresponding microbial class except *Candidatus* Handelsmanbacteria bacterium UWMA 0286. The chord starts from the donor microbe/class towards recipient microbe/class. The thickness of the chord represents the number of microbes participating in the interaction from the same class. The colours are mapped to microbial classes.

**Figure 5.**
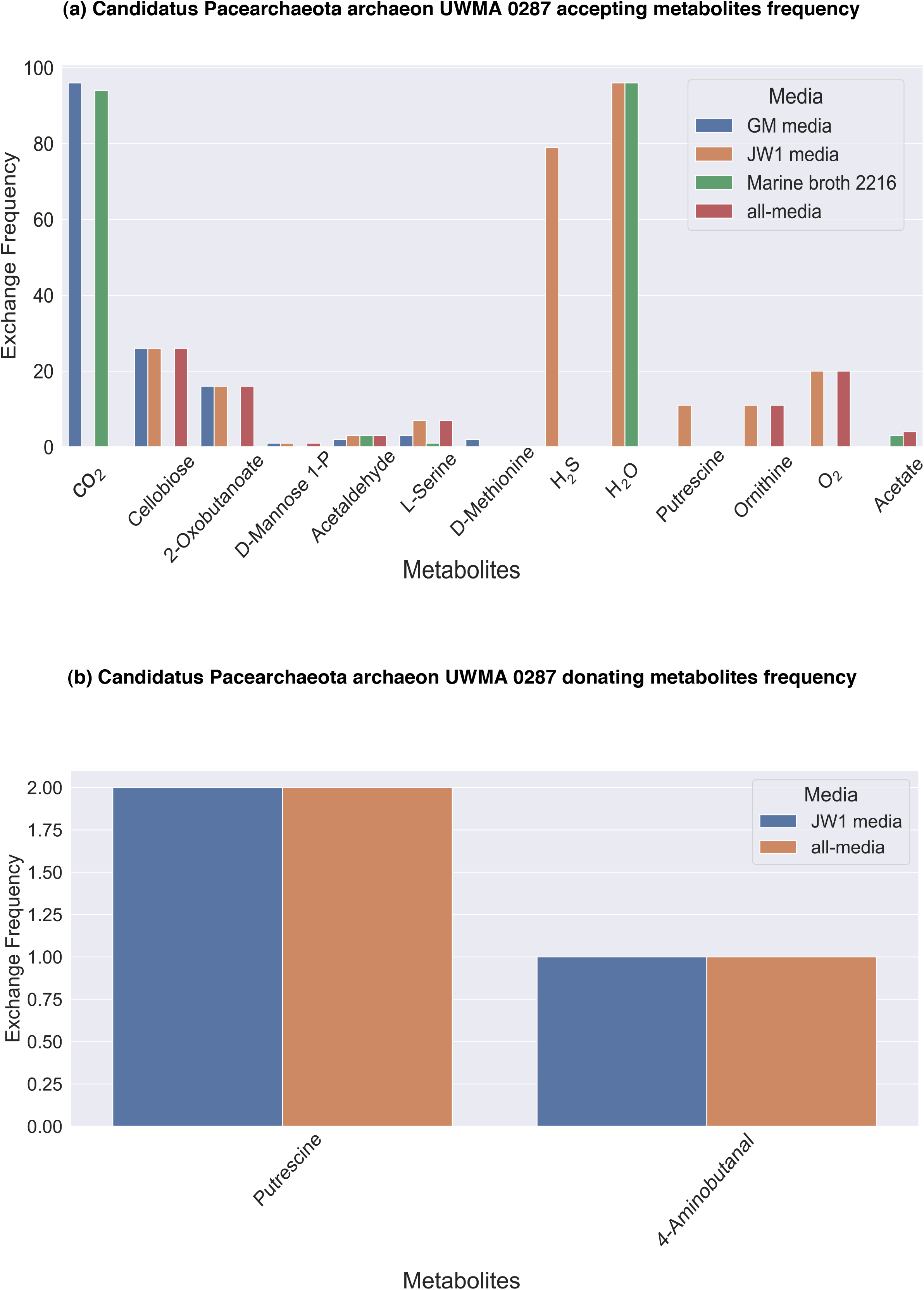
Metabolic support received and provided by *Candidatus* Pacearchaeota archaeon UWMA 0287. These plots depict the set of possible metabolites (a)accepted and (b)received by *Candidatus* Pacearchaeota archaeon UWMA 0287 from other microbes through the metabolic exchange. The Y-axis in this plot shows the number of interaction pairs in which that exchange has occurred, with 97 being the highest number of possible pairs for a microbe in a 98-member community.

### Interactions of bacteria in the Guaymas Basin microbiome

Bacteria in the Guaymas Basin microbiome exhibited a range of interactions from being able to interact with other classes of microbes to interacting with organisms from the same phylum/class. Members of the proteobacterial class *Gammaproteobacteria* are amongst the most abundant and dominant microbial populations in hydrothermal plumes^23^. In the Guaymas Basin microbiome, *Gammaproteobacteria* were predicted to have amongst the largest number of interactions.

*Candidatus* Lambdaproteobacteria bacterium UWMA 0318 (a member of the phylum SAR324) had the potential to interact with all other 23 microbial classes of the Guaymas Basin microbiome in JW1 media. *Candidatus* Lambdaproteobacteria bacterium UWMA 0298 showed the most interactions with microbes of the class *Gammaproteobacteria, Bacteroidia*, and *Alphaproteobacteria* in all the given media, except in GM media. The interactions were very minimal in GM media. *Candidatus* Lambdaproteobacteria bacterium UWMA 0298 acted as both donor and acceptor in all the four media. *Candidatus* Lambdaproteobacteria bacterium UWMA 0318 also showed interactions consistently with microbes of class *Gammaproteobacteria* in all the four media. Interactions with *Alphaproteobacteria* and *Bacteroidia* were seen in all-media and JW1 media while interactions with *Poseidonia* and *Marinisomatia* were prevalent in GM media. *Candidatus* Lambdaproteobacteria bacterium UWMA 0318 acted mostly as an acceptor in all the four media.

Meanwhile, *Candidatus* Handelsmanbacteria bacterium UWMA 0286 had the potential to interact with all other microbial classes as well as with the other member of its own class (*Candidatus* Handelsmanbacteria bacterium UWMA 0300) in marine broth 2216 (Figure 4 represents the interactions between *Candidatus* Handelsmanbacteria bacterium UWMA 0286 and other microbial classes in all four media). *Candidatus* Handelsmanbacteria bacteria showed most interactions with microbes of class *Gammaproteobacteria, Poseidonia, Bacteroidia*, and with *Alphaproteobacteria* in all the given media, except in GM media. The interactions were very minimal in GM media. In most cases *Candidatus* Handelsmanbacteria bacteria acts as a donor except in marine broth 2216 where all the interactions involved *Candidatus* Handelsmanbacteria bacteria were as receivers, though surprisingly the microbial class with which it interacted were the same in both cases.

Given the abundance of reduced sulfur species in hydrothermal plumes, we also identified sulfur oxidizing bacteria and observed their interactions. Bacteria from the SUP05 clade of *Gammaproteobacteria* (*Candidatus* Thioglobus) are amongst the most abundant and active members of plumes. In the Guaymas Basin microbiome, five different *Candidatus* Thioglobus members represented by *Candidatus* Thioglobus sp UWMA 0259, *Candidatus* Thioglobus sp UWMA 027, *Candidatus* Thioglobus sp UWMA 0322, *Candidatus* Thioglobus sp UWMA 0342, *Candidatus* Thioglobus sp UWMA 0360 interacted extensively with other organisms. First, *Candidatus* Thioglobus sp UWMA 0259 showed consistent interactions with other *Gammaproteobacteria* in all the four media, interactions with *Poseidonia* were observed in three media, except for marine broth 2216. *Candidatus* Thioglobus sp UWMA 0259 acts as acceptor in most cases. However in marine broth 2216, this bacterium acted as the donor but still maintained interactions with *Gammaproteobacteria, Alphaproteobacteria*, and *Bacterioidia* which were previously donating metabolites to *Candidatus* Thioglobus sp UWMA 0259 in other media. Second, *Candidatus* Thioglobus sp UWMA 0272 interacted with other *Gammaproteobacteria* and *Alphaproteobacteria* in three media except GM media where the only interaction was with *Nanoarchaeia* (*Candidatus* Pacearchaeota archaeon UWMA 0287). Third, *Candidatus* Thioglobus sp UWMA 0272 was observed as an acceptor in all-media while in other three media it acted as both an acceptor and donor. *Candidatus* Thioglobus sp UWMA 0322 showed most interactions with *Gammaproteobacteria* in all the four media, while interactions with *Poseidonia* were prevalent only in all-media and JW1 media. This microbe is a dominant a donor in all the four media conditions. Fourth, *Candidatus* Thioglobus sp UWMA 0342 interacted extensively with other *Gammaproteobacteria* in all four media, while interactions with *Bacterioidia* were seen in media except marine broth 2216 where the interactions were minimal. Interactions between *Alphaproteobacteria* and *Candidatus* Thioglobus sp UWMA 0342 were observed in only all-media and JW1 media. *Candidatus* Thioglobus sp UWMA 0342 was observed to act as both a donor and acceptor in all four media. Fifth, *Candidatus* Thioglobus sp UWMA 0360 interacted with other *Gammaproteobacteria* and *Alphaproteobacteria* in all-media and JW1 media. Interactions were minimal in the other two media, GM media and marine broth 2216. Interactions with *Nanoarchaeia* was observed in all four media. *Candidatus* Thioglobus sp UWMA 0360 was a dominant acceptor in all-media and JW1 media, while in marine broth 2216 it acted as a donor.

In addition to *Candidatus* Thioglobus, other abundant sulfur oxidizing bacteria in plumes were *Sulfitobacter* and *Thiotrichaceae* species. Sulfitobacter sp UWMA 0305 interacted predominantly with *Gammaproteobacteria* and *Bacterioidia* in media except GM media where the interactions were constrained to *Nanoarchaeia* and Planctomycetes. Interactions with *Poseidonia* were observed only in all-media and JW1 media. Sulfitobacter sp UWMA 0305 was a dominant donor except in marine broth 2216. Thiotrichaceae bacterium UWMA 0311 interacted predominantly with *Gammaproteobacteria* and *Poseidonia* in media except in marine broth 2216, while interactions with *Alphaproteobacteria* and *Bacteriodia* were common in media except GM media. Thiotrichaceae bacterium UWMA 0311 was observed to act as both an acceptor and donor in all four media.

### Key microbes in Guaymas Basin microbiome and unique contributors in the community

To determine the significance of microorganisms in a microbial community, we conducted CSI analyses (see Support offered by a group of microbes to the community in Materials and Methods) on the Guaymas Basin microbiome. First, the 98 microbes were clustered into 24 clusters based on the taxonomic class they belonged to. Secondly, each cluster was “knocked out” from the community to identify the metabolic capabilities lost by the community, a CSI value above zero indicates that the cluster has some significance to the community and an CSI score equal to zero indicates little to no significance to the community. This analysis was performed in all four media conditions and eight key microbial classes were identified based on the MSI scores (**Refer Supplementary Table 2** for the list of microbes in each media). These key microbial classes were *Alphaproteobacteria*, *Dehalococcoidia, Gammaproteobacteria, Nitrososphaeria*, Planctomycetes, *Poseidoniia, Rhodothermia*, and UBA8108. Only *Poseidoniia* were identified to be significant in all four media conditions.

Unique contributors in the community are microbes that have the capability to produce and donate certain metabolites that cannot be produced by any other microbe in the community. This was determined by performing CSI analysis where the metabolic capabilities of a community are studied before and after adding the microbe of interest (see Unique contributors of the community in Materials and Methods). Unique contributors were identified in Guaymas community in all four media conditions summing up to 10 different microbes (Refer Supplementary Table 3). Every one of these microbes is attributed to one or more unique metabolites that they can contribute to the community. The unique metabolites exchanged from these microbes to the community in different media conditions were agmantine, L-citrulline, L-ornithine, trans-4-hydroxy-L-proline, 2-oxoglutarate (Figure 6), which were mostly metabolites involved in amino acid synthesis or metabolism pathways. This shows that amino acid auxotrophies exist in the community and are an important driver of the exchange of these metabolites from producers to the auxotrophs. This is potentially explained by the abundance of DPANN archaea in the community which are known to be auxotrophic for amino acids^50^. In addition to these metabolites, dihydroxyacetone, dihydroxyacetone phosphate, and acetone were also exchanged by these contributors to the community.

**Figure 6.**
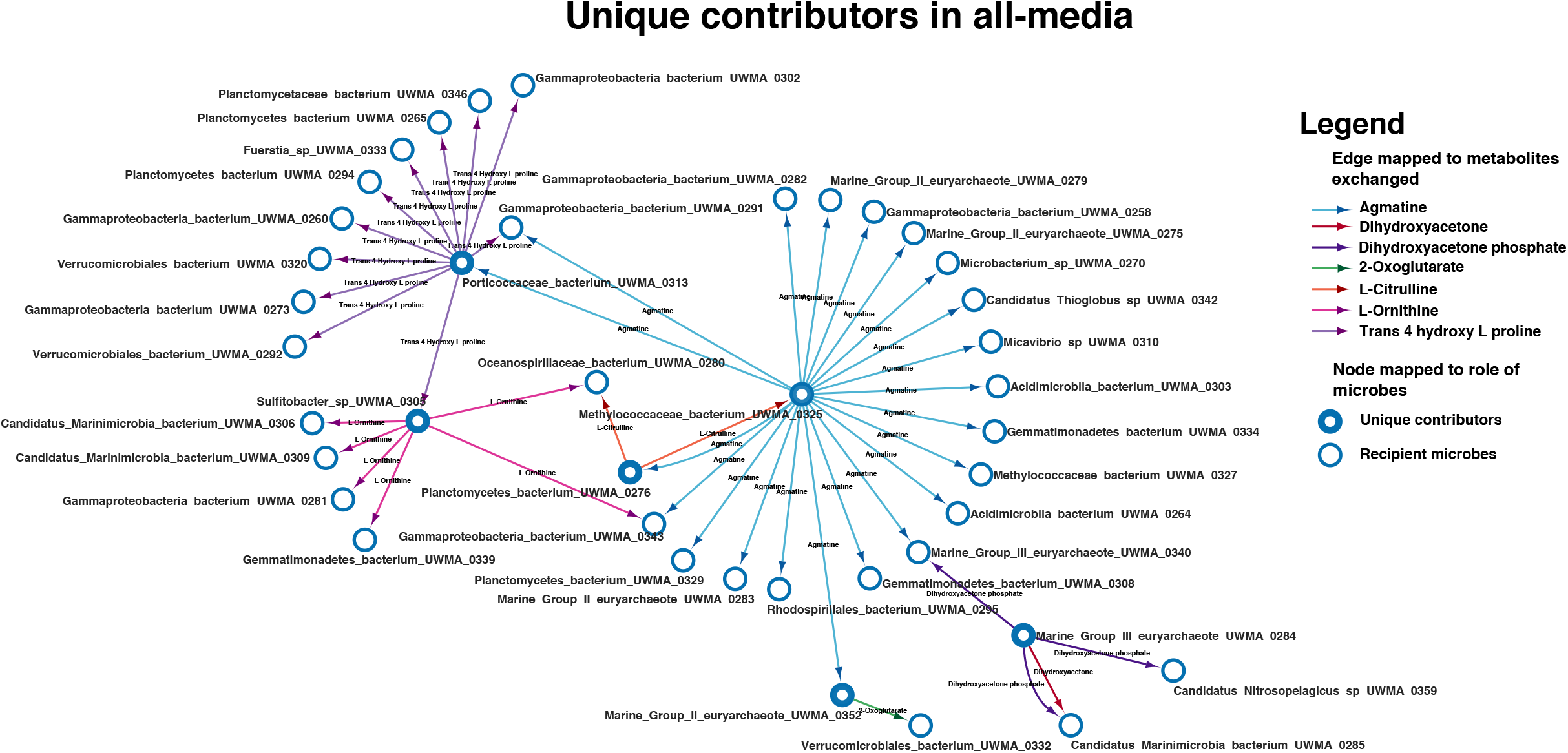
Unique microbial contributors in all-media. This network highlights the unique contributors to the Guaymas microbiome in all-media. Each node represents a microbe, and the arrows start from donating the microbe to the recipient microbe. The colour of the arrows is mapped to different unique metabolites exchanged by the unique contributors.

### Resource competition in the Guaymas Basin microbial community

To study metabolic resource competition in the community, we employed a metric called Metabolic Resource Overlap (MRO)^37, 51^. Briefly, MRO is the maximum possible overlap of the minimal metabolite set of all members of the community required for their growth. MRO is solely dependent on the metabolism of the microbes and hence the lesser the MRO, more complementary the microbial metabolisms to each other in the community. In this study, we have computed MRO for different communities including an anaerobic digestion microbiome (ADM)^38^, gut microbiome^39^, East Pacific Rise L hydrothermal vent microbiome, East Pacific Rise M hydrothermal vent microbiome, and Guaymas Basin hydrothermal plume microbiome. In each microbiome dataset, MRO was observed for community size ranging from 2 (pairwise community) to 10 (10-member community) (Refer Studying the level of competition in the community in Materials and Methods). On comparing the MRO values of diverse microbial communities pertaining to different metabolic niches, we observed the the MRO of ADM and gut microbiomes which belong to relatively similar niches were relatively close while that of hydrothermal vent microbiomes were significantly lower than that of former (Figure 7). Overall, these MRO values agree with our findings from the MSI analyses since lesser the overlap in metabolism, the higher the potential for interaction between microbes.

**Figure 7.**
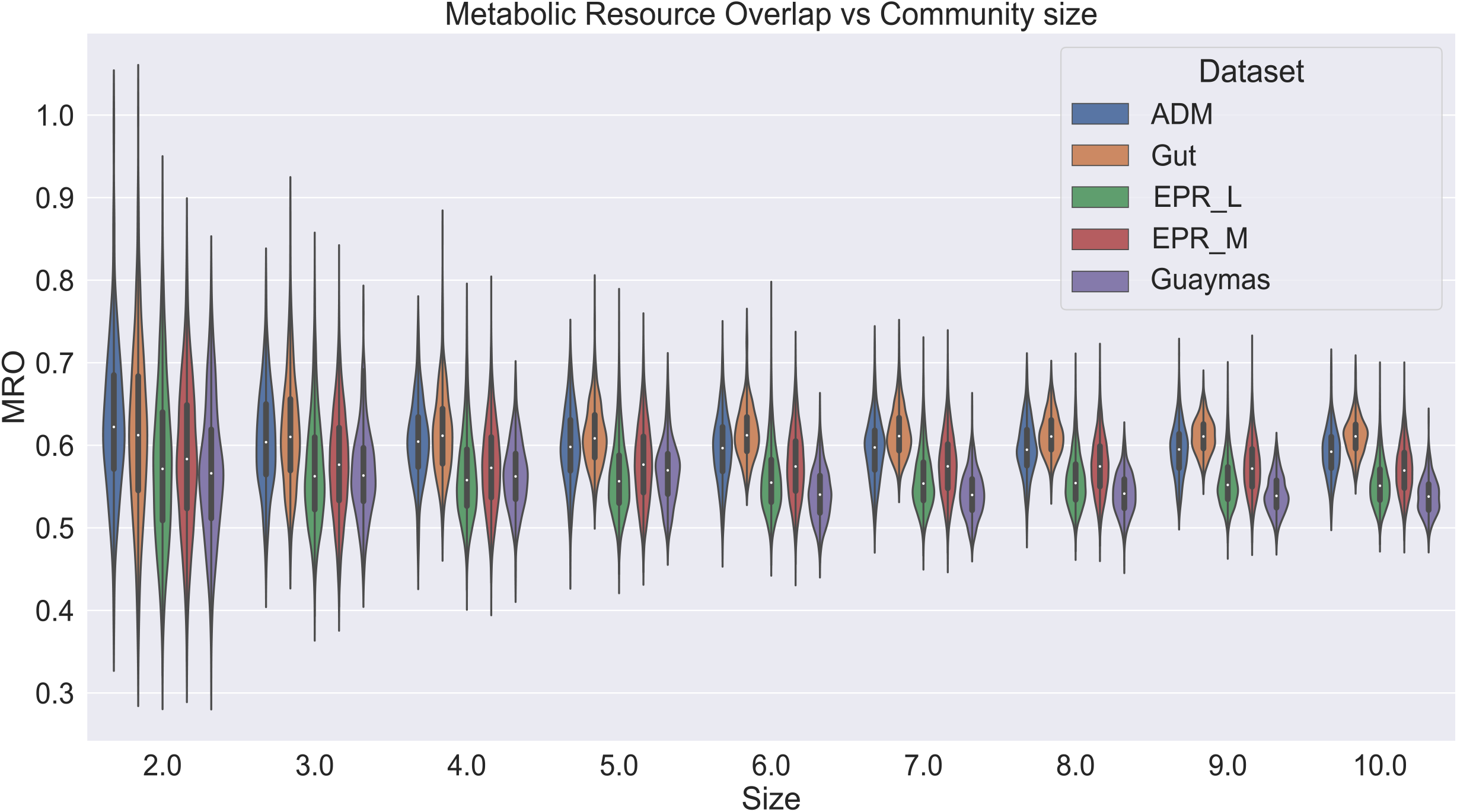
Metabolic Resource Overlap scores of different microbiomes compared to the Guaymas microbiome. This violin plot represents the distribution of the MRO score of four different microbial communities. 1. Anaerobic Digestion Microbiome (ADM) (Blue) 2. Gut mucribiome (Orange) 3. East Pacific Rise (EPR) L hydrothermal vent (active vent) microbiome (Green) 4. East Pacific Rise (EPR) M hydrothermal vent (inactive vent) microbiome (Red) 5. Guaymas microbiome (Violet). The MRO scores (Y-axis) are determined for different community sizes (X-axis), from a 2-member community to a 10-member community.

### Horizontal Gene Transfers (HGTs) in the Guaymas Basin microbiome

HGT is one of the survival strategies adopted by microbes to compete in challenging ecosystems^52^. During this process, microbes acquire novel DNA from their partners or from the environment and evolve their metabolic capabilities^53, 54^. Microbes coexisting as communities undergo HGT events to enforce cooperation and hence also helpful in structuring the communities^55^. Therefore, we studied HGT events in the community using a tool called MetaCHIP, which allowed for detecting HGT events in our metagenomic data. A list of 214 HGT events was detected in the community (Figure 8(a)). On functional annotation, we observed that most of the HGT genes were responsible for translation machinery, energy production and conversion, amino acid metabolism, and transport mechanisms. The gene transfers occurred across genera and species, but there were no specific patterns observed at that level. However, zooming out to the level of classes, HGTs were more frequently observed between *Gammaproteobacteria* and *Alphaproteobacteria* (Figure 8(b))

**Figure 8(a).**
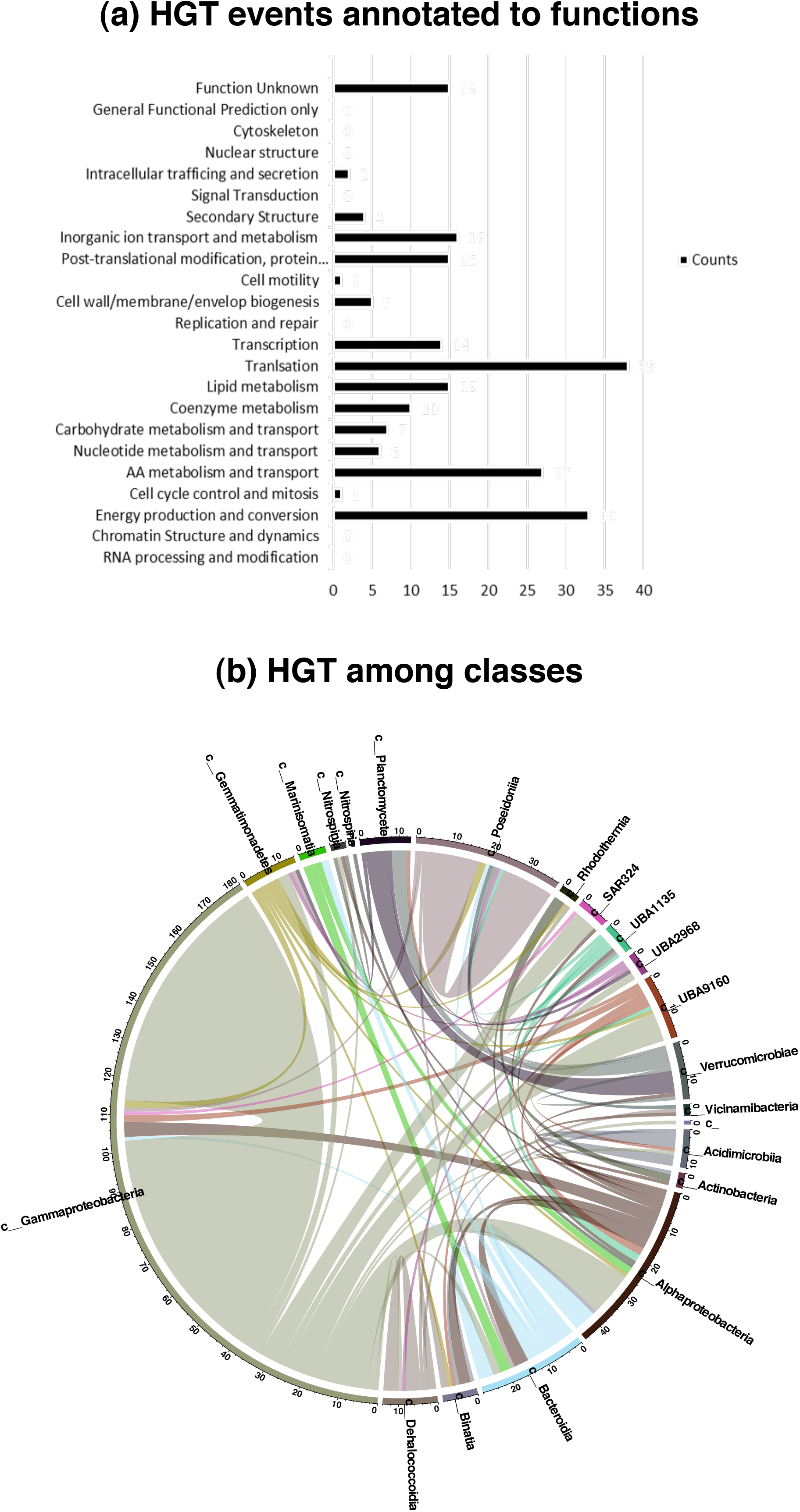
Horizontal gene transfer events annotated to functions. This figure represents the number of times a gene responsible for a particular function underwent horizontal transfer. **Figure 8(b) Horizontal gene transfer among microbial classes.** This figure traces the microbial classes participating in HGT events.

## Discussion

In this study, we explored the use of systems-level modelling of hydrothermal plume microbial communities in the Guaymas basin. Microbes in these extreme habitats are unique in different ways extending our knowledge of the diversity of life on earth. These microbes are adapted to chemoautotrophy due to their scarce exposure to sunlight. With metagenomic data corresponding to 98 microbes from Guaymas Basin hydrothermal plumes, genome-scale metabolic models were built using CarveMe. The main focus of our research was to shed light on the possible interactions that can be observed in these complex deep-sea microbiomes. Insights were obtained for metabolic interactions in the community by studying the metabolic exchanges and genetic interactions in the community by studying HGTs.

The major focus of our study was unveiling the possible interactions between archaea and bacteria. One of the interesting predictions was about archaeon *Candidatus* Pacearchaeota archaeon UWMA 0287 belonging to class Nanoarchaeia. This archaeon was one of the most dependent microbes in Guaymas microbiome (remains an acceptor in all high MSI pairwise interactions). We hypothesize that this observation is likely due to the microbes of class nanoarchaeota being devoid of core metabolic pathways as reported by^26, 46–48, 56^ and hence might lead a parasitic or symbiotic lifestyle.

At the same time, not all archaea in Guaymas microbiome are metabolically dependent on another microbe. Archaea of class *Poseidoniia* (Phylum Euryarchaeota) can act as supporters to bacteria and other archaea (**Refer Supplementary File S6**). This is likely because these microbes have greater metabolic capabilities in comparison to other microbes in the Guaymas microbiome. This can be due to the metabolic capabilities acquired through HGT events^57^. HGT analysis showed that *Poseidoniia* did take part in HGTs, and most of the genes transferred were related to metabolism. This is potential cause for *Poseidoniia* becoming dominant archaea in the Guaymas microbiome.

Microbes of class *Gammaproteobacteria* form the majority of the Guaymas microbiome, which might be due to their ability to interact with most microbes in the community. In most cases, *Gammaproteobacteria* act as donors due to the large metabolic capability of these microbes in the hydrothermal plume community. Results from genome-scale anlayses of MAGs generated using METABOLIC^9^ also confirmed that the metabolic contributions made by *Gammaproteobacteria* were the highest among the Guaymas community. *Gammaproteobacteria* are also recognised for their contributions towards nitrogen fixation, ammonia oxidation, and denitrification in hydrothermal vent ecosystems^4, 58, 59^.

Metabolic modelling showed a majority of microbial activity involves the exchange of oxygen, amino acids like serine, malate, methionine, amino acid intermediates like 4-Aminobutanoate, indole-3-acetaldehyde and elements involved in the carbon cycle like CO_2_, acetaldehyde, sulfur-based compounds like methanethiol, H_2_S. This might suggest that metabolites like CO_2_, H_2_ and H_2_S are important to the microbes in this environment. Microbes in hydrothermal vent ecosystems rely on the oxidation of sulfur, and sulfur-based reduced compounds, and hydrogen oxidation for energy metabolism^60, 61^. Thus, these metabolites are likley to play major roles in this ecosystem. It was also observed that the absence of these metabolites in media was always compensated by exchange from other microbes. The absence of CO_2_ in GM media and Marine broth 2216 and the absence of H_2_S in JW1 media were all compensated by metabolic exchanges.

The unique metabolites predicted to be exchanged in the community in different media conditions, were agmantine, L-citrulline, L-ornithine, trans-4-hydroxy-L-proline, and 2-oxoglutarate, which were mostly the metabolites involved in amino acid synthesis or metabolism pathways. This likely implies that amino acid auxotrophies exist in the community and drives the exchange of these metabolites from producers to auxotrophs. The extensive occurrence of metabolic handoffs in hydrothermal plume communities provides functional interdependency between microbes, leading to auxotrophies. Thus, the community achieves efficient energy and substrate transformations^62^.

In summary, this research focused on unveiling the possible interactions between archaea and bacteria in the Guaymas hydrothermal plume microbiome by constructing metabolic networks of corresponding microbes. This approach allowed us to predict possible metabolic exchanges between individual microbes, and the metabolic capabilities of microbes in different media conditions, which are indecipherable to this extent by experimental approaches. The approaches described herein have led to many interesting hypotheses, providing a fertile ground for future wet lab experiments to further understand the organisation of the Guaymas hydrothermal plume microbiome, and deep-sea microbiomes broadly, to gain better insights into the cultivation of uncultivated organisms in consortia. Studying higher-order interactions of microbes in this community has highlighted unique metabolic contributors amongst microbes in the community. While metabolic modelling provides insights into metabolic interactions, HGT analysis helped explore gene transfers between microbes in the community. Overall, the approach here is fairly generic and can be applied to any microbial community to generate testable hypotheses on experimentally unculturable microbes.

## Code availability

All code used in this study is publicly available from GitHub.

## Data availability

All genomes (MAGs) used in this study are publicly available through NCBI BioProject PRJNA522654. Metagenomic reads are avaialble through NCBI SRA accession number SRR3577362.

## Acknowledgements

DKKB acknowledges the HTRA fellowship from the Ministry of Education, Government of India. SU acknowledges the IBSE post-baccalaureate fellowship. KR acknowledges support from the Science and Engineering Board (SERB) MATRICS Grant MTR/2020/000490. KA acknowledges support from the US National Science Foundation under grant number OCE 2049478.

## References

1. Dick, G. J. The microbiomes of deep-sea hydrothermal vents: distributed globally, shaped locally. Nat. Rev. Microbiol. 17, 271–283, 10.1038/s41579-019-0160-2 (2019).

2. Dick, G. et al. The microbiology of deep-sea hydrothermal vent plumes: ecological and biogeographic linkages to seafloor and water column habitats. Front. Microbiol. 4, 124, 10.3389/fmicb.2013.00124 (2013).

3. Lesniewski, R. A., Jain, S., Anantharaman, K., Schloss, P. D. & Dick, G. J. The metatranscriptome of a deep-sea hydrothermal plume is dominated by water column methanotrophs and lithotrophs. The ISME J. 6, 2257–2268, 10.1038/ismej.2012.63 (2012).

4. Baker, B. J. et al. Community transcriptomic assembly reveals microbes that contribute to deep-sea carbon and nitrogen cycling. The ISME J. 7, 1962–1973, 10.1038/ismej.2013.85 (2013). Number: 10 Publisher: Nature Publishing Group.

5. Abreu, N. A. & Taga, M. E. Decoding molecular interactions in microbial communities. FEMS Microbiol. Rev. 40, 648–663, 10.1093/femsre/fuw019 (2016).

6. Bosse, M. et al. Interaction networks for identifying coupled molecular processes in microbial communities. BioData Min. 8, 21, 10.1186/s13040-015-0054-4 (2015).

7. Morris, B. E., Henneberger, R., Huber, H. & Moissl-Eichinger, C. Microbial syntrophy: interaction for the common good. FEMS Microbiol. Rev. 37, 384–406, 10.1111/1574-6976.12019 (2013).

8. Wankel, S. D. et al. Influence of subsurface biosphere on geochemical fluxes from diffuse hydrothermal fluids. Nat. Geosci. 4, 461–468, 10.1038/ngeo1183 (2011).

9. Zhou, Z. et al. METABOLIC: high-throughput profiling of microbial genomes for functional traits, metabolism, biogeochemistry, and community-scale functional networks. Microbiome 10, 33, 10.1186/s40168-021-01213-8 (2022).

10. Xu, H. S. et al. Survival and viability of nonculturableEscherichia coli andVibrio cholerae in the estuarine and marine environment. Microb. Ecol. 8, 313–323, 10.1007/BF02010671 (1982).

11. Shoaie, S. et al. Understanding the interactions between bacteria in the human gut through metabolic modeling. Sci. Reports 3, 2532, 10.1038/srep02532 (2013).

12. Thommes, M., Wang, T., Zhao, Q., Paschalidis, I. C. & Segrè, D. Designing Metabolic Division of Labor in Microbial Communities. mSystems 4, e00263–18, 10.1128/mSystems.00263-18 (2019).

13. Ponomarova, O. & Patil, K. R. Metabolic interactions in microbial communities: Untangling the Gordian knot. Curr. Opin. Microbiol. 27, 37–44, 10.1016/j.mib.2015.06.014 (2015).

14. Ang, K. S., Lakshmanan, M., Lee, N.-R. & Lee, D.-Y. Metabolic Modeling of Microbial Community Interactions for Health, Environmental and Biotechnological Applications. Curr. Genomics 19, 712–722, 10.2174/1389202919666180911144055 (2018).

15. Blasche, S., Kim, Y., Oliveira, A. P. & Patil, K. R. Model microbial communities for ecosystems biology. Curr. Opin. Syst. Biol. 6, 51–57, 10.1016/j.coisb.2017.09.002 (2017).

16. McCloskey, D., Palsson, B. Ø. & Feist, A. M. Basic and applied uses of genome-scale metabolic network reconstructions of Escherichia coli. Mol. Syst. Biol. 9, 661, 10.1038/msb.2013.18 (2013).

17. Ravikrishnan, A. & Raman, K. Systems-Level Modelling of Microbial Communities: Theory and Practice (CRC Press, Boca Raton, 2018).

18. Kumar, M., Ji, B., Zengler, K. & Nielsen, J. Modelling approaches for studying the microbiome. Nat. Microbiol. 4, 1253–1267, 10.1038/s41564-019-0491-9 (2019).

19. Ibrahim, M., Raajaraam, L. & Raman, K. Modelling microbial communities: Harnessing consortia for biotechnological applications. Comput. Struct. Biotechnol. J. 19, 3892–3907, 10.1016/j.csbj.2021.06.048 (2021).

20. Gu, C., Kim, G. B., Kim, W. J., Kim, H. U. & Lee, S. Y. Current status and applications of genome-scale metabolic models. Genome Biol. 20, 121, 10.1186/s13059-019-1730-3 (2019).

21. Machado, D., Andrejev, S., Tramontano, M. & Patil, K. R. Fast automated reconstruction of genome-scale metabolic models for microbial species and communities. Nucleic Acids Res. 46, 7542–7553, 10.1093/nar/gky537 (2018).

22. Thiele, I. & Palsson, B. Ø. A protocol for generating a high-quality genome-scale metabolic reconstruction. Nat. Protoc. 5, 93–121, 10.1038/nprot.2009.203 (2010).

23. Zhou, Z., Tran, P. Q., Kieft, K. & Anantharaman, K. Genome diversification in globally distributed novel marine Proteobacteria is linked to environmental adaptation. The ISME J. 14, 2060–2077, 10.1038/s41396-020-0669-4 (2020).

24. Bowers, R. M. et al. Minimum information about a single amplified genome (MISAG) and a metagenome-assembled genome (MIMAG) of bacteria and archaea. Nat. Biotechnol. 35, 725–731, 10.1038/nbt.3893 (2017). Number: 8 Publisher: Nature Publishing Group.

25. Anantharaman, K., Breier, J. A., Sheik, C. S. & Dick, G. J. Evidence for hydrogen oxidation and metabolic plasticity in widespread deep-sea sulfur-oxidizing bacteria. Proc. Natl. Acad. Sci. 110, 330–335, 10.1073/pnas.1215340110 (2013).

26. Hou, J. et al. Microbial succession during the transition from active to inactive stages of deep-sea hydrothermal vent sulfide chimneys. Microbiome 8, 102, 10.1186/s40168-020-00851-8 (2020).

27. Takai, K. et al. Isolation and phylogenetic diversity of members of previously uncultivated-Proteobacteria in deep-sea hydrothermal fields. FEMS Microbiol. Lett. 218, 167–174, 10.1111/j.1574-6968.2003.tb11514.x (2003).

28. Henson, M. W. et al. Artificial Seawater Media Facilitate Cultivating Members of the Microbial Majority from the Gulf of Mexico. mSphere 1, e00028–16, 10.1128/mSphere.00028-16 (2016).

29. Giovannoni, S. & Stingl, U. The importance of culturing bacterioplankton in the ‘omics’ age. Nat. Rev. Microbiol. 5, 820–826, 10.1038/nrmicro1752 (2007). Number: 10 Publisher: Nature Publishing Group.

30. Ravikrishnan, A., Nasre, M. & Raman, K. Enumerating all possible biosynthetic pathways in metabolic networks. Sci. Reports 8, 9932, 10.1038/s41598-018-28007-7 (2018).

31. Ravikrishnan, A., Blank, L. M., Srivastava, S. & Raman, K. Investigating metabolic interactions in a microbial co-culture through integrated modelling and experiments. Comput. Struct. Biotechnol. J. 18, 1249–1258, 10.1016/j.csbj.2020.03.019 (2020).

32. Shannon, P. et al. Cytoscape: A Software Environment for Integrated Models of Biomolecular Interaction Networks. Genome Res. 13, 2498–2504, 10.1101/gr.1239303 (2003).

33. Kumar, R. K. et al. Metabolic modeling of the International Space Station microbiome reveals key microbial interactions. Microbiome 10, In Press (2022).

34. Song, W., Wemheuer, B., Zhang, S., Steensen, K. & Thomas, T. MetaCHIP: community-level horizontal gene transfer identification through the combination of best-match and phylogenetic approaches. Microbiome 7, 36, 10.1186/s40168-019-0649-y (2019).

35. Bansal, M. S., Kellis, M., Kordi, M. & Kundu, S. RANGER-DTL 2.0: Rigorous reconstruction of gene-family evolution by duplication, transfer and loss. Bioinforma. (Oxford, England) 34, 3214–3216, 10.1093/bioinformatics/bty314 (2018).

36. Huerta-Cepas, J. et al. Fast Genome-Wide Functional Annotation through Orthology Assignment by eggNOG-Mapper. Mol. Biol. Evol. 34, 2115–2122, 10.1093/molbev/msx148 (2017).

37. Zelezniak, A. et al. Metabolic dependencies drive species co-occurrence in diverse microbial communities. Proc. Natl. Acad. Sci. 112, 6449–6454, 10.1073/pnas.1421834112 (2015).

38. Basile, A. et al. Revealing metabolic mechanisms of interaction in the anaerobic digestion microbiome by flux balance analysis. Metab. Eng. 62, 138–149, 10.1016/j.ymben.2020.08.013 (2020).

39. Magnúsdóttir, S. et al. Generation of genome-scale metabolic reconstructions for 773 members of the human gut microbiota. Nat. Biotechnol. 35, 81–89, 10.1038/nbt.3703 (2017).

40. Rinke, C. et al. Insights into the phylogeny and coding potential of microbial dark matter. Nature 499, 431–437, 10.1038/nature12352 (2013).

41. Castelle, C. J. et al. Genomic Expansion of Domain Archaea Highlights Roles for Organisms from New Phyla in Anaerobic Carbon Cycling. Curr. Biol. 25, 690–701, 10.1016/j.cub.2015.01.014 (2015).

42. Ortiz-Alvarez, R. & Casamayor, E. O. High occurrence of Pacearchaeota and Woesearchaeota (Archaea superphylum DPANN) in the surface waters of oligotrophic high-altitude lakes. Environ. Microbiol. Reports 8, 210–217, 10.1111/1758-2229.12370 (2016).

43. Karner, M. B., DeLong, E. F. & Karl, D. M. Archaeal dominance in the mesopelagic zone of the Pacific Ocean. Nature 409, 507–510, 10.1038/35054051 (2001). Number: 6819 Publisher: Nature Publishing Group.

44. Francis, C. A., Roberts, K. J., Beman, J. M., Santoro, A. E. & Oakley, B. B. Ubiquity and diversity of ammonia-oxidizing archaea in water columns and sediments of the ocean. Proc. Natl. Acad. Sci. 102, 14683–14688, 10.1073/pnas.0506625102 (2005). Publisher: Proceedings of the National Academy of Sciences.

45. Baker, B. J., Lesniewski, R. A. & Dick, G. J. Genome-enabled transcriptomics reveals archaeal populations that drive nitrification in a deep-sea hydrothermal plume. The ISME J. 6, 2269–2279, 10.1038/ismej.2012.64 (2012). Number: 12 Publisher: Nature Publishing Group.

46. Casanueva, A. et al. Nanoarchaeal 16S rRNA gene sequences are widely dispersed in hyperthermophilic and mesophilic halophilic environments. Extremophiles 12, 651–656, 10.1007/s00792-008-0170-x (2008).

47. Wurch, L. et al. Genomics-informed isolation and characterization of a symbiotic Nanoarchaeota system from a terrestrial geothermal environment. Nat. Commun. 7, 12115, 10.1038/ncomms12115 (2016).

48. St. John, E. et al. A new symbiotic nanoarchaeote (Candidatus Nanoclepta minutus) and its host (Zestosphaera tikiterensis gen. nov., sp. nov.) from a New Zealand hot spring. Syst. Appl. Microbiol. 42, 94–106, 10.1016/j.syapm.2018.08.005 (2019).

49. Koning, S. M., Elferink, M. G. L., Konings, W. N. & Driessen, A. J. M. Cellobiose Uptake in the Hyperthermophilic Archaeon Pyrococcus furiosus Is Mediated by an Inducible, High-Affinity ABC Transporter (2001).

50. Sakai, H. D. et al. Insight into the symbiotic lifestyle of DPANN archaea revealed by cultivation and genome analyses. Proc. Natl. Acad. Sci. 119, e2115449119, 10.1073/pnas.2115449119 (2022). Publisher: Proceedings of the National Academy of Sciences.

51. Machado, D. et al. Polarization of microbial communities between competitive and cooperative metabolism. bioRxiv 2020.01.28.922583, 10.1101/2020.01.28.922583 (2020).

52. Brito, I. L. Examining horizontal gene transfer in microbial communities. Nat. Rev. Microbiol. 19, 442–453, 10.1038/s41579-021-00534-7 (2021).

53. Hehemann, J.-H. et al. Transfer of carbohydrate-active enzymes from marine bacteria to Japanese gut microbiota. Nature 464, 908–912, 10.1038/nature08937 (2010).

54. Jain, R., Rivera, M. C., Moore, J. E. & Lake, J. A. Horizontal gene transfer in microbial genome evolution. Theor. Popul. Biol. 61, 489–495, 10.1006/tpbi.2002.1596 (2002).

55. Polz, M. F., Alm, E. J. & Hanage, W. P. Horizontal gene transfer and the evolution of bacterial and archaeal population structure. Trends Genet. 29, 170–175, 10.1016/j.tig.2012.12.006 (2013).

56. McCliment, E. A. et al. Colonization of nascent, deep-sea hydrothermal vents by a novel Archaeal and Nanoarchaeal assemblage. Environ. Microbiol. 8, 114–125, 10.1111/j.1462-2920.2005.00874.x (2006). _eprint: https://onlinelibrary.wiley.com/doi/pdf/10.1111/j.1462-2920.2005.00874.x.

57. Deschamps, P., Zivanovic, Y., Moreira, D., Rodriguez-Valera, F. & López-García, P. Pangenome Evidence for Extensive Interdomain Horizontal Transfer Affecting Lineage Core and Shell Genes in Uncultured Planktonic Thaumarchaeota and Euryarchaeota. Genome Biol. Evol. 6, 1549–1563, 10.1093/gbe/evu127 (2014).

58. Li, J. et al. Oxidative Weathering and Microbial Diversity of an Inactive Seafloor Hydrothermal Sulfide Chimney. Front. Microbiol. 8 (2017).

59. Adam, N. & Perner, M. Microbially Mediated Hydrogen Cycling in Deep-Sea Hydrothermal Vents. Front. Microbiol. 9 (2018).

60. Zhou, Z. et al. Gammaproteobacteria mediating utilization of methyl-, sulfur- and petroleum organic compounds in deep ocean hydrothermal plumes. The ISMEJ. 14, 3136–3148, 10.1038/s41396-020-00745-5 (2020). Number: 12 Publisher: Nature Publishing Group.

61. Zierenberg, R. A., Adams, M. W. W. & Arp, A. J. Life in extreme environments: Hydrothermal vents. Proc. Natl. Acad. Sci. 97, 12961–12962, 10.1073/pnas.210395997 (2000).

62. Zhou, Z., John, E. S., Anantharaman, K. & Reysenbach, A.-L. Global patterns of diversity and metabolism of microbial communities in deep-sea hydrothermal vent deposits, 10.1101/2022.08.08.503129 (2022). Pages: 2022.08.08.503129 Section: New Results.

