## Supplementary Tables 1-3 for "Metagenome-based metabolic modelling predicts unique microbial interactions in deep-sea hydrothermal plume microbiomes"


| | |
|---|---|
| Table 1 | List of archaea present in the Guaymas microbiome |
| Table 2 | List of unique contributing microbial classes in Guaymas microbiome across media |
| Table 3 | List of unique contributors in Guaymas microbiome across media |


**Supplementary Table 1.** List of archaea present in the Guaymas microbiome.

| S. No | Archaea |
|---|---|
| 1 | Candidatus Nitrosopelagicus sp UWMA 0359 |
| 2 | Candidatus Pacearchaeota archaeon UWMA 0287 |
| 3 | Marine Group II euryarchaeote UWMA 0266 |
| 4 | Marine Group II euryarchaeote UWMA 0275 |
| 5 | Marine Group II euryarchaeote UWMA 0279 |
| 6 | Marine Group II euryarchaeote UWMA 0283 |
| 7 | Marine Group II euryarchaeote UWMA 0323 |
| 8 | Marine Group II euryarchaeote UWMA 0328 |
| 9 | Marine Group II euryarchaeote UWMA 0344 |
| 10 | Marine Group II euryarchaeote UWMA 0350 |
| 11 | Marine Group II euryarchaeote UWMA 0352 |
| 12 | Marine Group II euryarchaeote UWMA 0357 |
| 13 | Marine Group III euryarchaeote UWMA 0284 |
| 14 | Marine Group III euryarchaeote UWMA 0340 |
| 15 | Nitrosopumilus sp UWMA 0263 |

**Supplementary Table 2.** List of unique contributing microbial classes in Guaymas microbiome across media.

| S. No | allmedia | GM media | JW1 media | Marine broth 2216 |
|---|---|---|---|---|
| 1 | Alphaproteobacteria | | Alphaproteobacteria | |
| 2 | | | Dehalococcoidia | |
| 3 | Gammaproteobacteria | | Gammaproteobacteria | Gammaproteobacteria |
| 4 | | | | Nitrososphaeria |
| 5 | | Planctomycetes | | |
| 6 | Poseidoniia | Poseidoniia | Poseidoniia | Poseidoniia |
| 7 | | | | Rhodothermia |
| 8 | UBA8108 | | UBA8108 | |

**Supplementary Table 3.** List of unique contributors in Guaymas microbiome across media.

| S.No | allmedia | GM media | JW1 media | Marine broth 2216 |
|---|---|---|---|---|
| 1 | Planctomycetes bacterium UWMA 0276 | | Planctomycetes bacterium UWMA 0276 | |
| 2 | Porticoccaceae bacterium UWMA 0313 | | Porticoccaceae bacterium UWMA 0313 | |
| 3 | Sulfitobacter sp UWMA 0305 | | Sulfitobacter sp UWMA 0305 | |
| 4 | Methylococcaceae bacterium UWMA 0325 | | Methylococcaceae bacterium UWMA 0325 | |
| 5 | Marine Group II euryarchaeote UWMA 0352 | | Marine Group II euryarchaeote UWMA 0352 | Marine Group II euryarchaeote UWMA 0352 |
| 6 | Marine Group III euryarchaeote UWMA 0284 | Marine Group III euryarchaeote UWMA 0284 | Marine Group III euryarchaeote UWMA 0284 | |
| 7 | | | Dehalococcoidia bacterium UWMA 0267 | |
| 8 | | | | Nitrosopumilus sp UWMA 0263 |
| 9 | | | | Bacteroidetes bacterium UWMA 0293 |
| 10 | | | | Gammaproteobacteria bacterium UWMA 0299 |
