## Supplementary File S2 for "Metagenome-based metabolic modelling predicts unique microbial interactions in deep-sea hydrothermal plume microbiomes": Supplementary Figure S0 legend.pdf

### Legend for pairwise interaction plots

- Microbe under study
- Class Acidimicrobiia
- Class Actinobacteria
- Class Alphaproteobacteria
- Class Bacteroidia
- Class Binatia
- Class Dehalococcoidia
- Class Gammaproteobacteria
- Class Gemmatimondates
- Class Marinisomatia
- Class Nanoarchaeia
- Class Nitrososphaeria
- Class Nitrospinia
- Class Nitrospiria
- Class Planctomycetes
- Class Poseidoniia
- Class Rhodothermia
- Class SAR324
- Class UBA1135
- Class UBA2968
- Class UBA8108
- Class UBA9160
- Class unclassified candidate division Zixibacteria
- Class Verrucomicrobiae
- Class Vicinamibacteria
