## Supplementary figures and images for "Metagenome-based metabolic modelling predicts unique microbial interactions in deep-sea hydrothermal plume microbiomes"

### Supplementary Figure S1 Acidimicrobiia bacterium UWMA 0264 accepting metabolites.pdf

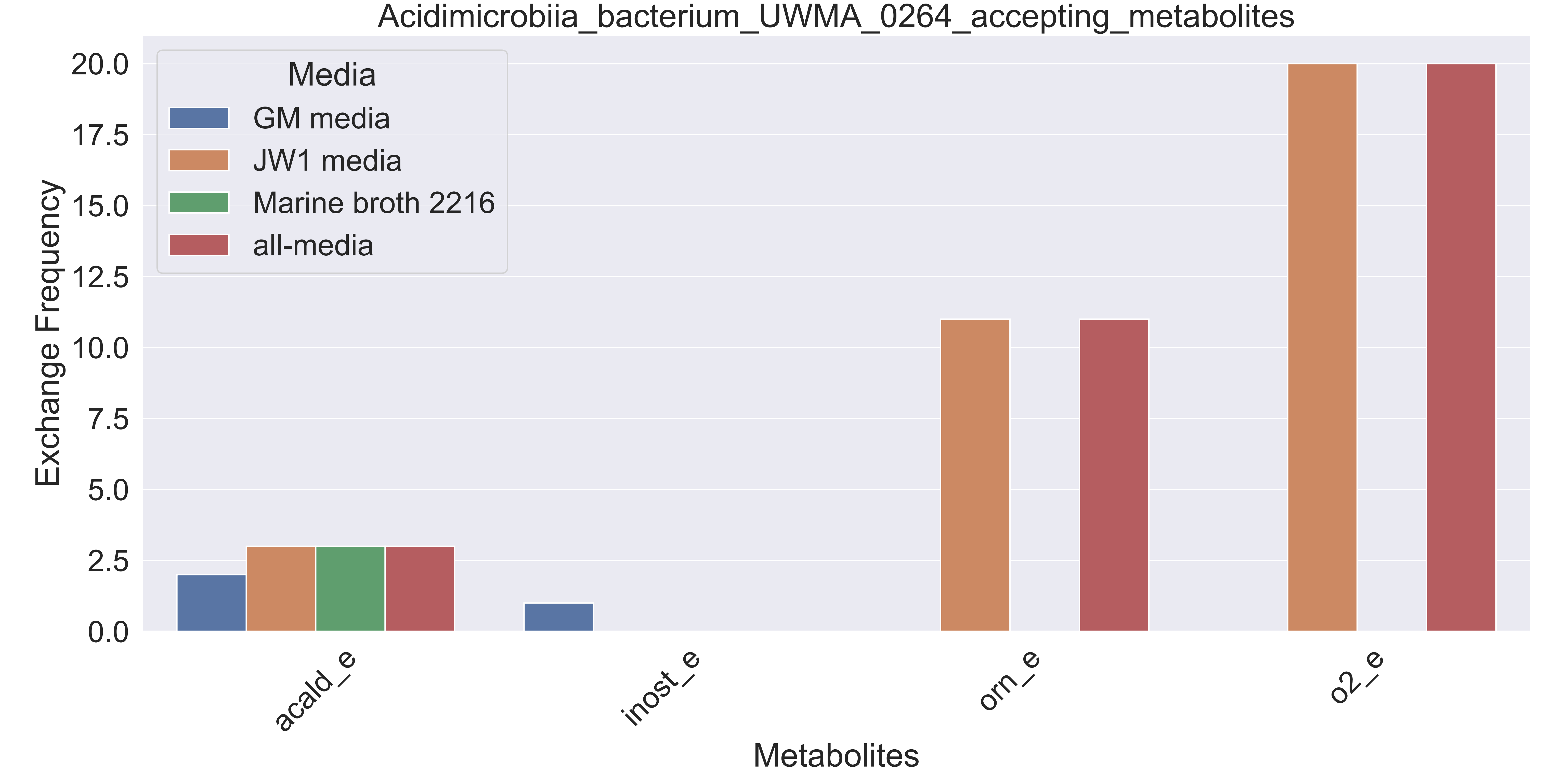

### Supplementary Figure S2 Acidimicrobiia bacterium UWMA 0303 accepting metabolites.pdf

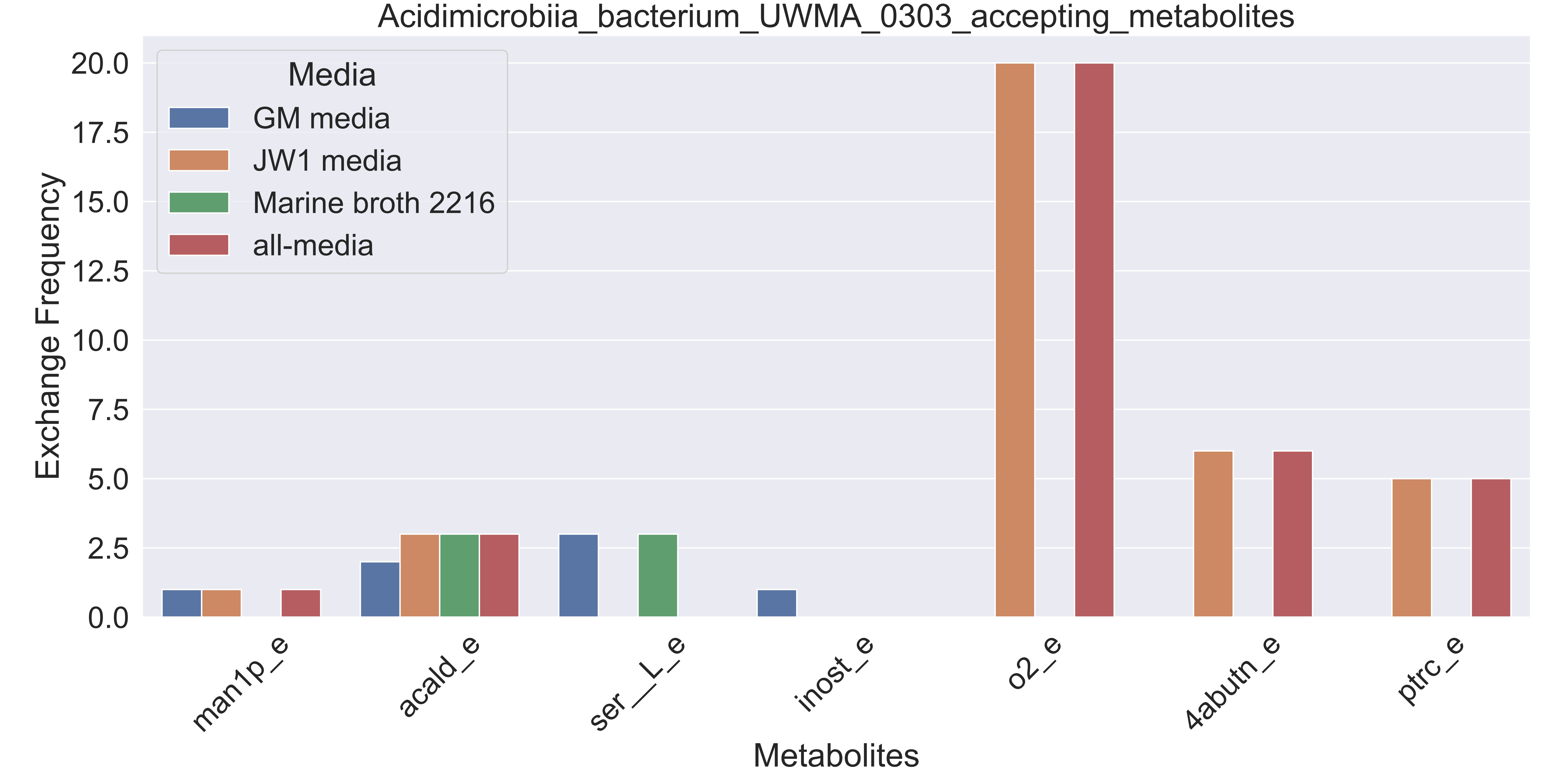

### Supplementary Figure S3 Acidimicrobiia bacterium UWMA 0319 accepting metabolites.pdf

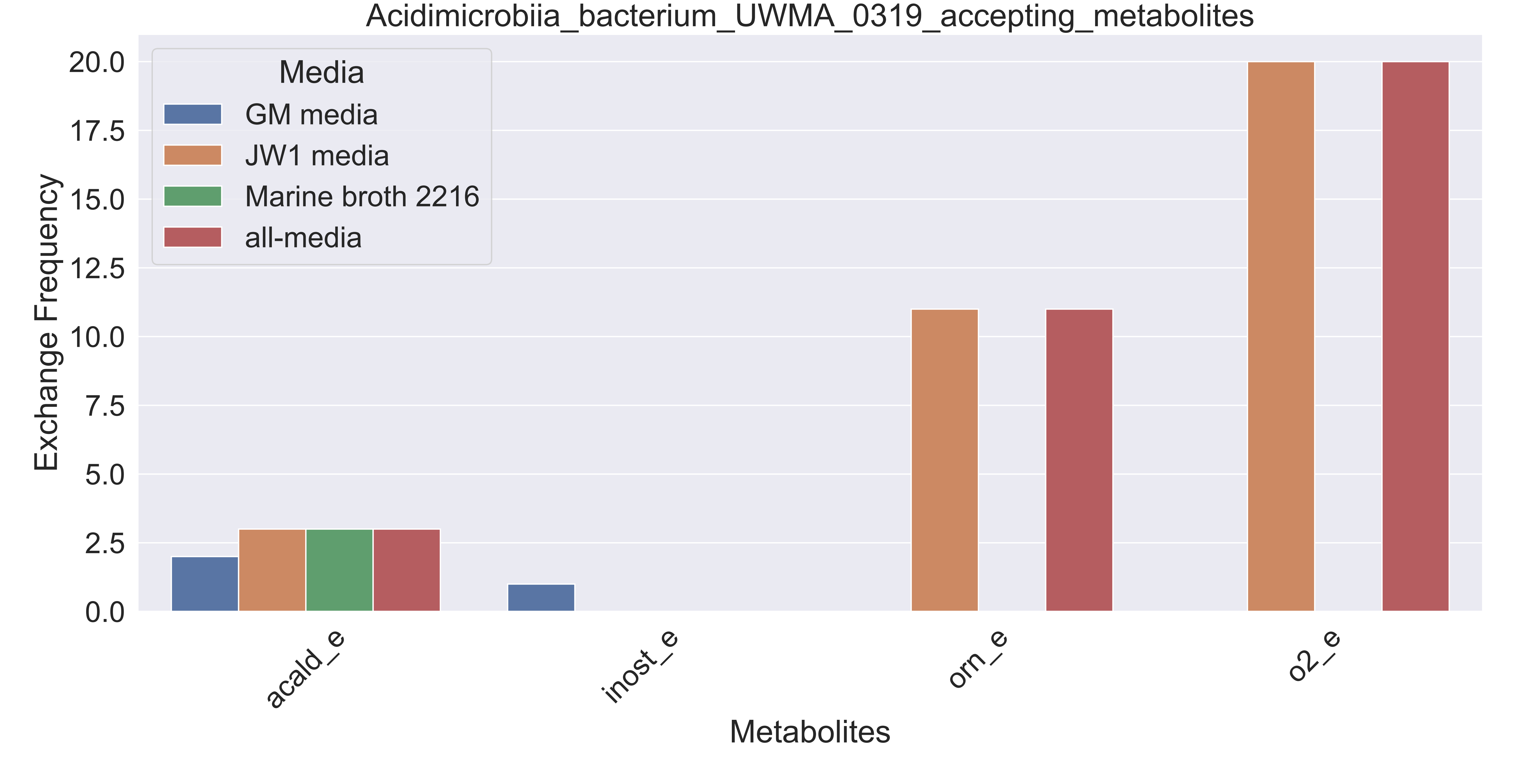

### Supplementary Figure S4 Acidobacteria bacterium UWMA 0355 accepting metabolites.pdf

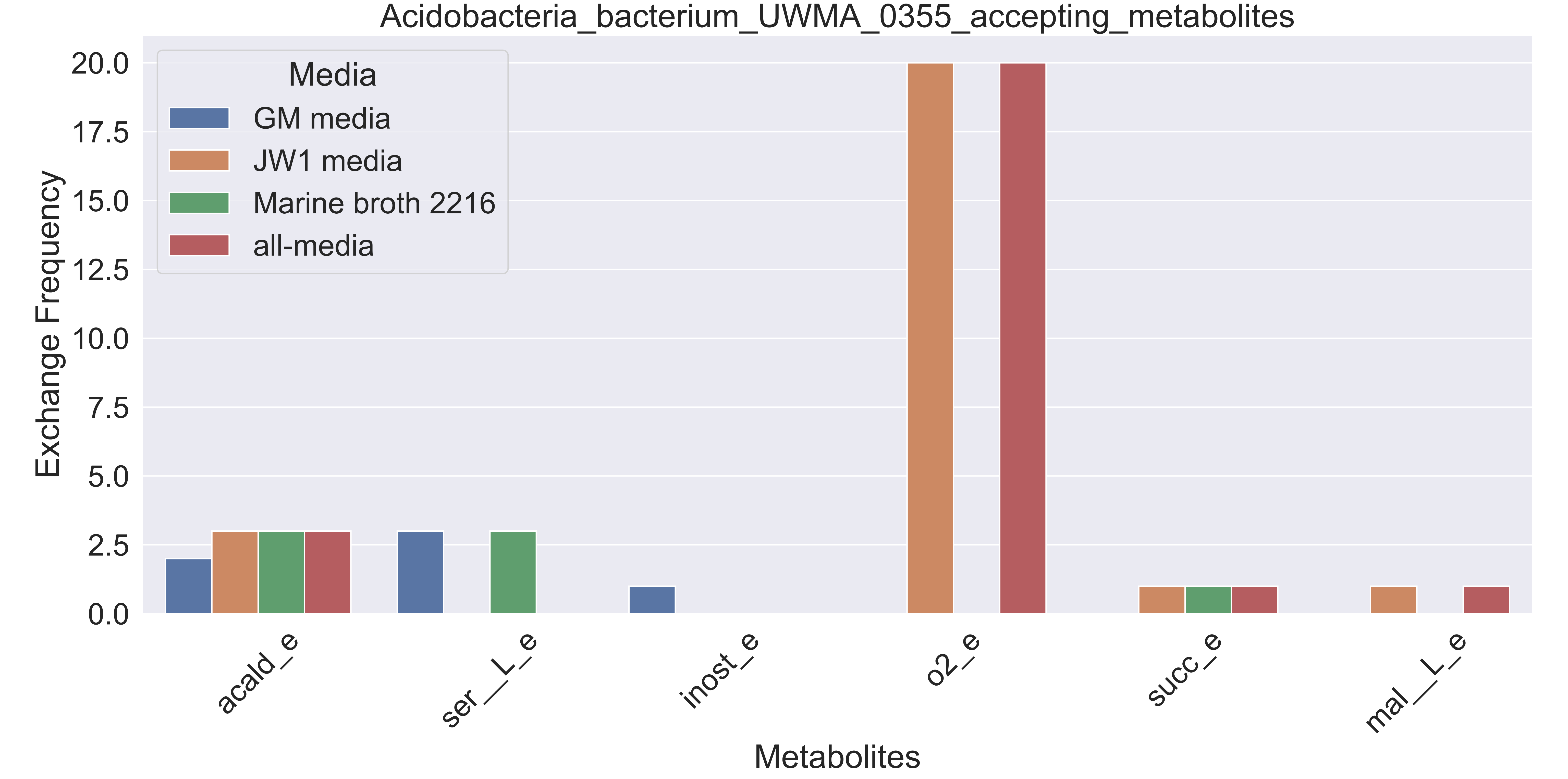

### Supplementary Figure S5 Alphaproteobacteria bacterium UWMA 0321 accepting metabolites.pdf

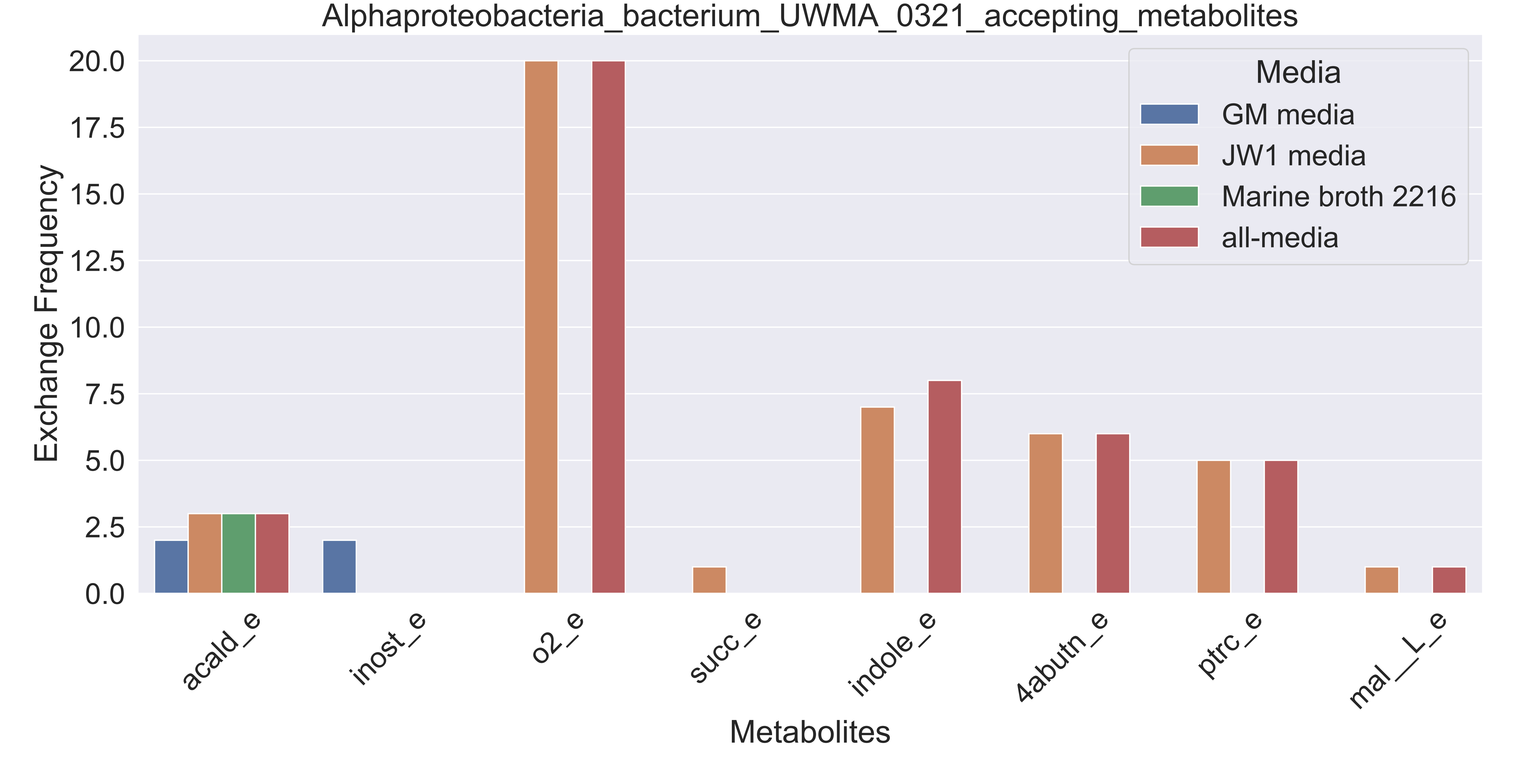

### Supplementary Figure S6 Bacteroidetes bacterium UWMA 0293 accepting metabolites.pdf

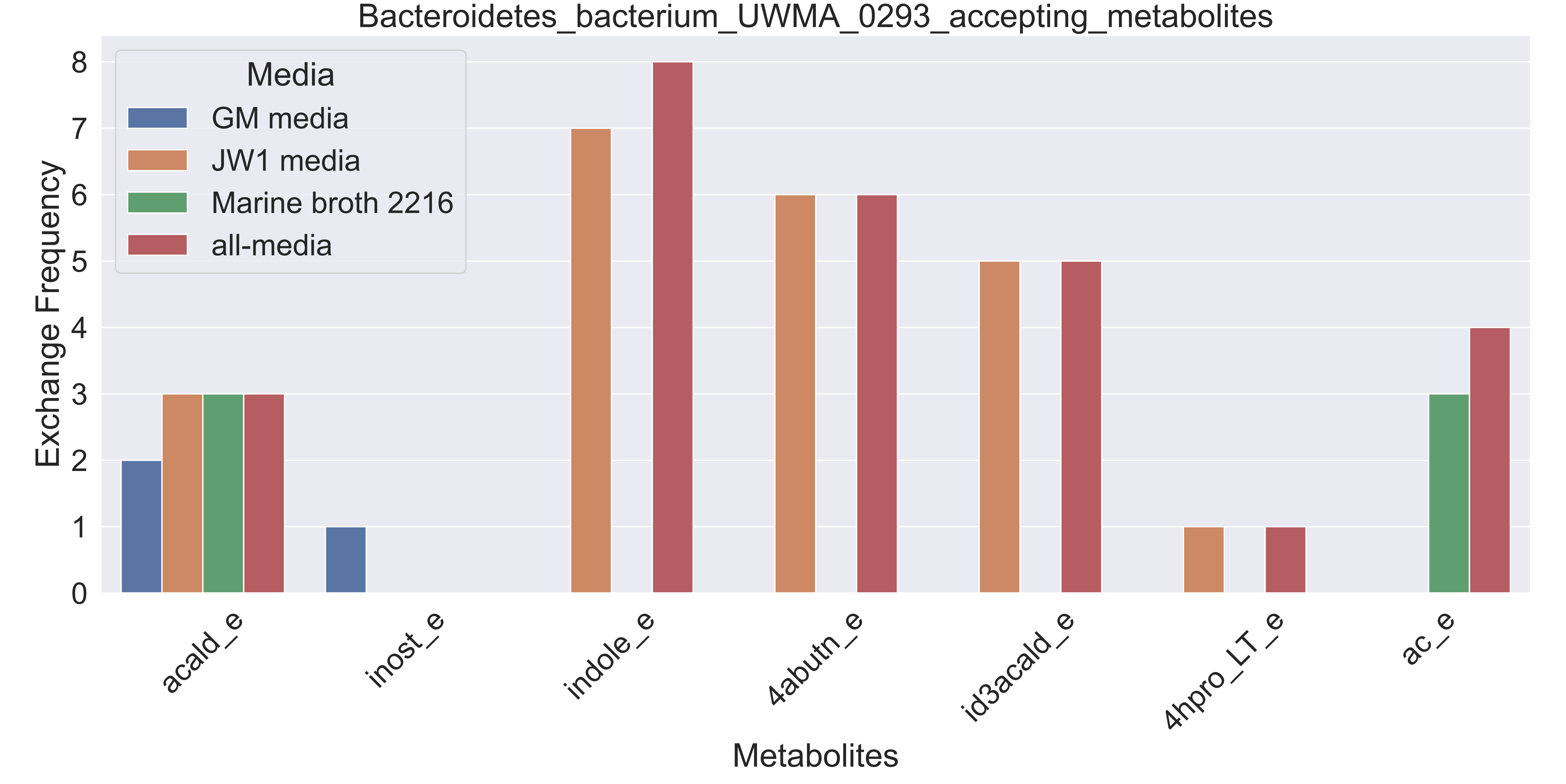

### Supplementary Figure S7 Bacteroidetes bacterium UWMA 0358 accepting metabolites.pdf

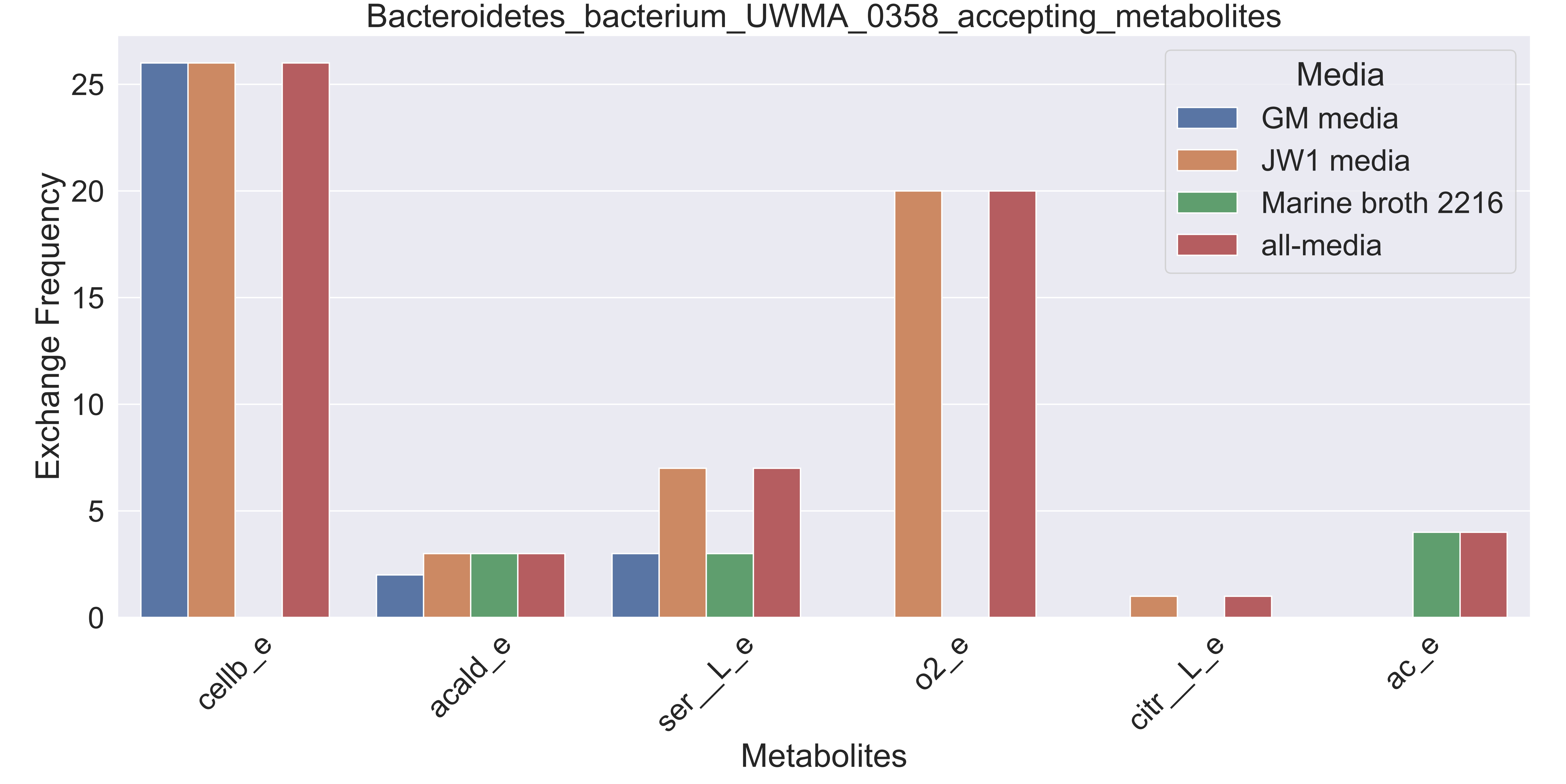

### Supplementary Figure S8 candidate division UBP10 bacterium UWMA 0304 accepting metabolites.pdf

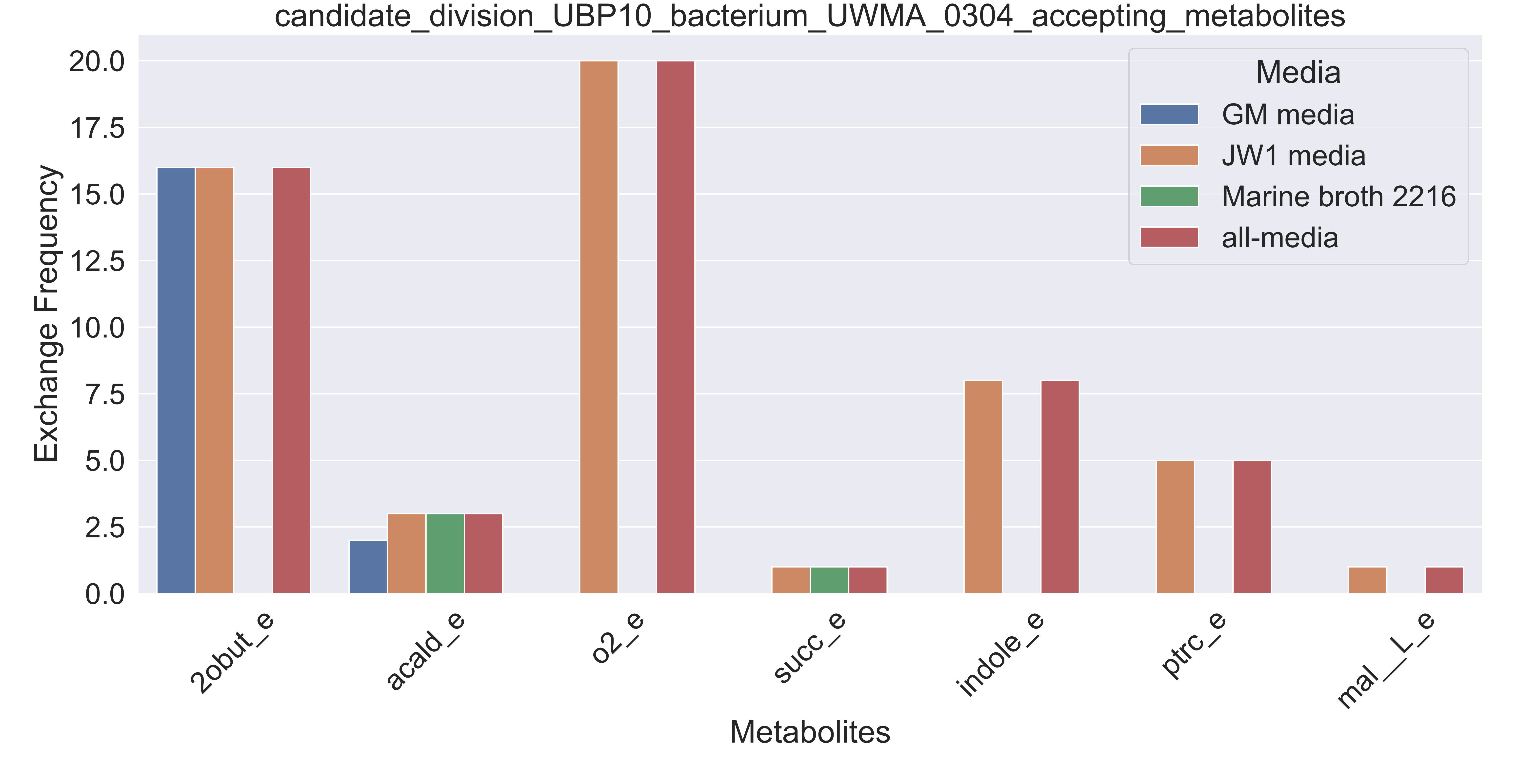

### Supplementary Figure S9 candidate division Zixibacteria bacterium UWMA 0288 accepting metabolites.pdf

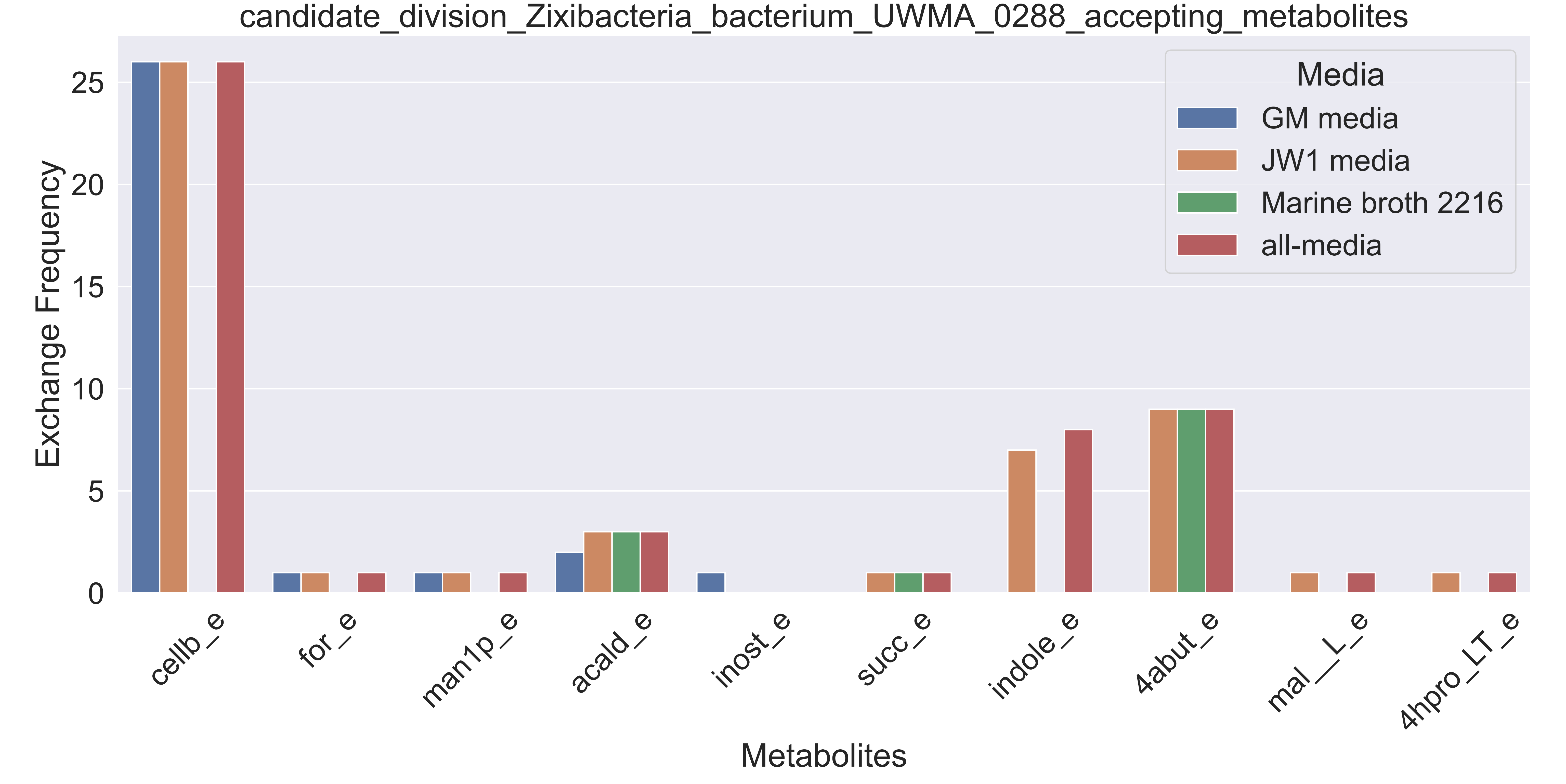

### Supplementary Figure S10 Candidatus Handelsmanbacteria bacterium UWMA 0286 accepting metabolites.pdf

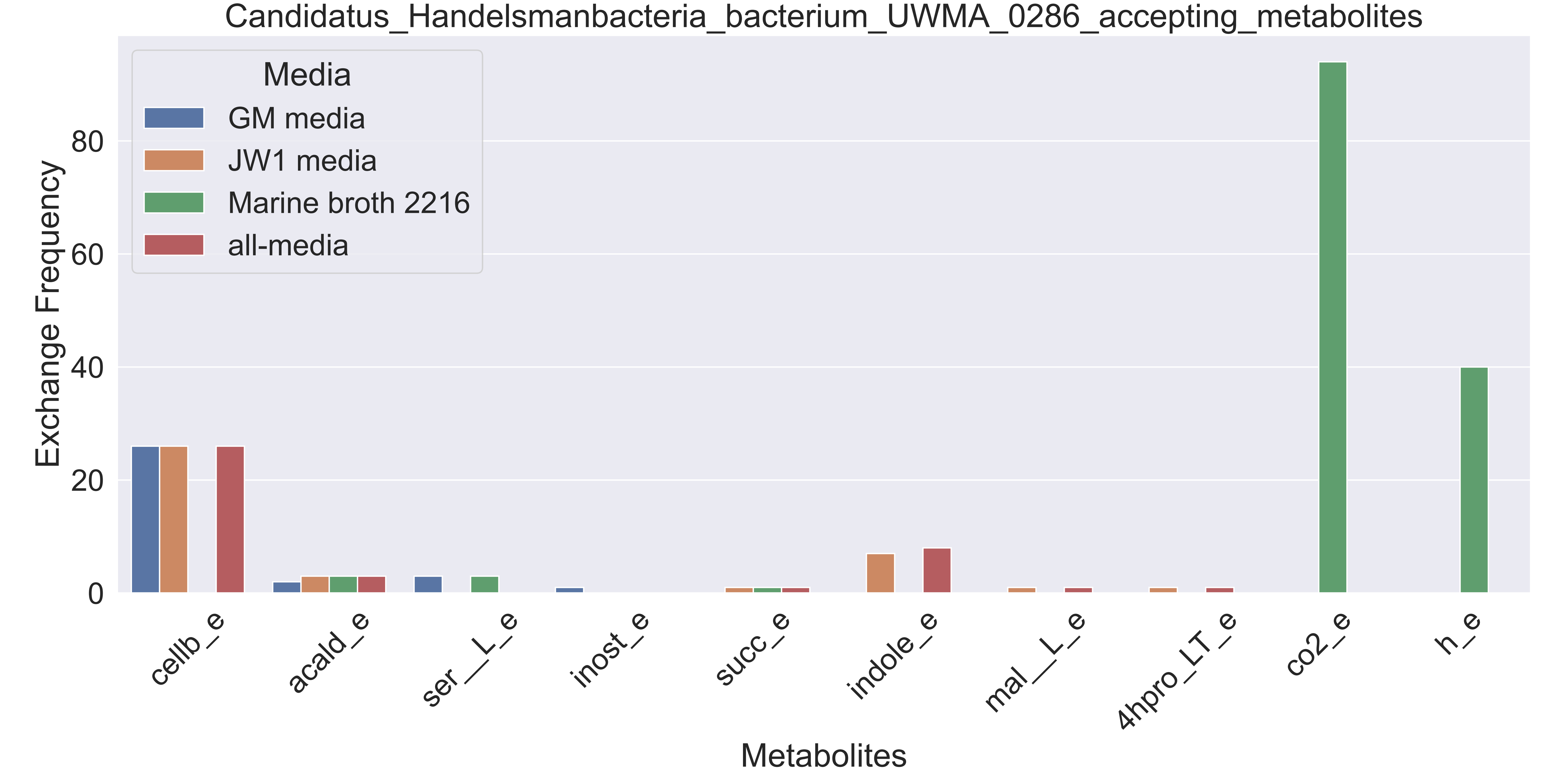

### Supplementary Figure S11 Candidatus Handelsmanbacteria bacterium UWMA 0300 accepting metabolites.pdf

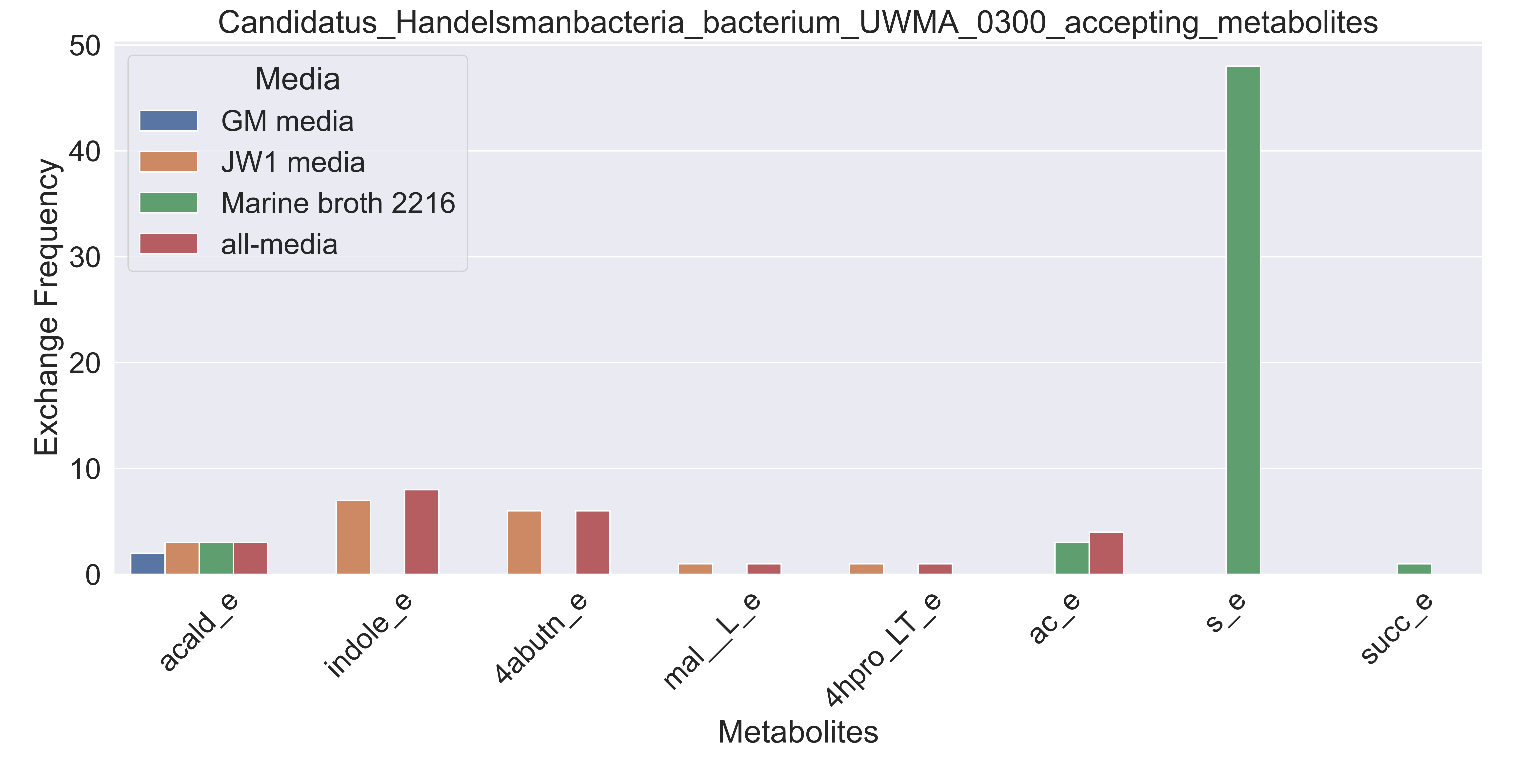

### Supplementary Figure S12 Candidatus Lambdaproteobacteria bacterium UWMA 0298 accepting metabolites.pdf

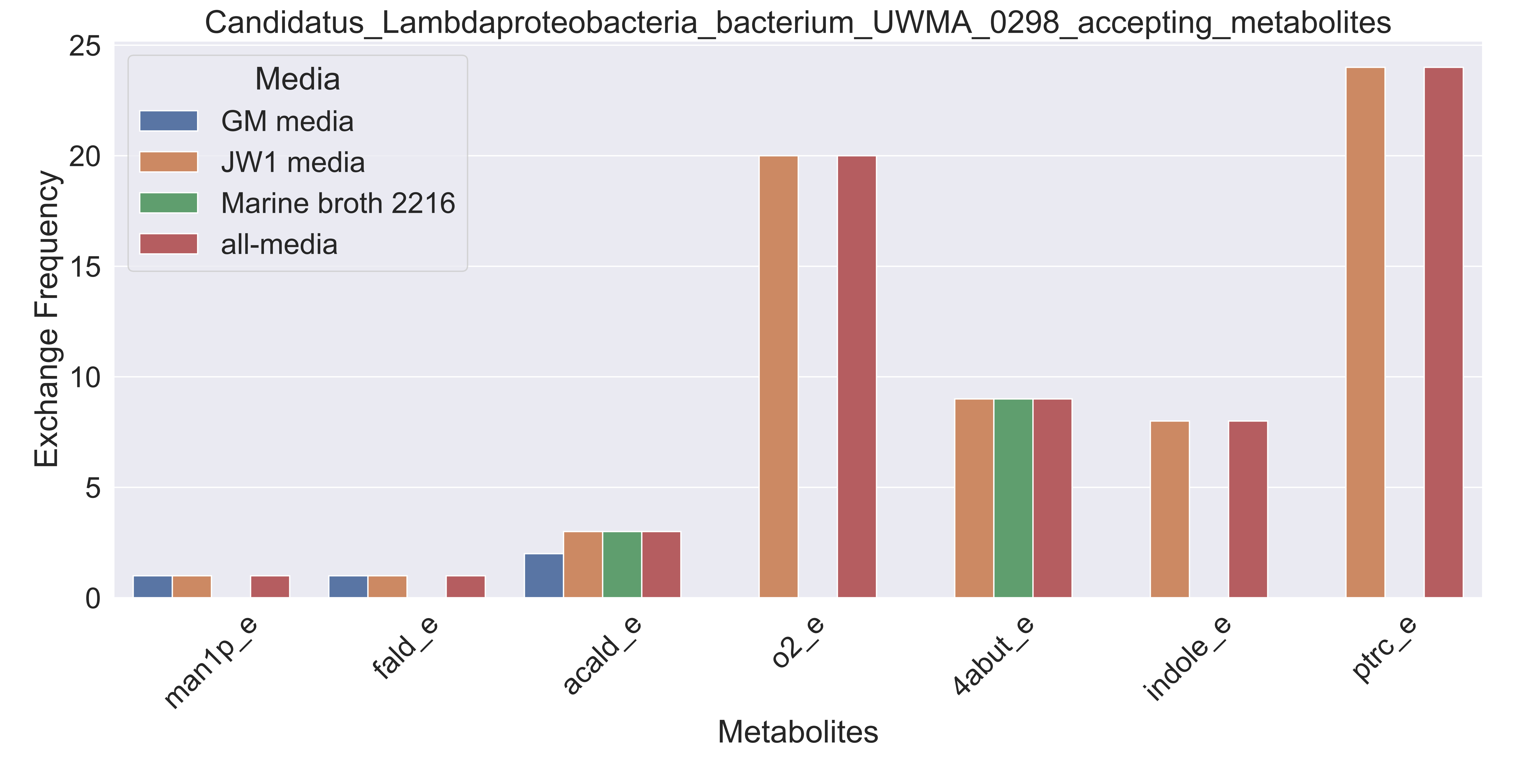

### Supplementary Figure S13 Candidatus Lambdaproteobacteria bacterium UWMA 0318 accepting metabolites.pdf

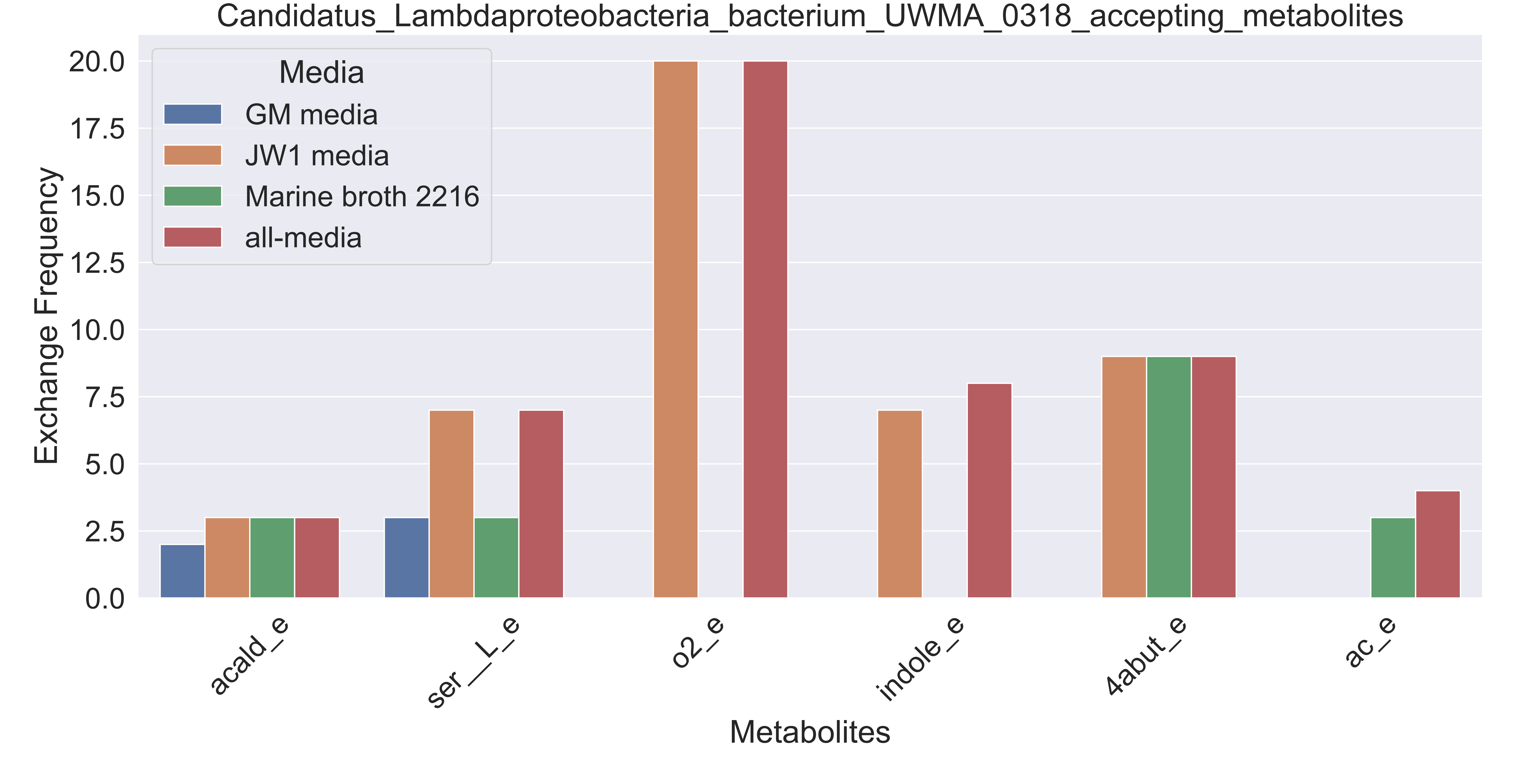

### Supplementary Figure S14 Candidatus Marinimicrobia bacterium UWMA 0285 accepting metabolites.pdf

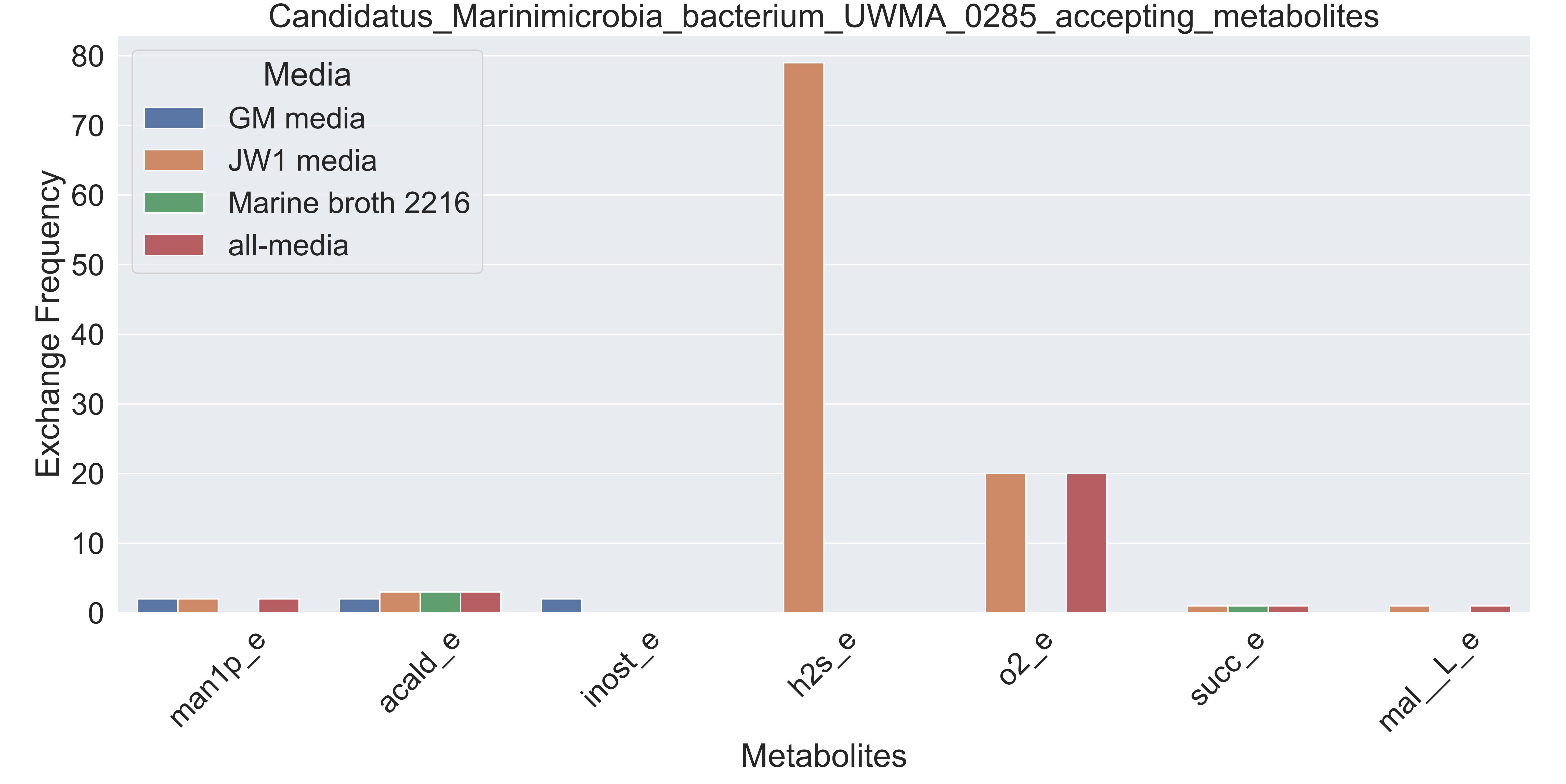

### Supplementary Figure S15 Candidatus Marinimicrobia bacterium UWMA 0306 accepting metabolites.pdf

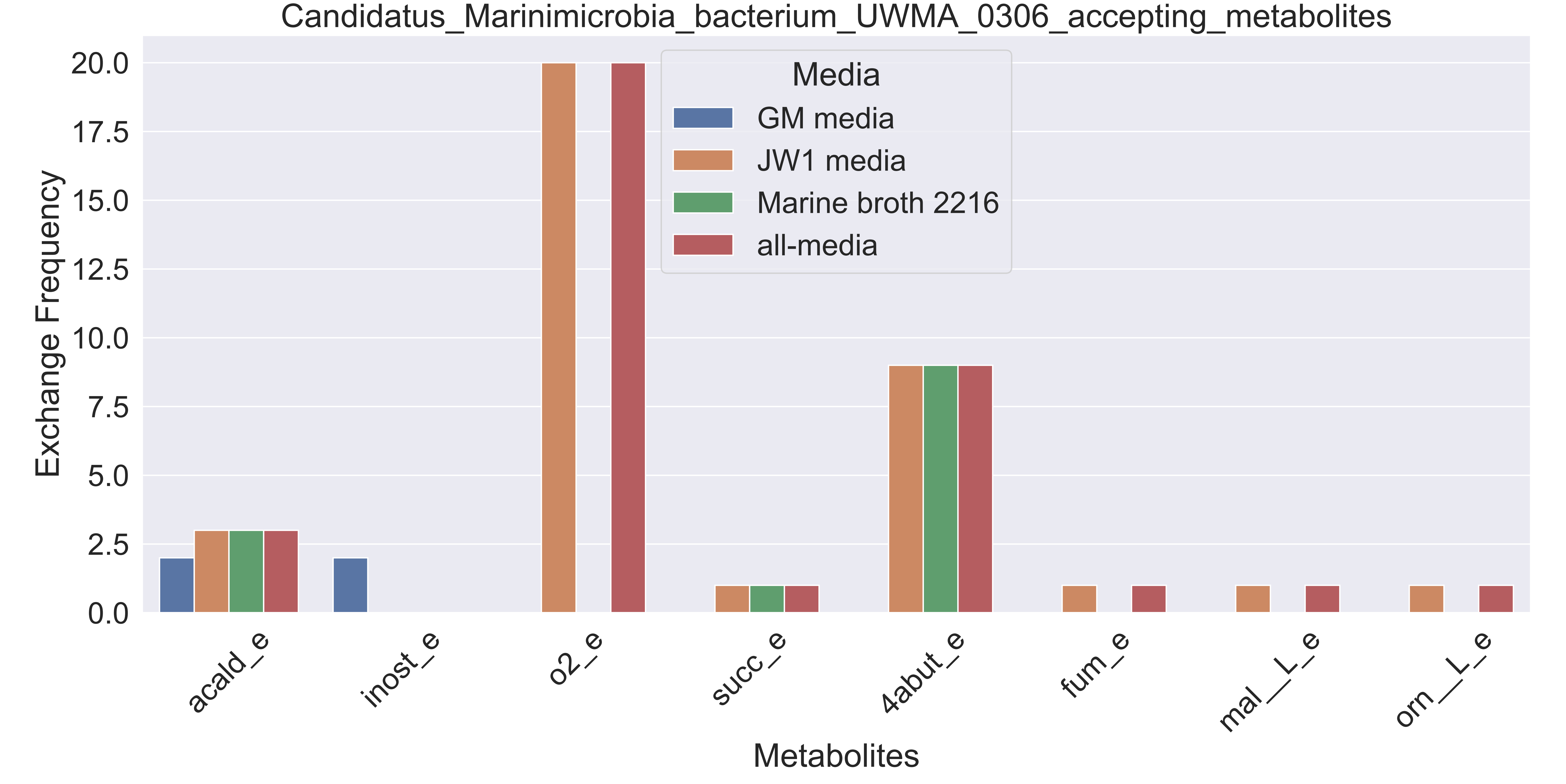

### Supplementary Figure S16 Candidatus Marinimicrobia bacterium UWMA 0309 accepting metabolites.pdf

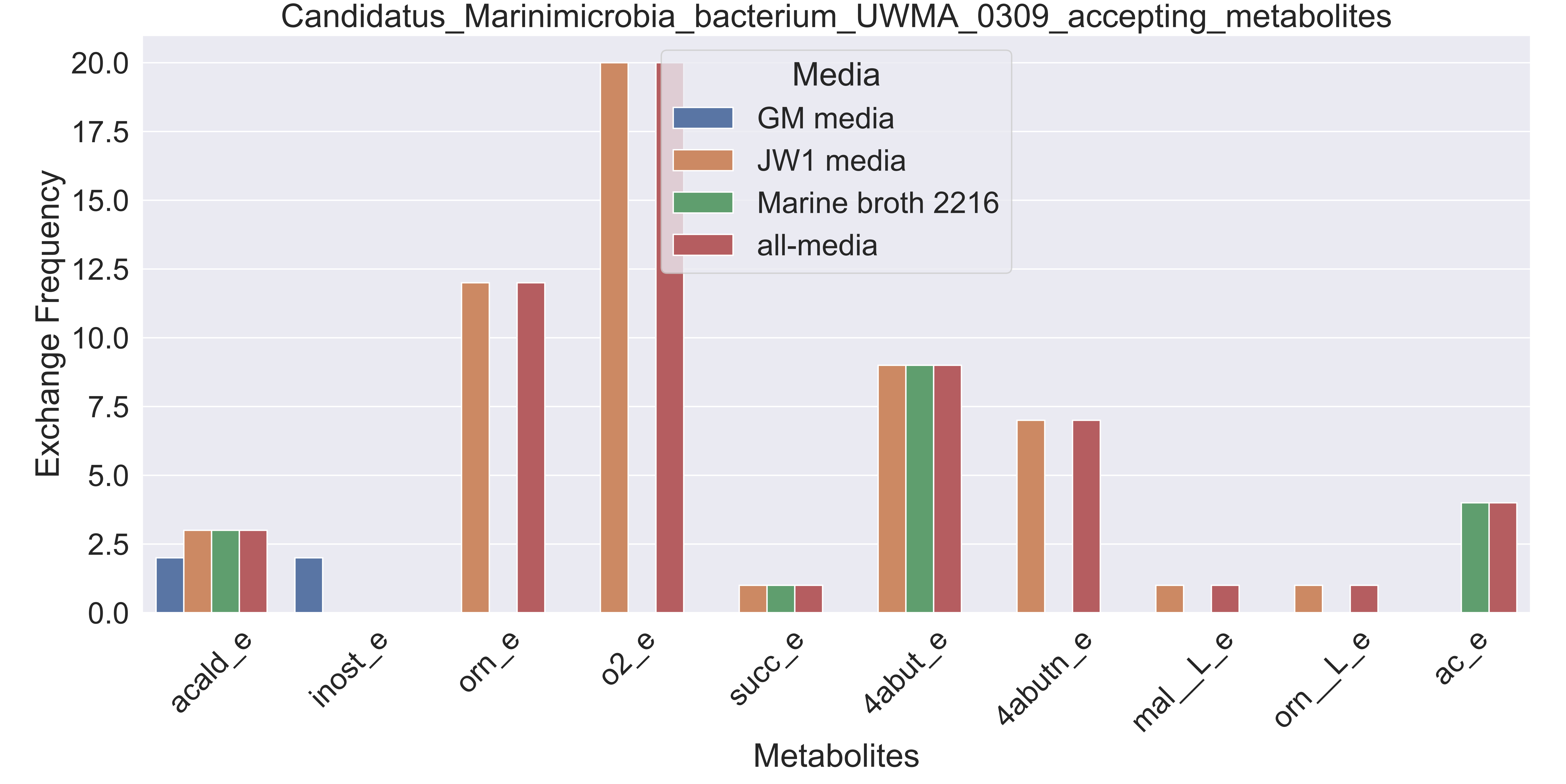

### Supplementary Figure S17 Candidatus Marinimicrobia bacterium UWMA 0347 accepting metabolites.pdf

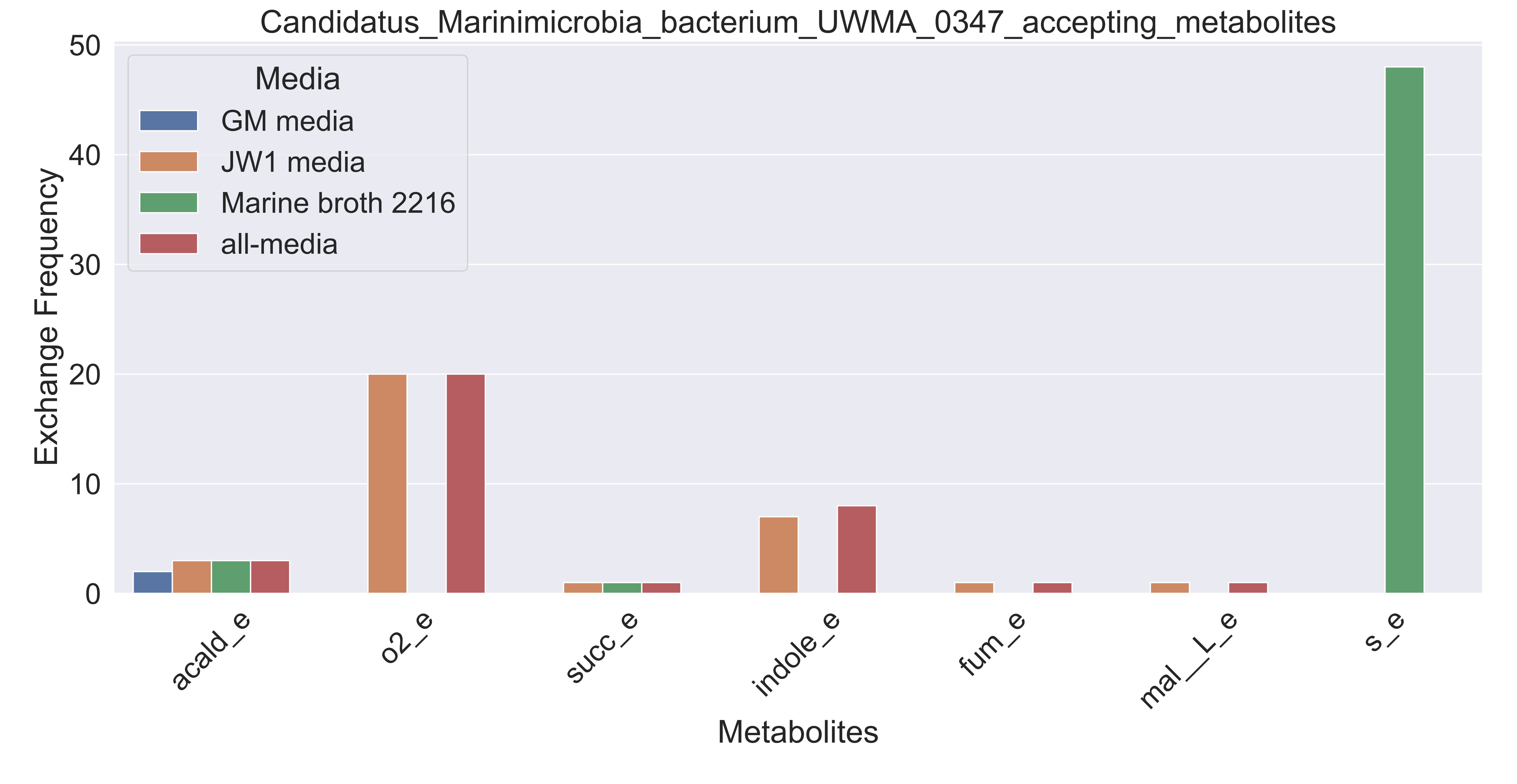

### Supplementary Figure S18 Candidatus Nitrosopelagicus sp UWMA 0359 accepting metabolites.pdf

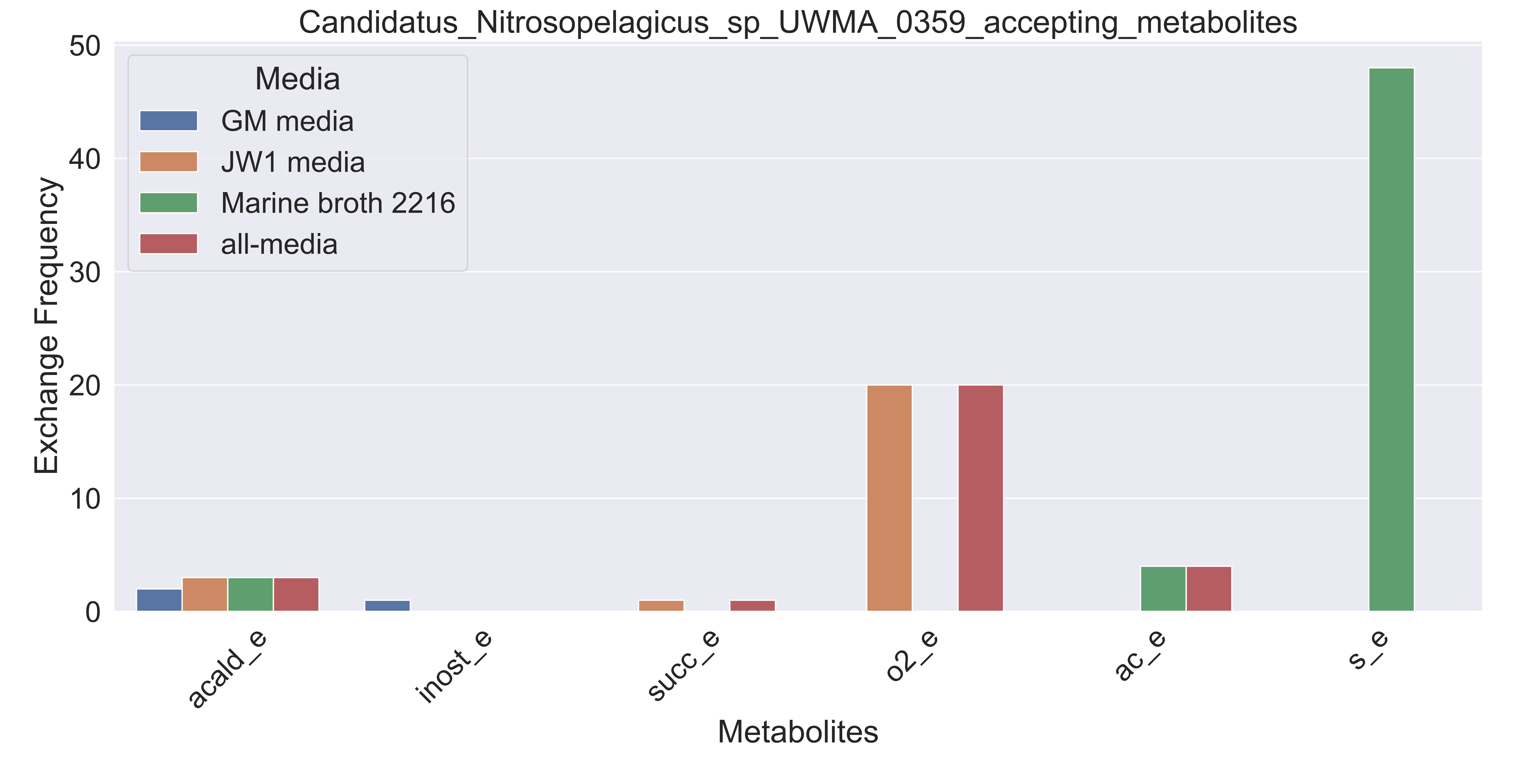

### Supplementary Figure S19 Candidatus Pacearchaeota archaeon UWMA 0287 accepting metabolites.pdf

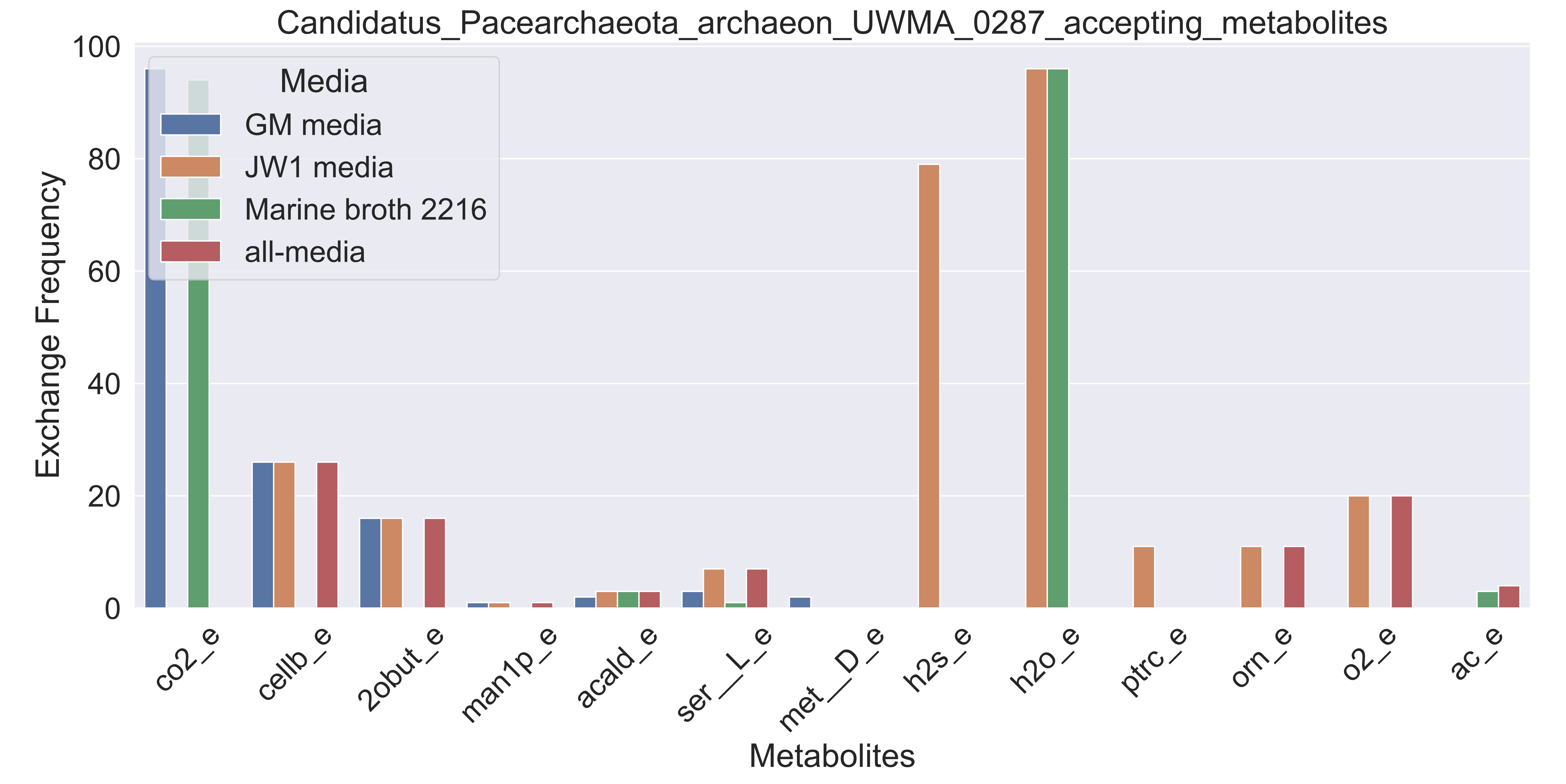

### Supplementary Figure S20 Candidatus Pelagibacter sp UWMA 0307 accepting metabolites.pdf

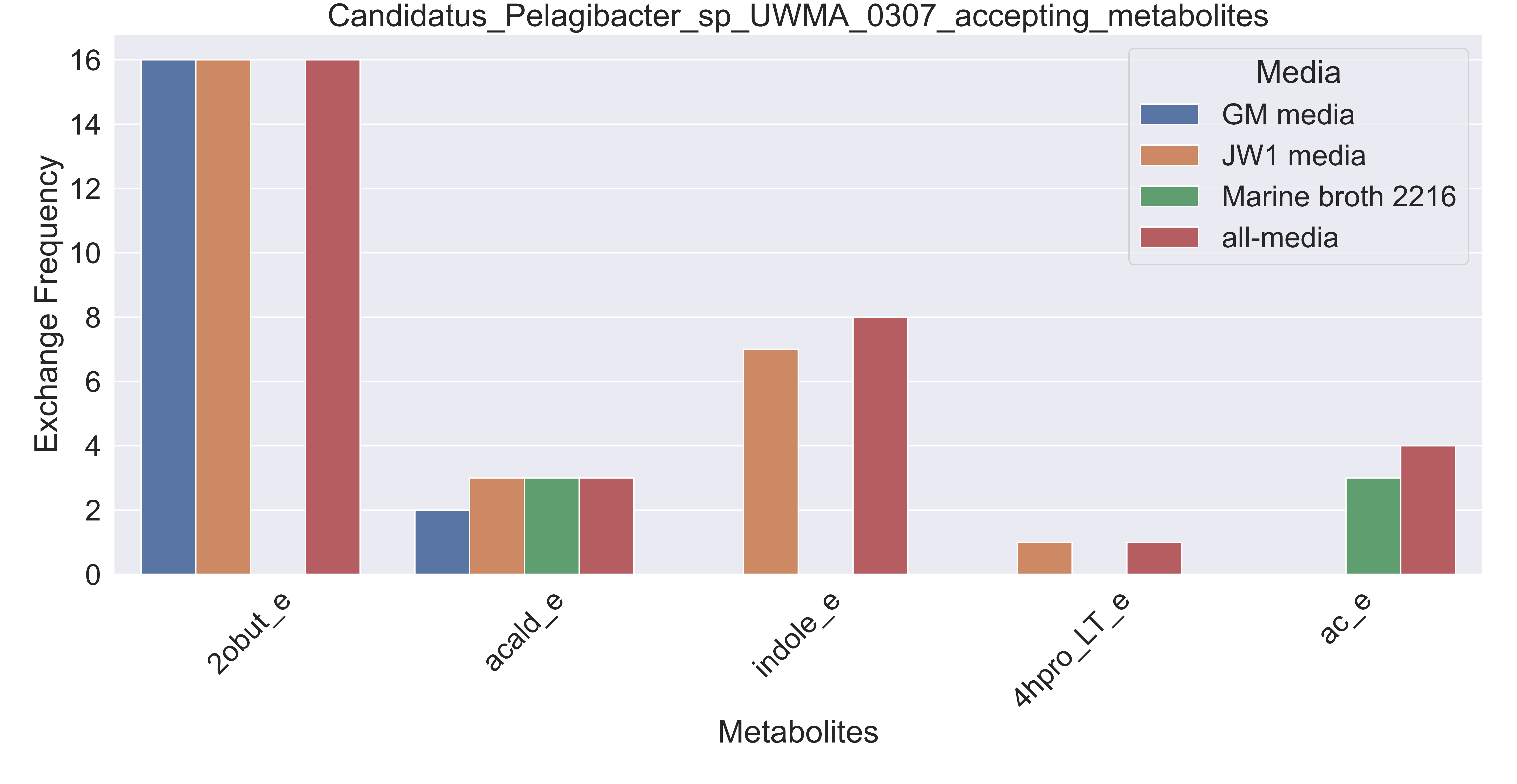

### Supplementary Figure S21 Candidatus Thioglobus sp UWMA 0259 accepting metabolites.pdf

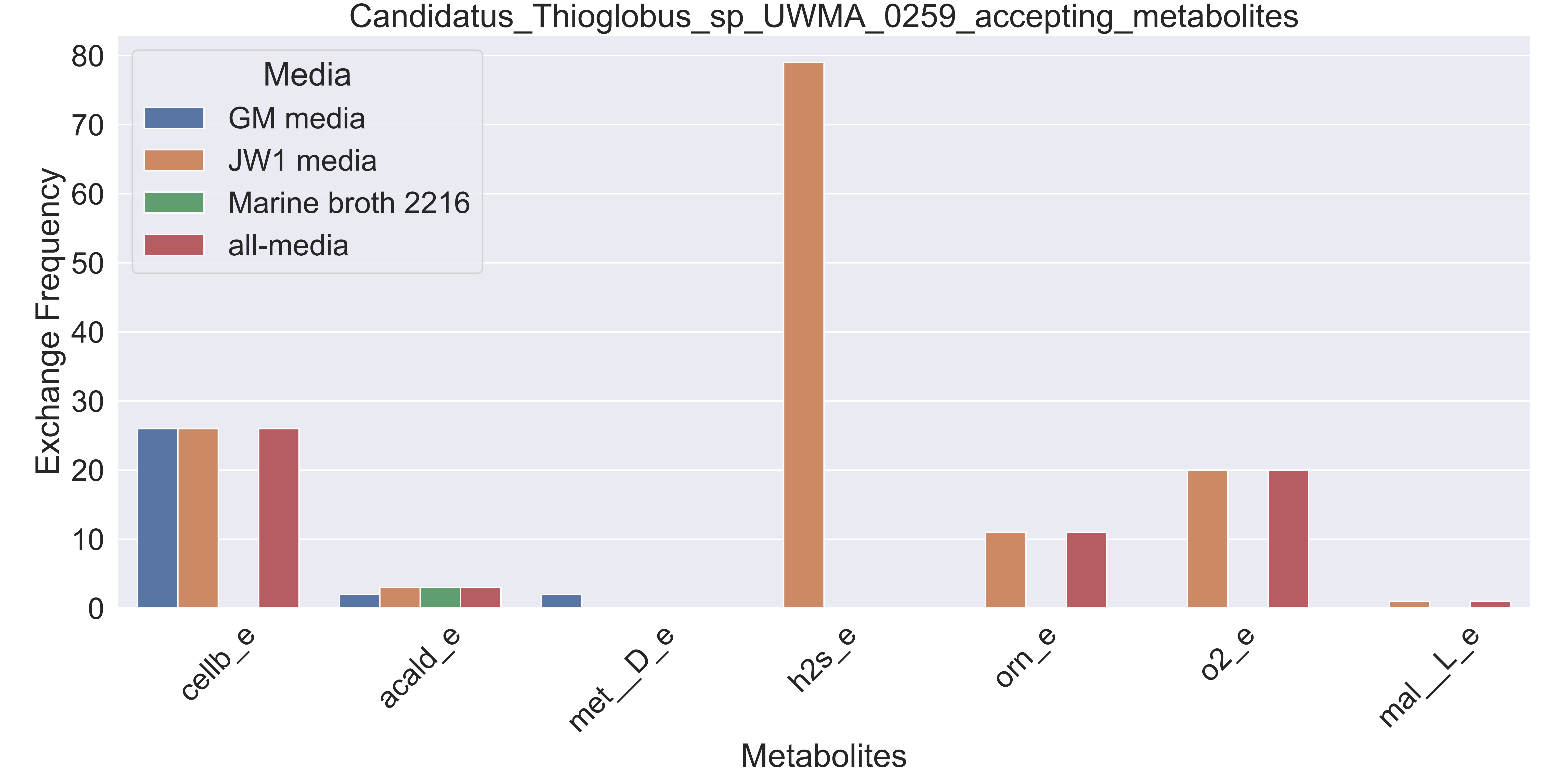

### Supplementary Figure S22 Candidatus Thioglobus sp UWMA 0272 accepting metabolites.pdf

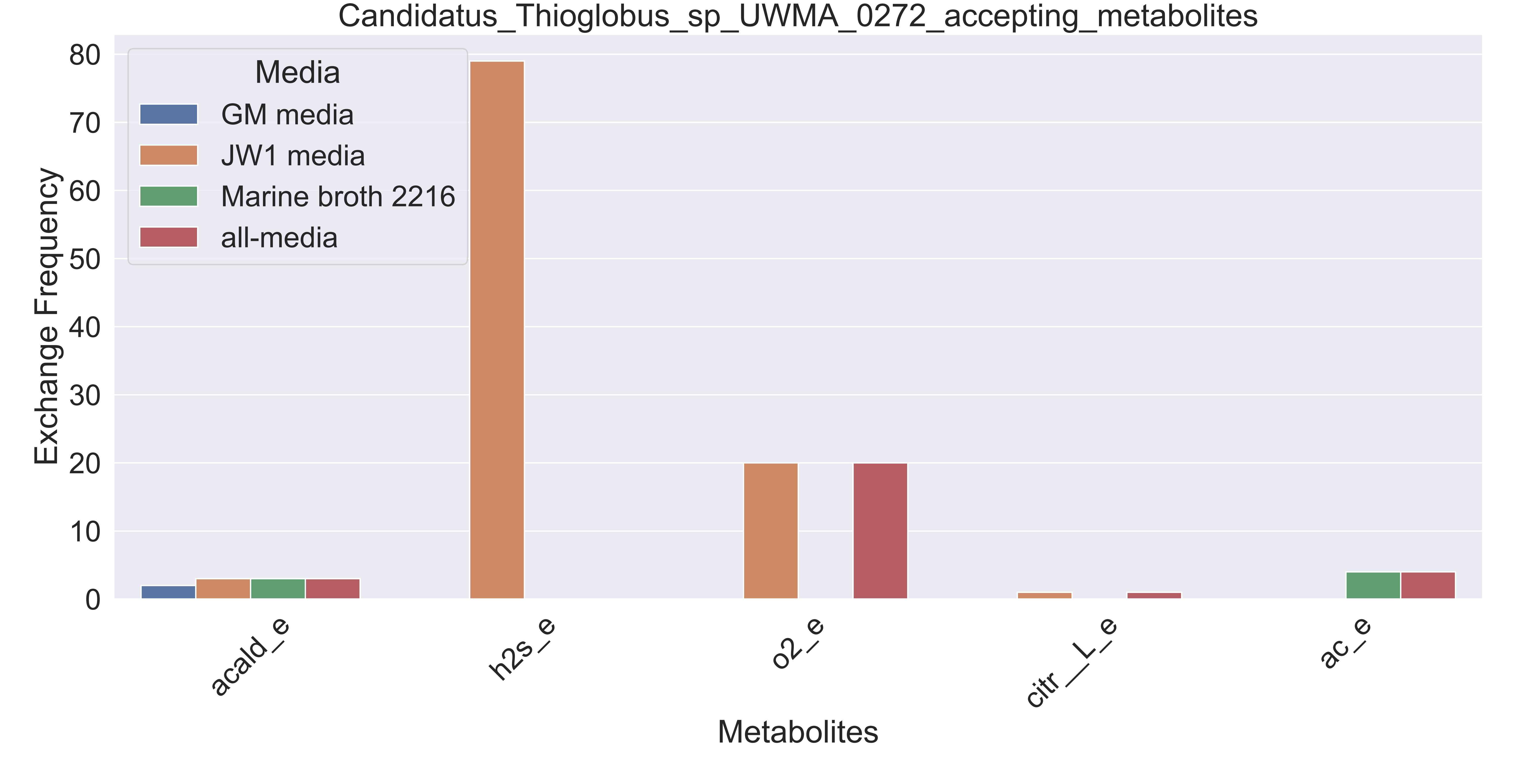

### Supplementary Figure S23 Candidatus Thioglobus sp UWMA 0322 accepting metabolites.pdf

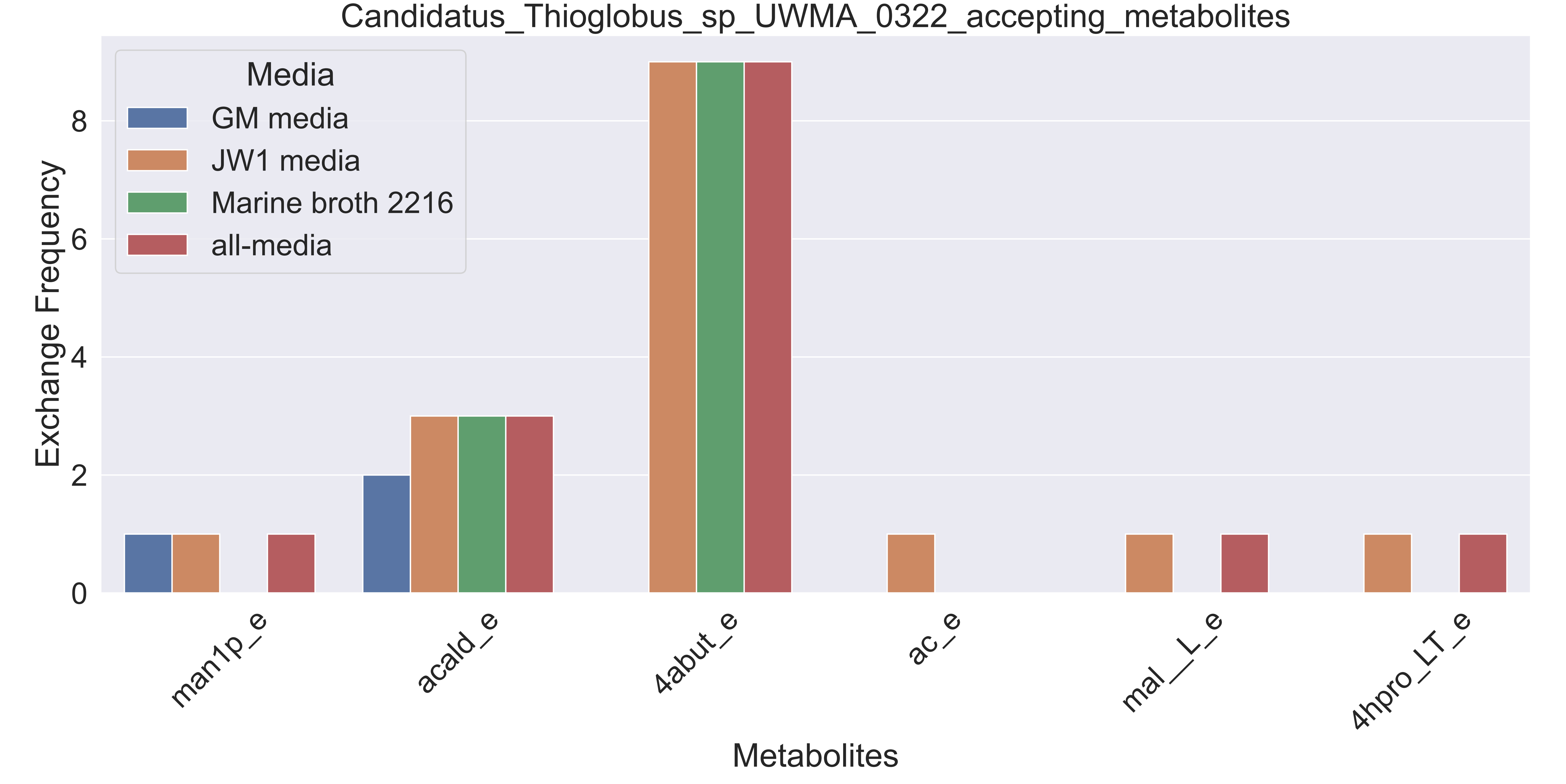

### Supplementary Figure S24 Candidatus Thioglobus sp UWMA 0342 accepting metabolites.pdf

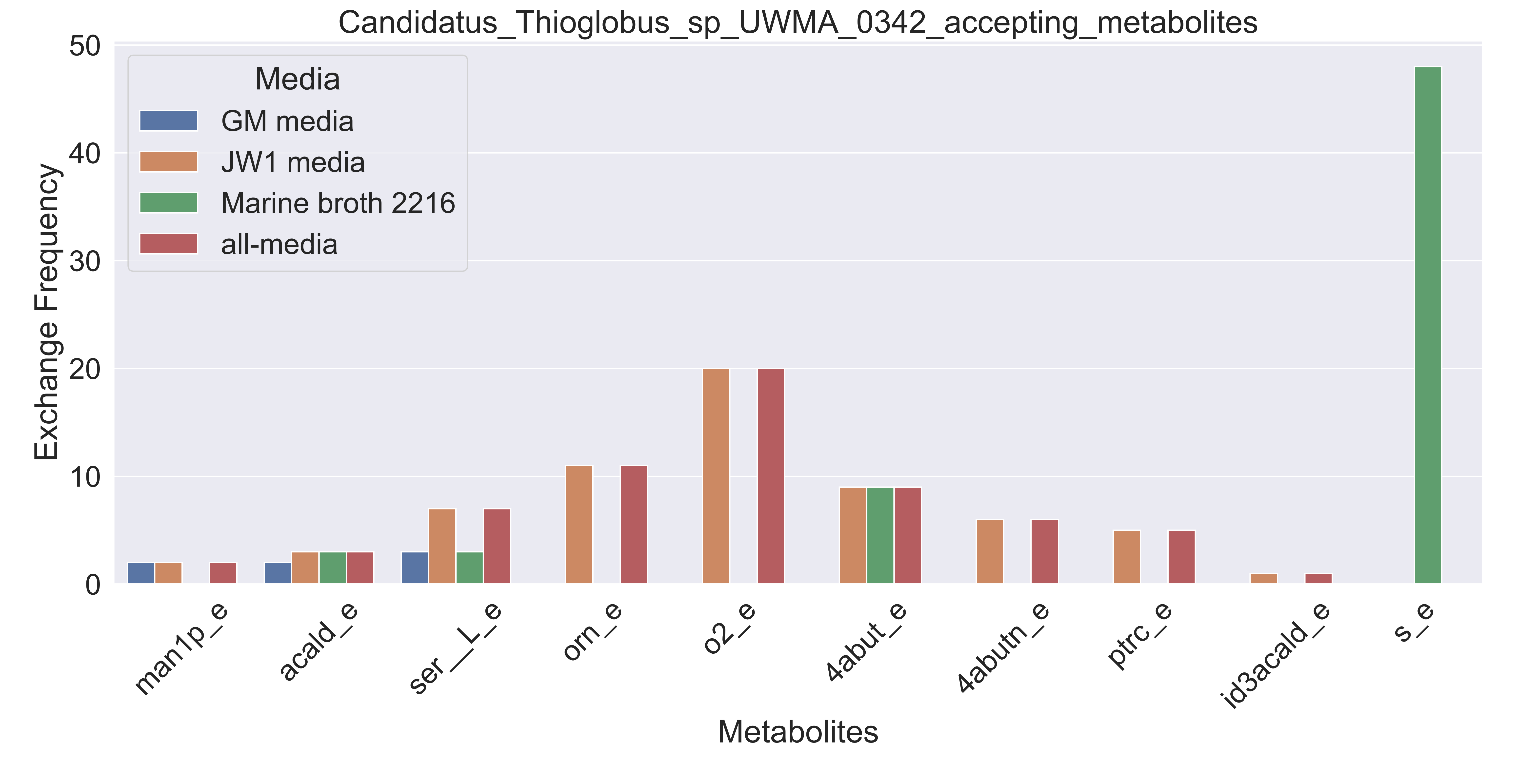

### Supplementary Figure S25 Candidatus Thioglobus sp UWMA 0360 accepting metabolites.pdf

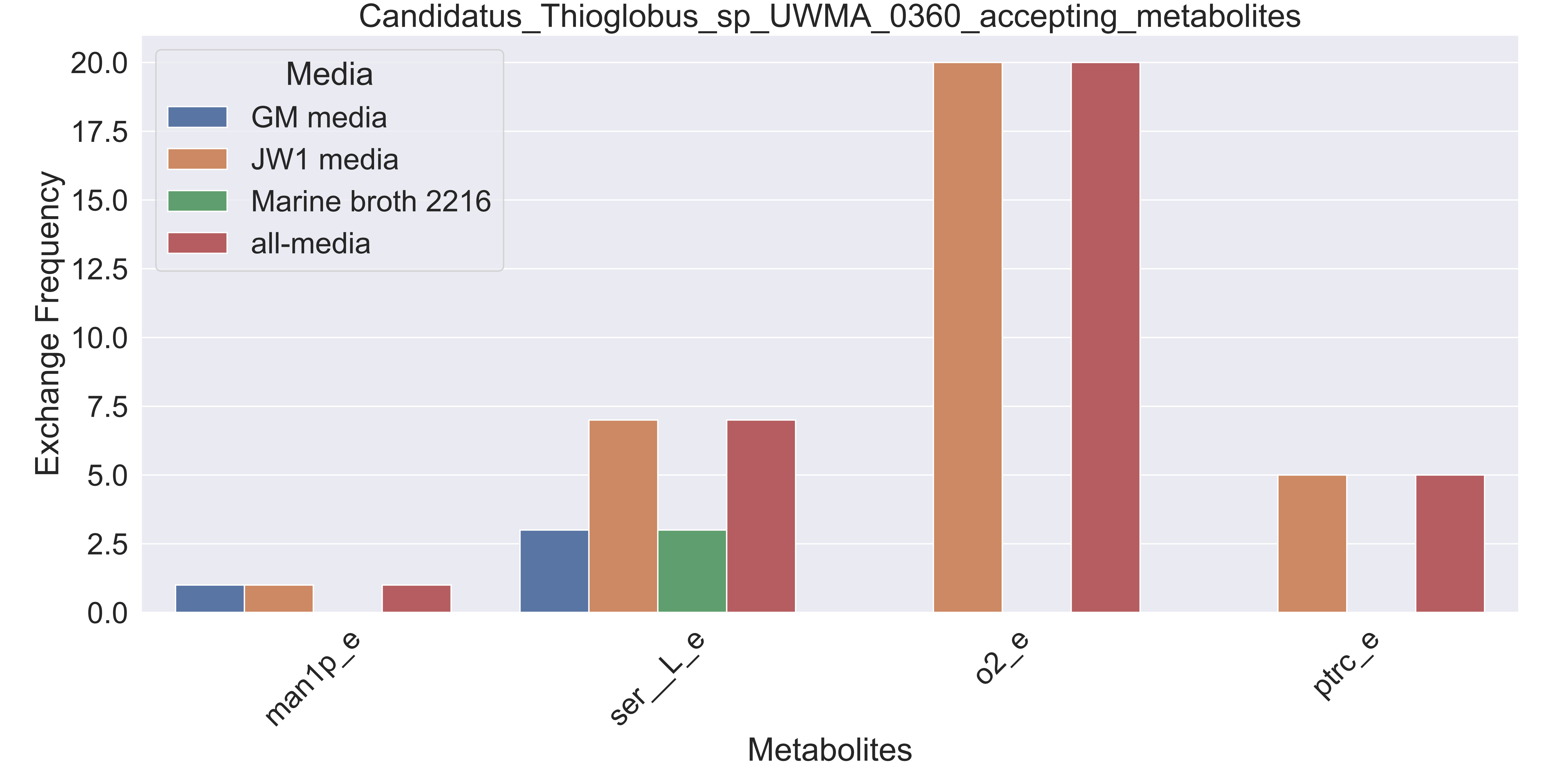

### Supplementary Figure S26 Cycloclasticus sp UWMA 0316 accepting metabolites.pdf

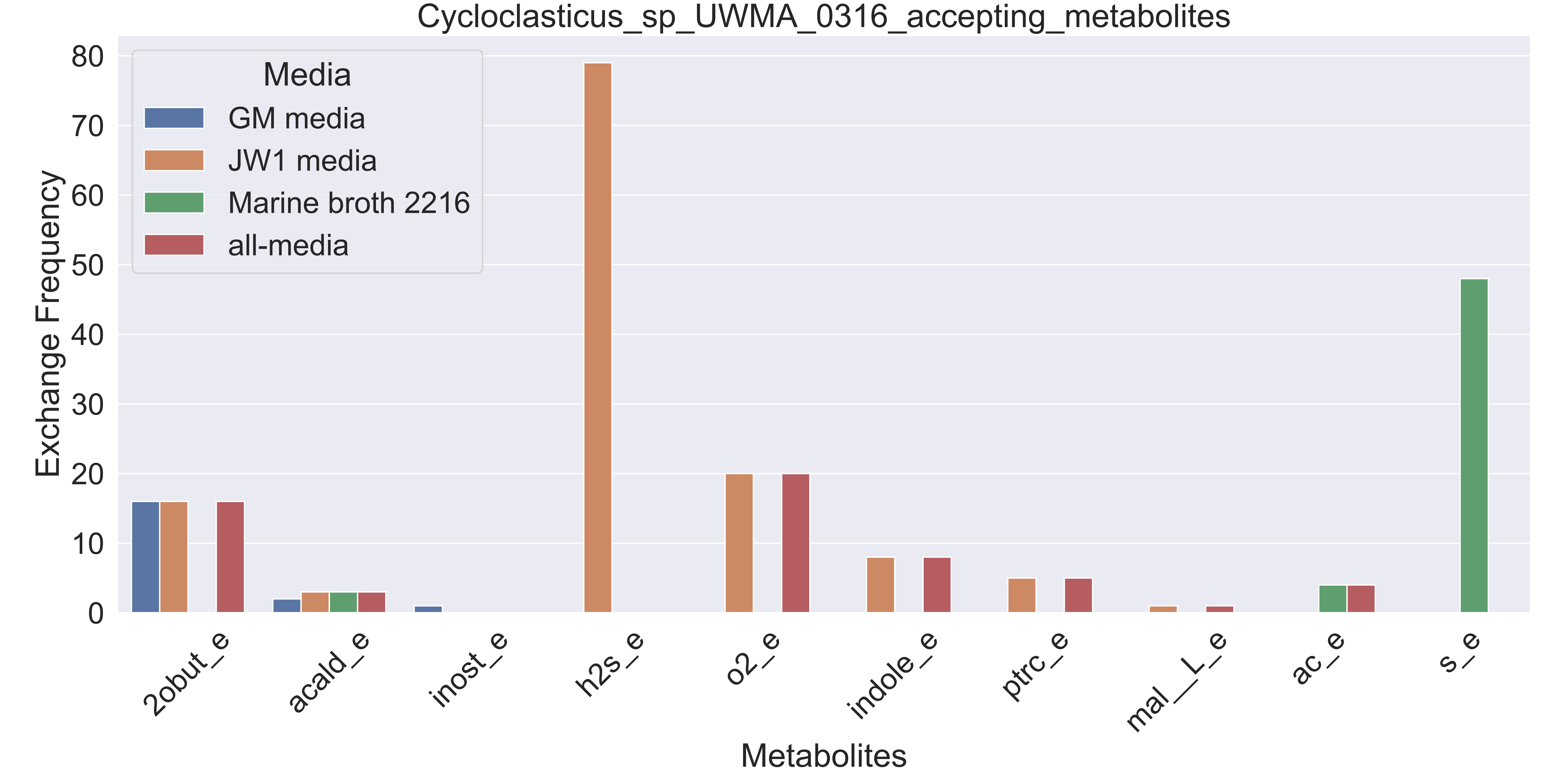

### Supplementary Figure S27 Cycloclasticus sp UWMA 0335 accepting metabolites.pdf

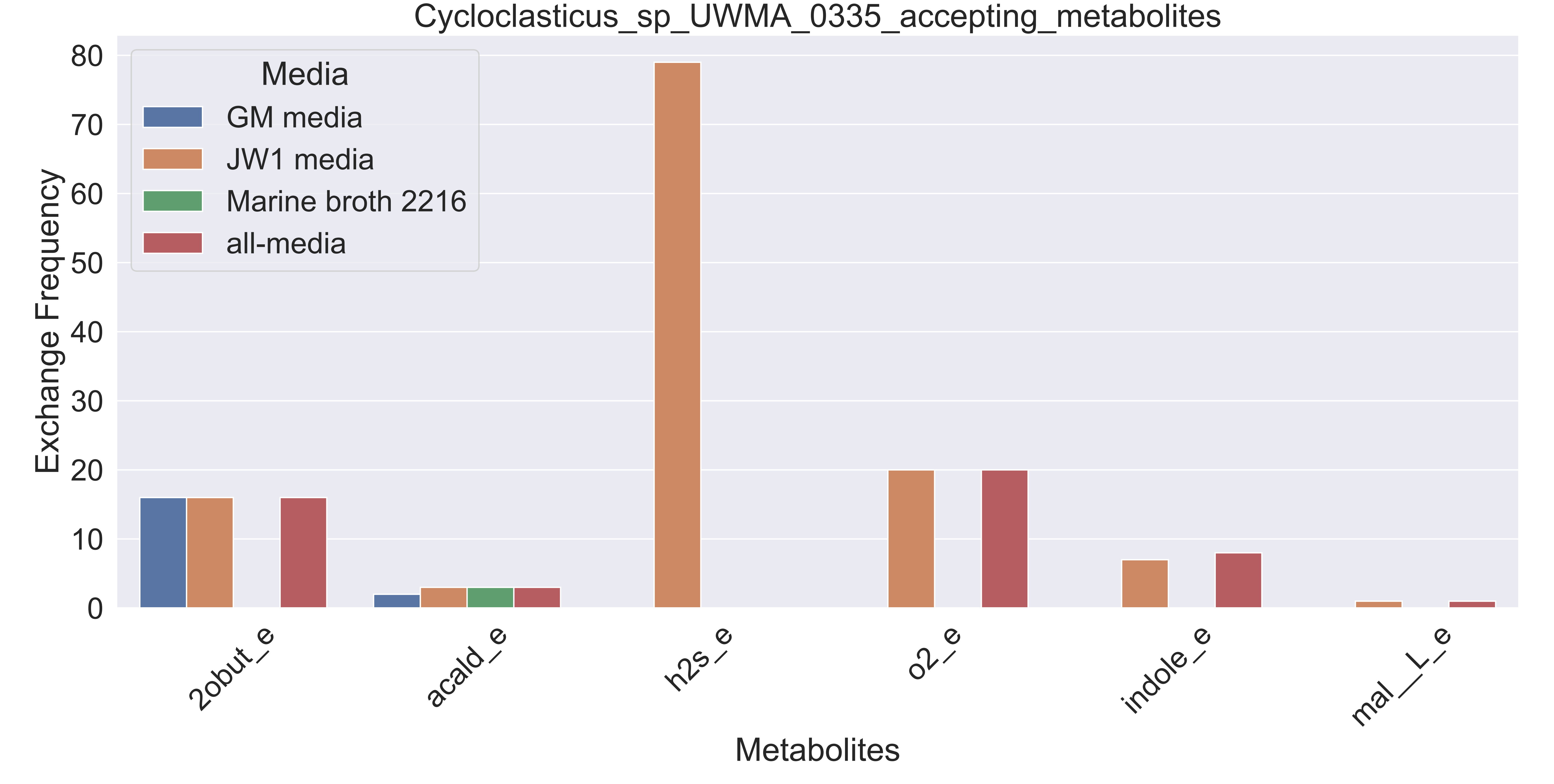

### Supplementary Figure S28 Cytophagales bacterium UWMA 0338 accepting metabolites.pdf

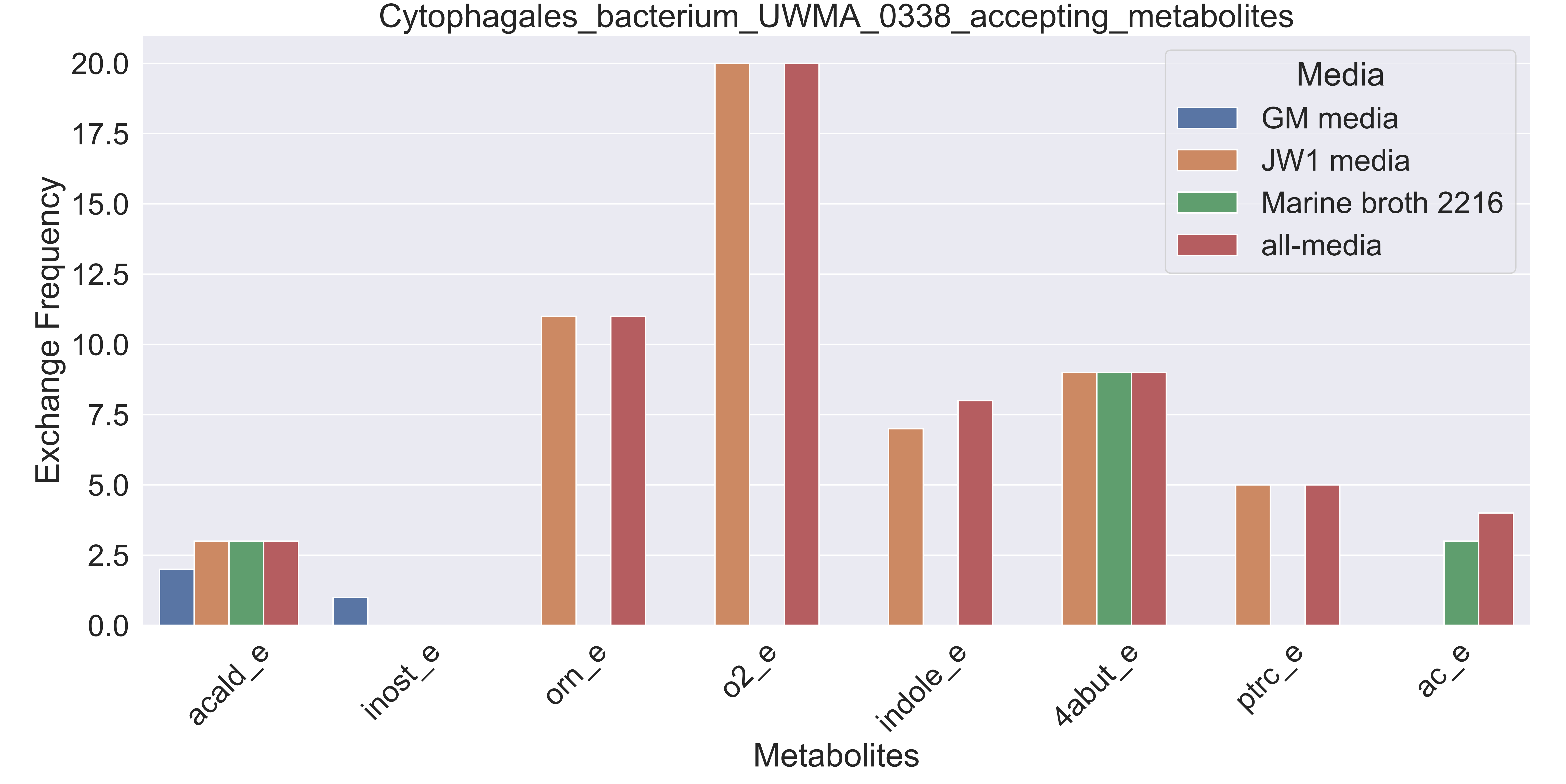

### Supplementary Figure S29 Dehalococcoidia bacterium UWMA 0267 accepting metabolites.pdf

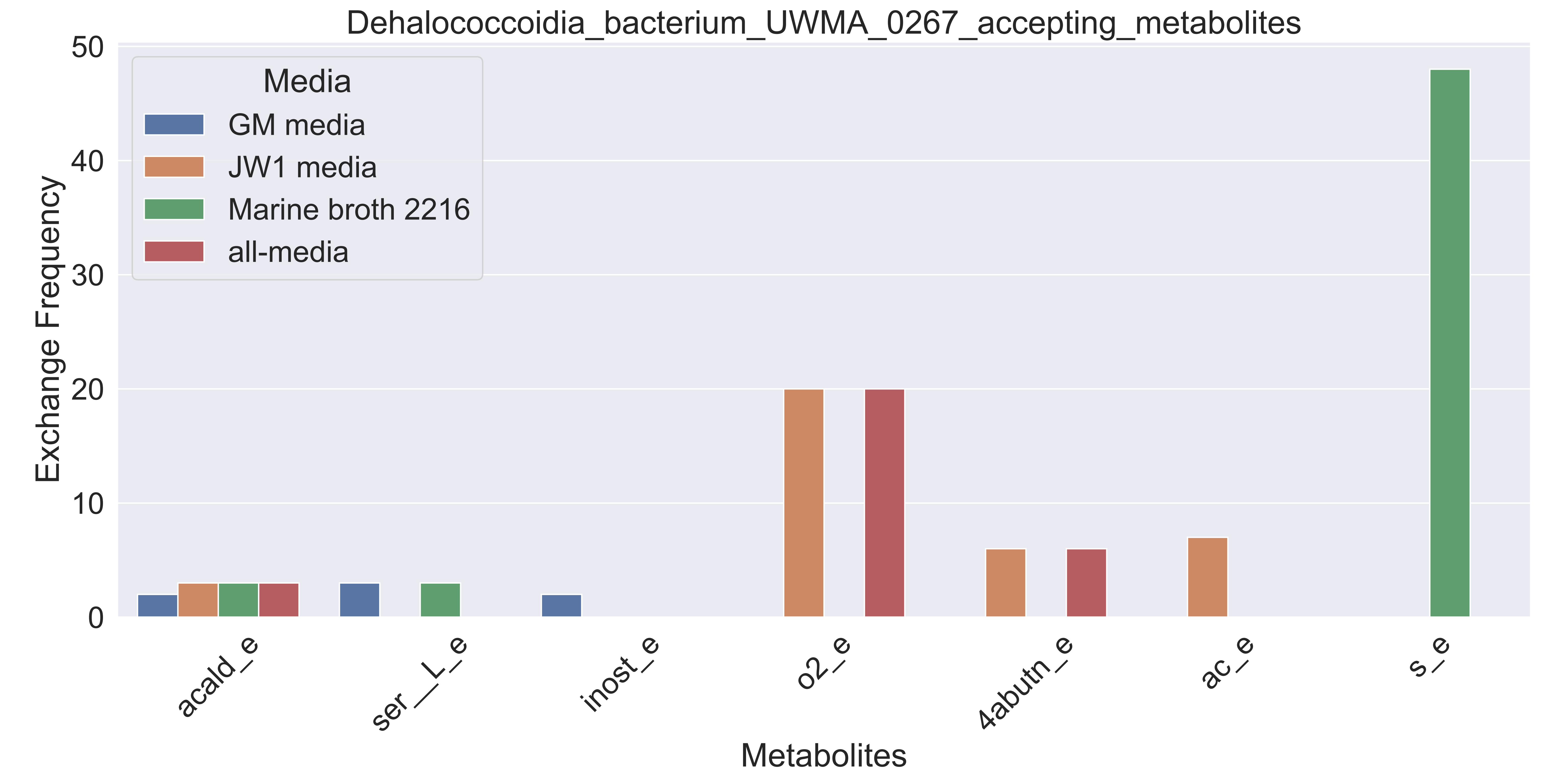

### Supplementary Figure S30 Dehalococcoidia bacterium UWMA 0331 accepting metabolites.pdf

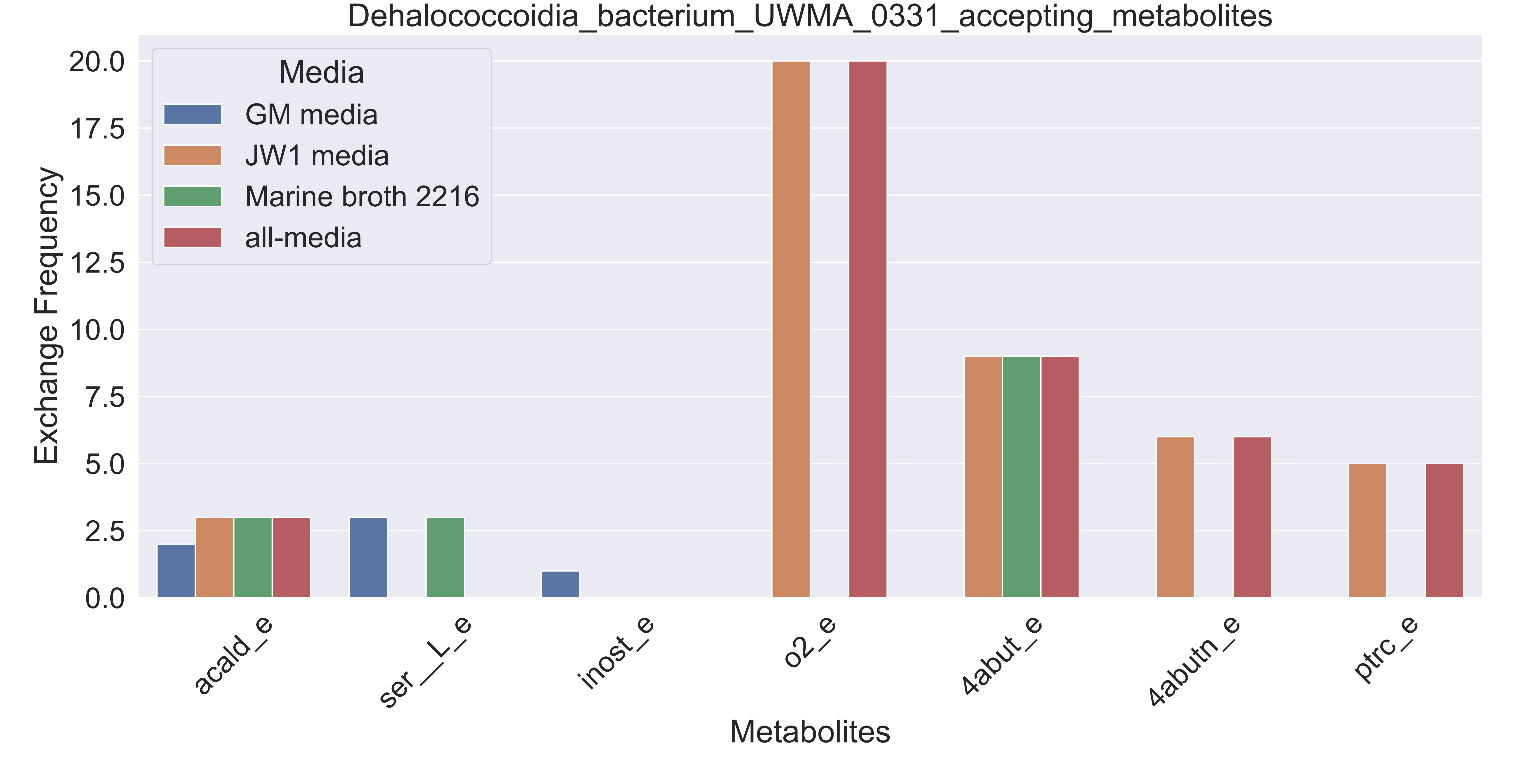

### Supplementary Figure S31 Dehalococcoidia bacterium UWMA 0337 accepting metabolites.pdf

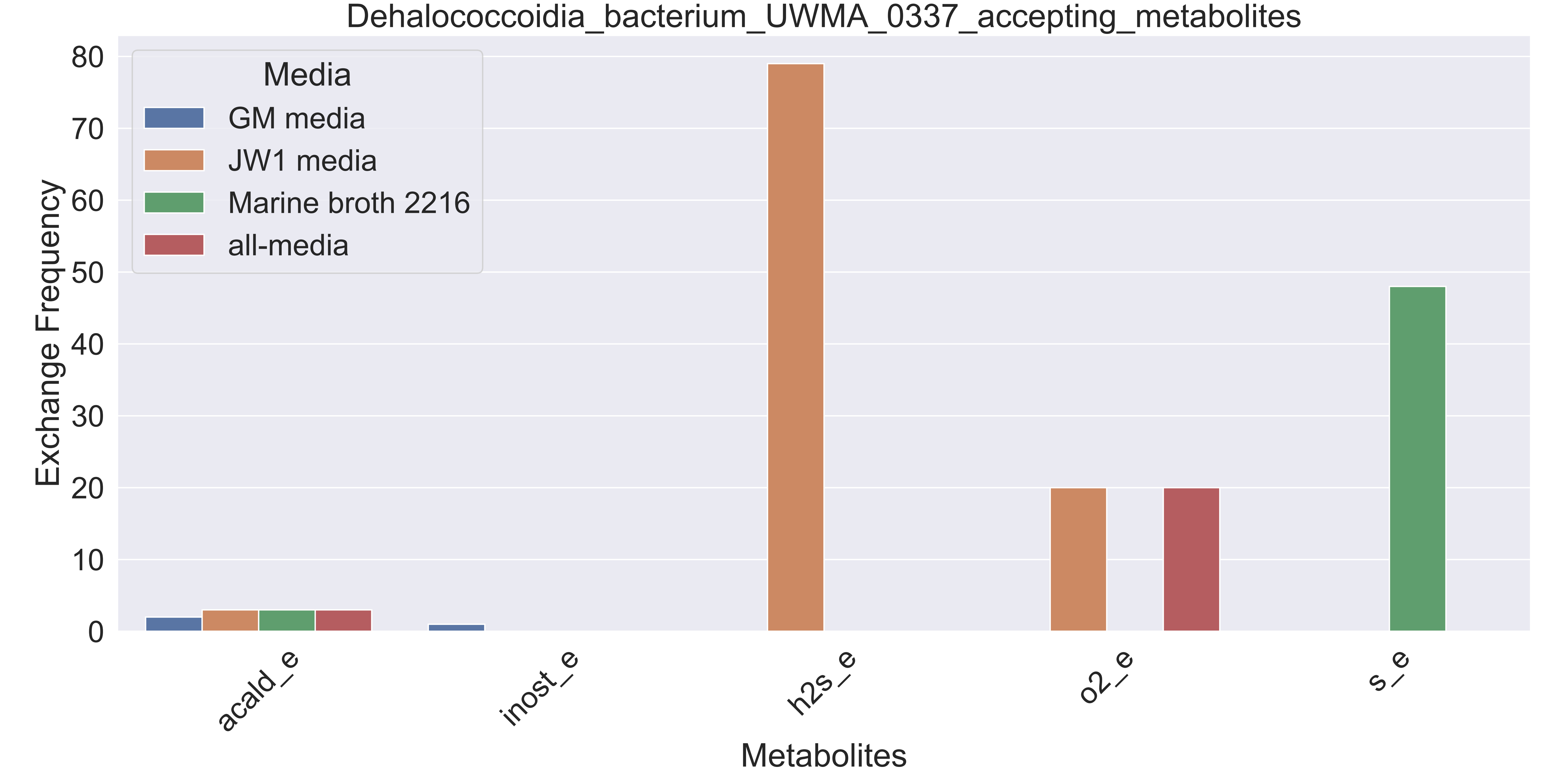

### Supplementary Figure S32 Dehalococcoidia bacterium UWMA 0354 accepting metabolites.pdf

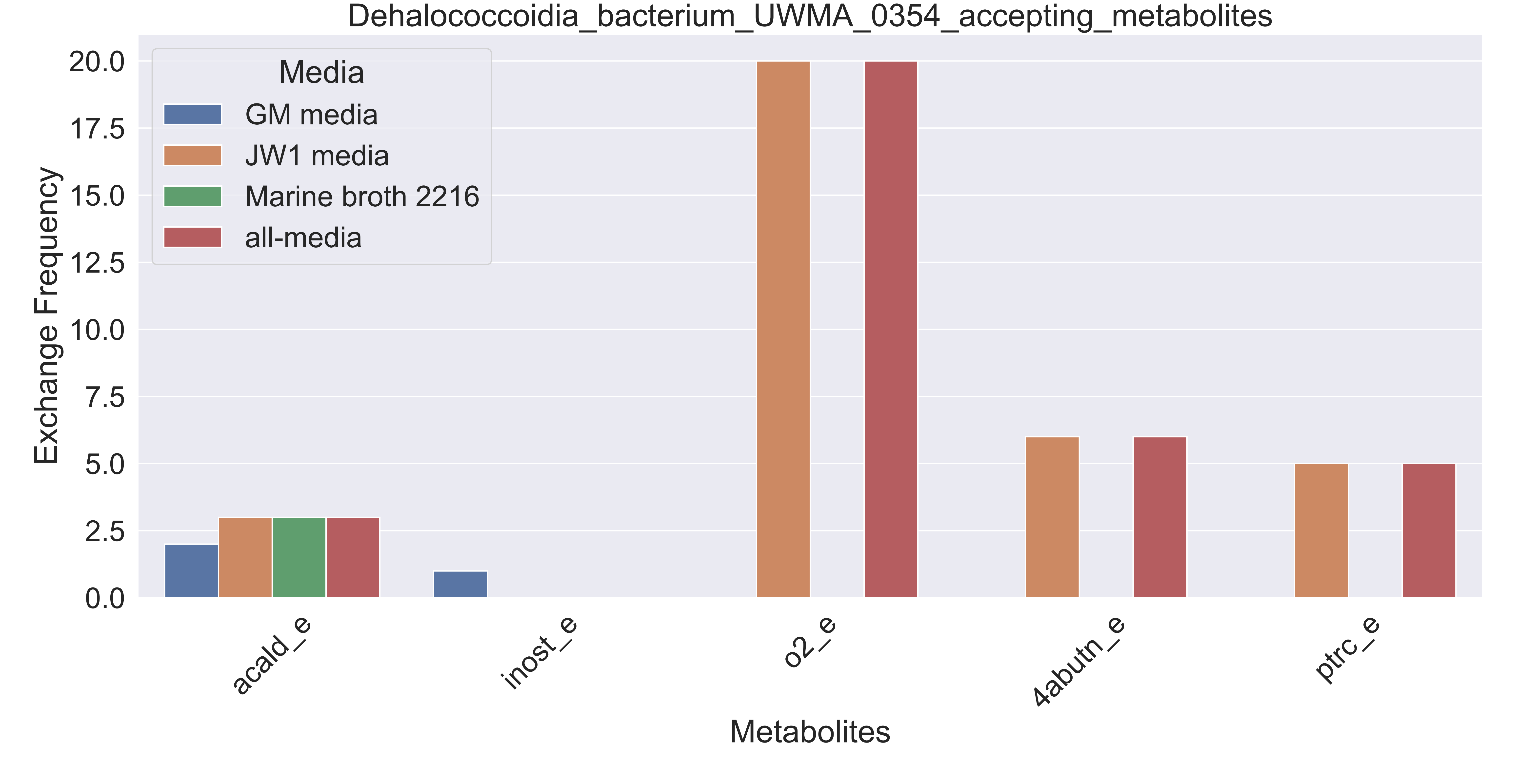

### Supplementary Figure S33 Flavobacteriaceae bacterium UWMA 0314 accepting metabolites.pdf

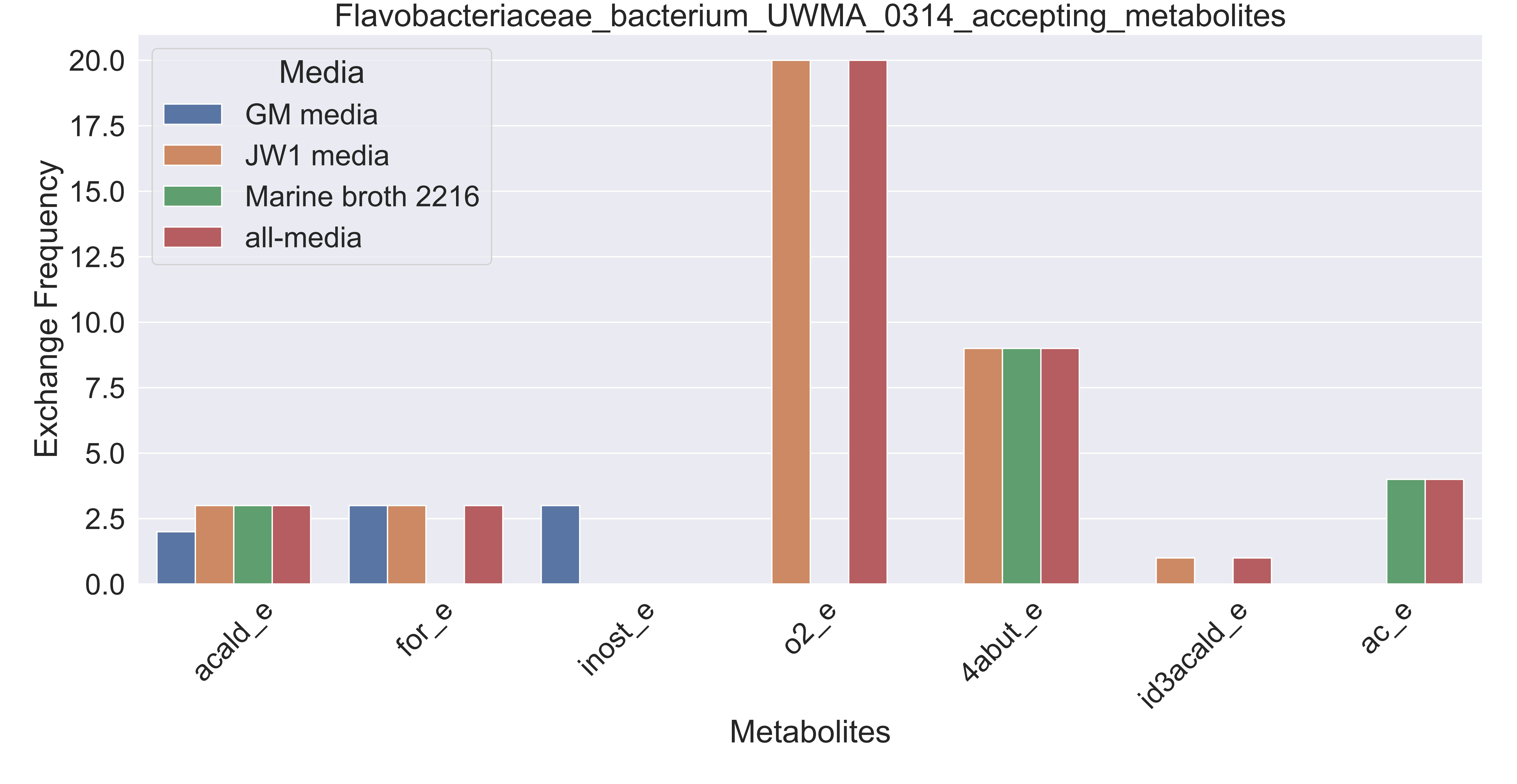

### Supplementary Figure S34 Flavobacteriaceae bacterium UWMA 0351 accepting metabolites.pdf

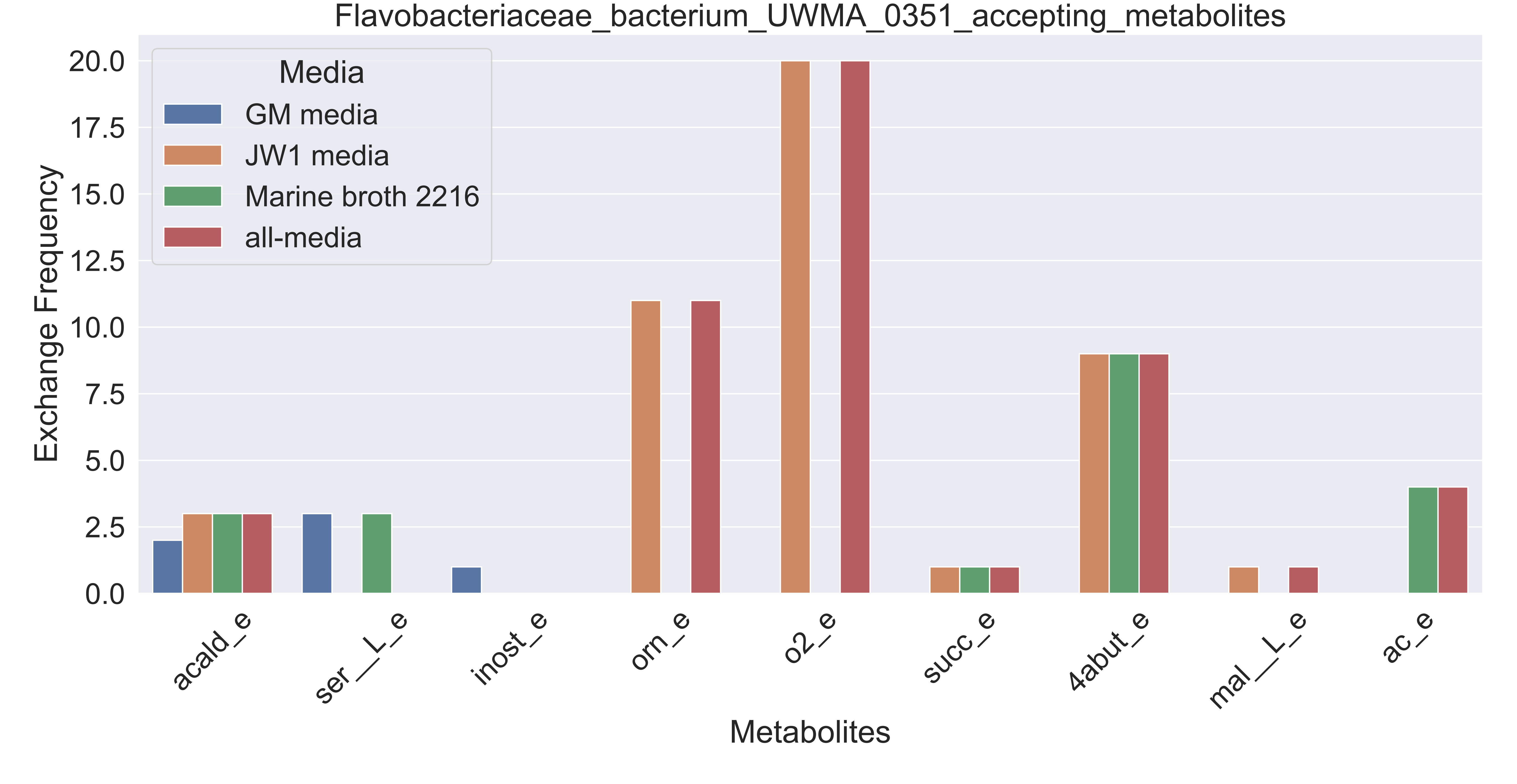

### Supplementary Figure S35 Flavobacteriales bacterium UWMA 0274 accepting metabolites.pdf

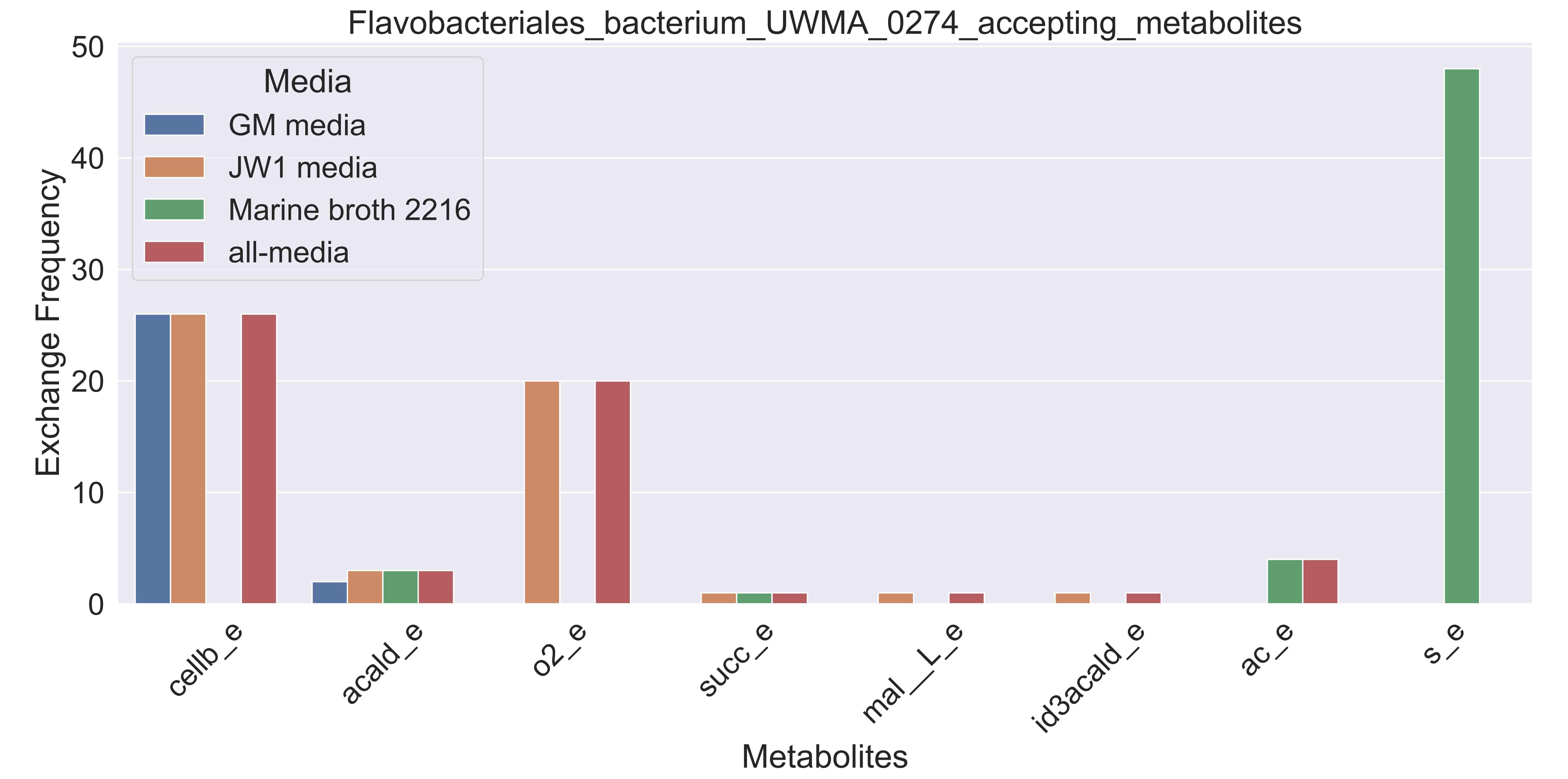

### Supplementary Figure S36 Flavobacteriales bacterium UWMA 0277 accepting metabolites.pdf

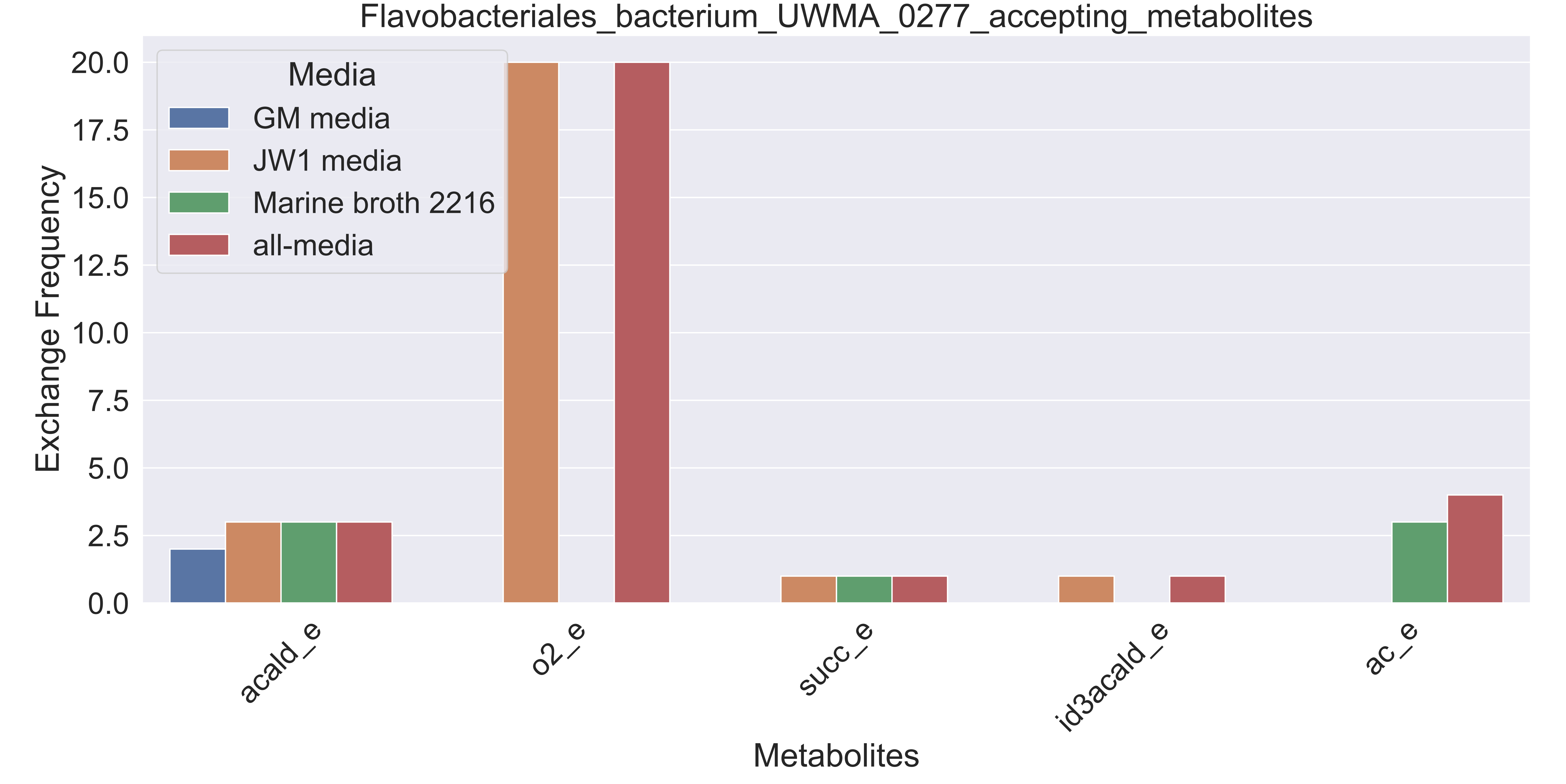

### Supplementary Figure S37 Flavobacteriales bacterium UWMA 0289 accepting metabolites.pdf

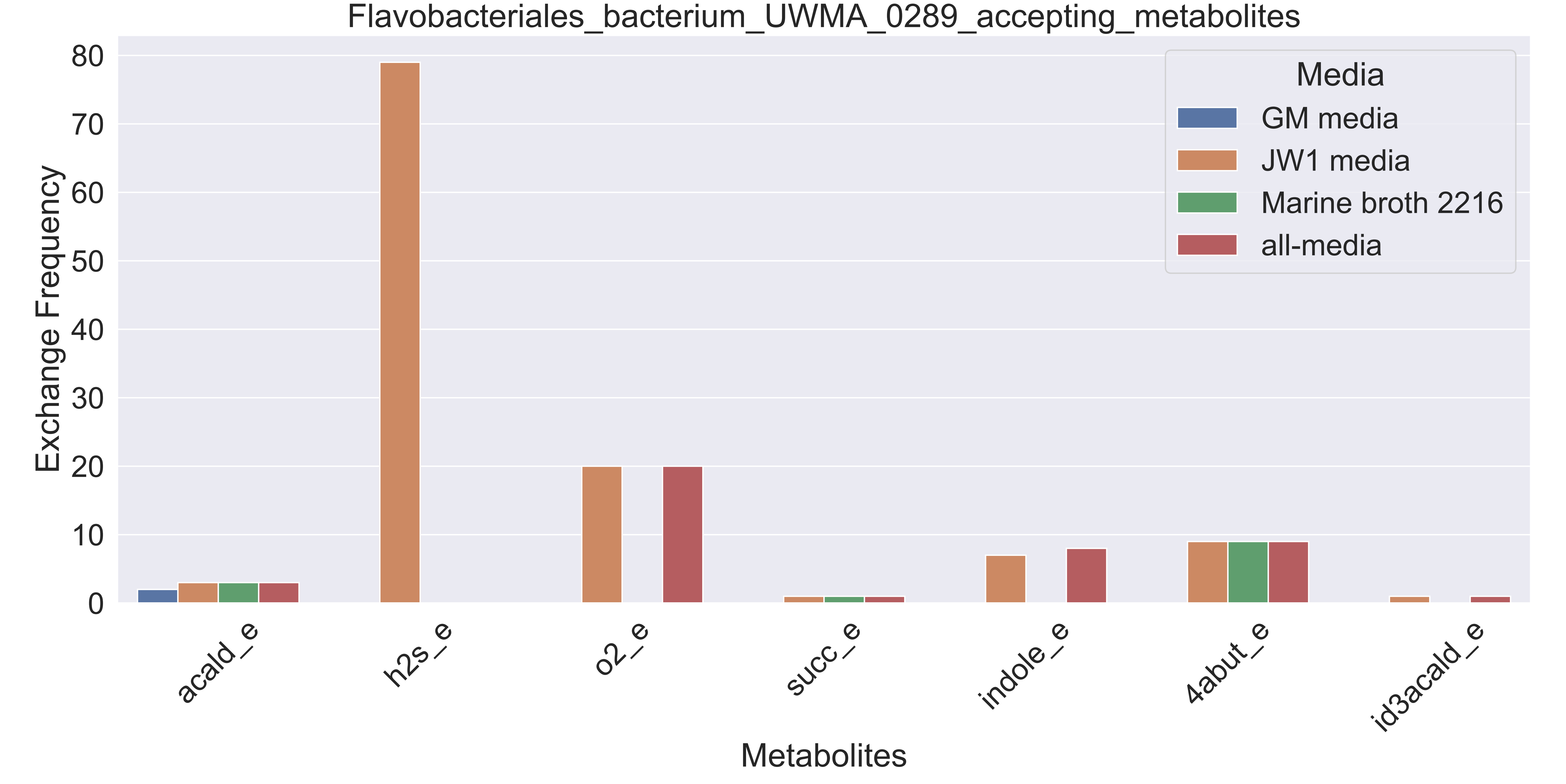

### Supplementary Figure S38 Fuerstia sp UWMA 0333 accepting metabolites.pdf

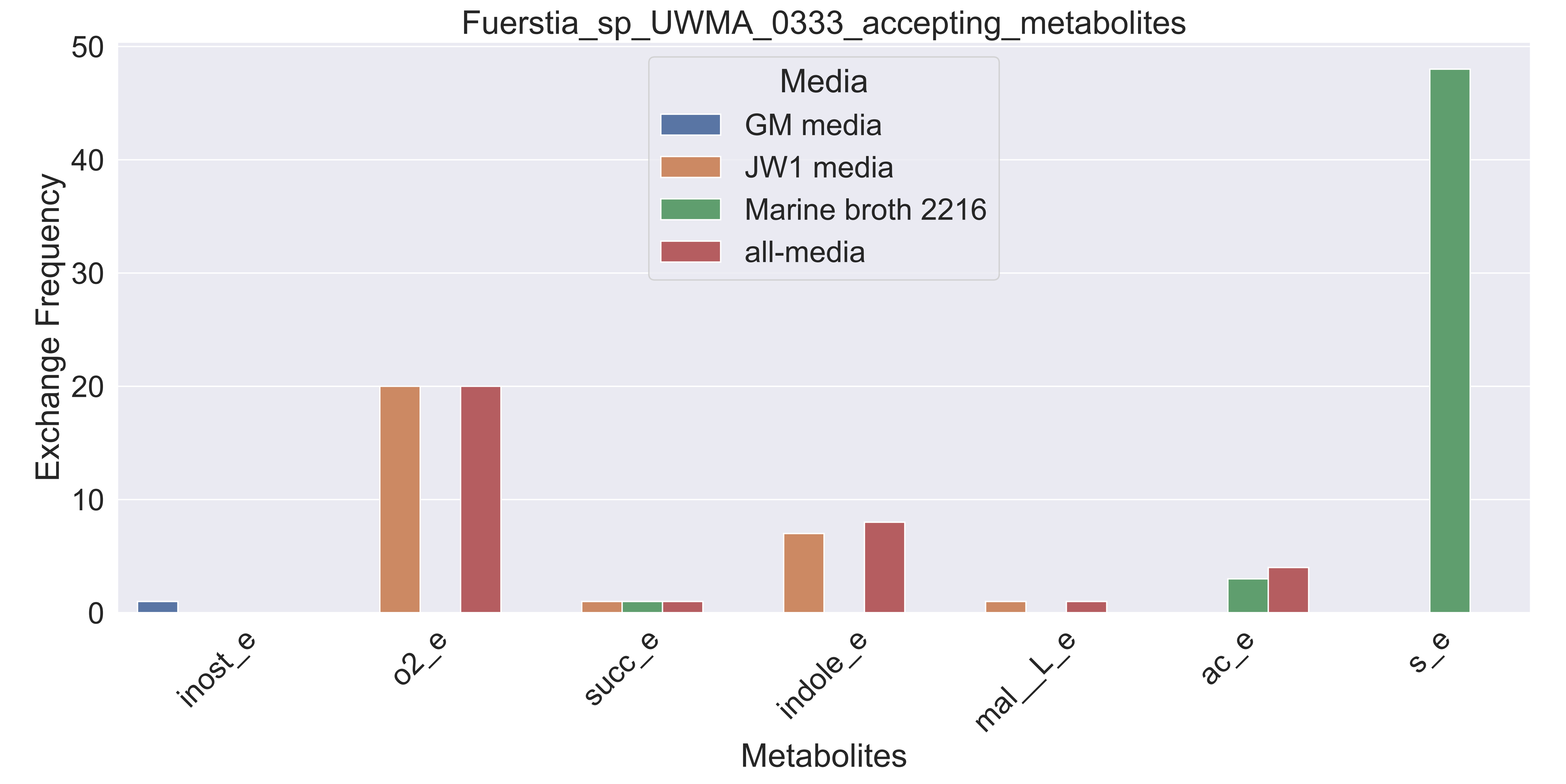

### Supplementary Figure S39 Gammaproteobacteria bacterium UWMA 0258 accepting metabolites.pdf

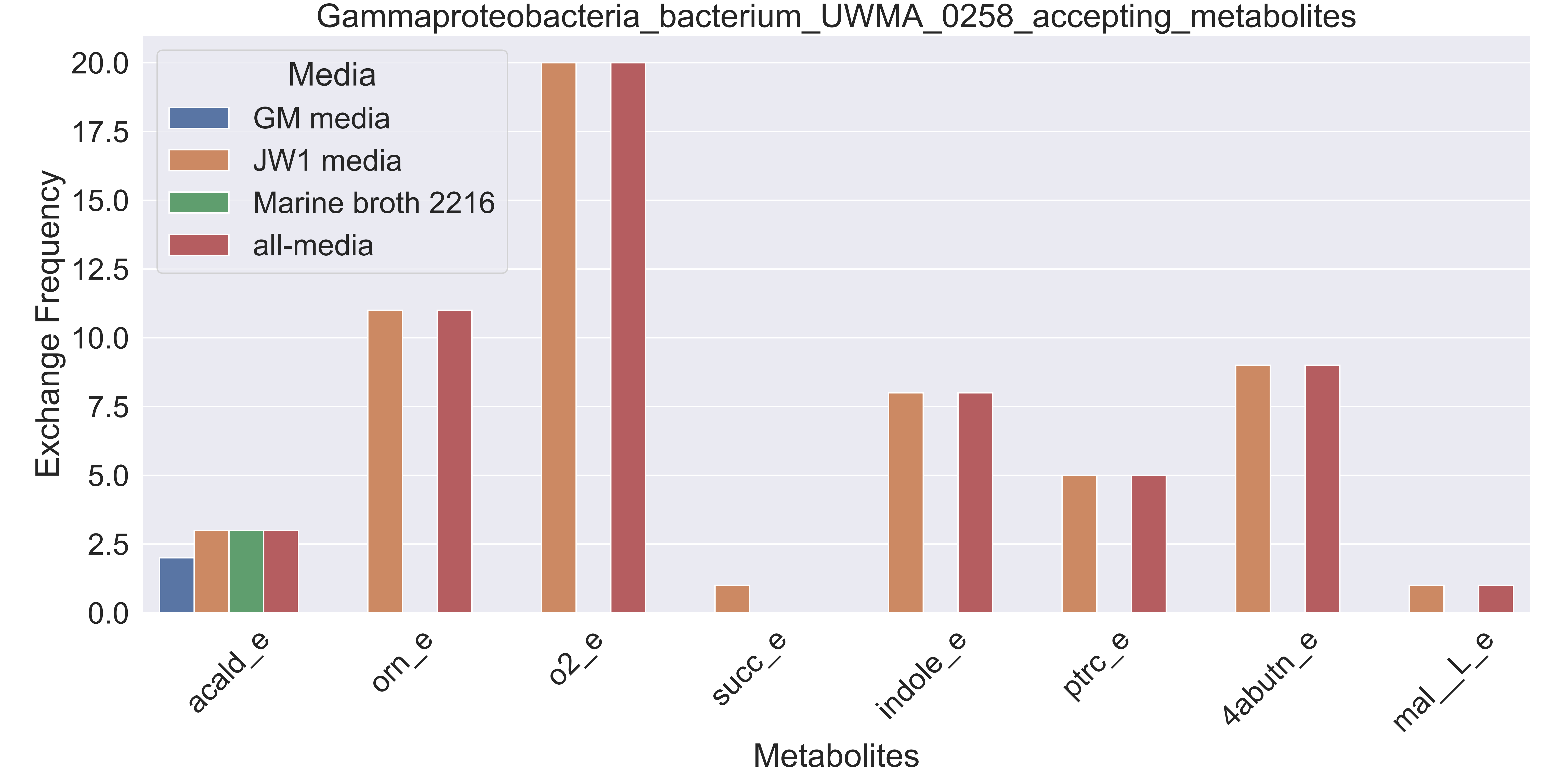

### Supplementary Figure S40 Gammaproteobacteria bacterium UWMA 0260 accepting metabolites.pdf

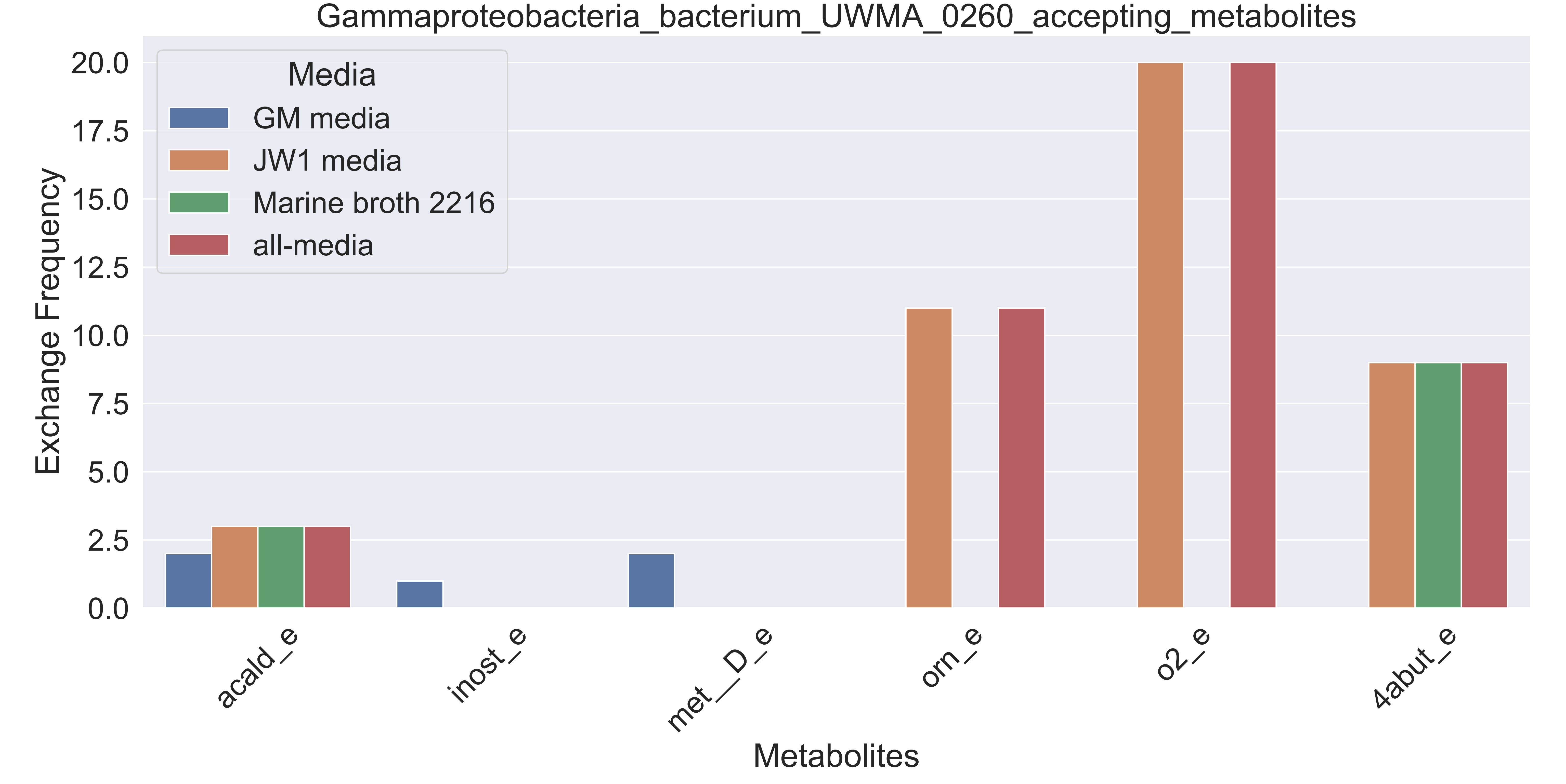

### Supplementary Figure S41 Gammaproteobacteria bacterium UWMA 0261 accepting metabolites.pdf

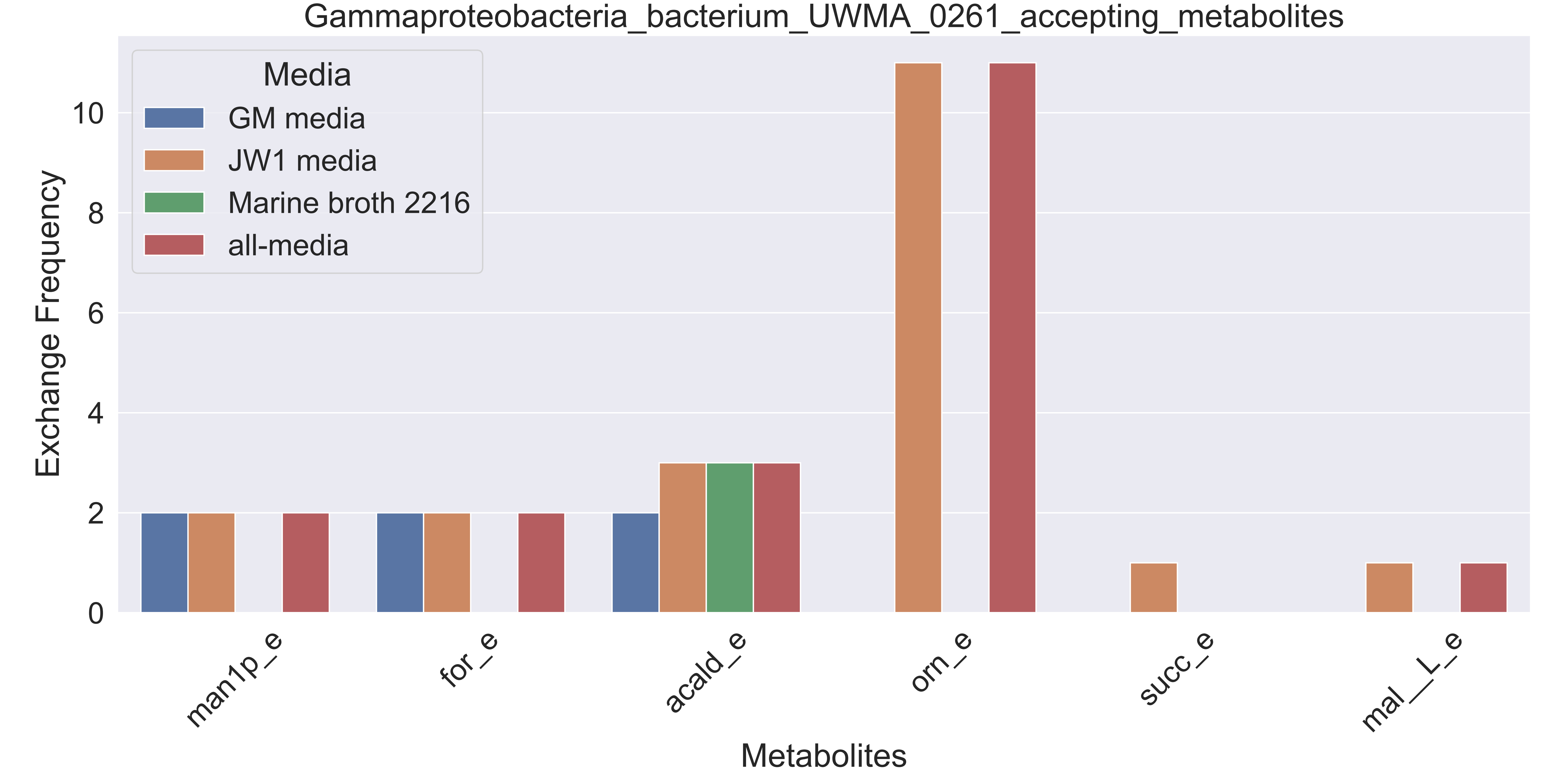

### Supplementary Figure S42 Gammaproteobacteria bacterium UWMA 0273 accepting metabolites.pdf

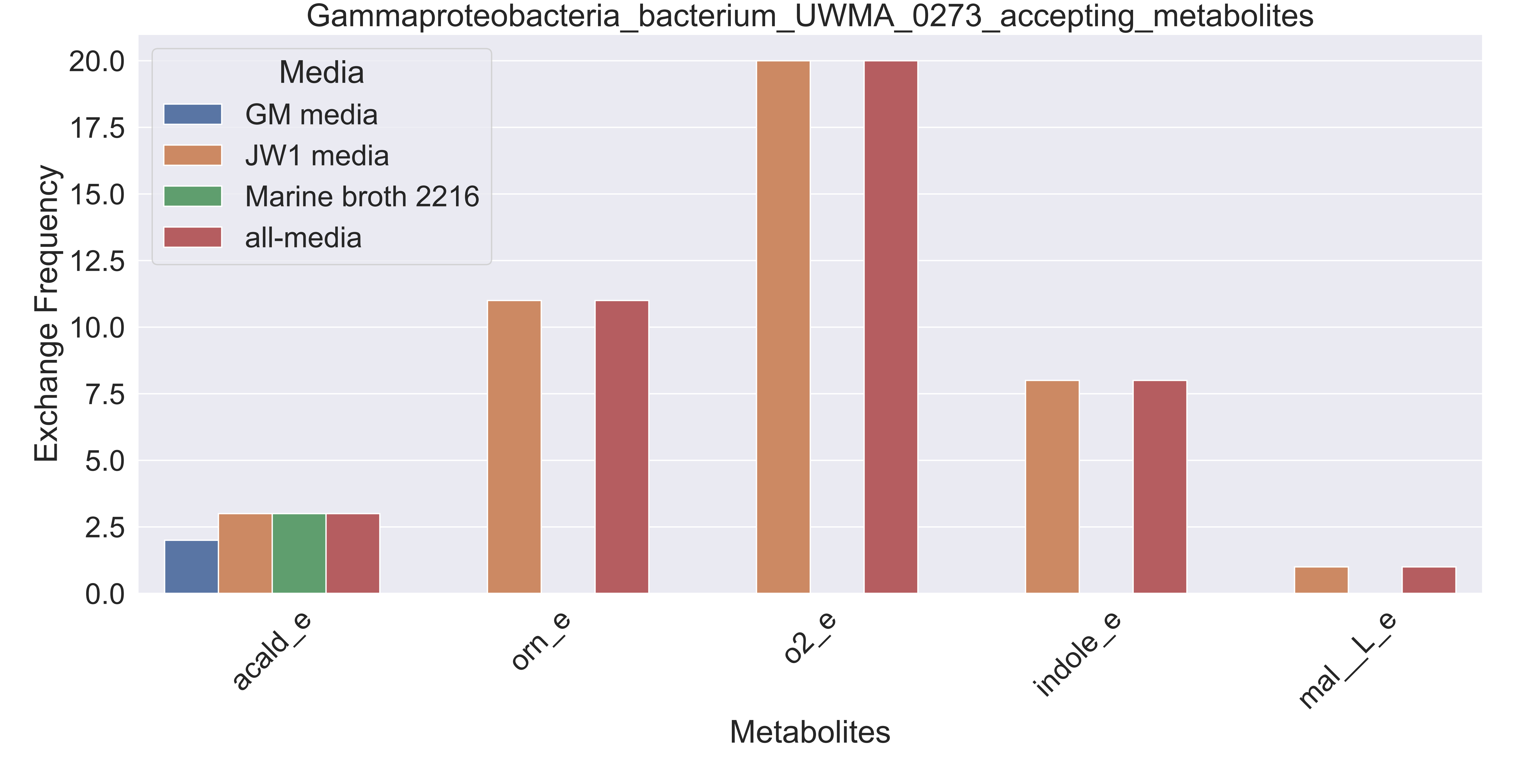

### Supplementary Figure S43 Gammaproteobacteria bacterium UWMA 0278 accepting metabolites.pdf

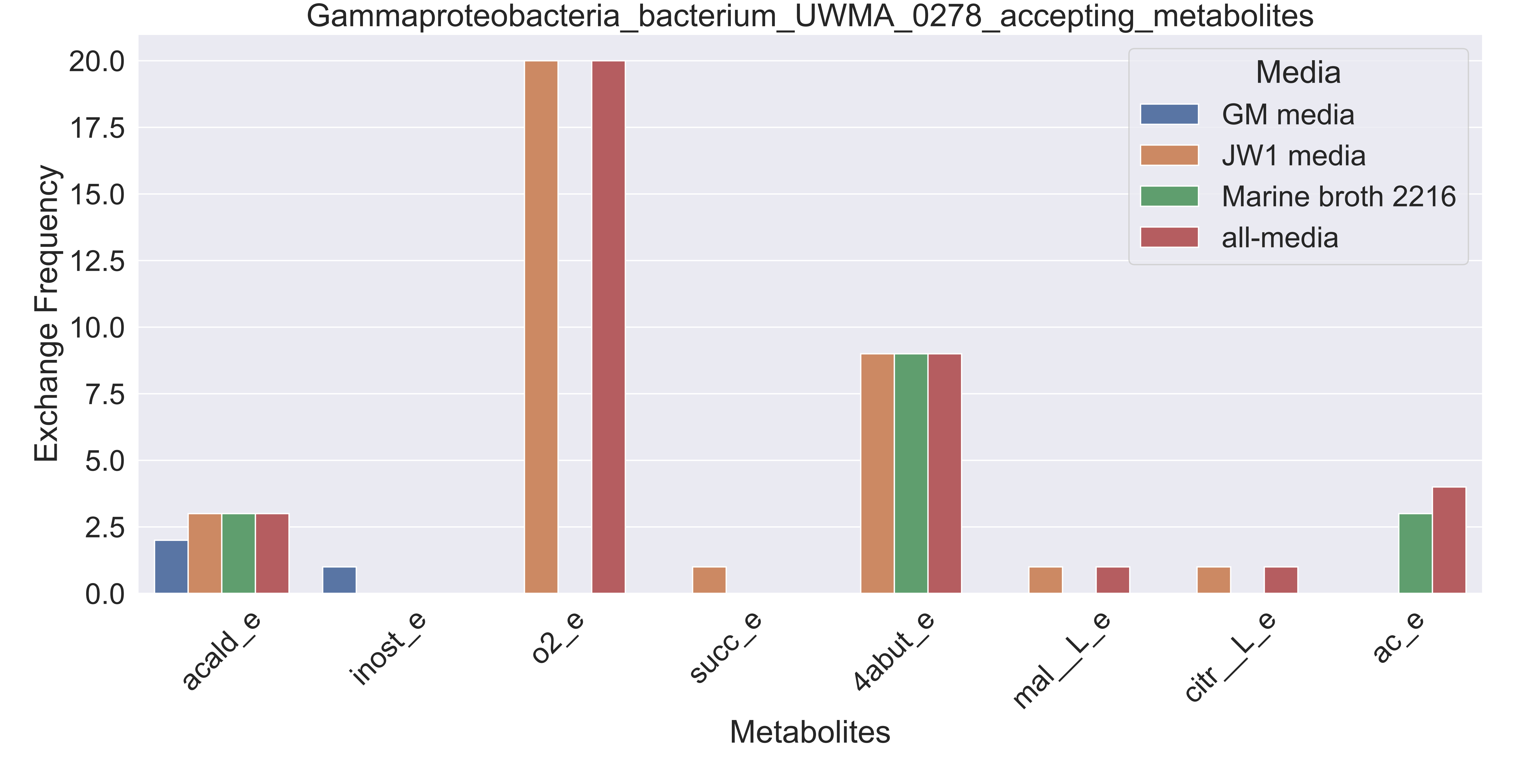

### Supplementary Figure S44 Gammaproteobacteria bacterium UWMA 0281 accepting metabolites.pdf

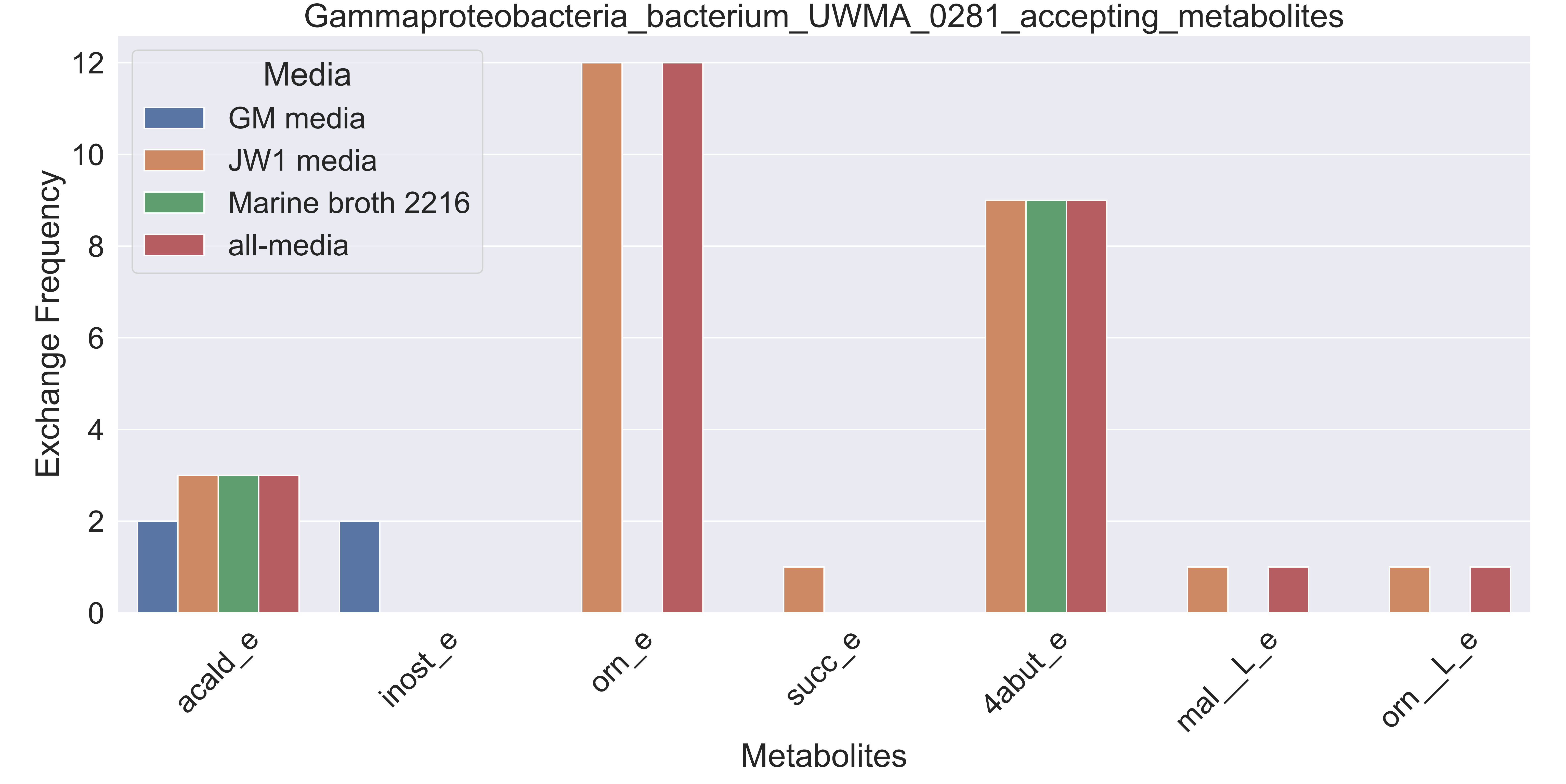

### Supplementary Figure S45 Gammaproteobacteria bacterium UWMA 0282 accepting metabolites.pdf

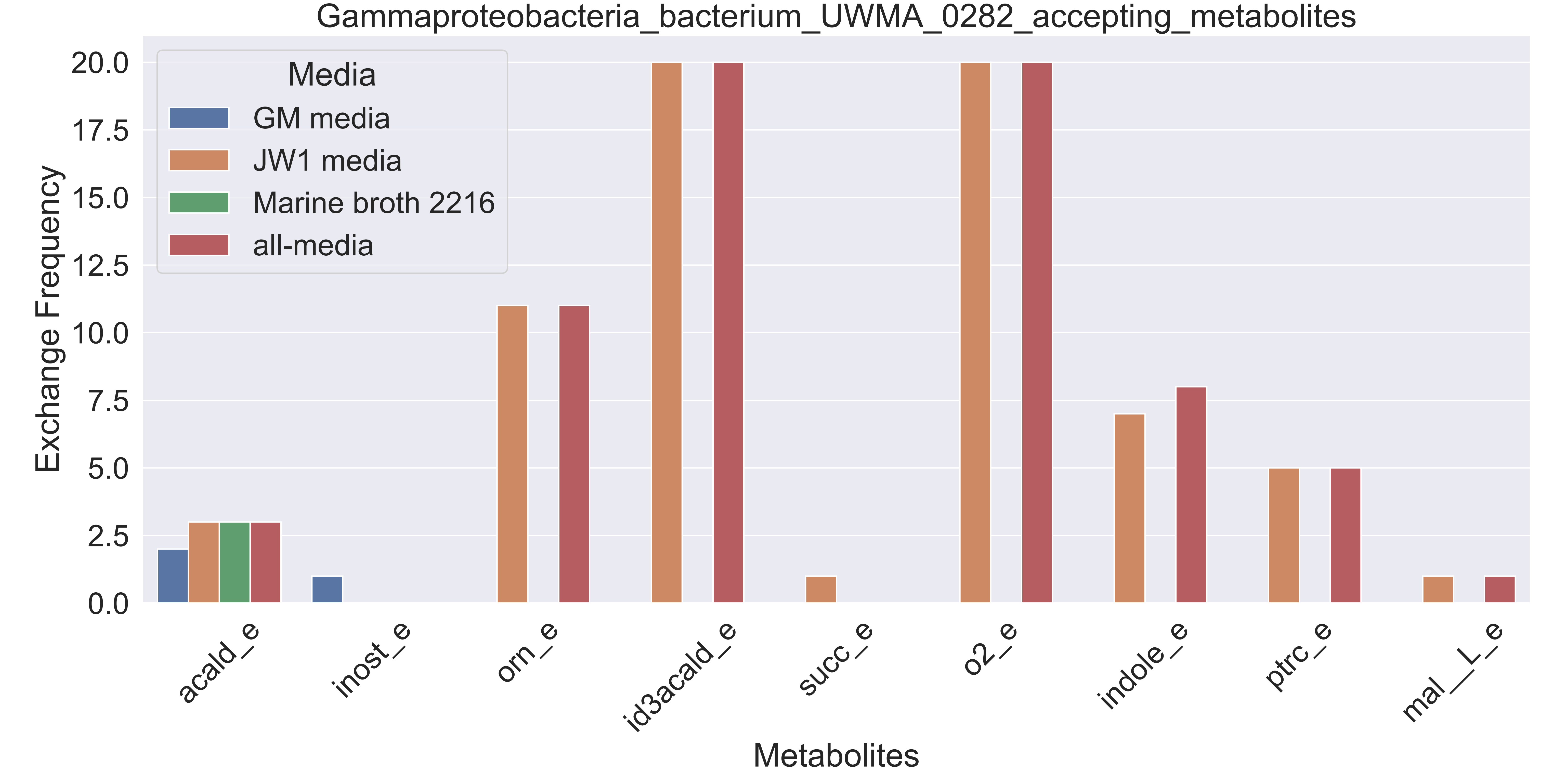

### Supplementary Figure S46 Gammaproteobacteria bacterium UWMA 0291 accepting metabolites.pdf

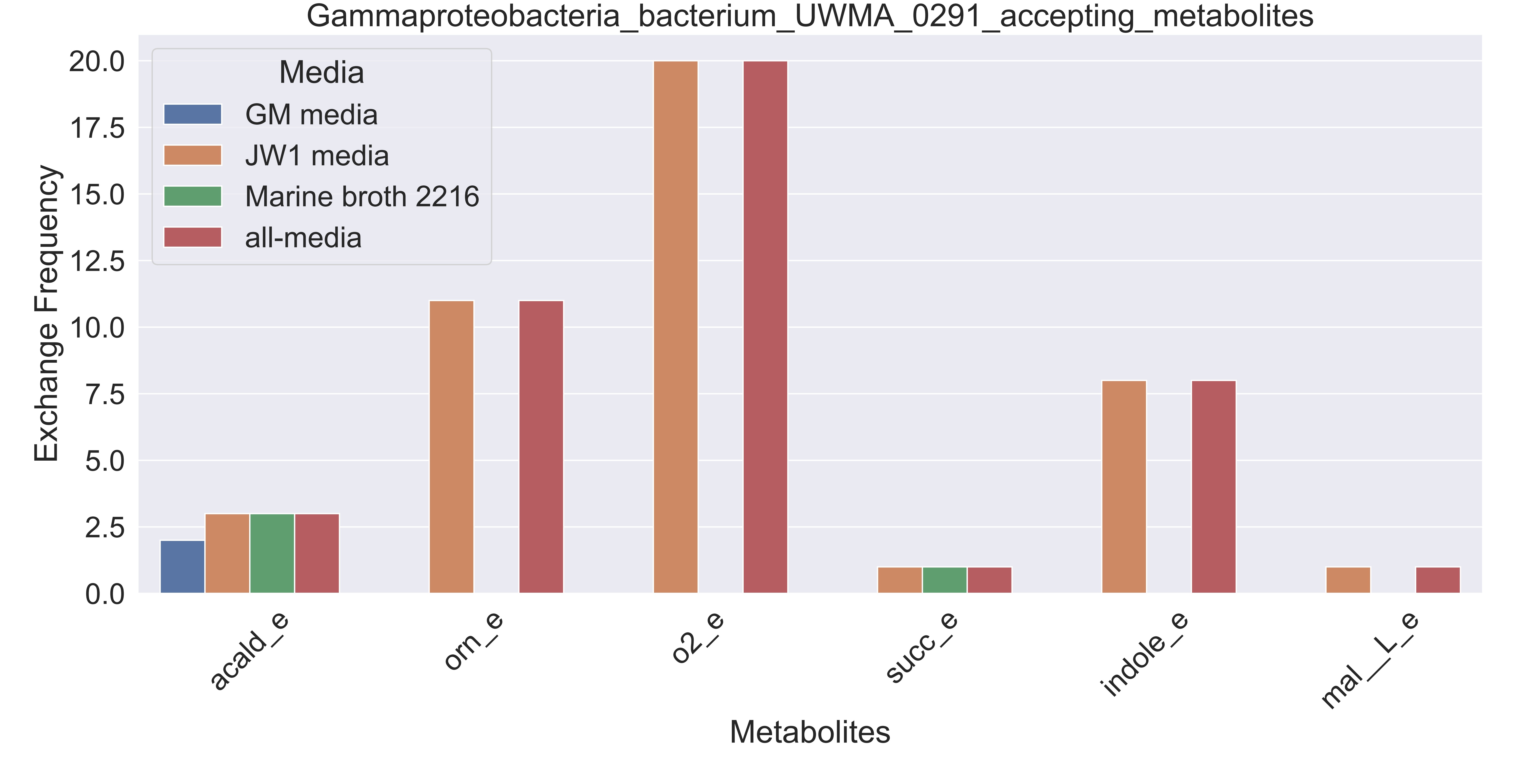

### Supplementary Figure S47 Gammaproteobacteria bacterium UWMA 0299 accepting metabolites.pdf

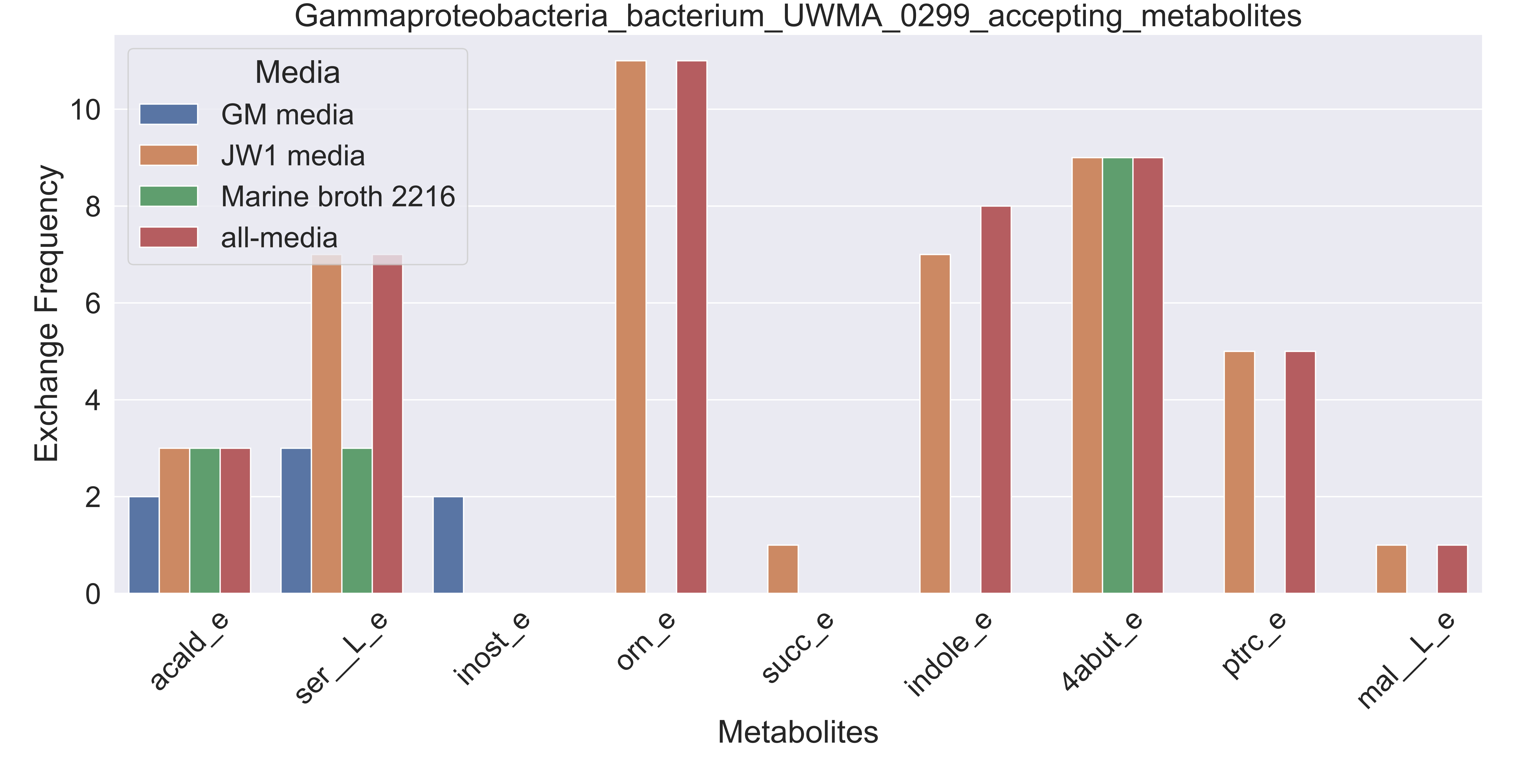

### Supplementary Figure S48 Gammaproteobacteria bacterium UWMA 0301 accepting metabolites.pdf

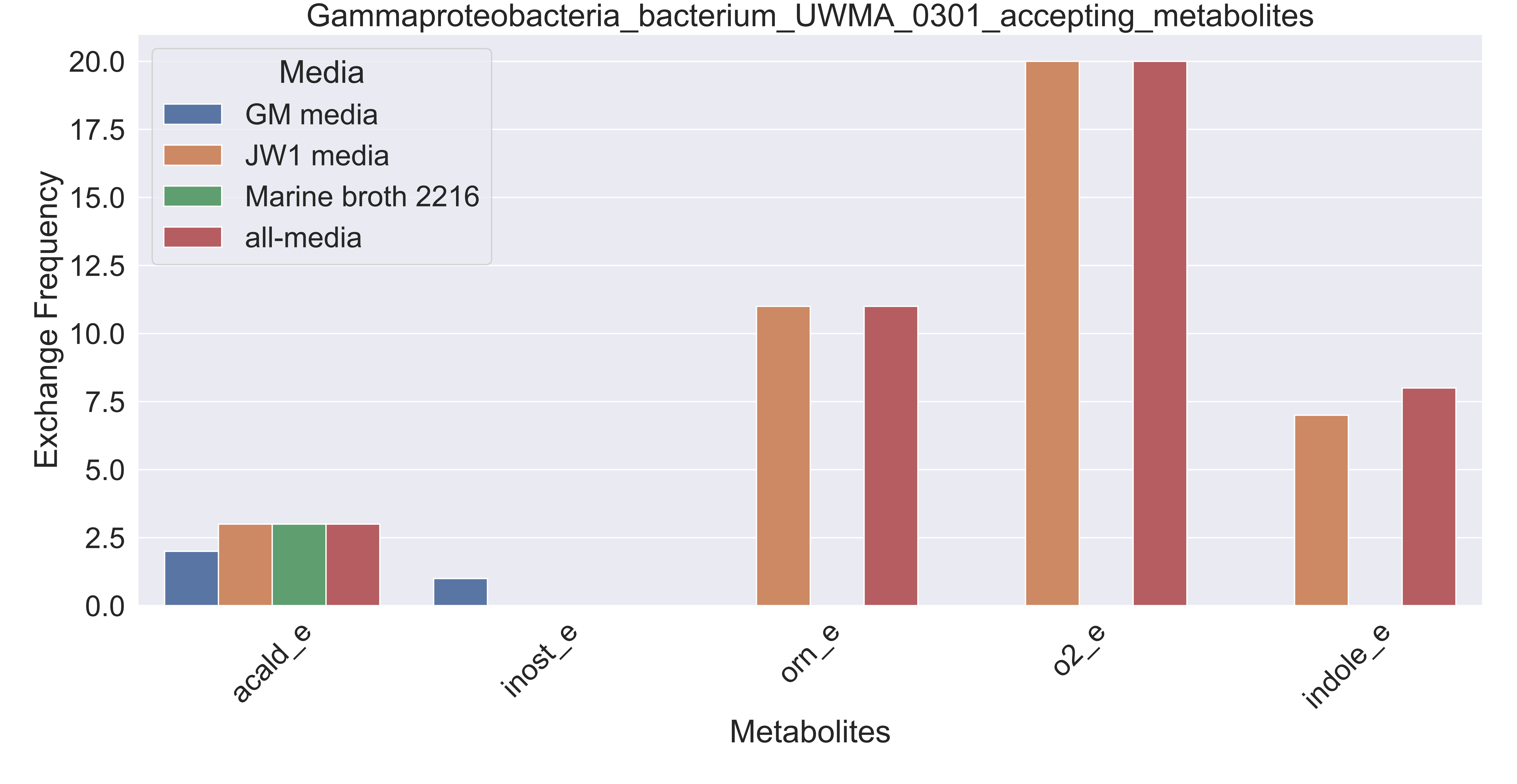

### Supplementary Figure S49 Gammaproteobacteria bacterium UWMA 0302 accepting metabolites.pdf

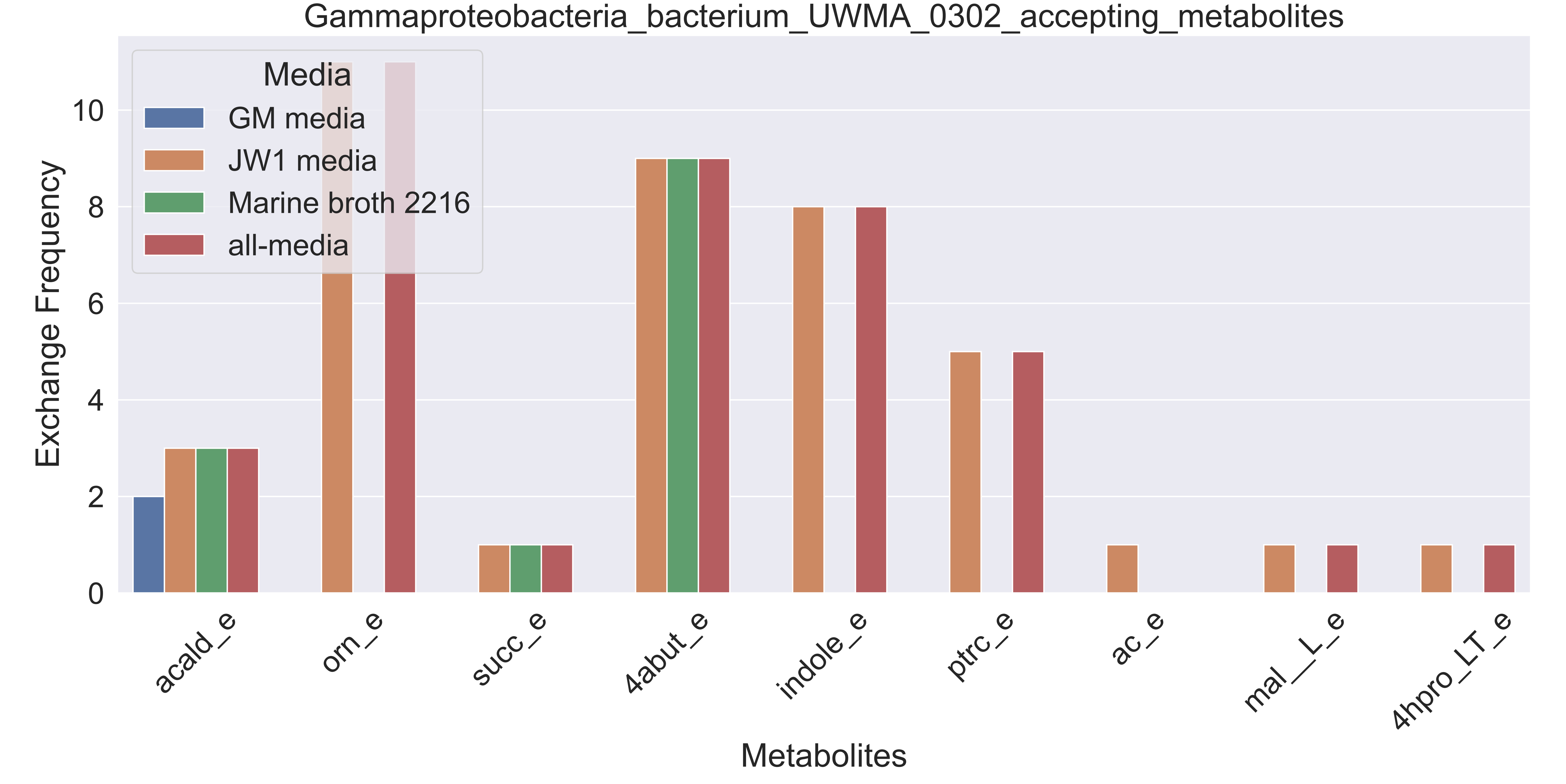

### Supplementary Figure S50 Gammaproteobacteria bacterium UWMA 0343 accepting metabolites.pdf

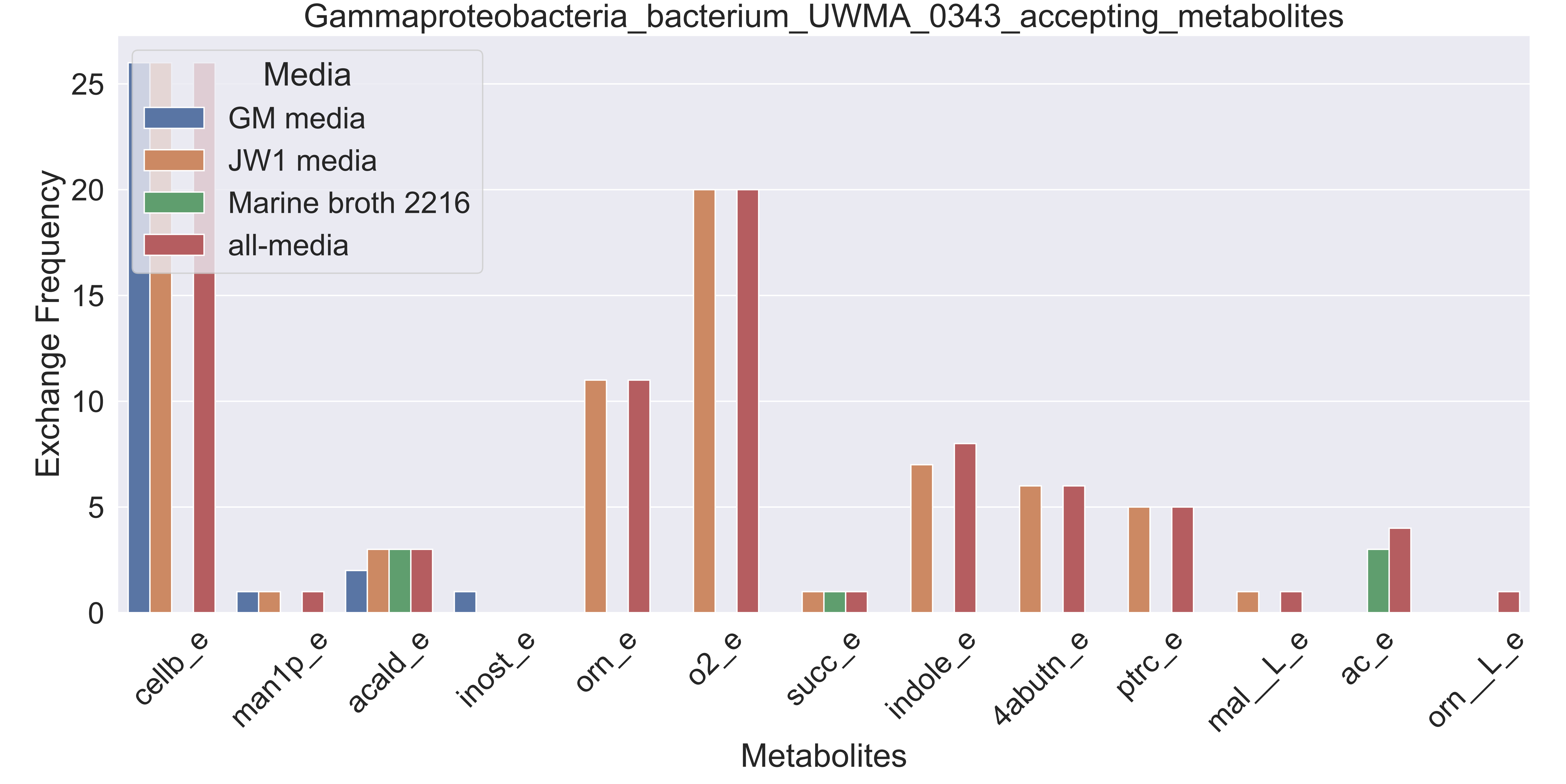

### Supplementary Figure S51 Gemmatimonadetes bacterium UWMA 0296 accepting metabolites.pdf

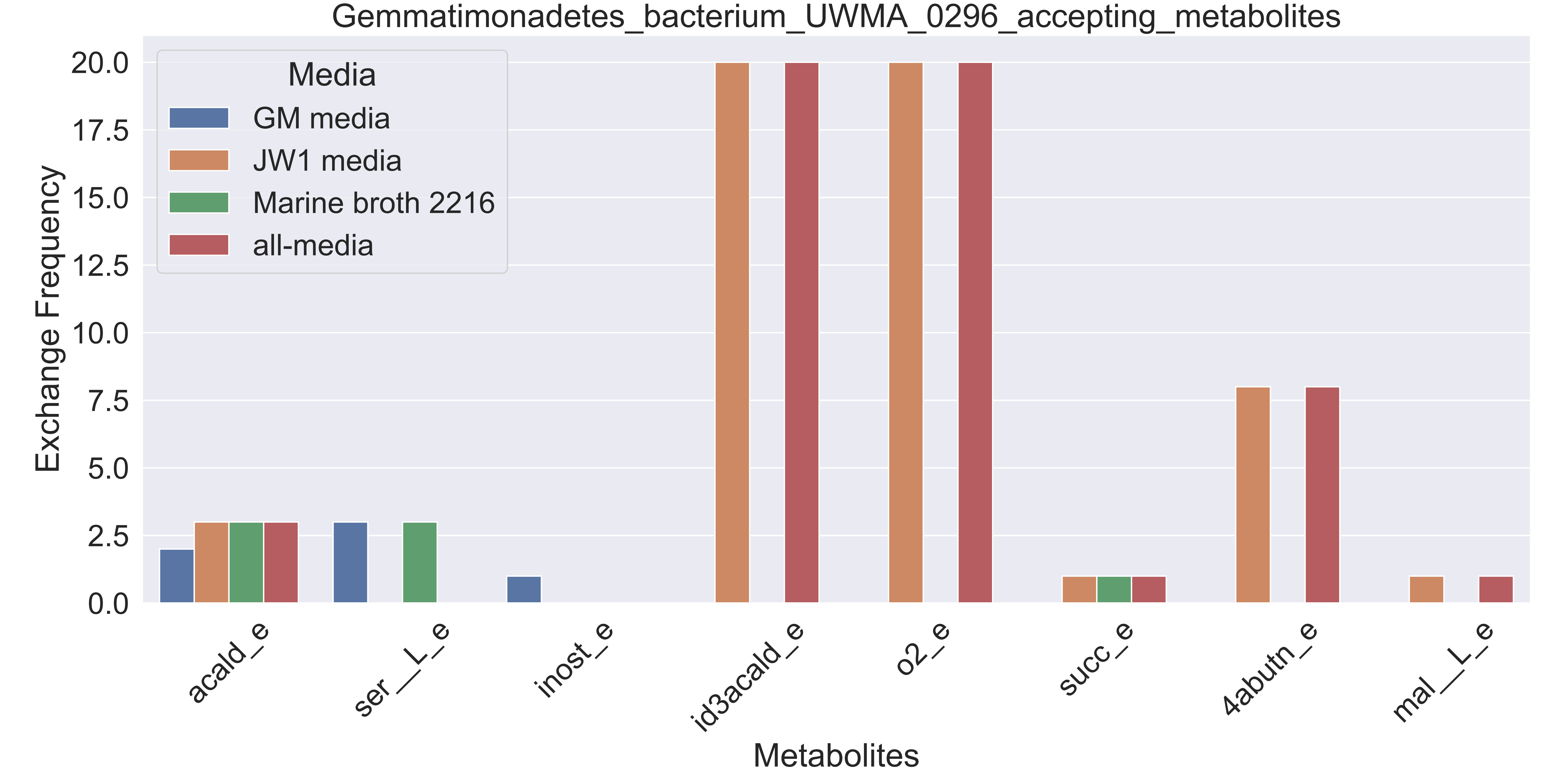

### Supplementary Figure S52 Gemmatimonadetes bacterium UWMA 0308 accepting metabolites.pdf

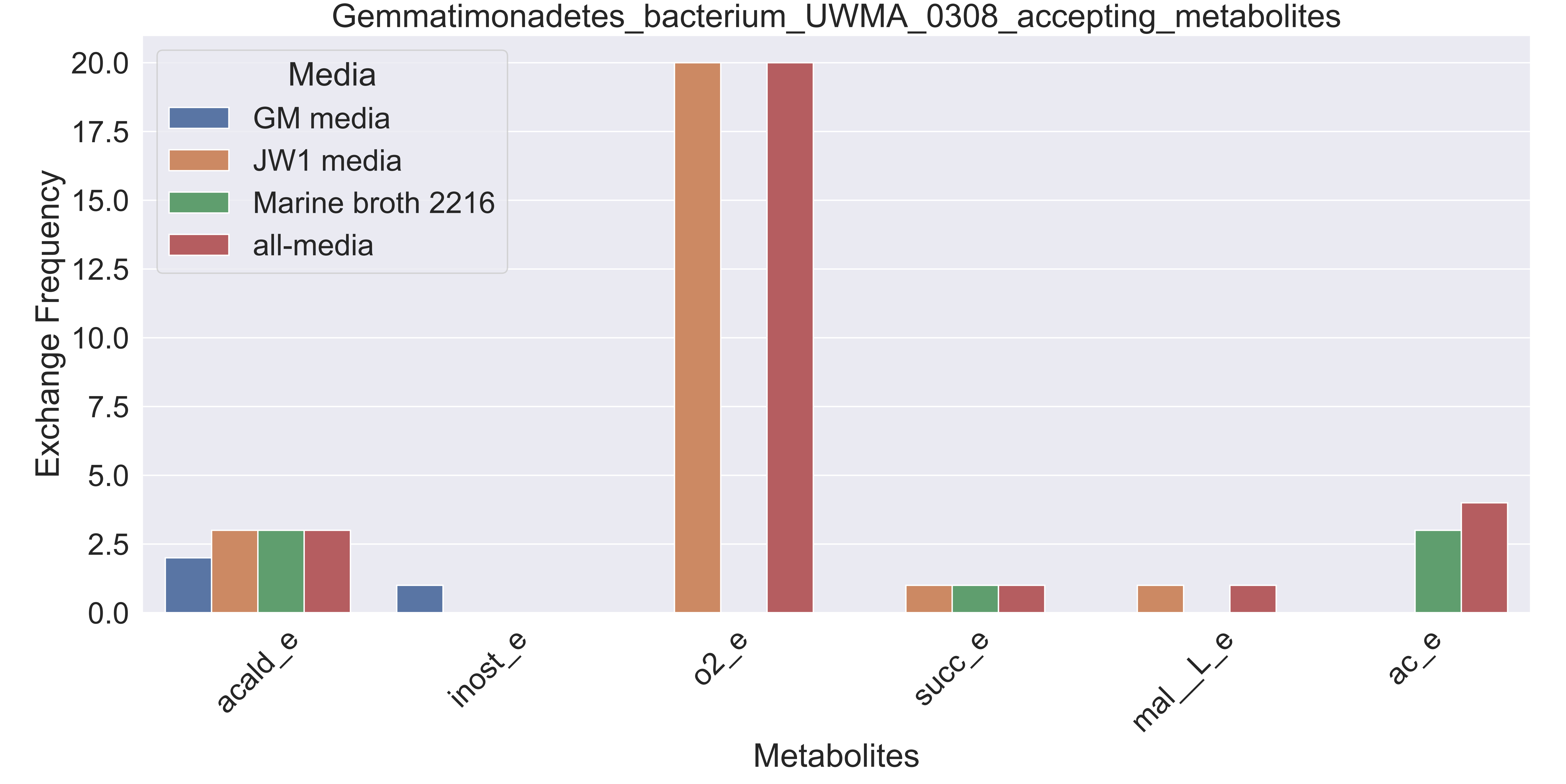

### Supplementary Figure S53 Gemmatimonadetes bacterium UWMA 0334 accepting metabolites.pdf

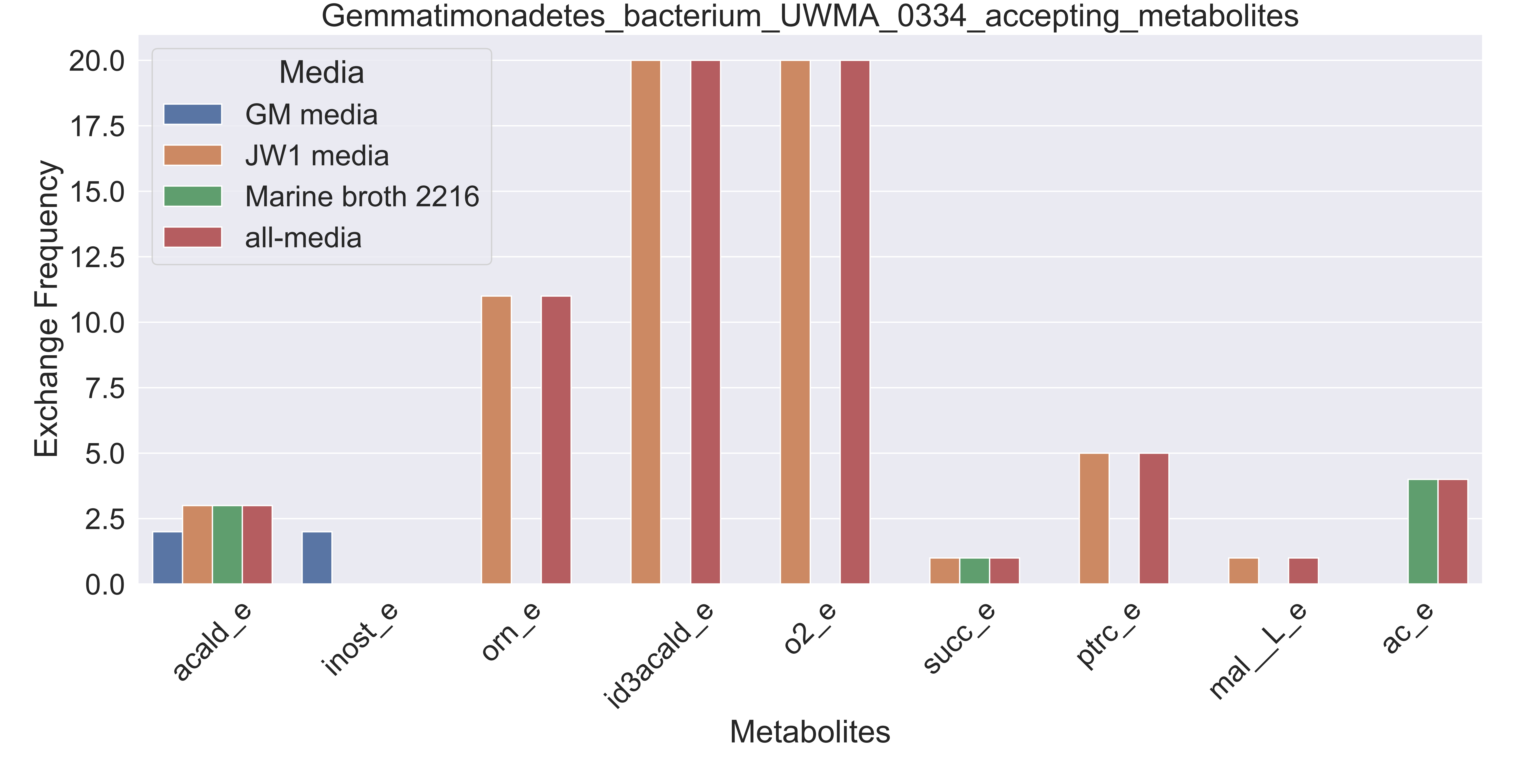

### Supplementary Figure S54 Gemmatimonadetes bacterium UWMA 0339 accepting metabolites.pdf

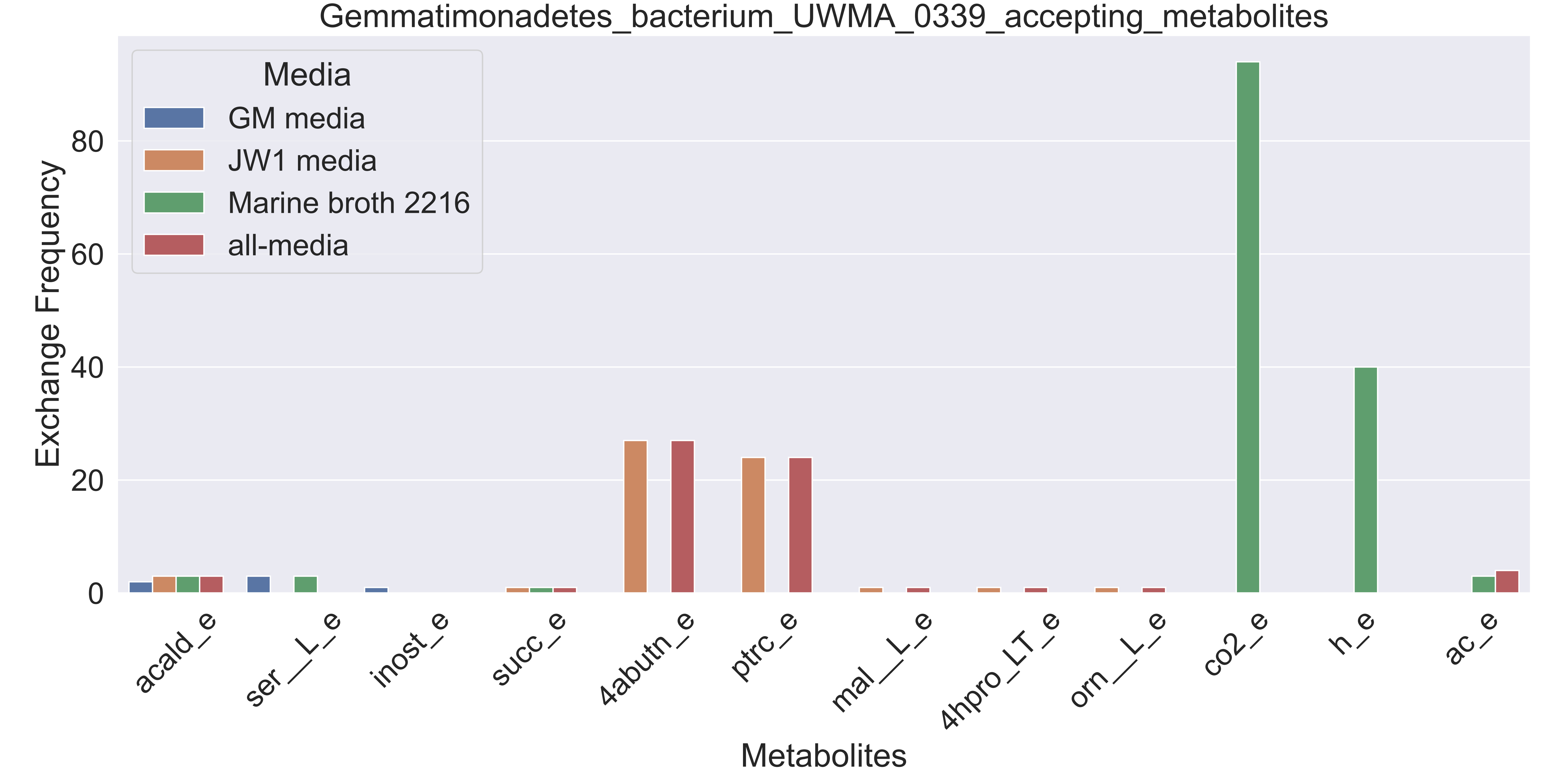

### Supplementary Figure S55 Henriciella sp UWMA 0297 accepting metabolites.pdf

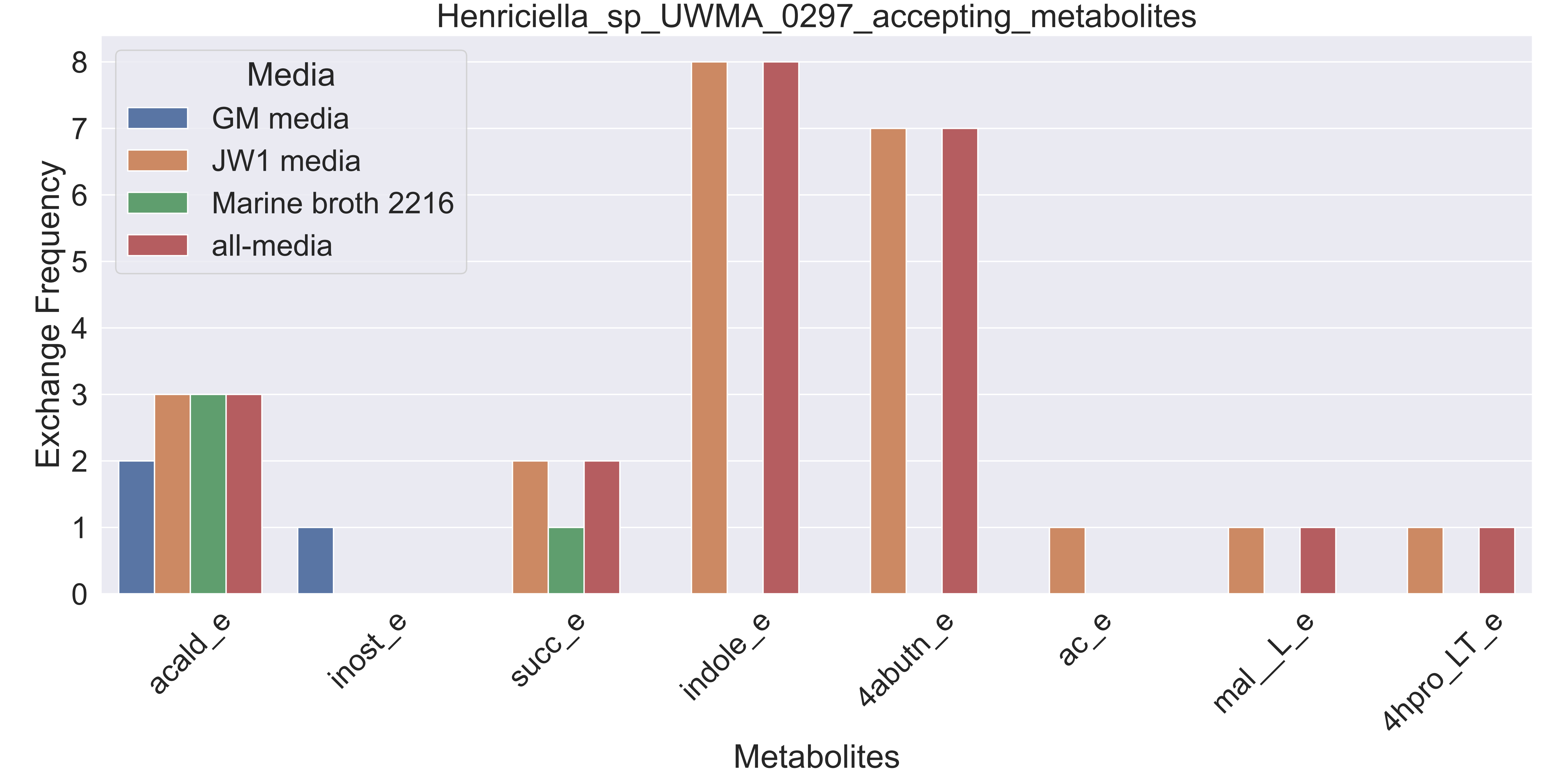

### Supplementary Figure S56 Marine Group III euryarchaeote UWMA 0284 accepting metabolites.pdf

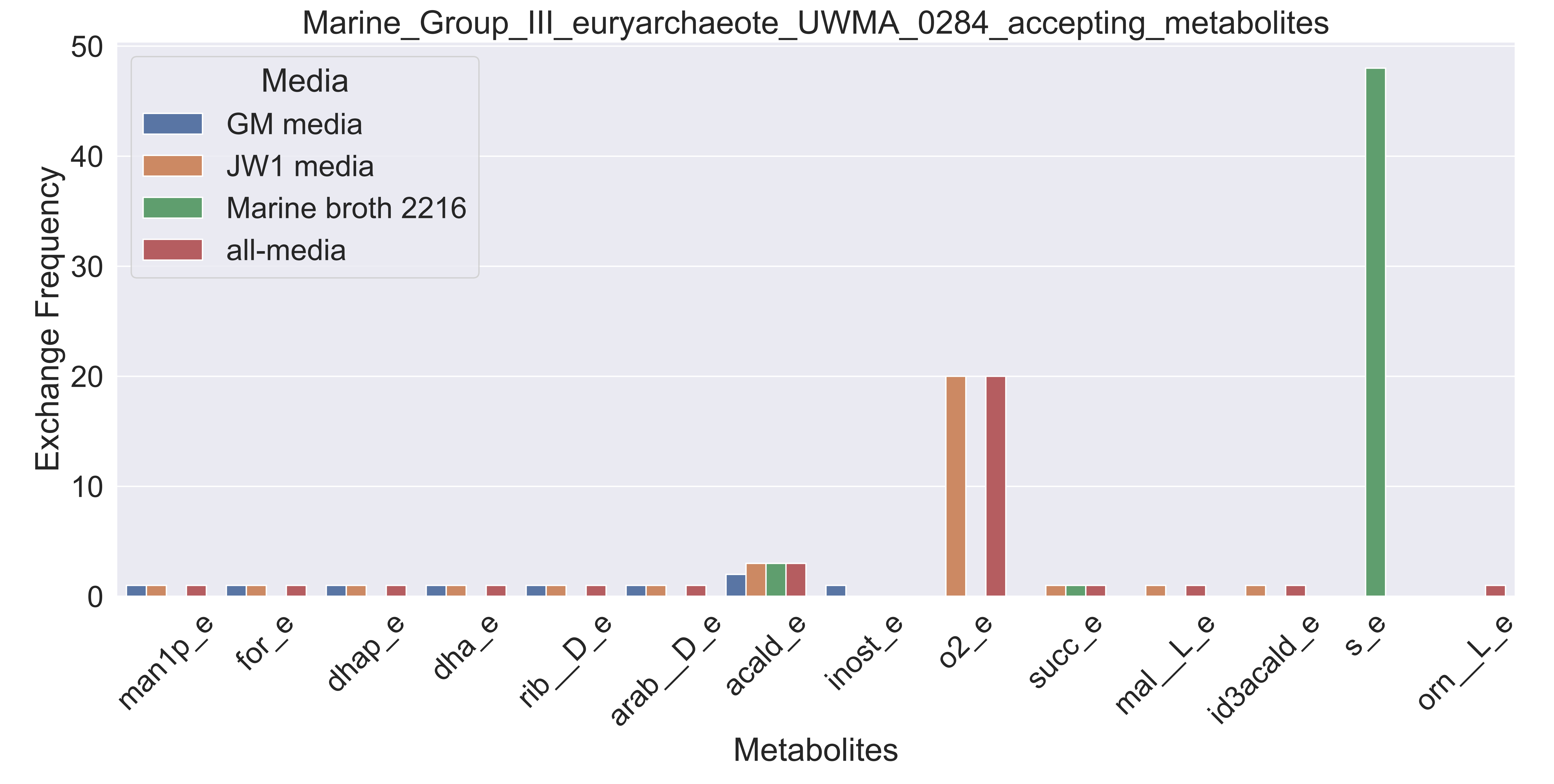

### Supplementary Figure S57 Marine Group III euryarchaeote UWMA 0340 accepting metabolites.pdf

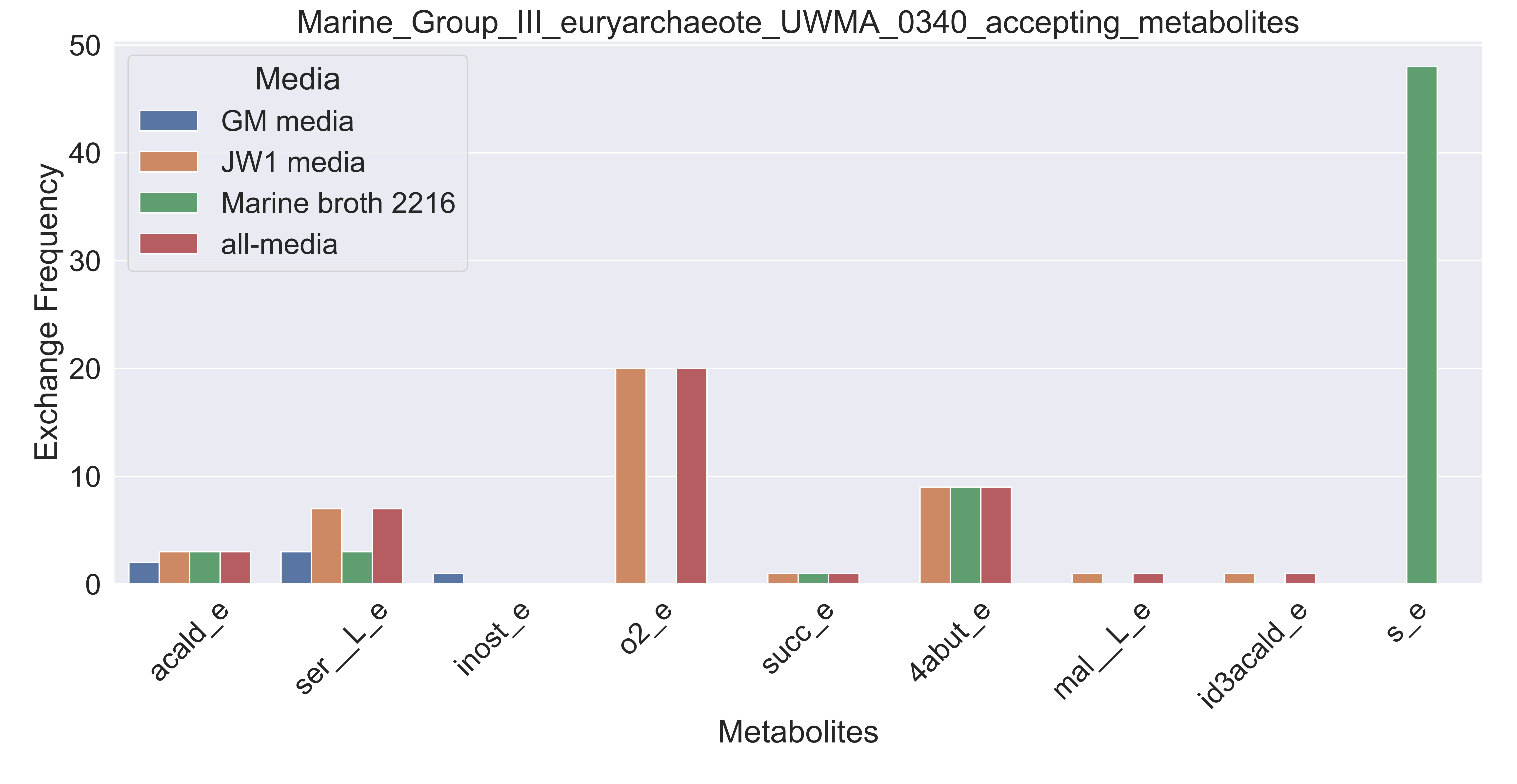

### Supplementary Figure S58 Marine Group II euryarchaeote UWMA 0266 accepting metabolites.pdf

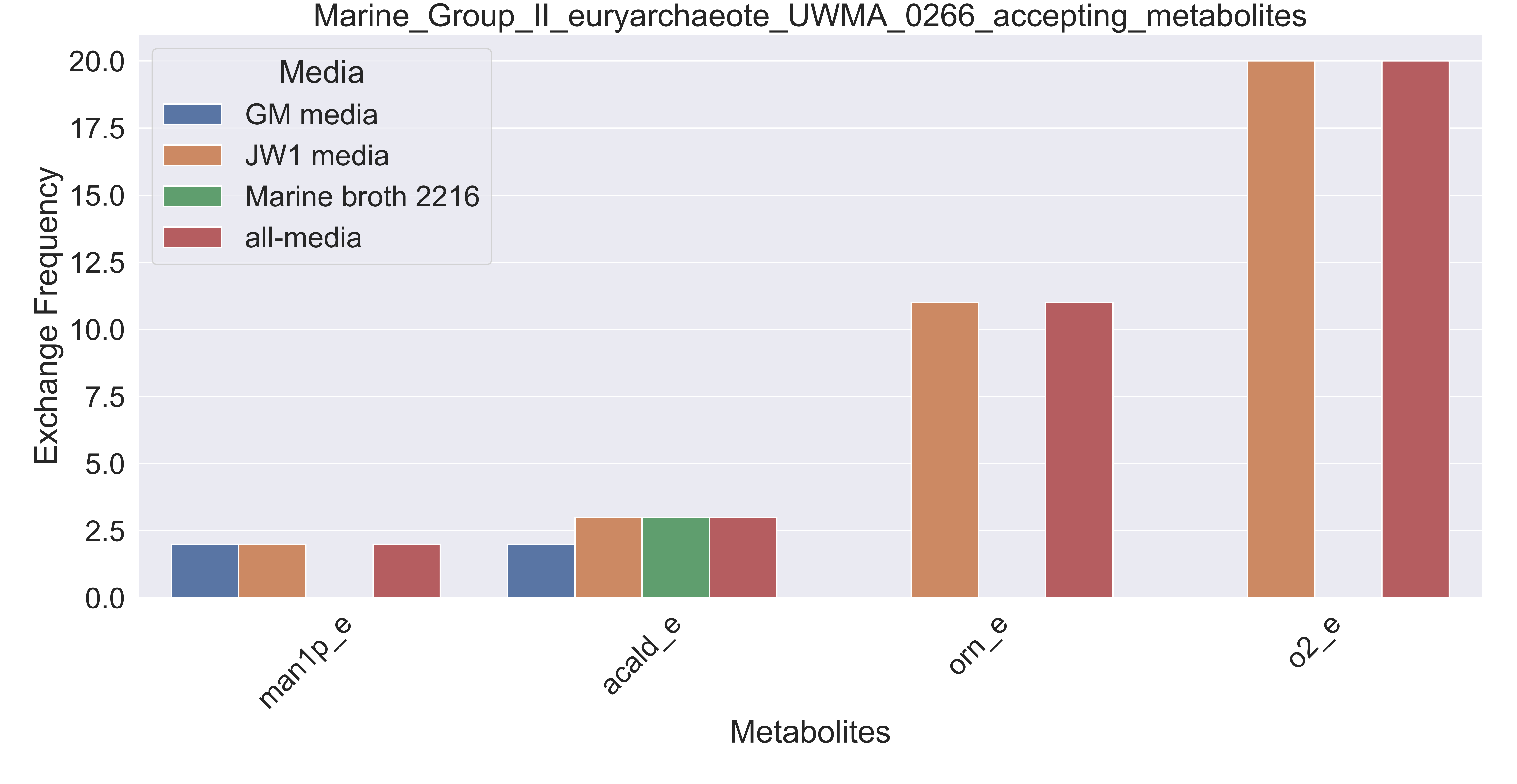

### Supplementary Figure S59 Marine Group II euryarchaeote UWMA 0275 accepting metabolites.pdf

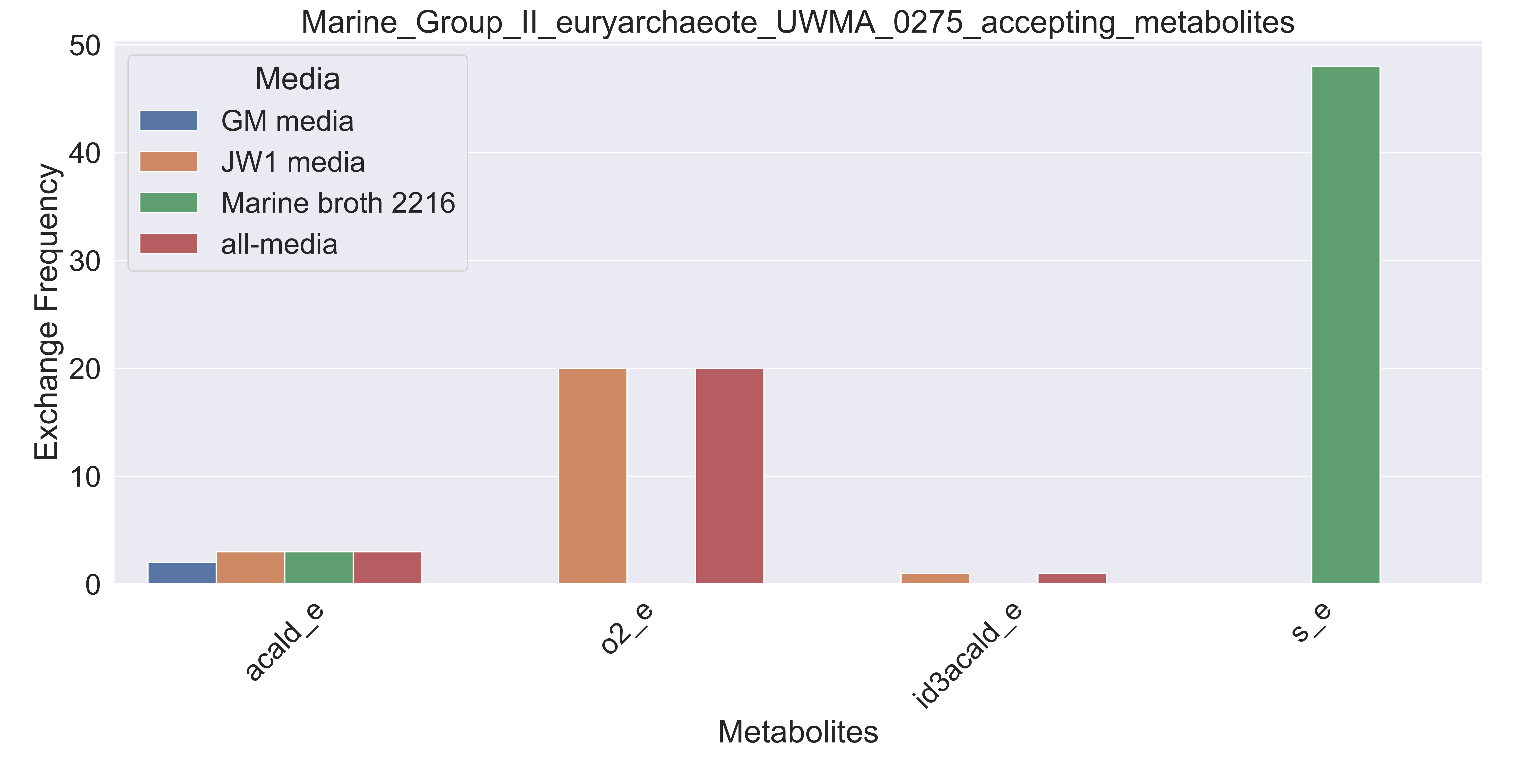

### Supplementary Figure S60 Marine Group II euryarchaeote UWMA 0279 accepting metabolites.pdf

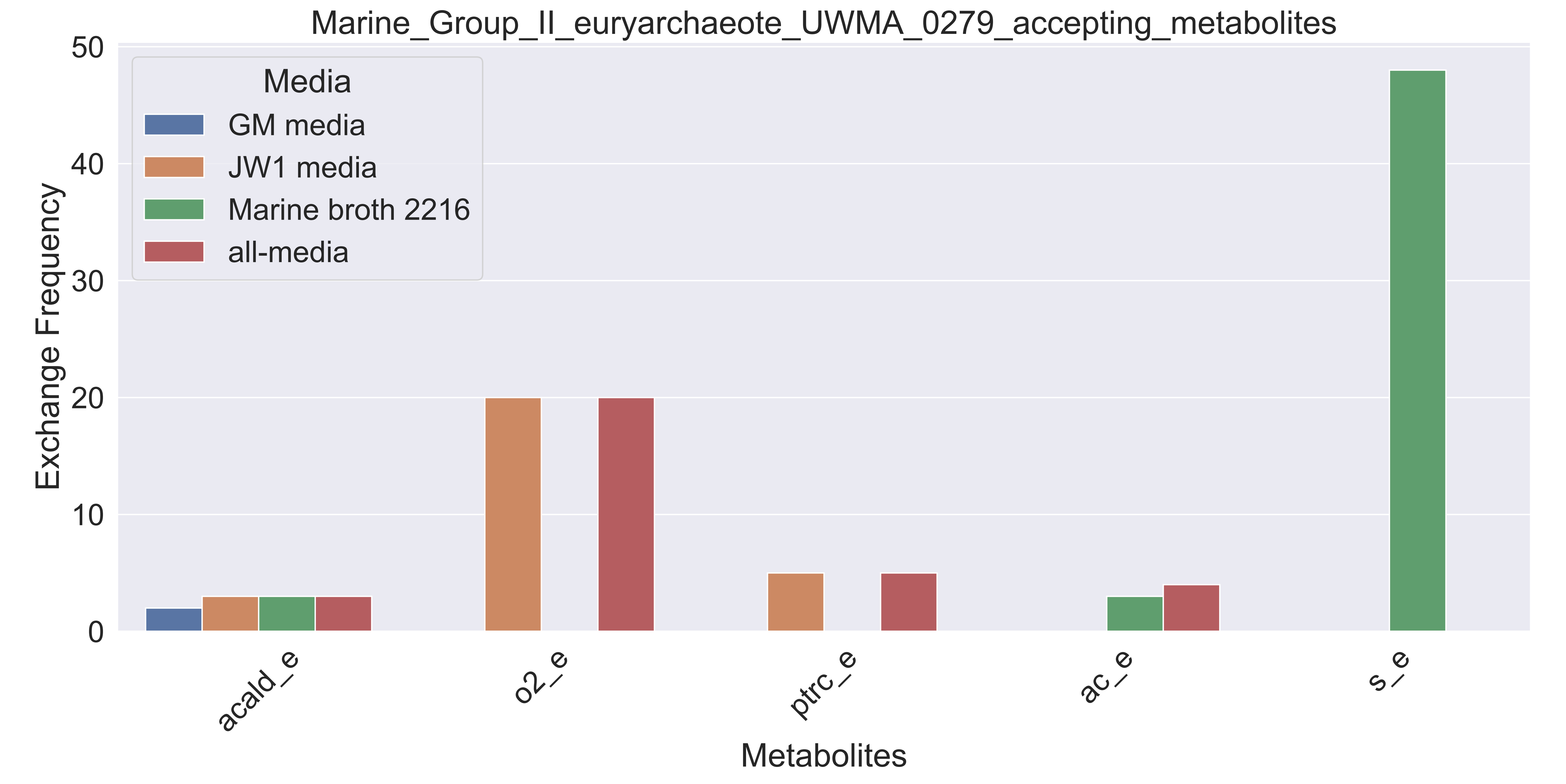

### Supplementary Figure S61 Marine Group II euryarchaeote UWMA 0283 accepting metabolites.pdf

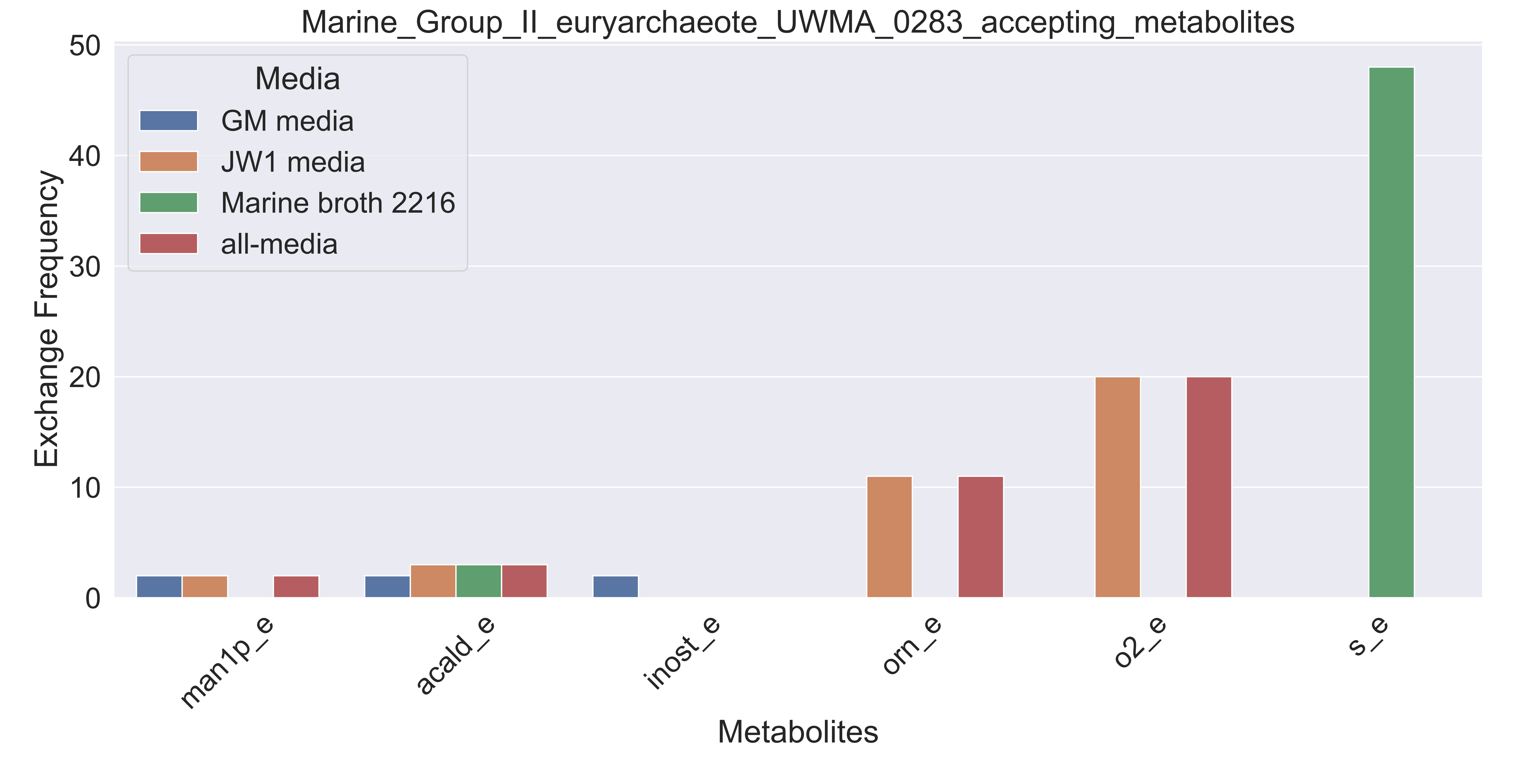

### Supplementary Figure S62 Marine Group II euryarchaeote UWMA 0323 accepting metabolites.pdf

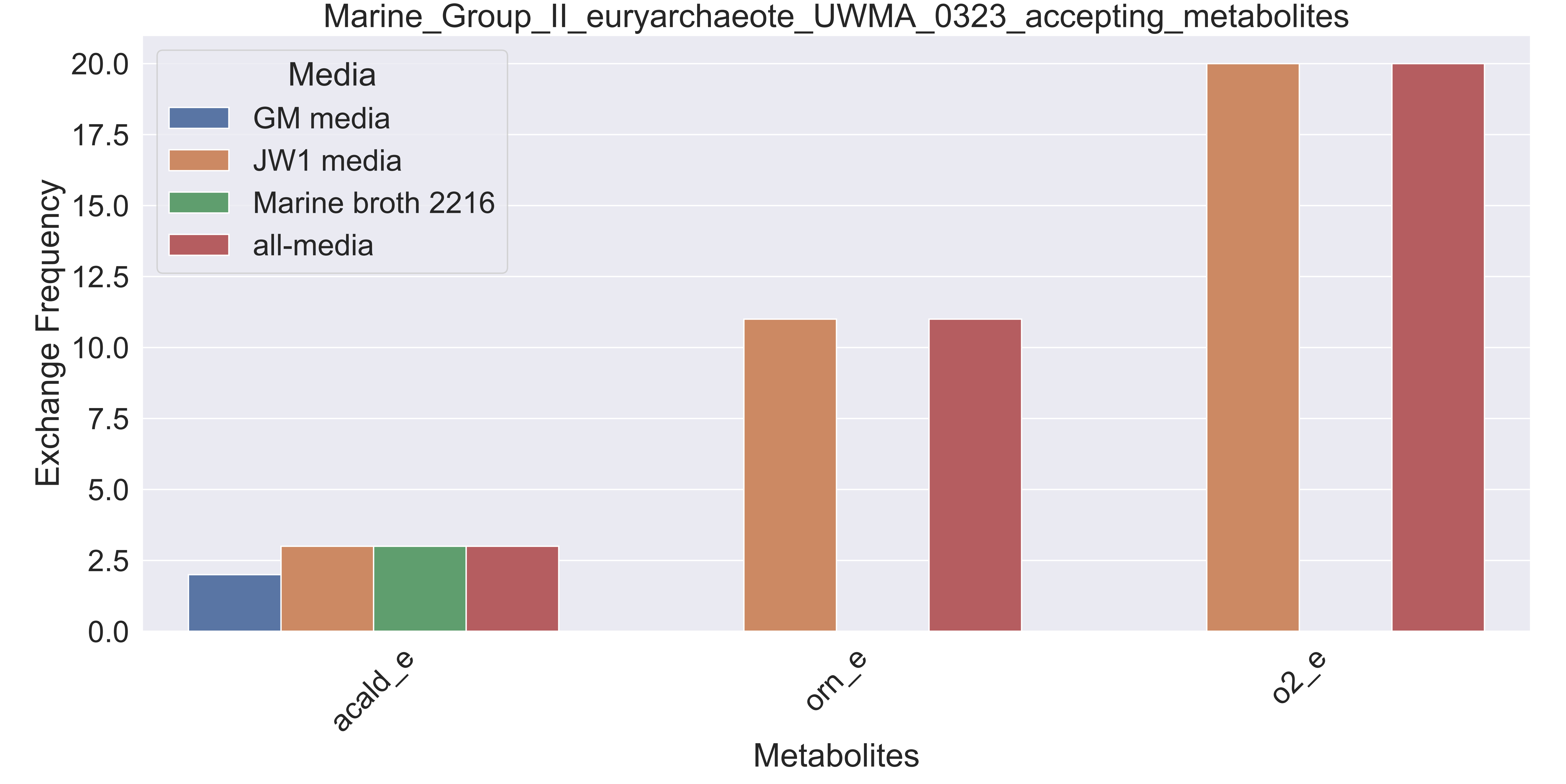

### Supplementary Figure S63 Marine Group II euryarchaeote UWMA 0328 accepting metabolites.pdf

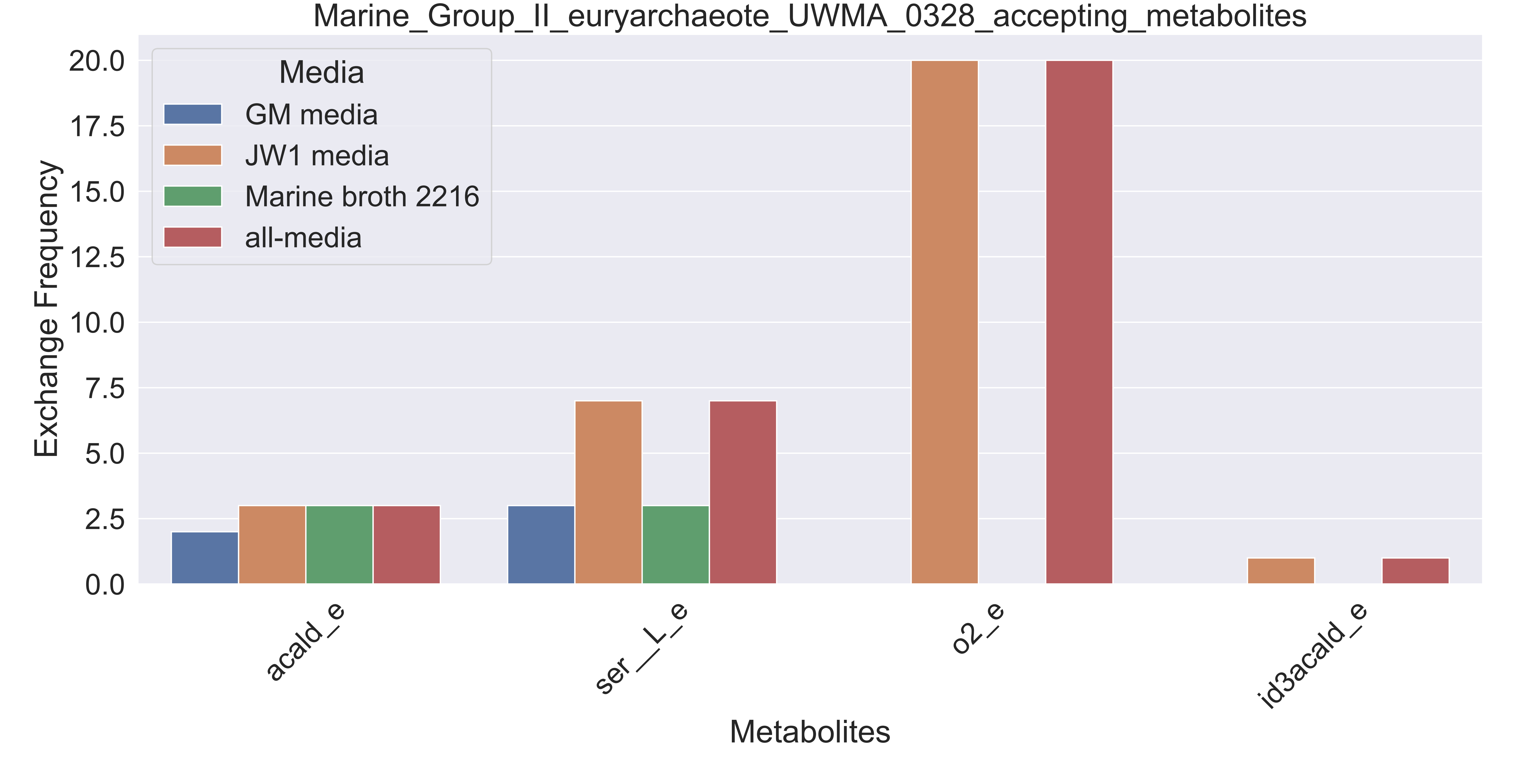

### Supplementary Figure S64 Marine Group II euryarchaeote UWMA 0344 accepting metabolites.pdf

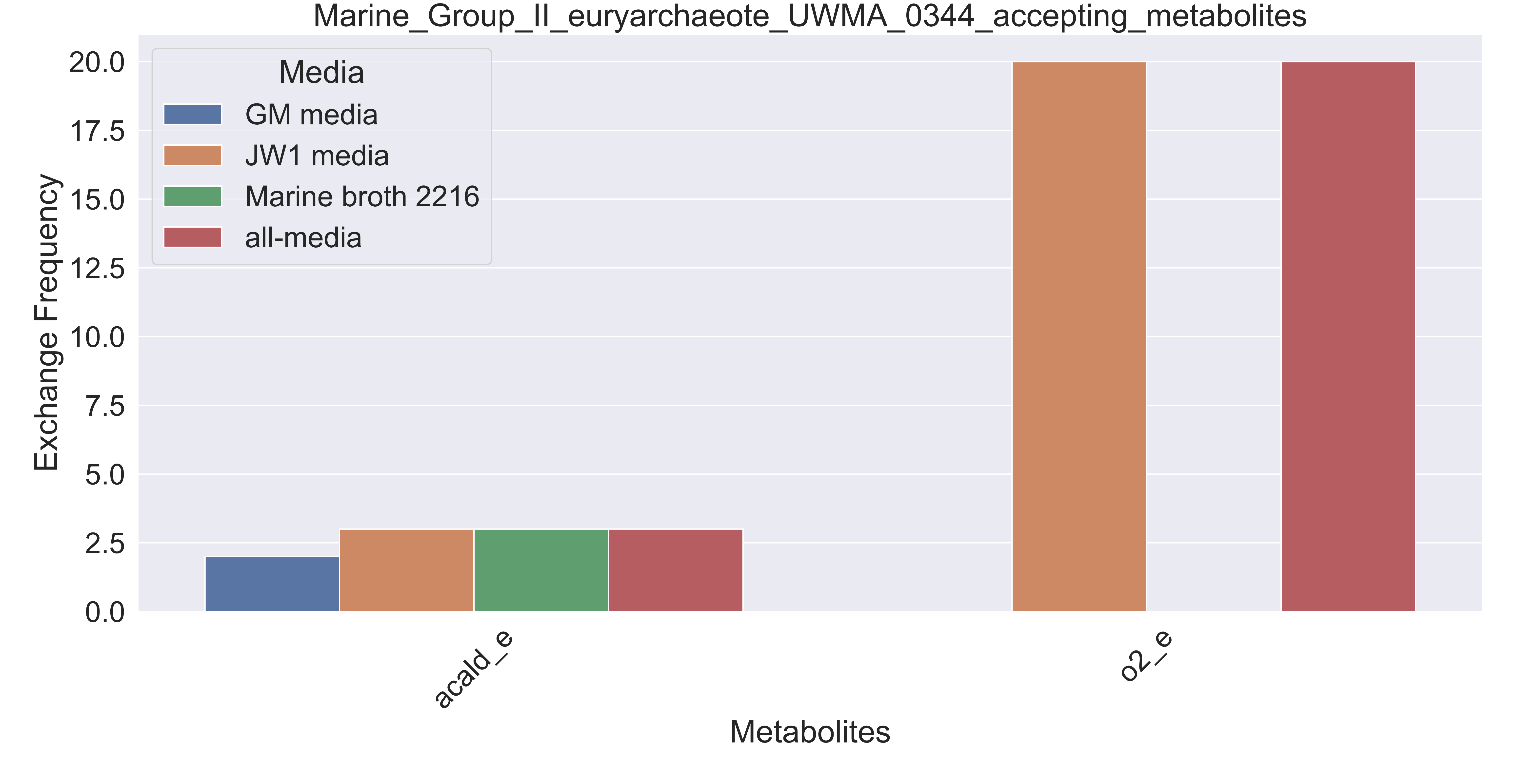

### Supplementary Figure S65 Marine Group II euryarchaeote UWMA 0350 accepting metabolites.pdf

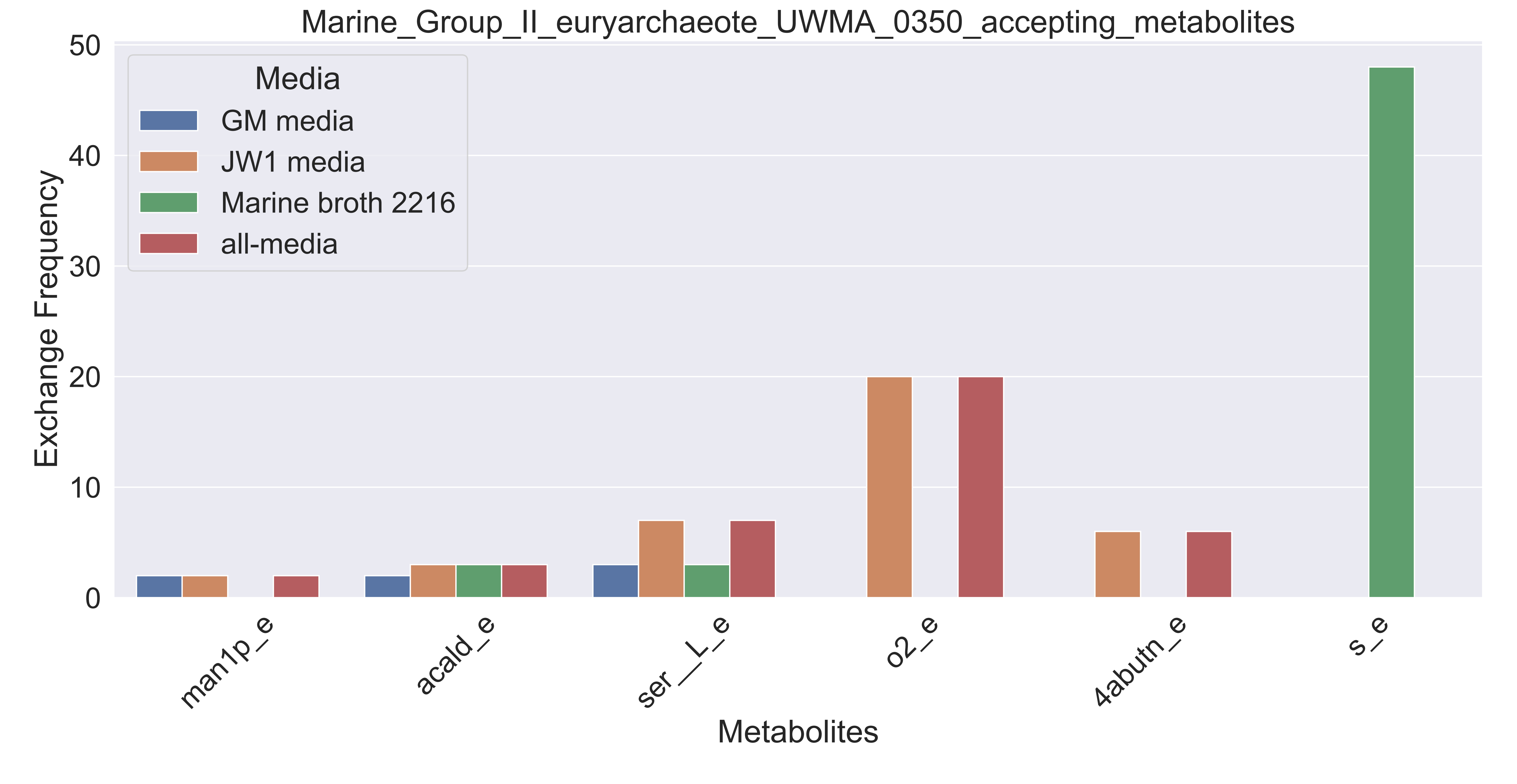

### Supplementary Figure S66 Marine Group II euryarchaeote UWMA 0352 accepting metabolites.pdf

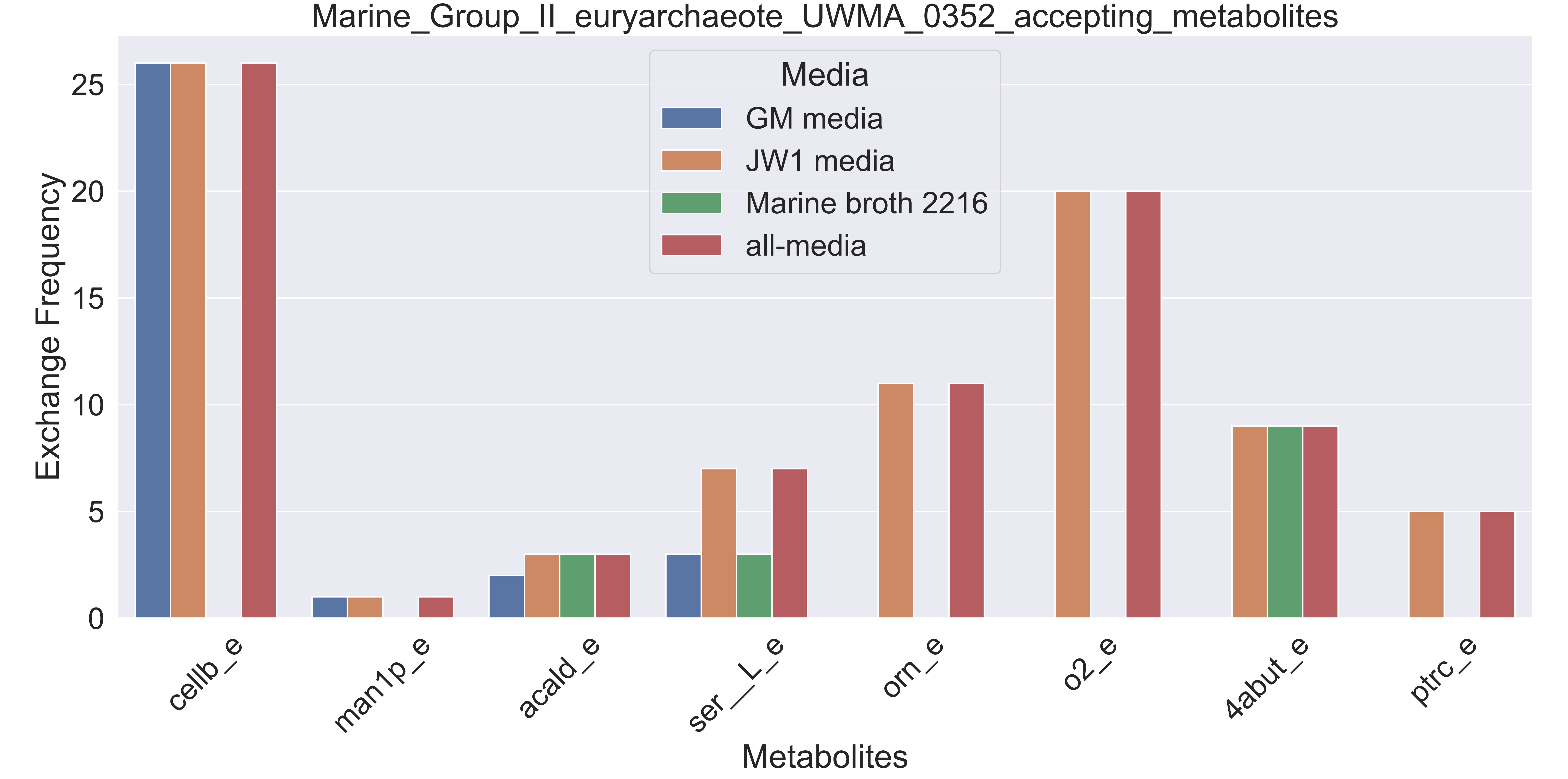

### Supplementary Figure S67 Marine Group II euryarchaeote UWMA 0357 accepting metabolites.pdf

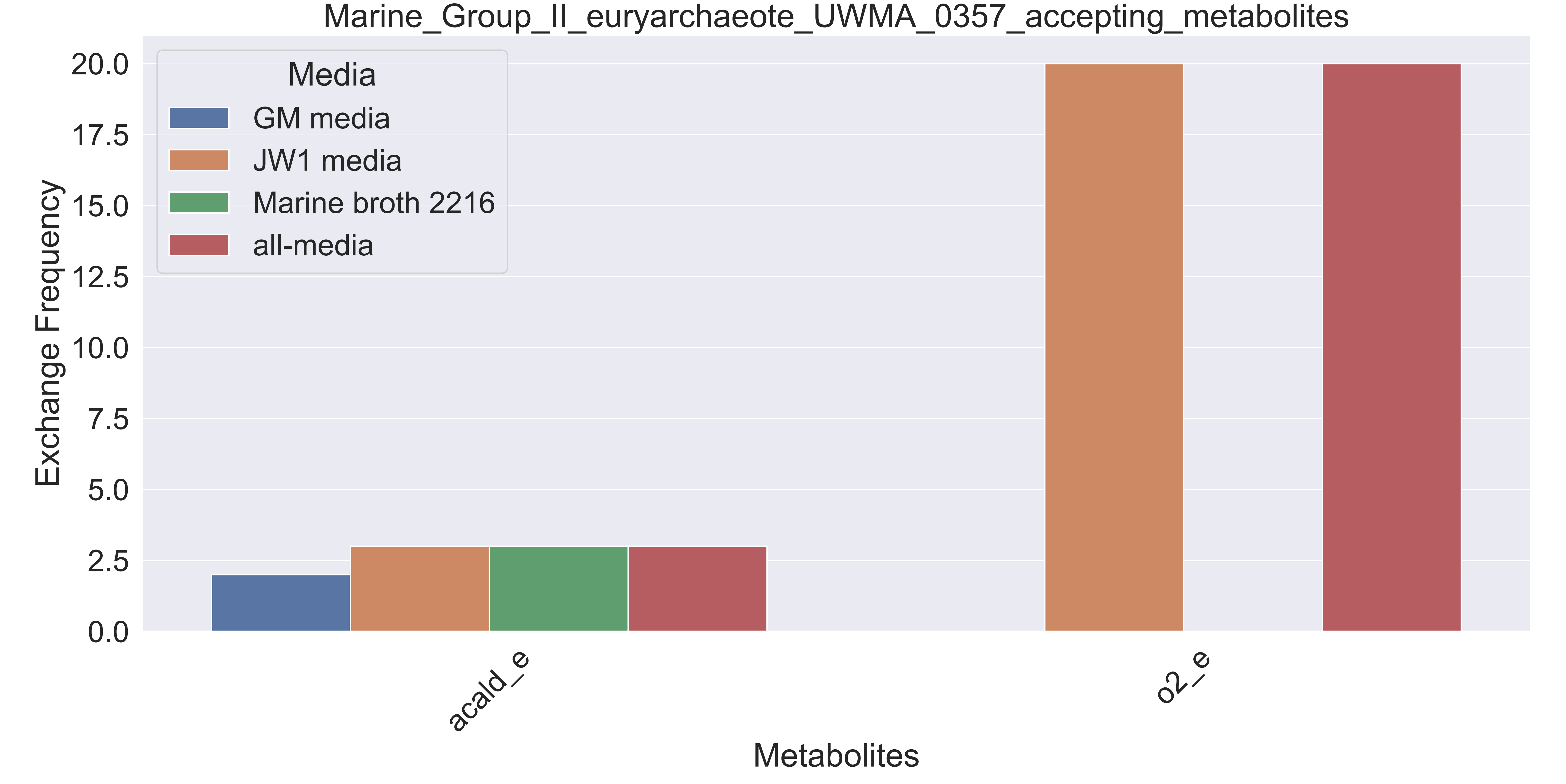

### Supplementary Figure S68 Methylococcaceae bacterium UWMA 0325 accepting metabolites.pdf

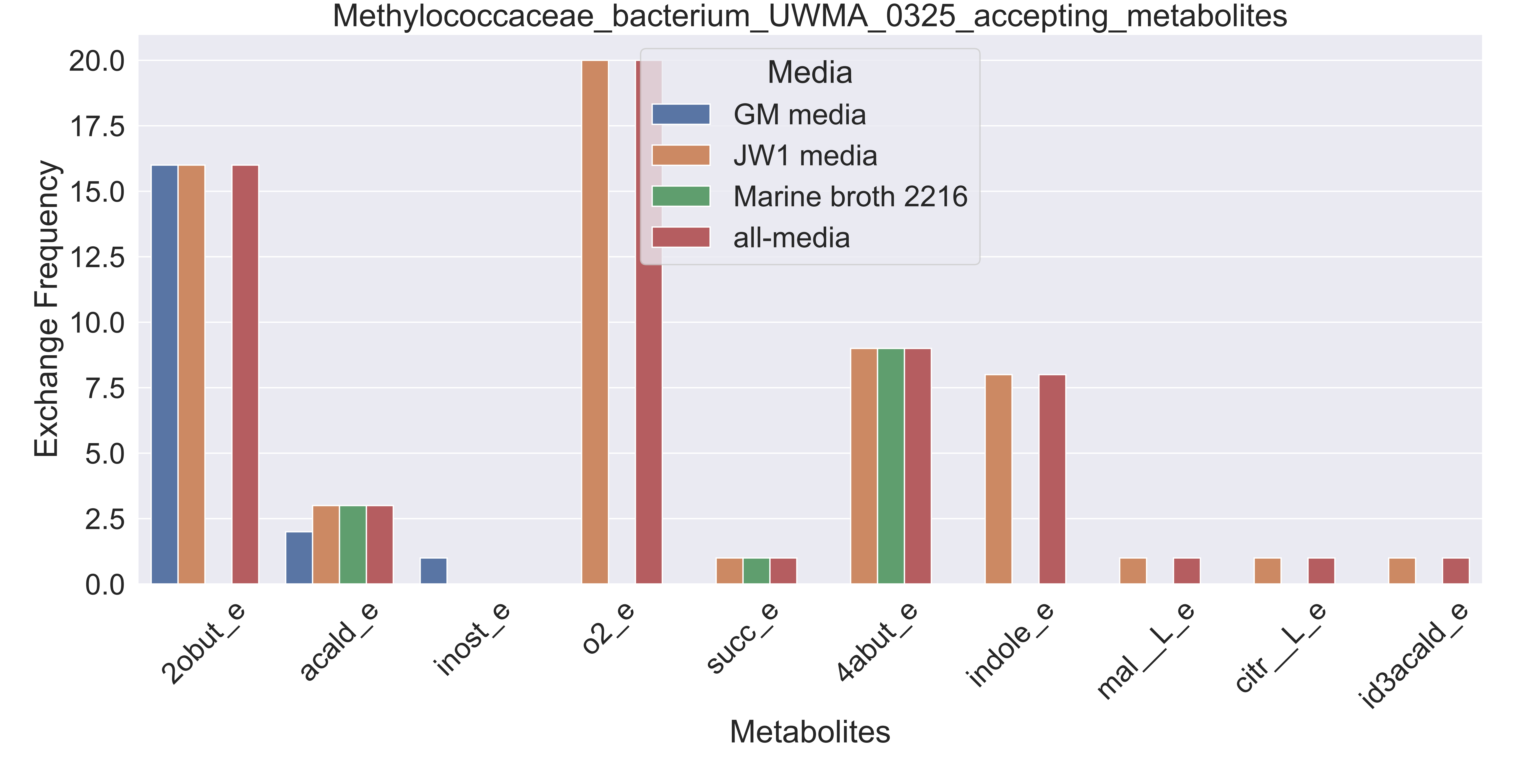

### Supplementary Figure S69 Methylococcaceae bacterium UWMA 0327 accepting metabolites.pdf

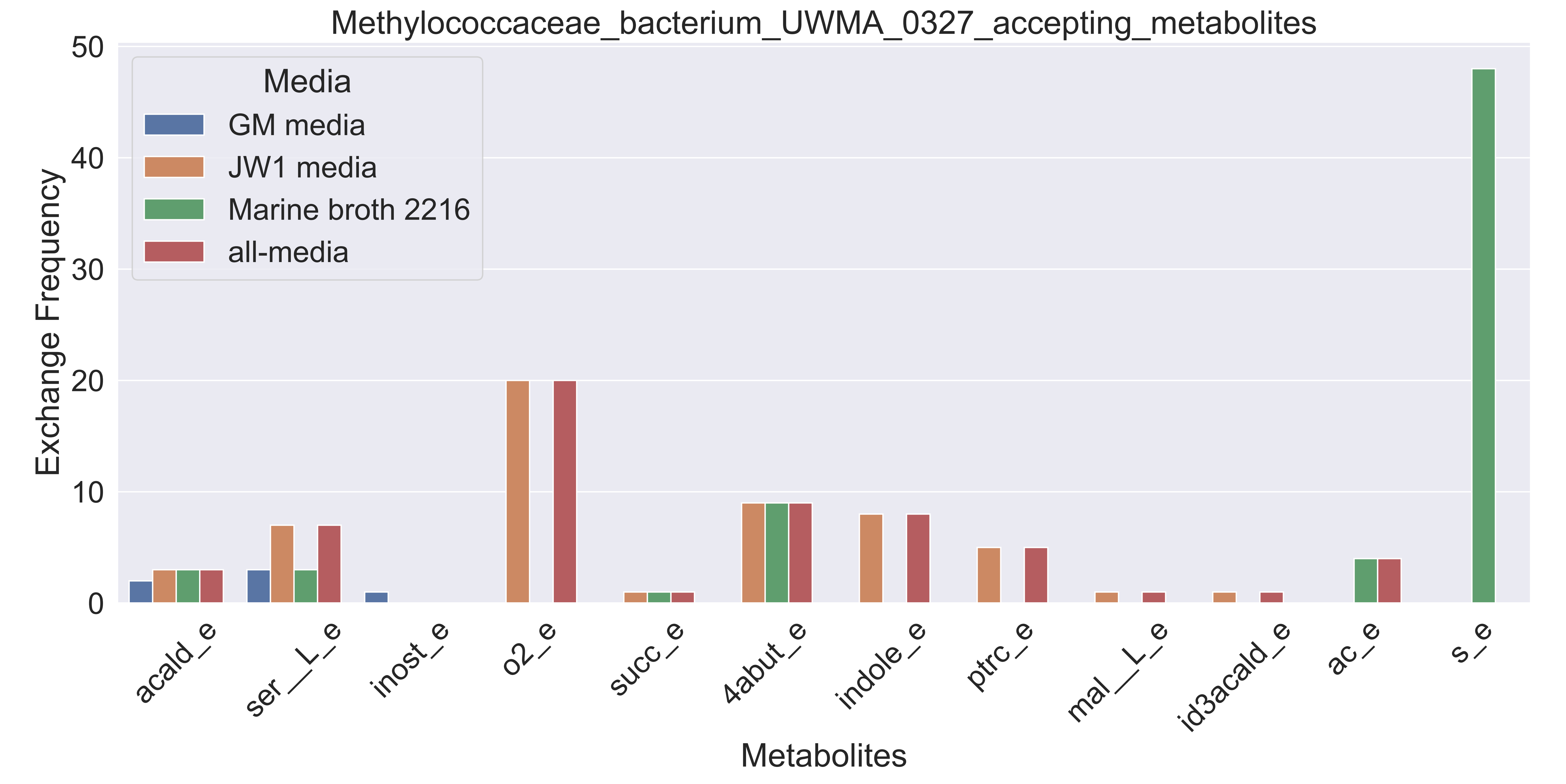

### Supplementary Figure S70 Methylococcaceae bacterium UWMA 0348 accepting metabolites.pdf

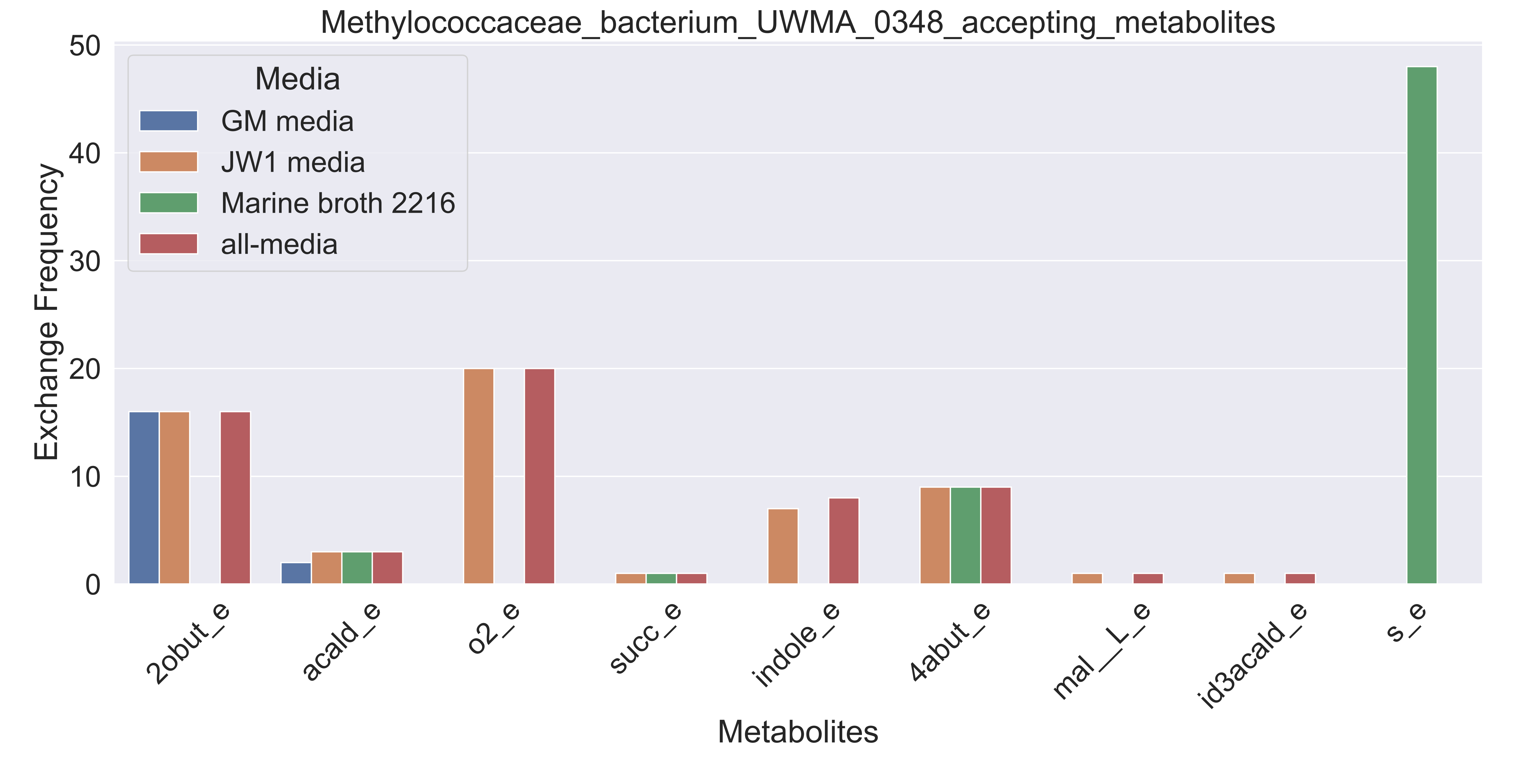

### Supplementary Figure S71 Methylophilaceae bacterium UWMA 0326 accepting metabolites.pdf

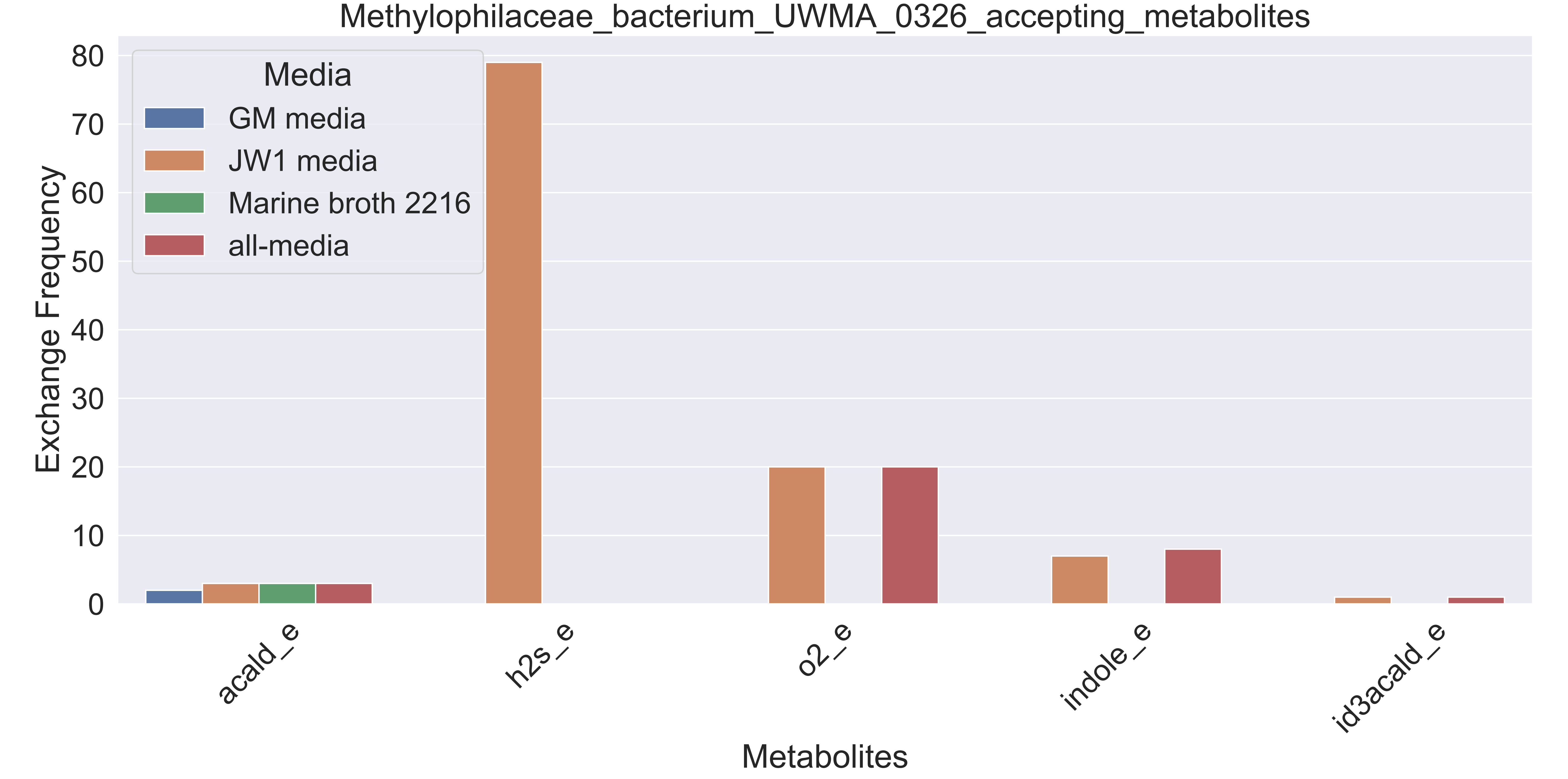

### Supplementary Figure S72 Methyloprofundus sp UWMA 0312 accepting metabolites.pdf

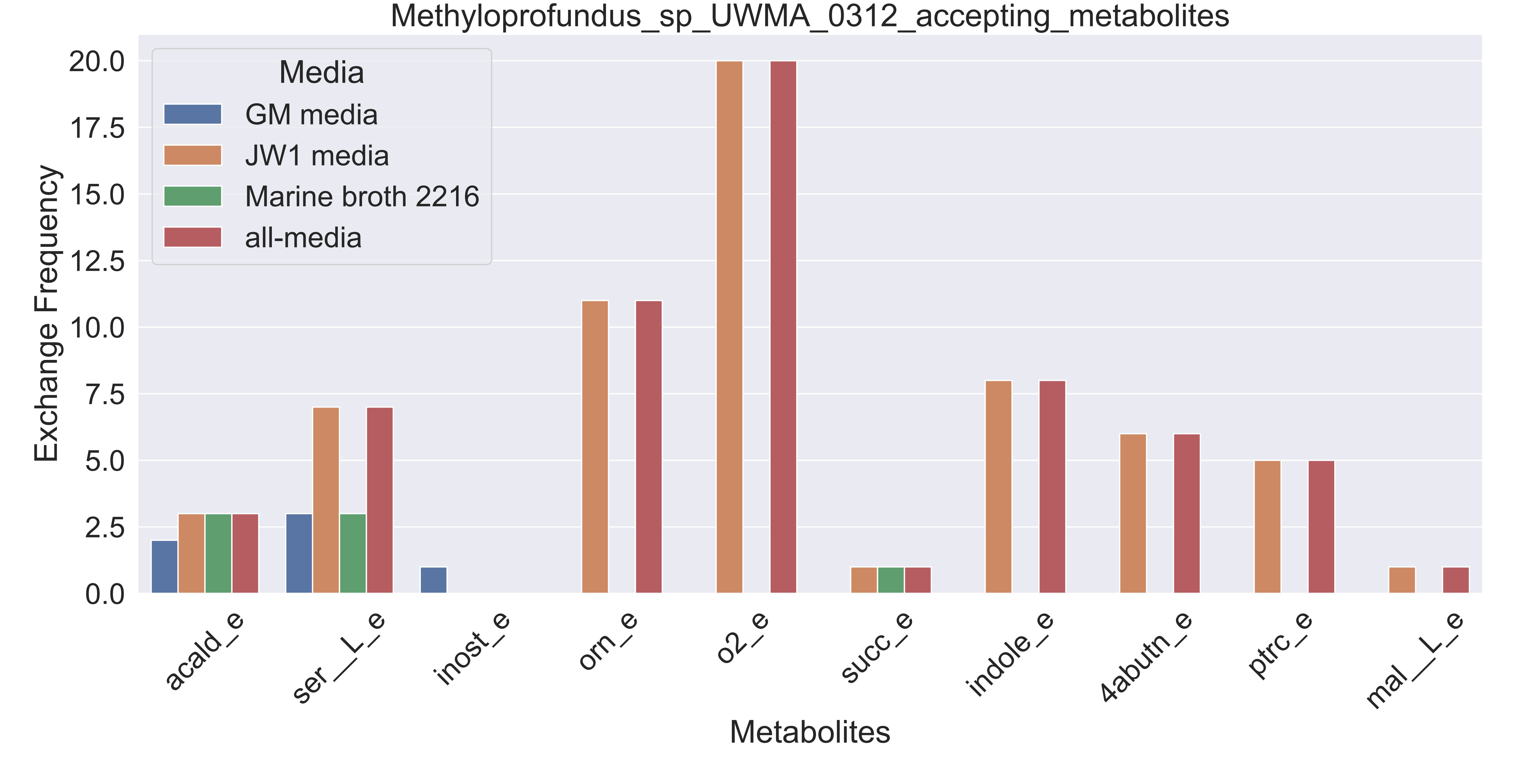

### Supplementary Figure S73 Micavibrio sp UWMA 0310 accepting metabolites.pdf

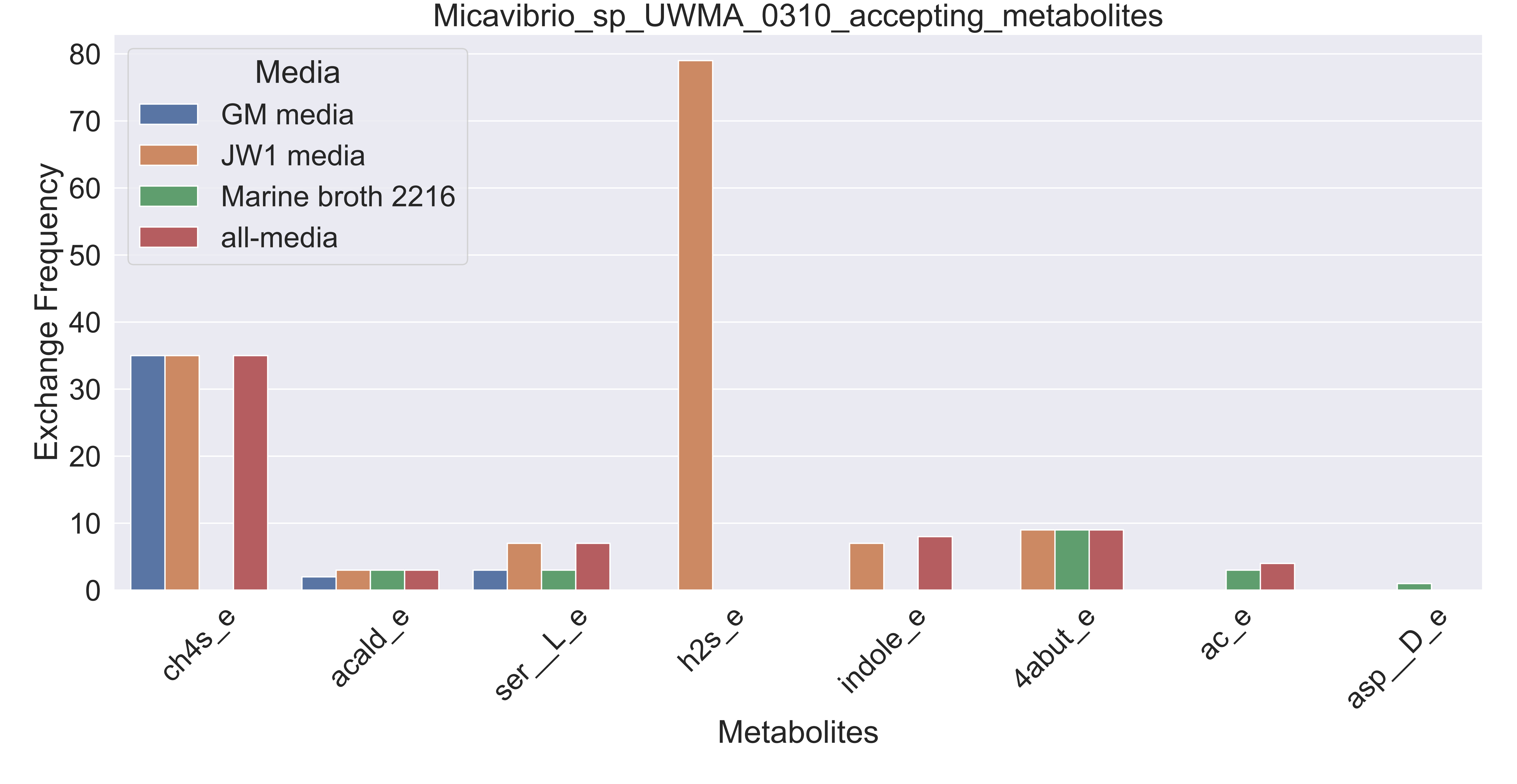

### Supplementary Figure S74 Microbacterium sp UWMA 0270 accepting metabolites.pdf

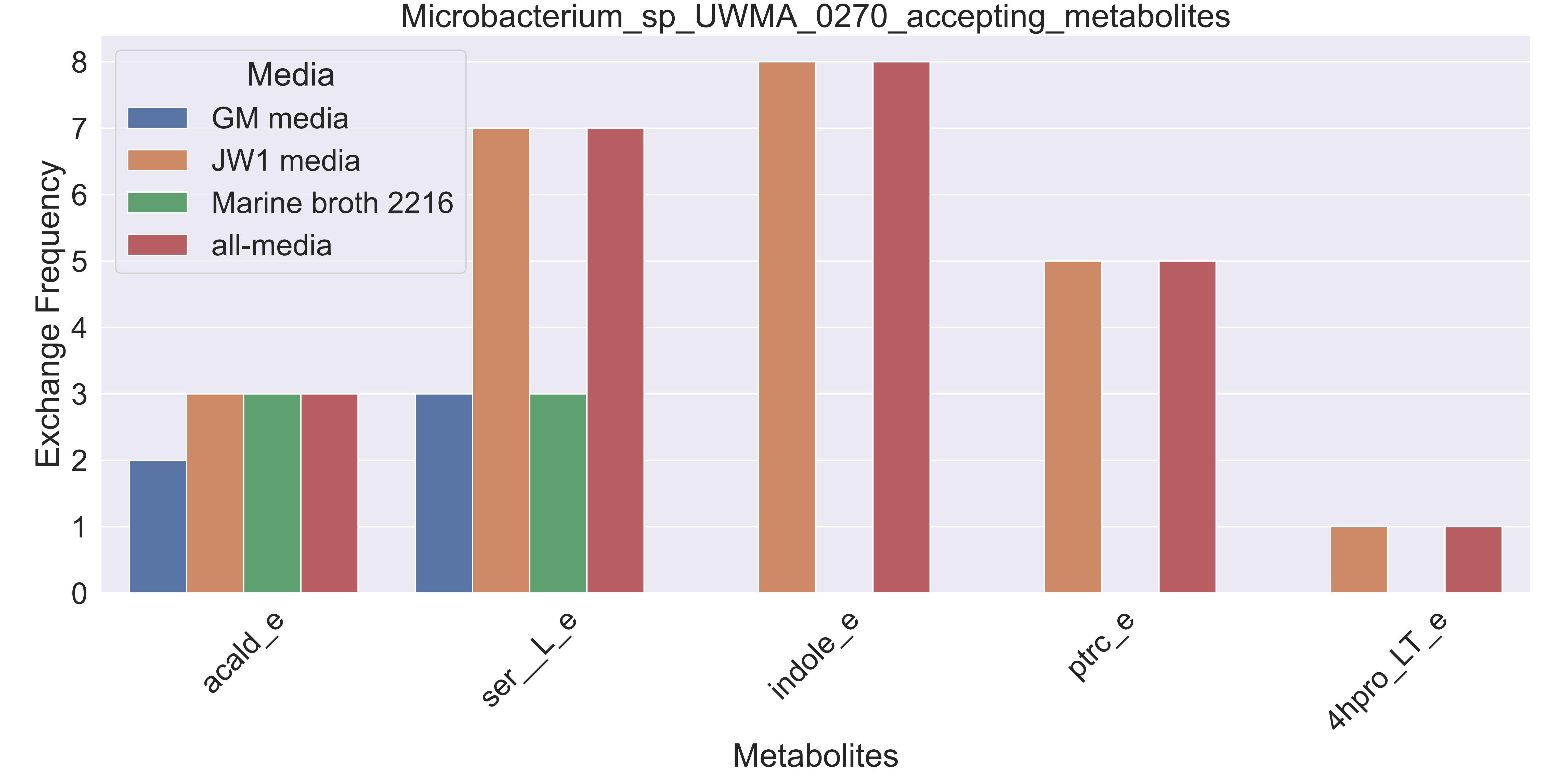

### Supplementary Figure S75 Myxococcales bacterium UWMA 0315 accepting metabolites.pdf

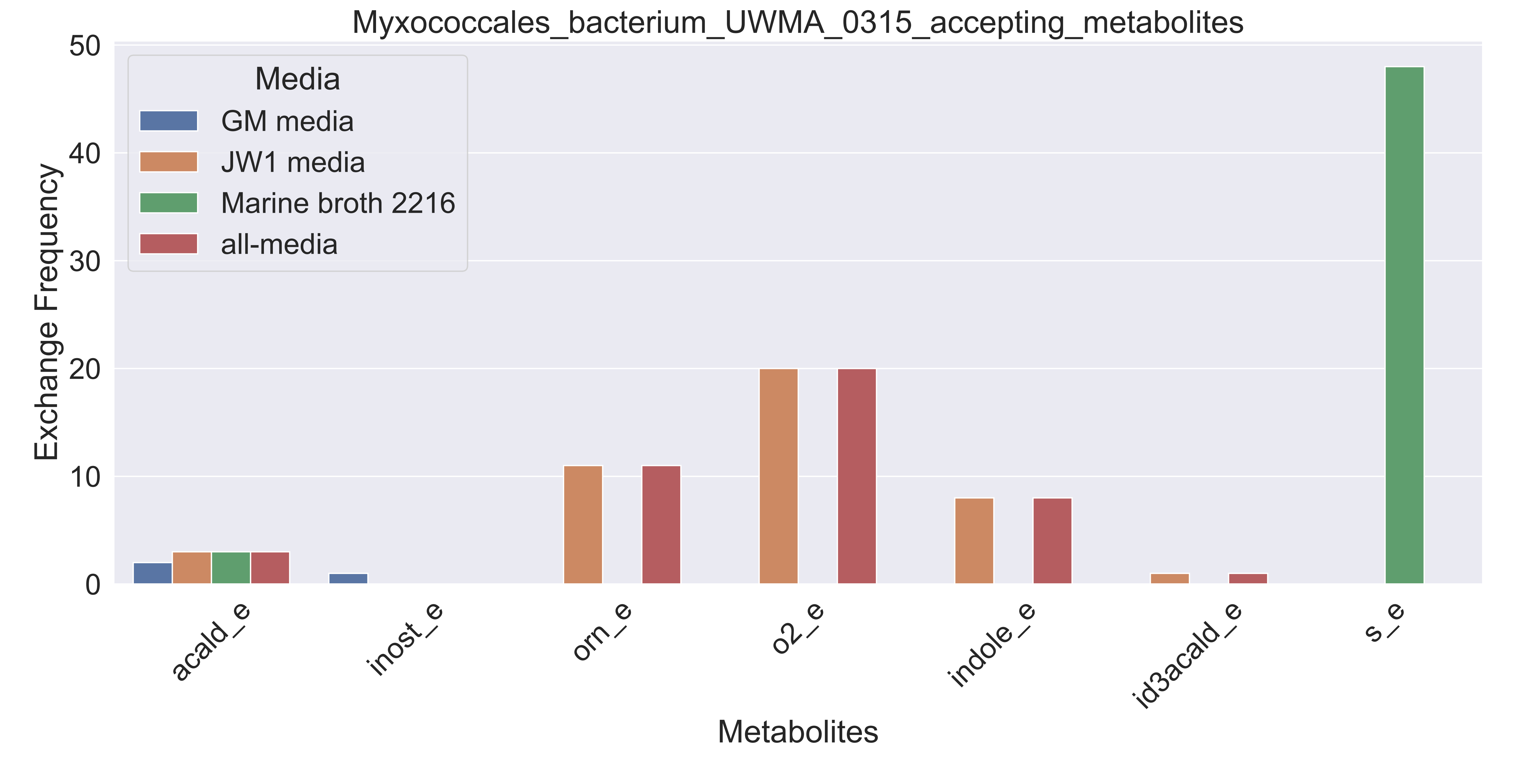

### Supplementary Figure S76 Myxococcales bacterium UWMA 0324 accepting metabolites.pdf

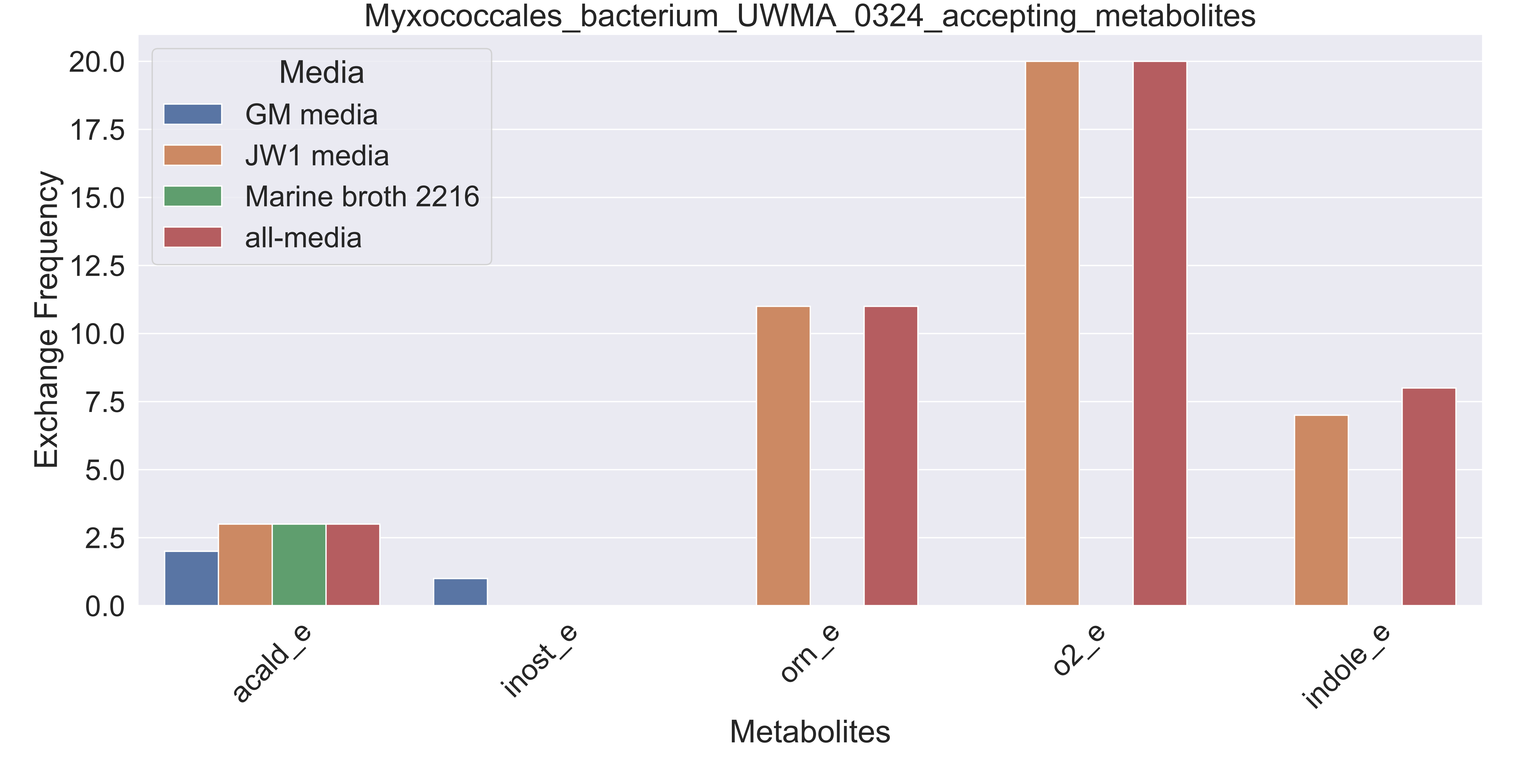

### Supplementary Figure S77 Myxococcales bacterium UWMA 0330 accepting metabolites.pdf

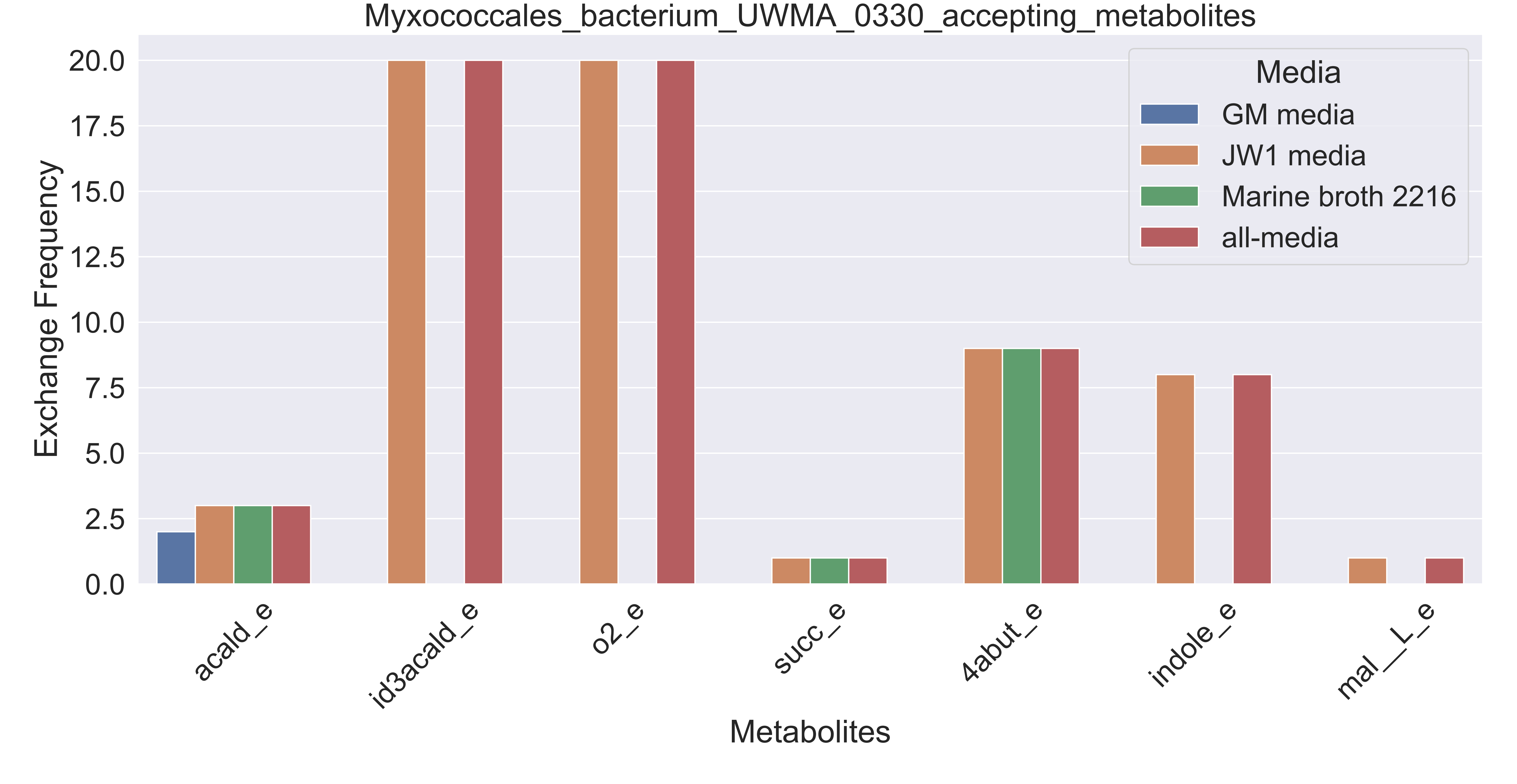

### Supplementary Figure S78 Myxococcales bacterium UWMA 0336 accepting metabolites.pdf

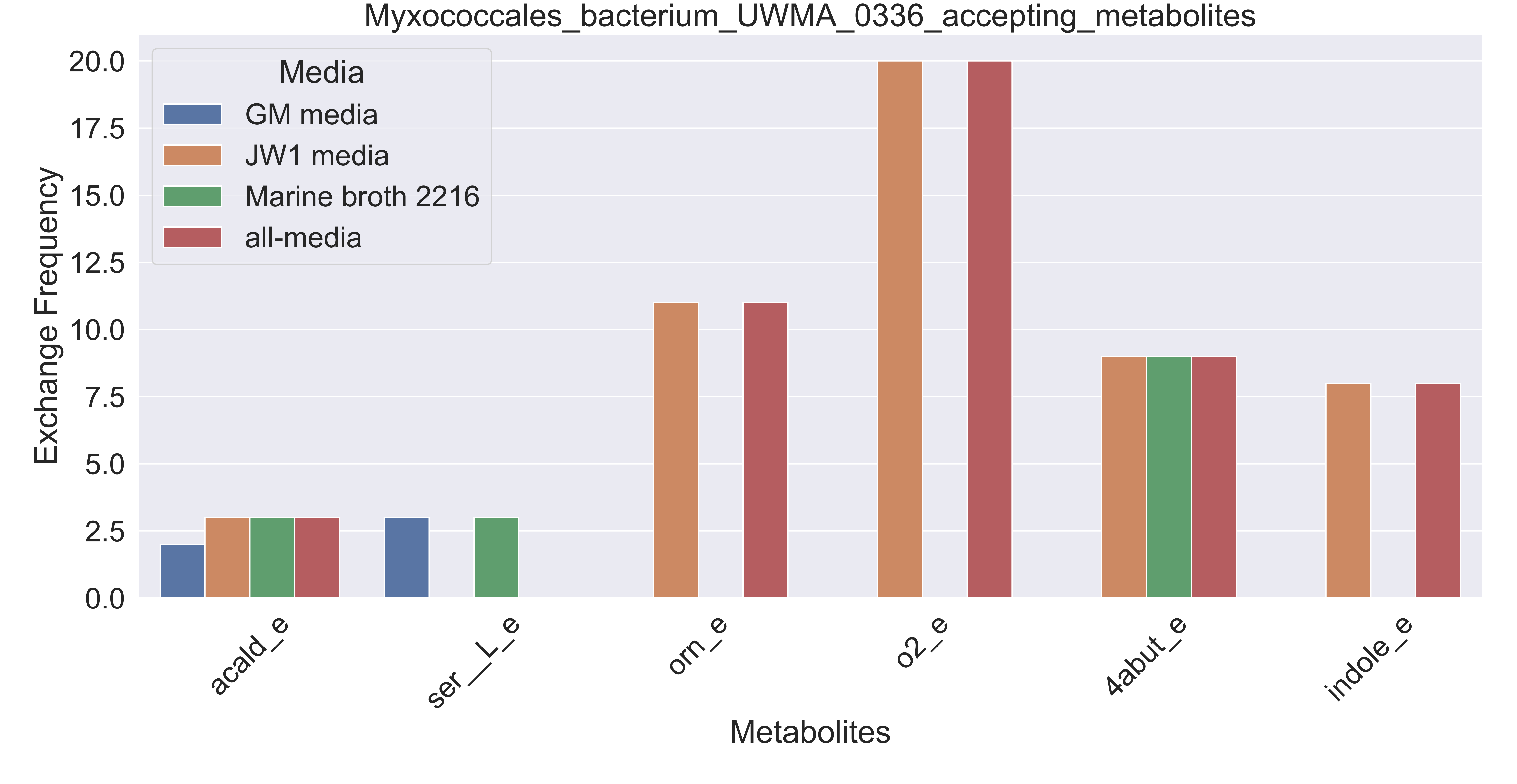

### Supplementary Figure S79 Nitrosopumilus sp UWMA 0263 accepting metabolites.pdf

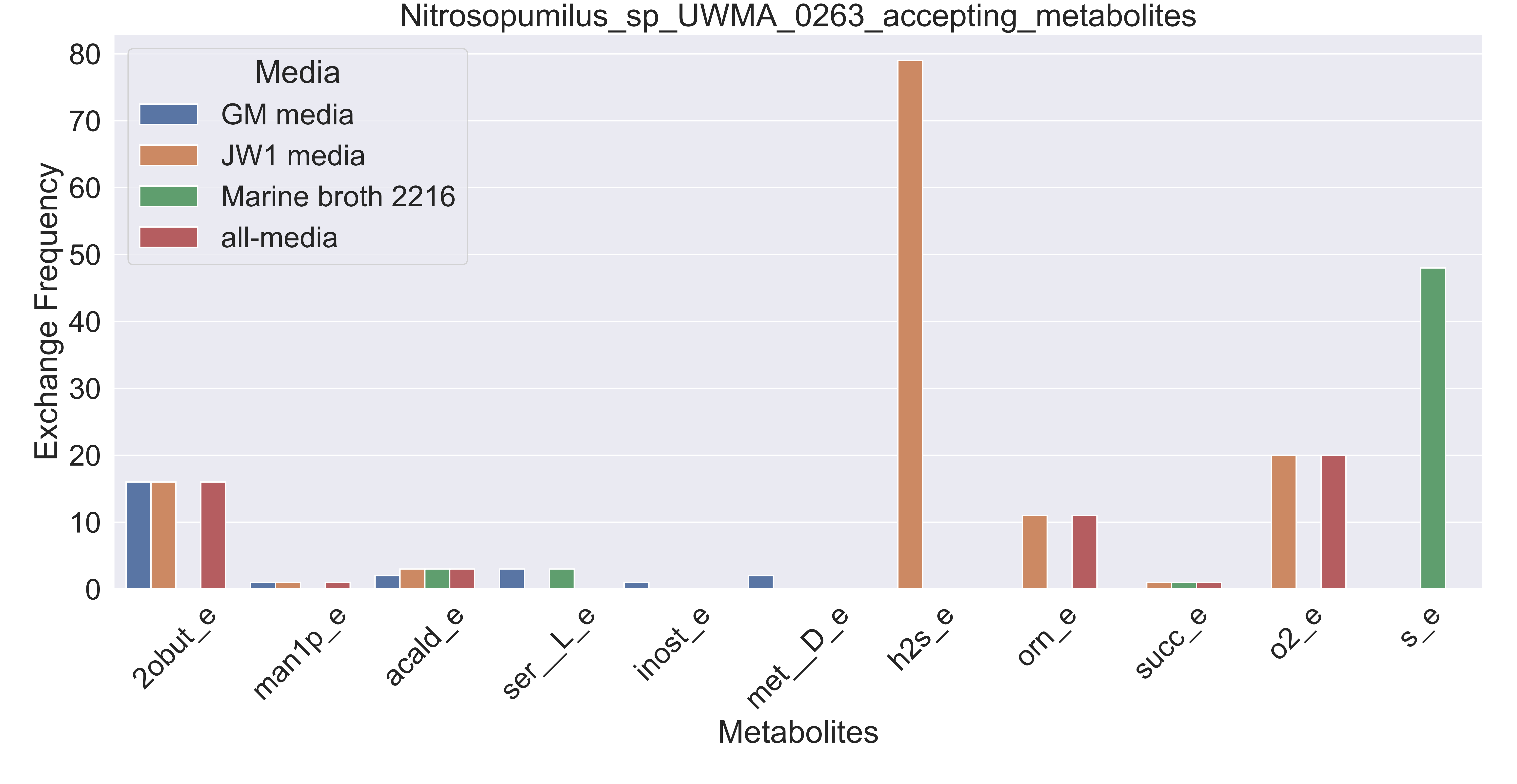

### Supplementary Figure S80 Nitrospinaceae bacterium UWMA 0262 accepting metabolites.pdf

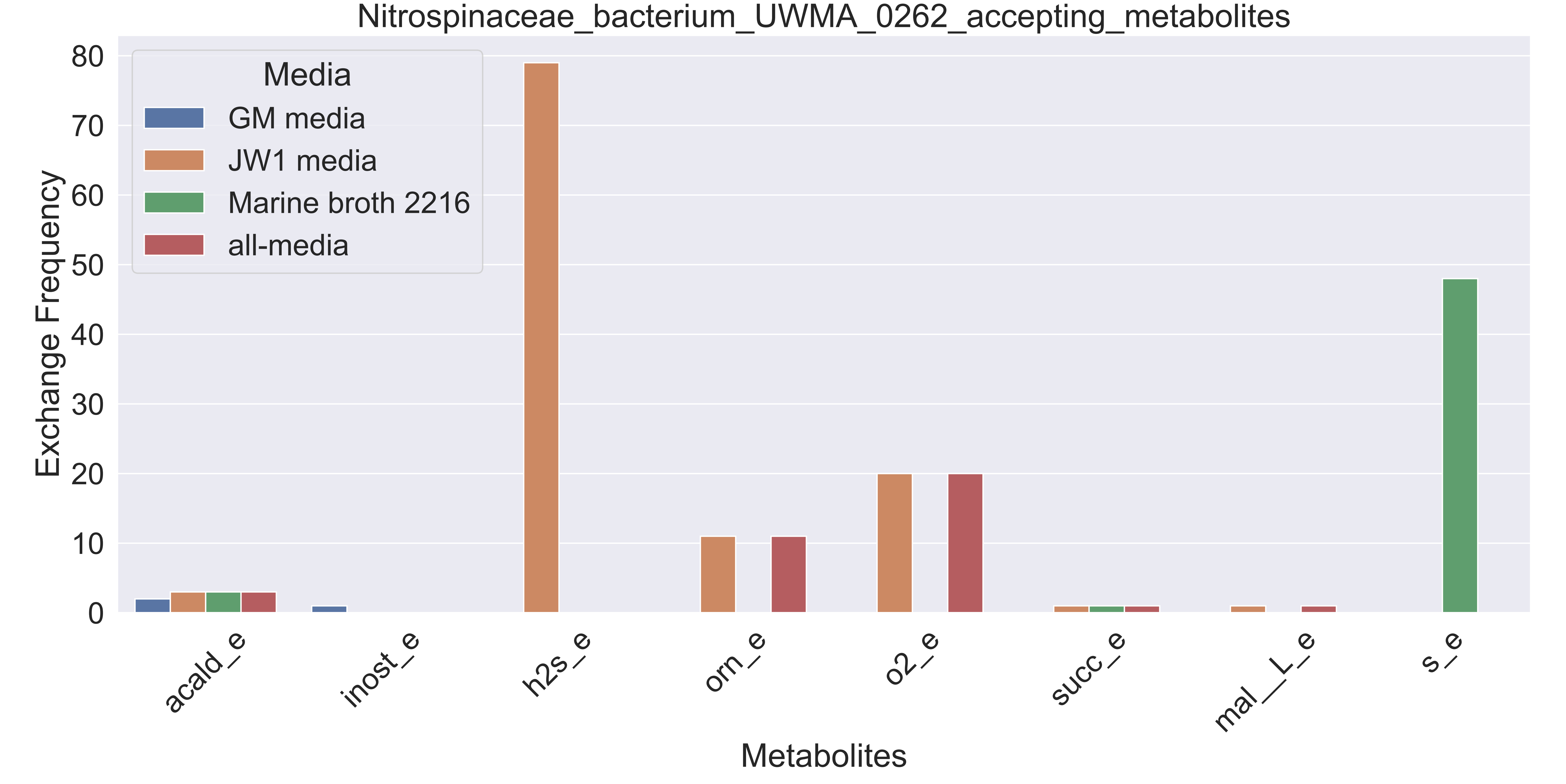

### Supplementary Figure S81 Nitrospinaceae bacterium UWMA 0271 accepting metabolites.pdf

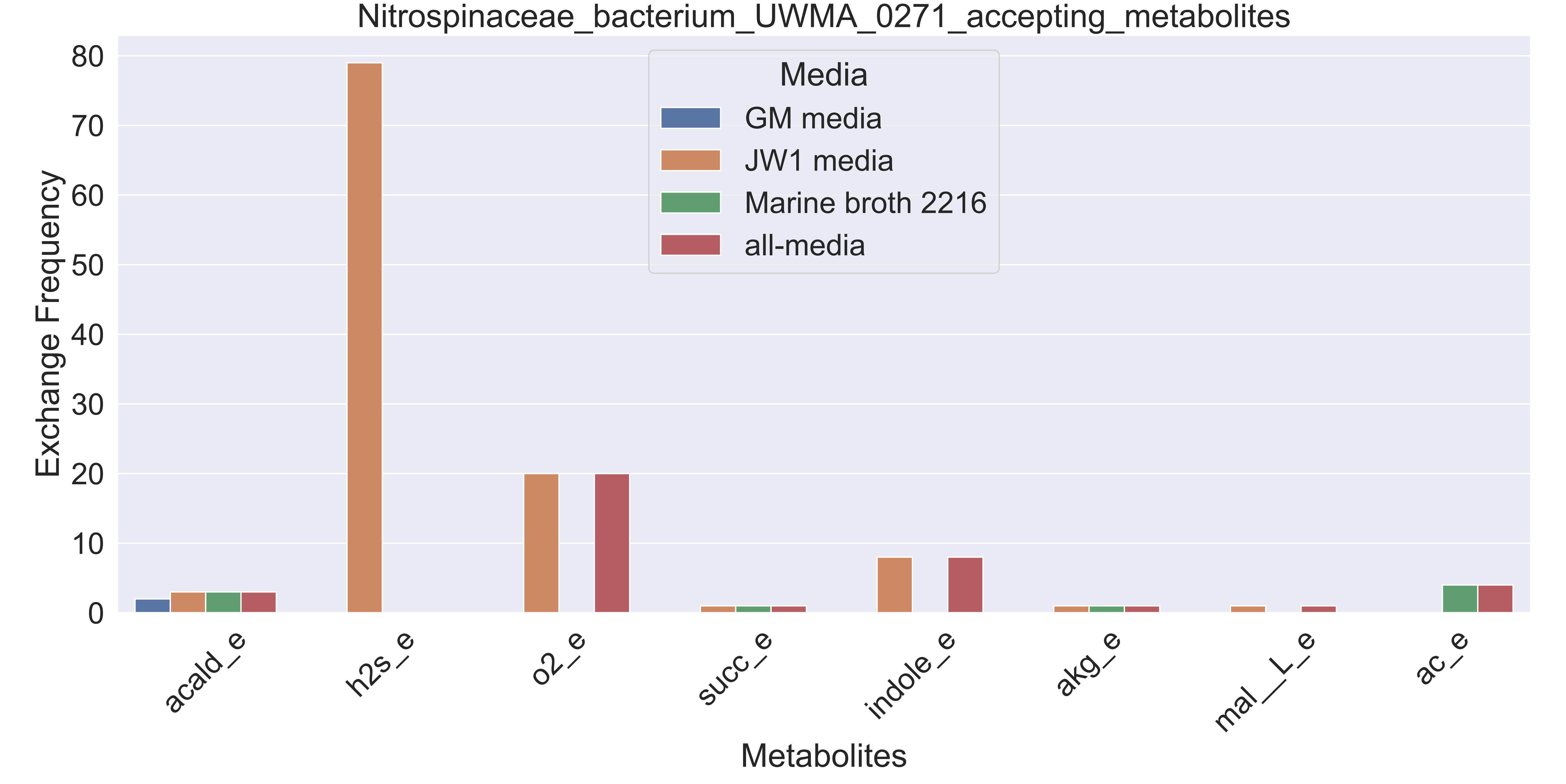

### Supplementary Figure S82 Nitrospinaceae bacterium UWMA 0356 accepting metabolites.pdf

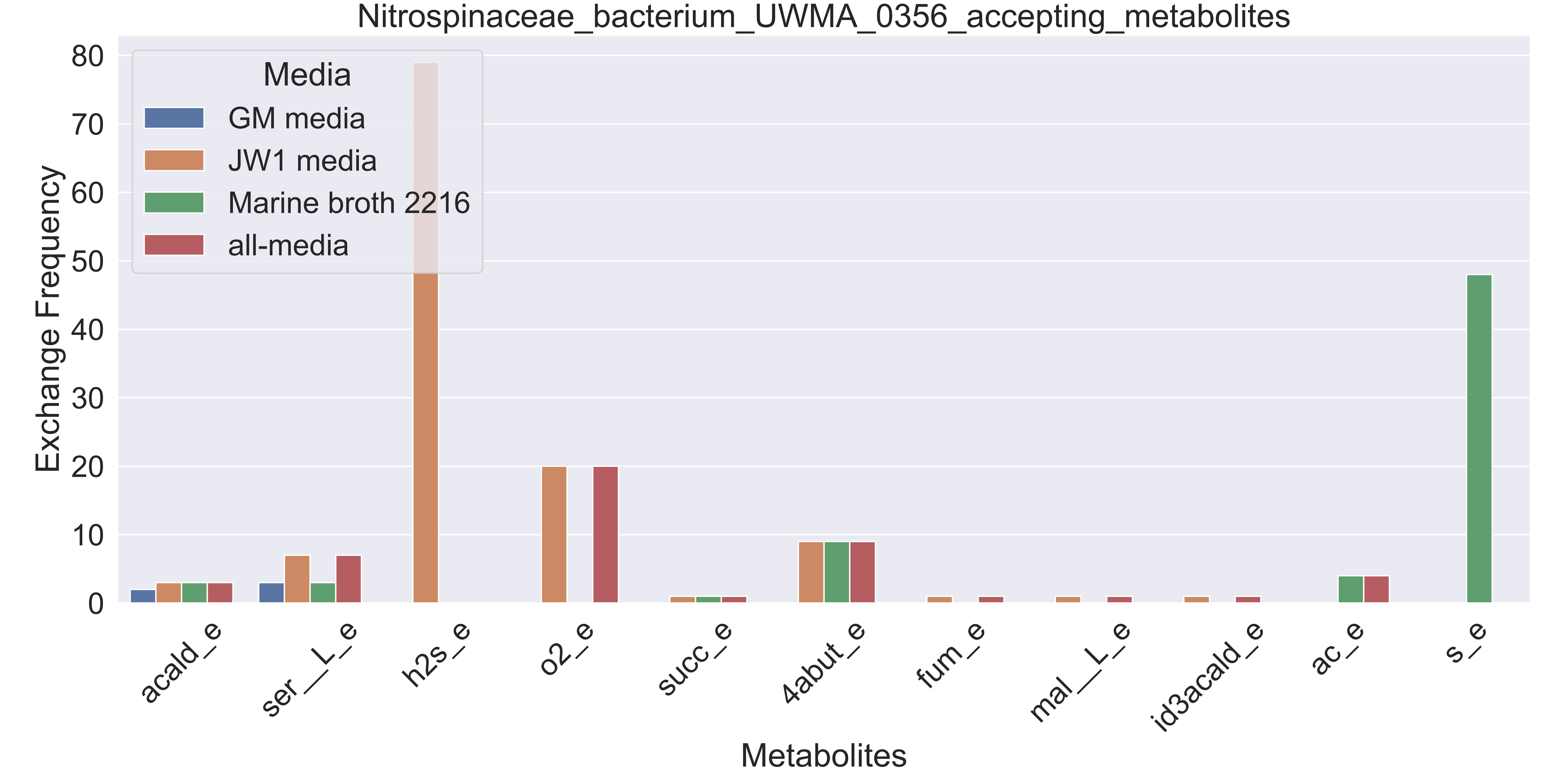

### Supplementary Figure S83 Nitrospira sp UWMA 0268 accepting metabolites.pdf

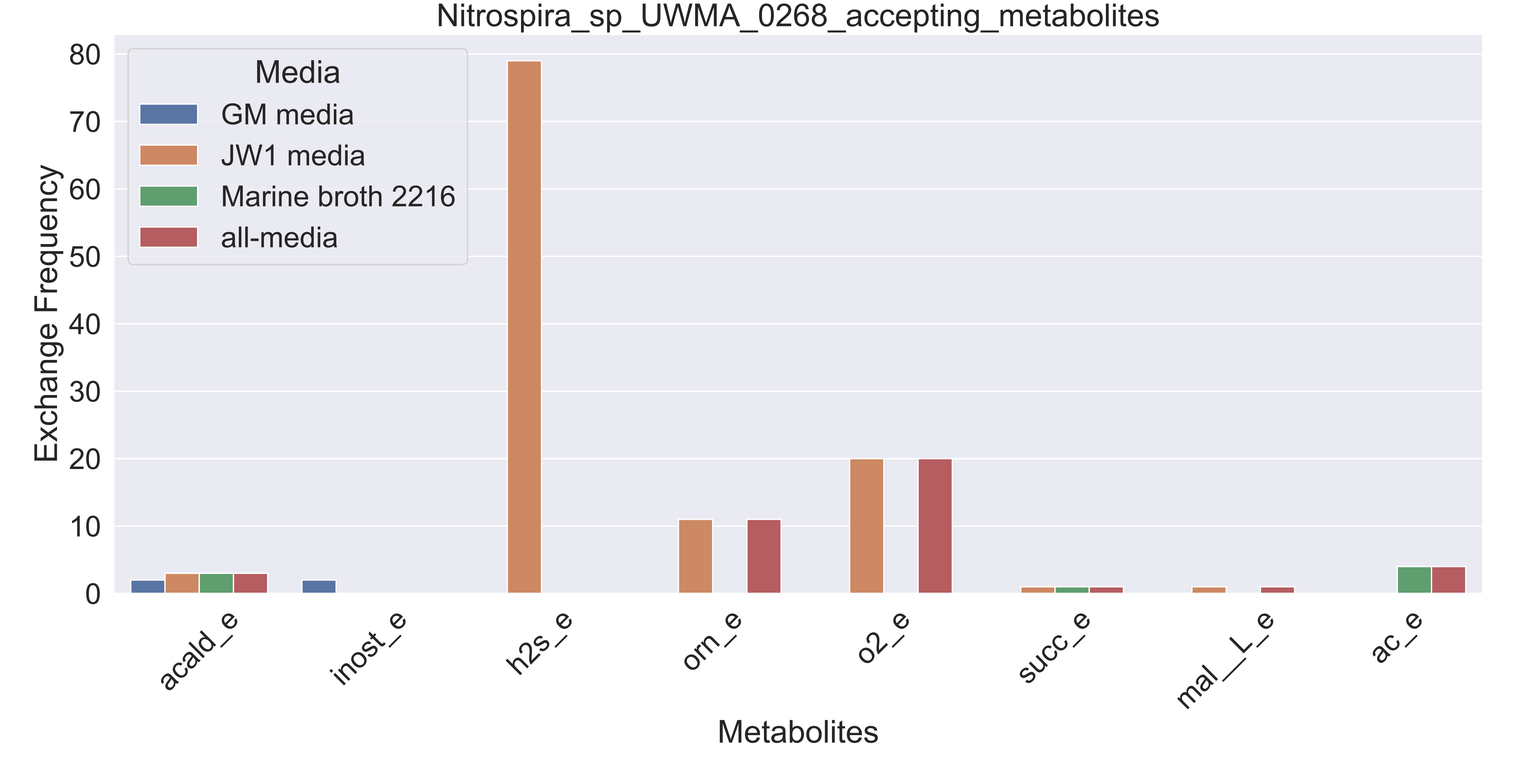

### Supplementary Figure S84 Oceanospirillaceae bacterium UWMA 0280 accepting metabolites.pdf

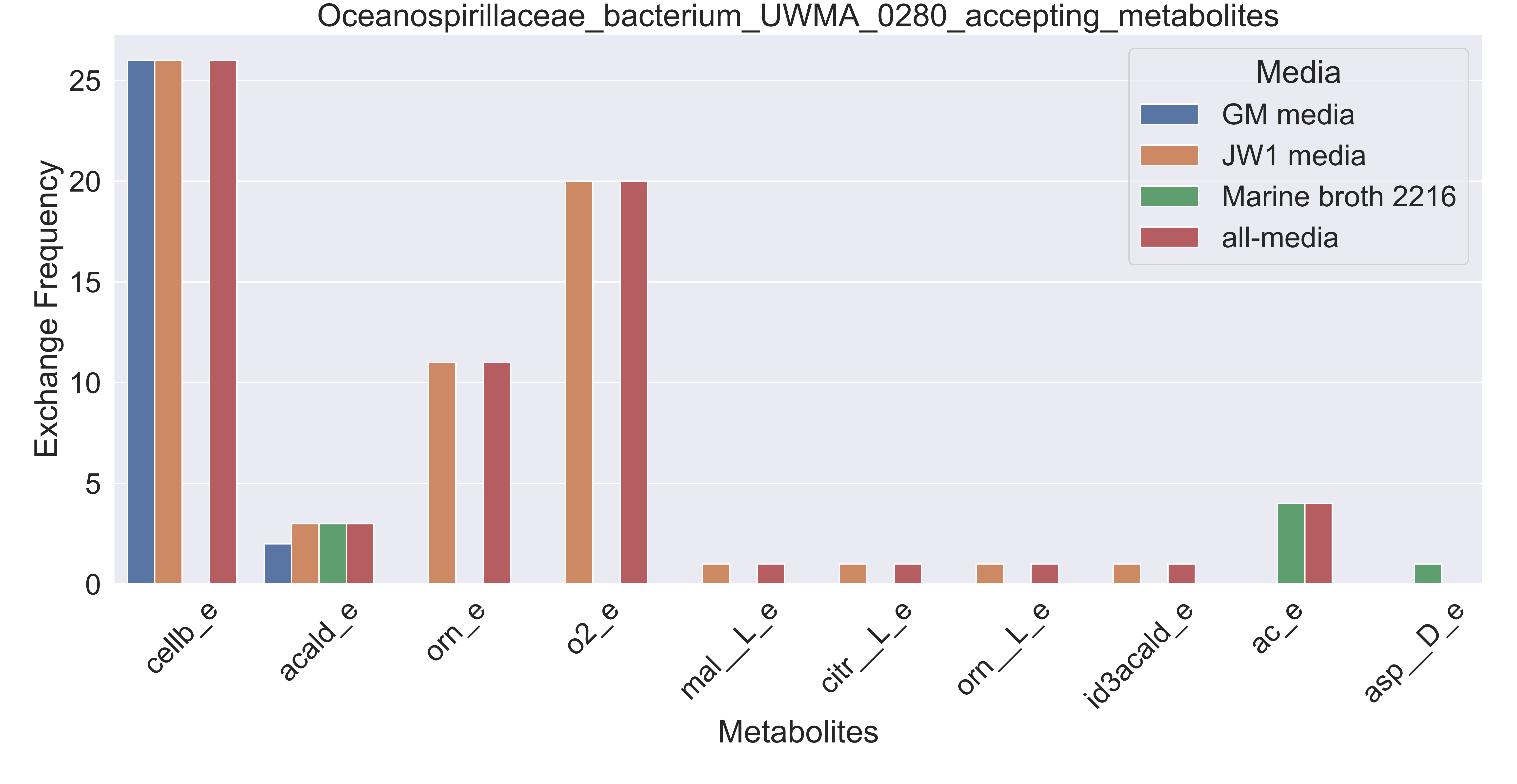

### Supplementary Figure S85 Planctomycetaceae bacterium UWMA 0346 accepting metabolites.pdf

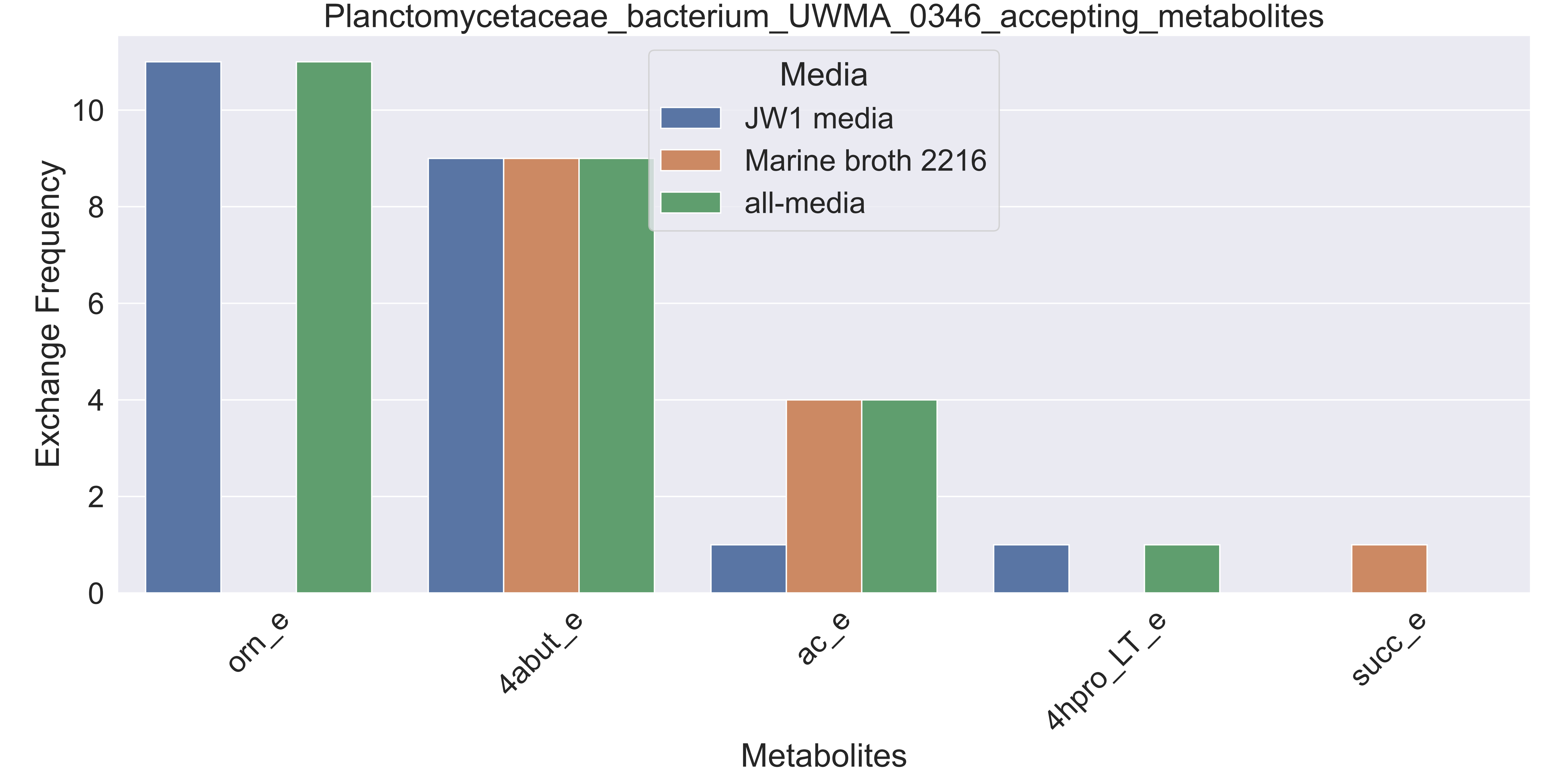

### Supplementary Figure S86 Planctomycetes bacterium UWMA 0265 accepting metabolites.pdf

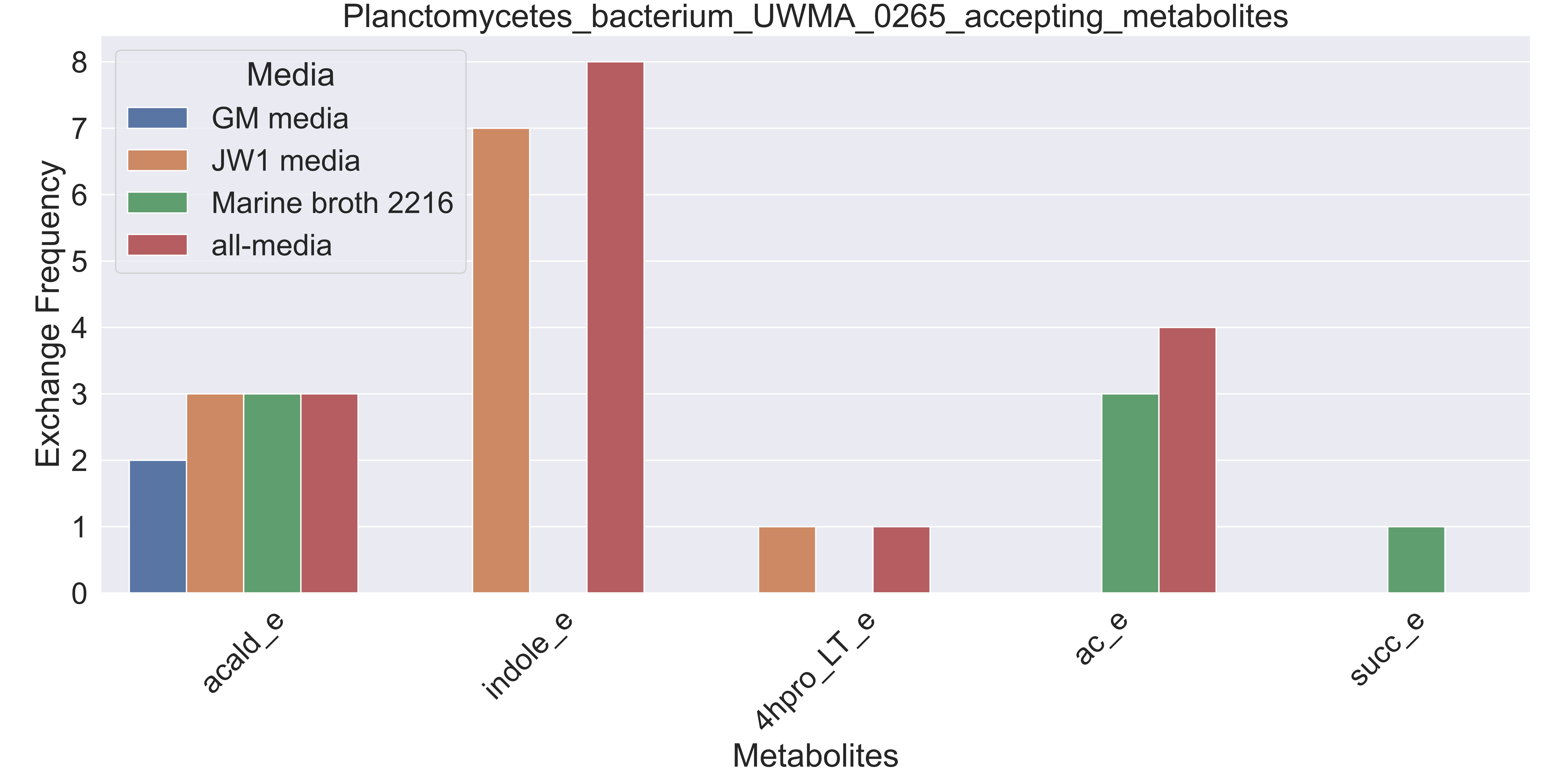

### Supplementary Figure S87 Planctomycetes bacterium UWMA 0276 accepting metabolites.pdf

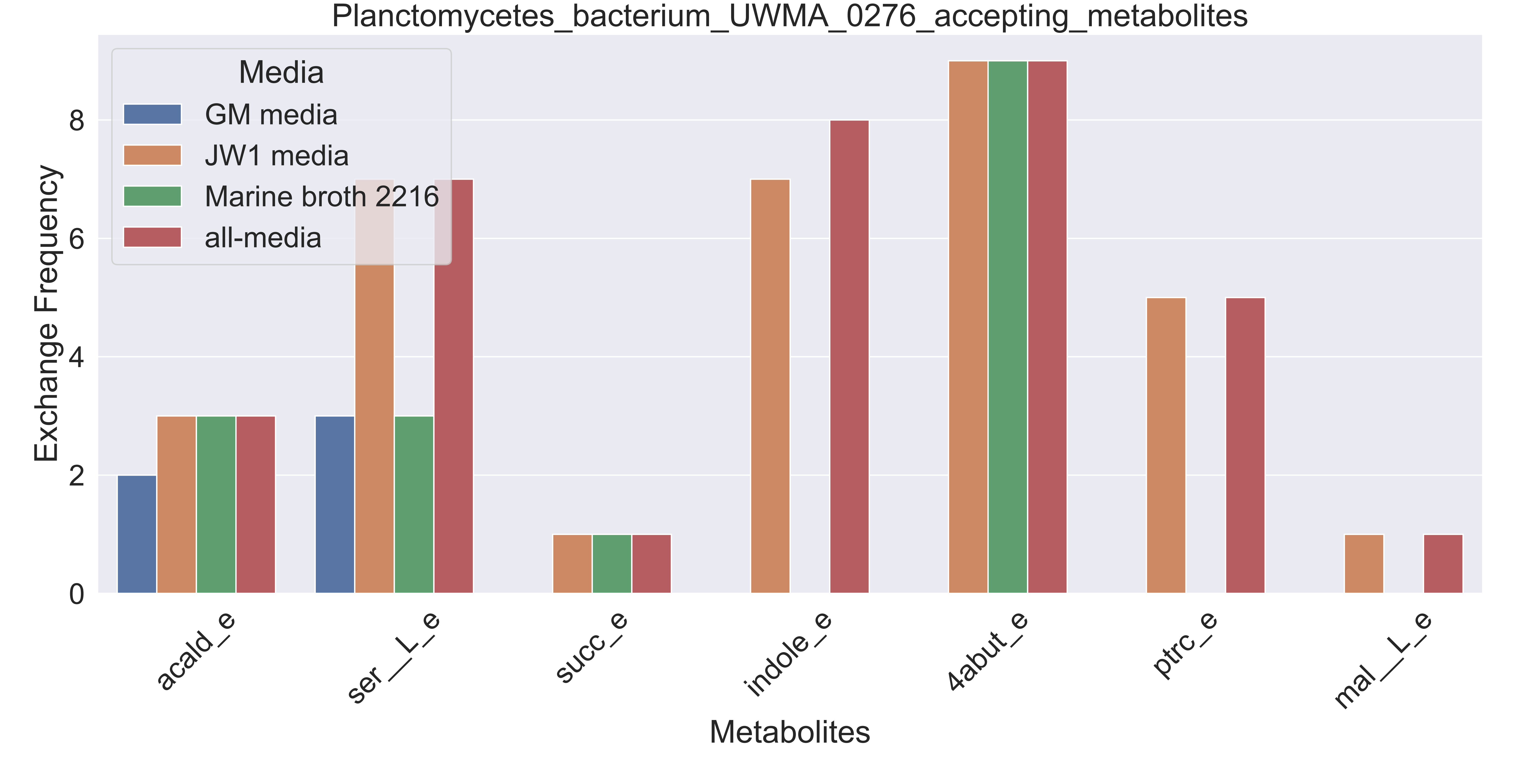

### Supplementary Figure S88 Planctomycetes bacterium UWMA 0294 accepting metabolites.pdf

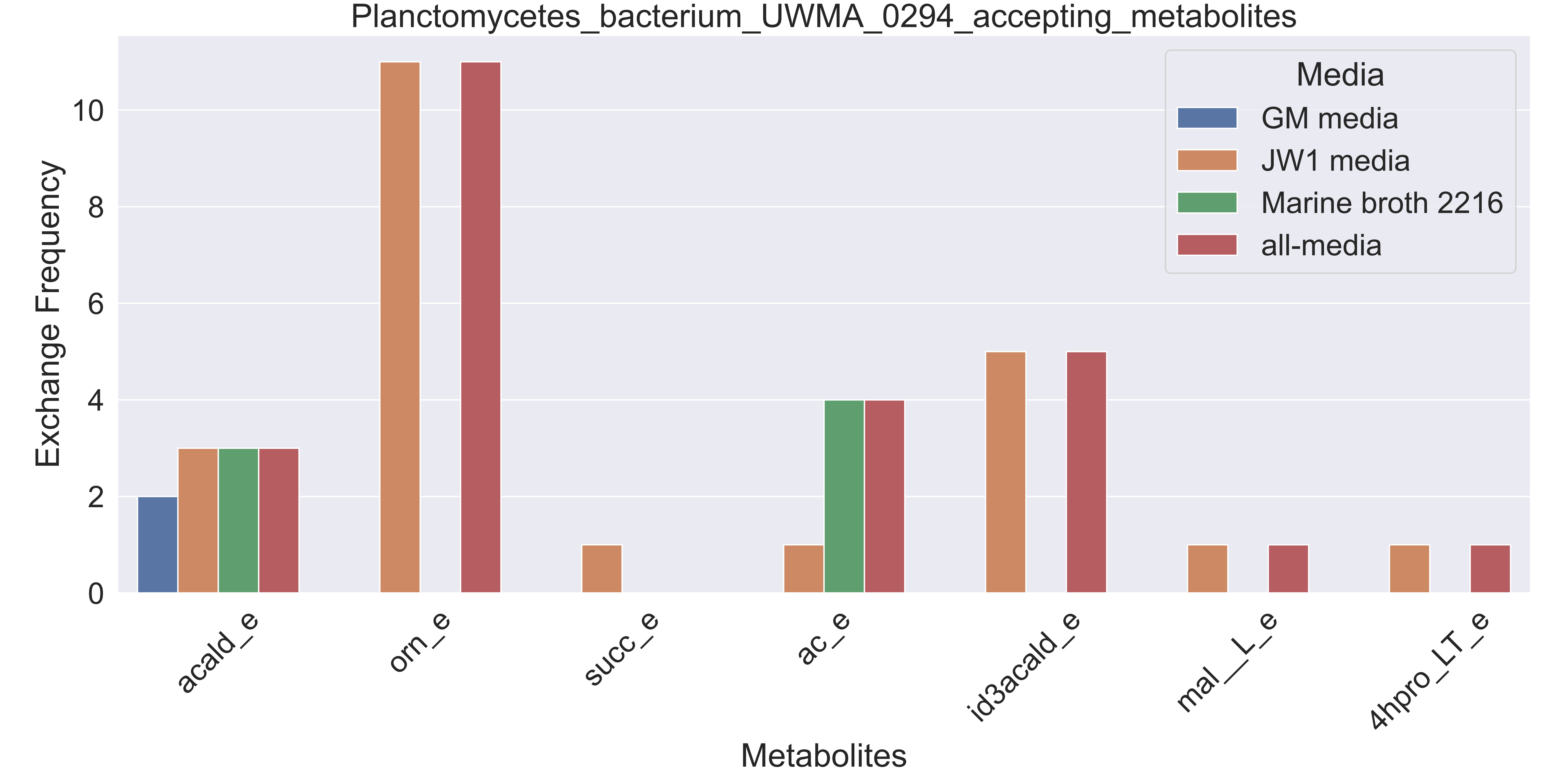

### Supplementary Figure S89 Planctomycetes bacterium UWMA 0329 accepting metabolites.pdf

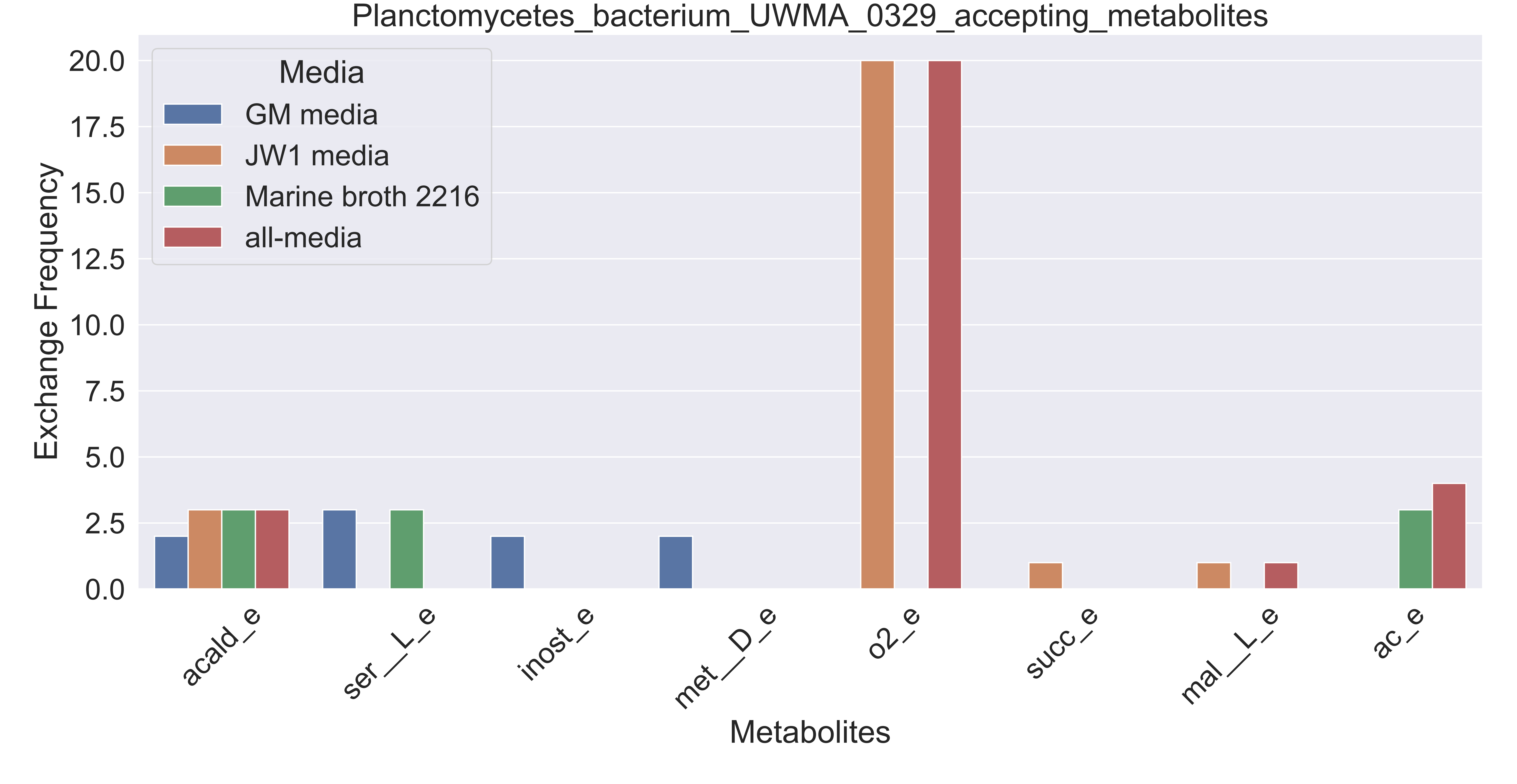

### Supplementary Figure S90 Planctomycetes bacterium UWMA 0345 accepting metabolites.pdf

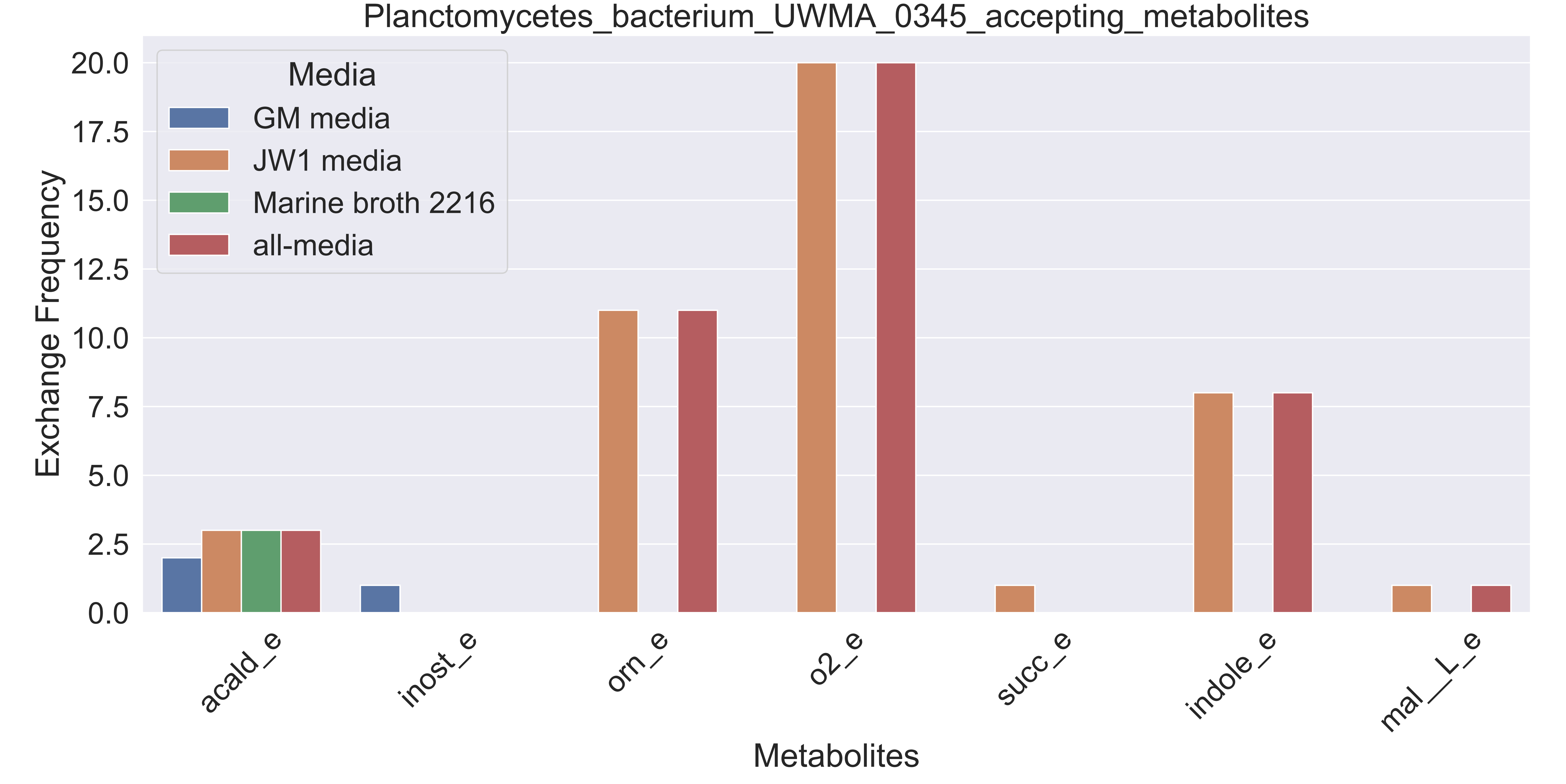

### Supplementary Figure S91 Porticoccaceae bacterium UWMA 0313 accepting metabolites.pdf

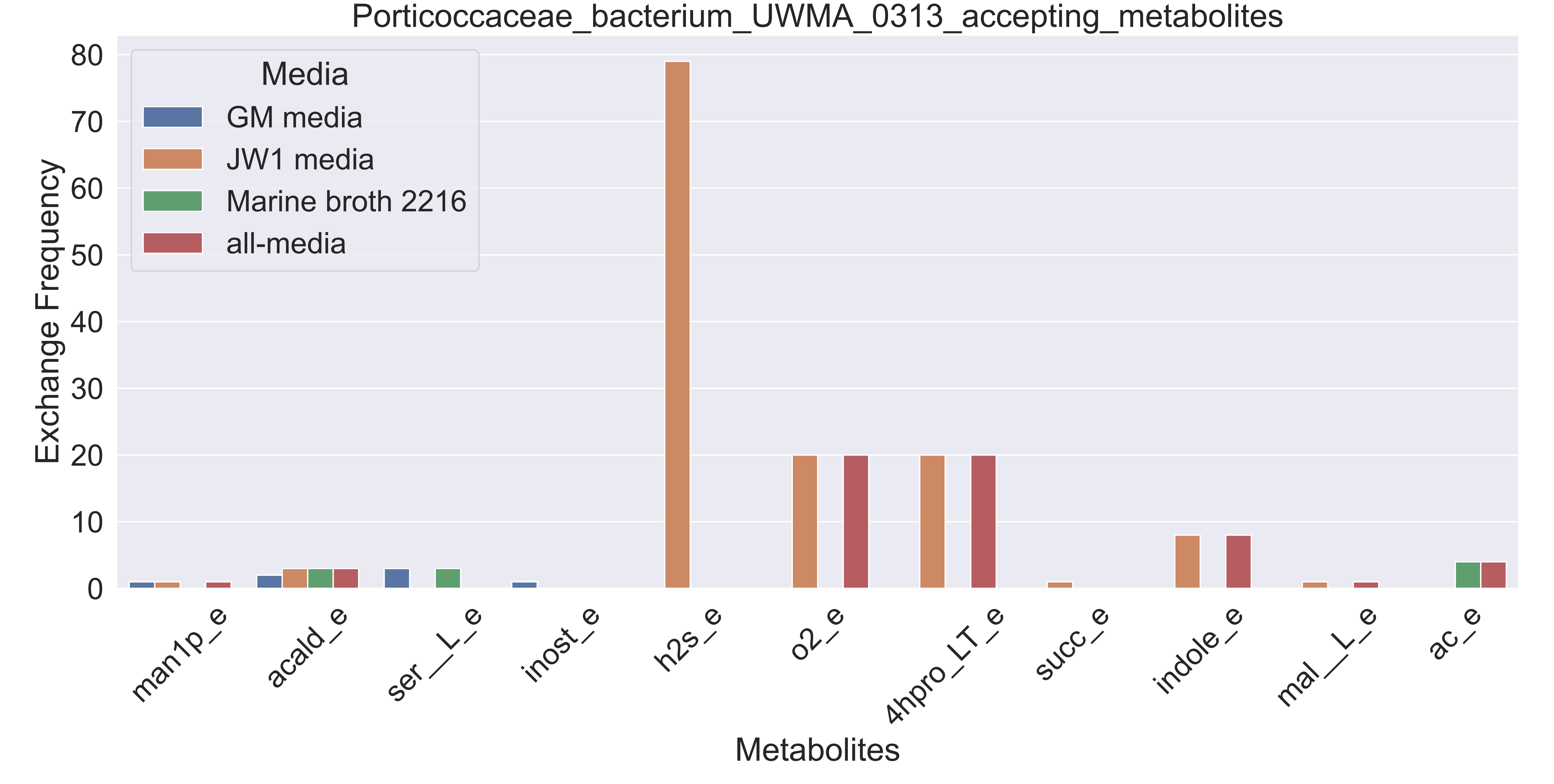

### Supplementary Figure S92 Rhodospirillales bacterium UWMA 0295 accepting metabolites.pdf

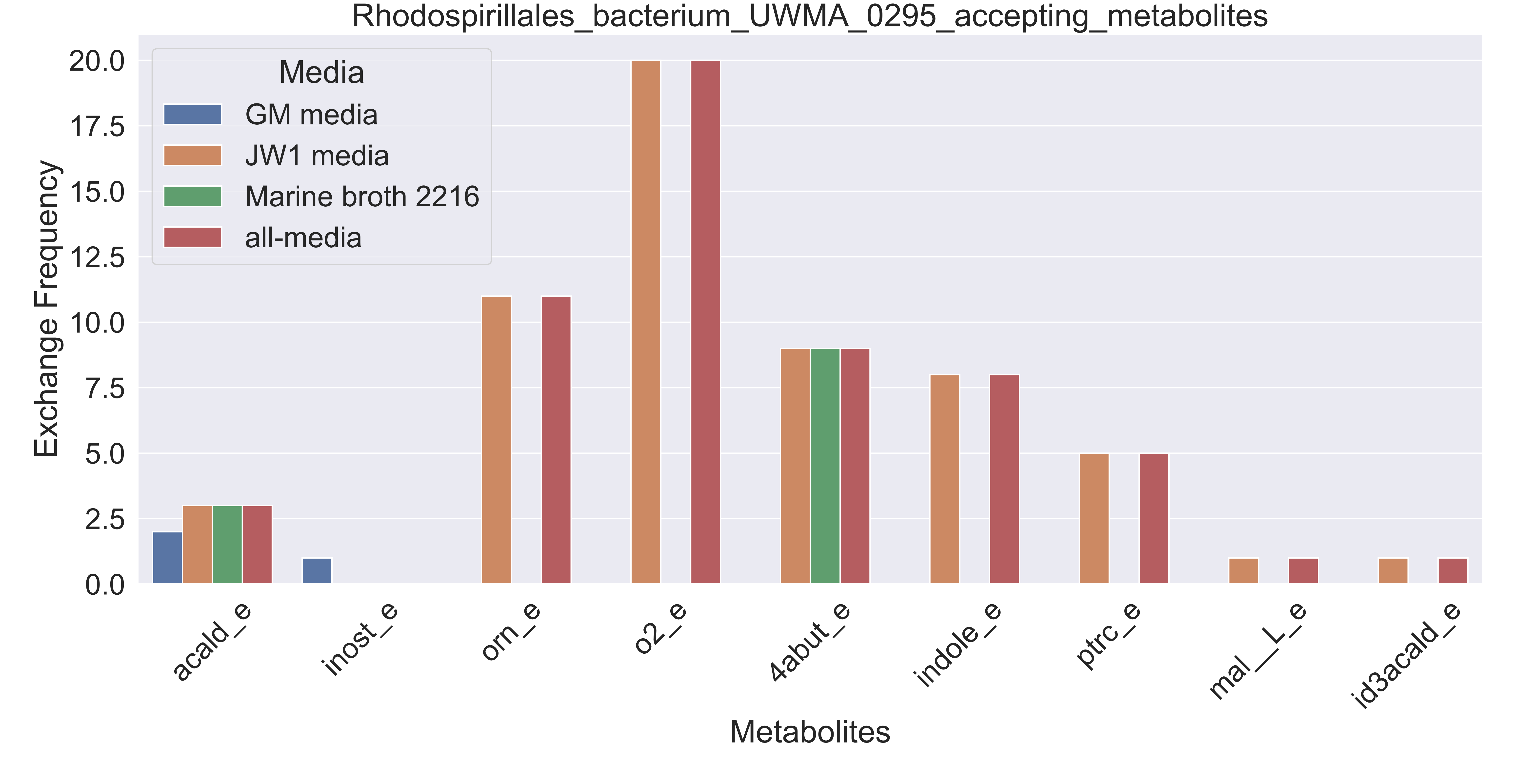

### Supplementary Figure S93 Sneathiellales bacterium UWMA 0353 accepting metabolites.pdf

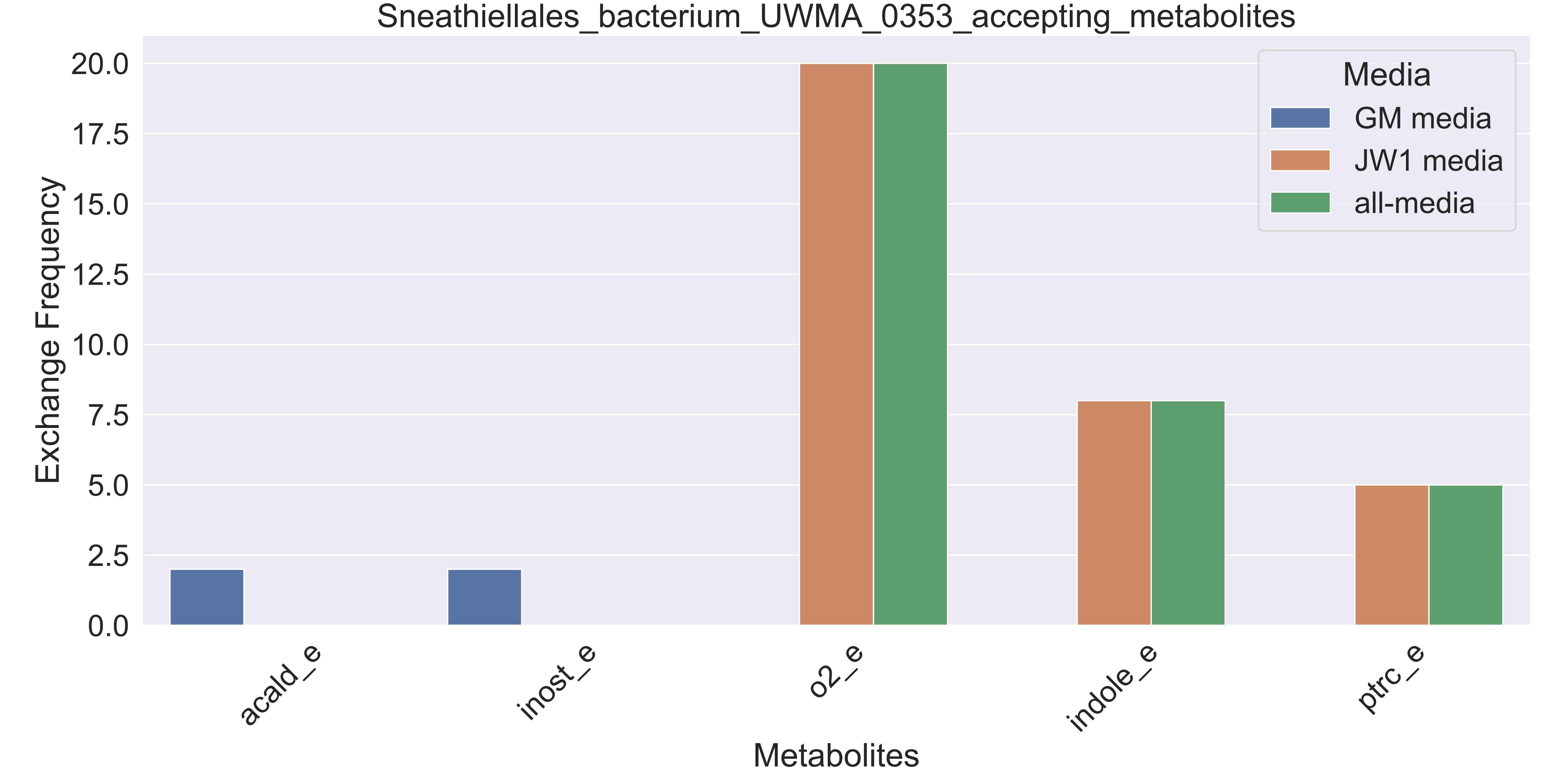

### Supplementary Figure S94 Sulfitobacter sp UWMA 0305 accepting metabolites.pdf

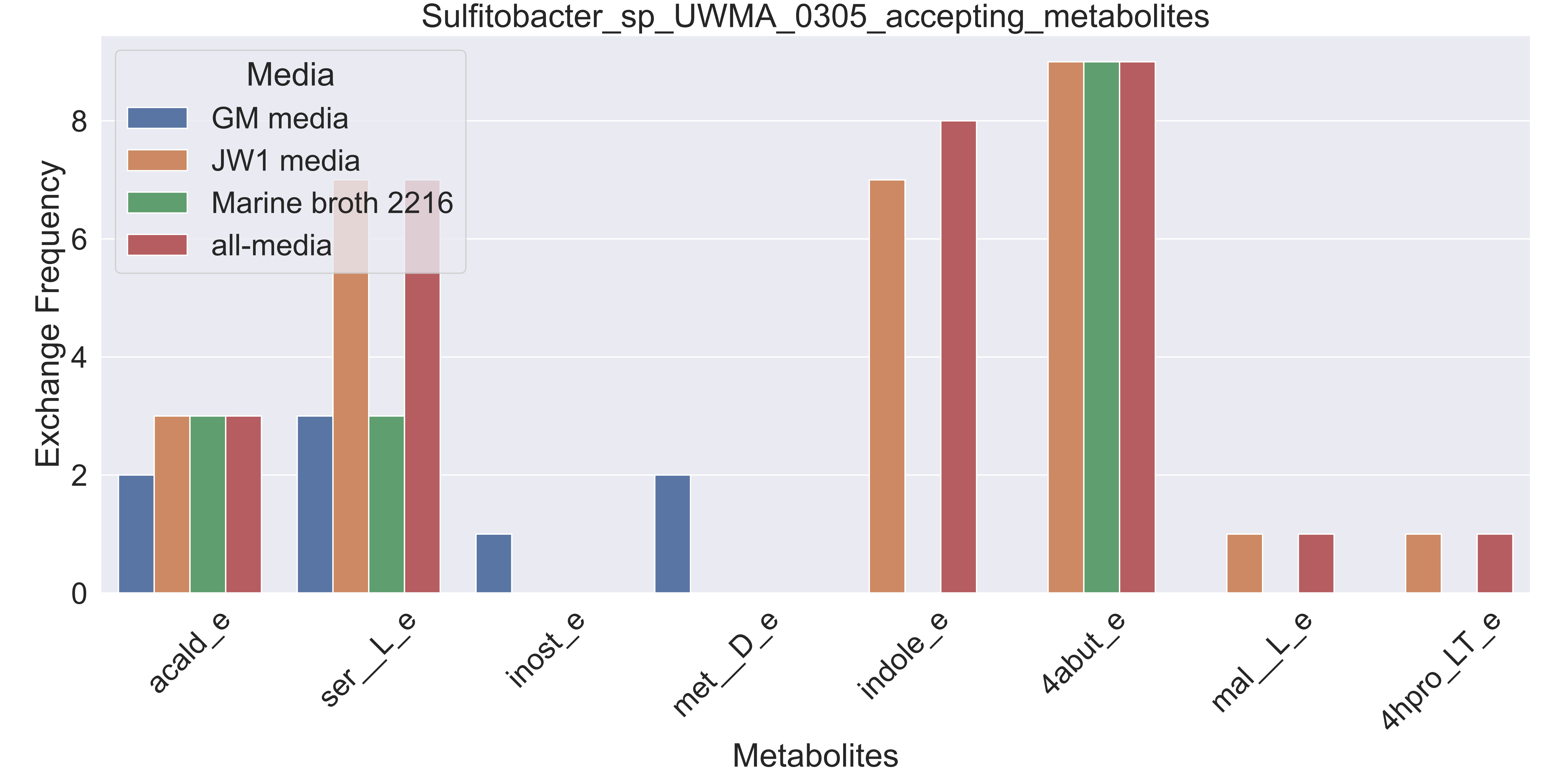

### Supplementary Figure S95 Thiotrichaceae bacterium UWMA 0311 accepting metabolites.pdf

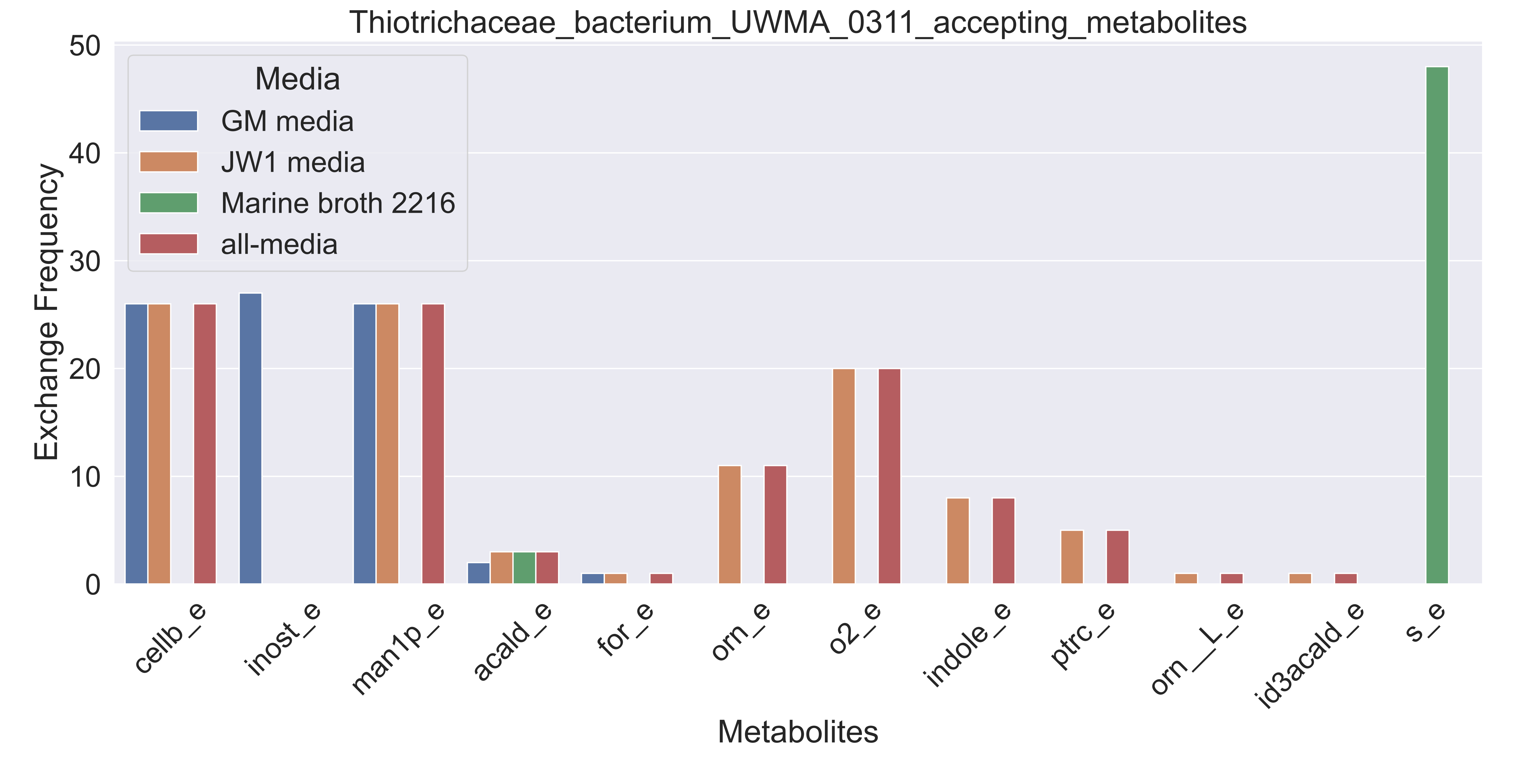

### Supplementary Figure S96 Verrucomicrobiales bacterium UWMA 0292 accepting metabolites.pdf

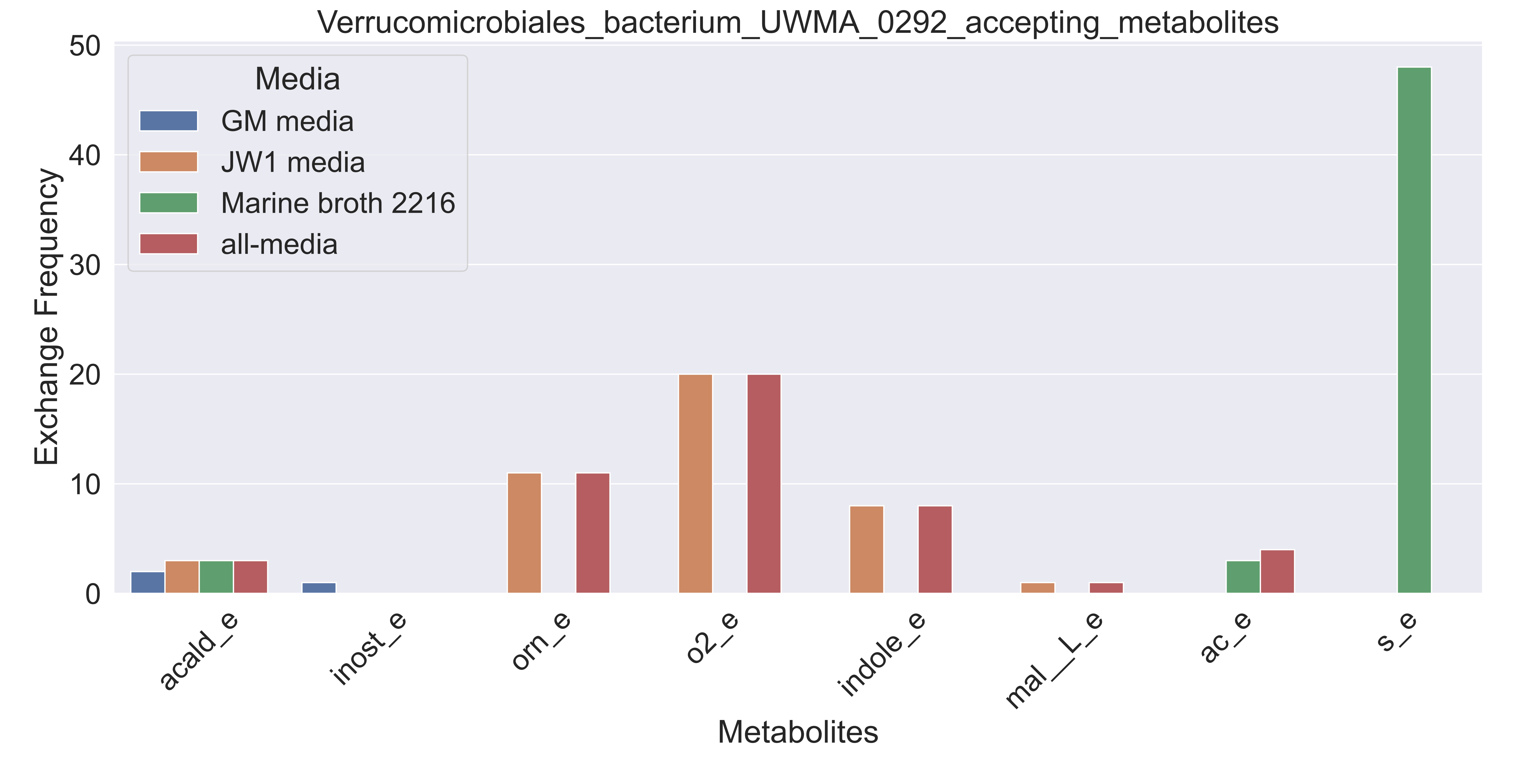

### Supplementary Figure S97 Verrucomicrobiales bacterium UWMA 0320 accepting metabolites.pdf

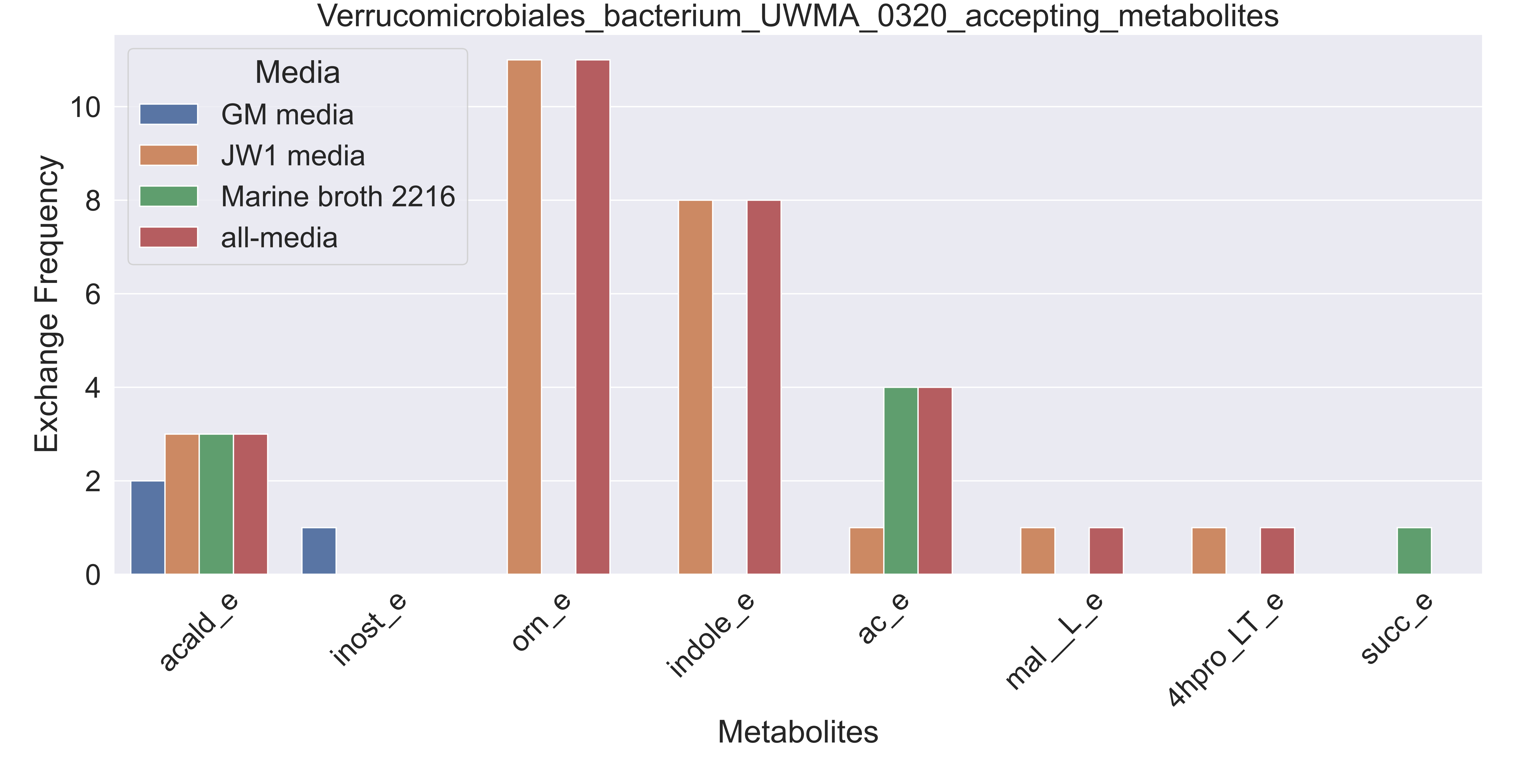

### Supplementary Figure S98 Verrucomicrobiales bacterium UWMA 0332 accepting metabolites.pdf

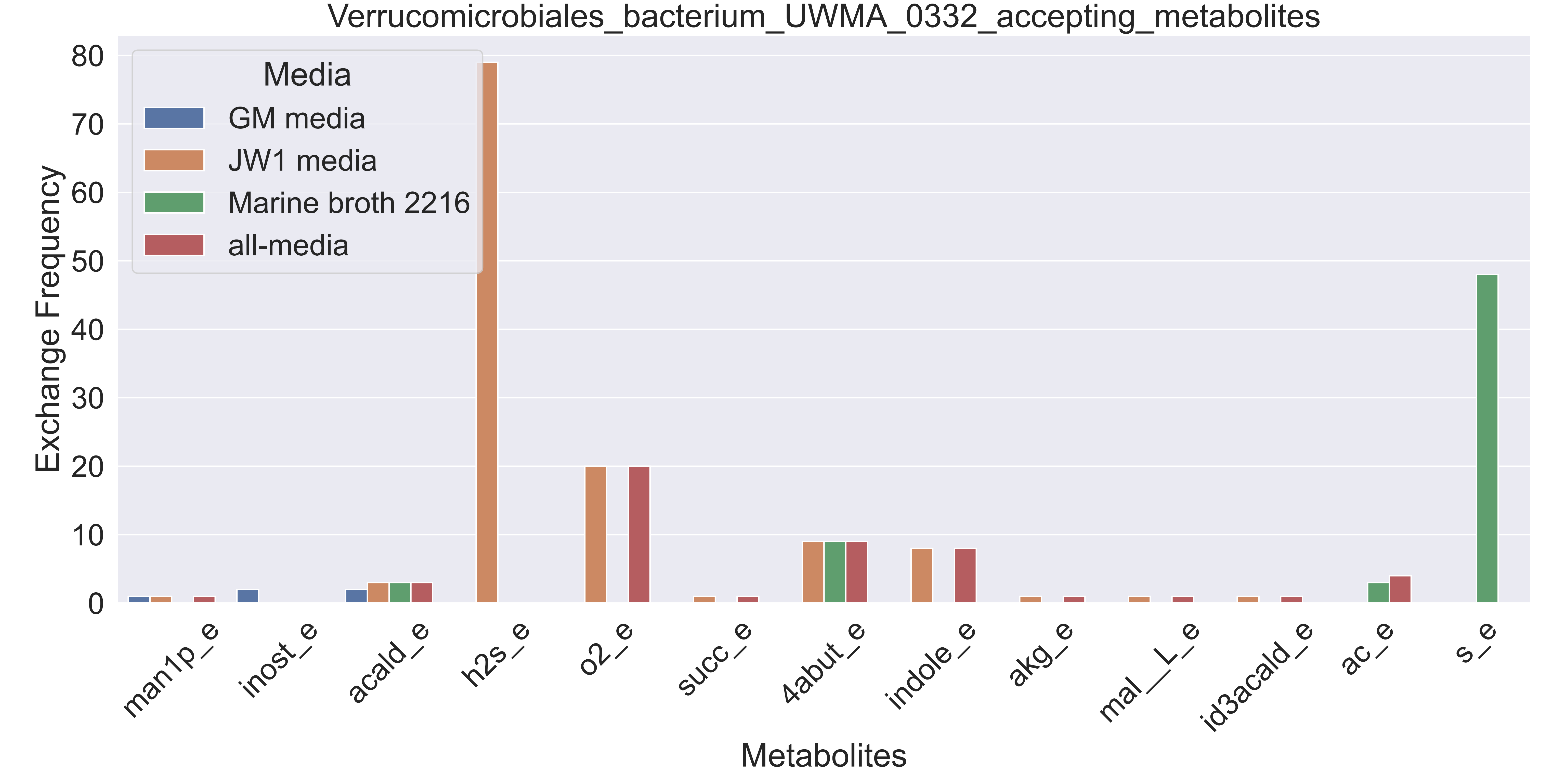

### Supplementary Figure S100 Acidimicrobiia bacterium UWMA 0303 donating metabolites.pdf

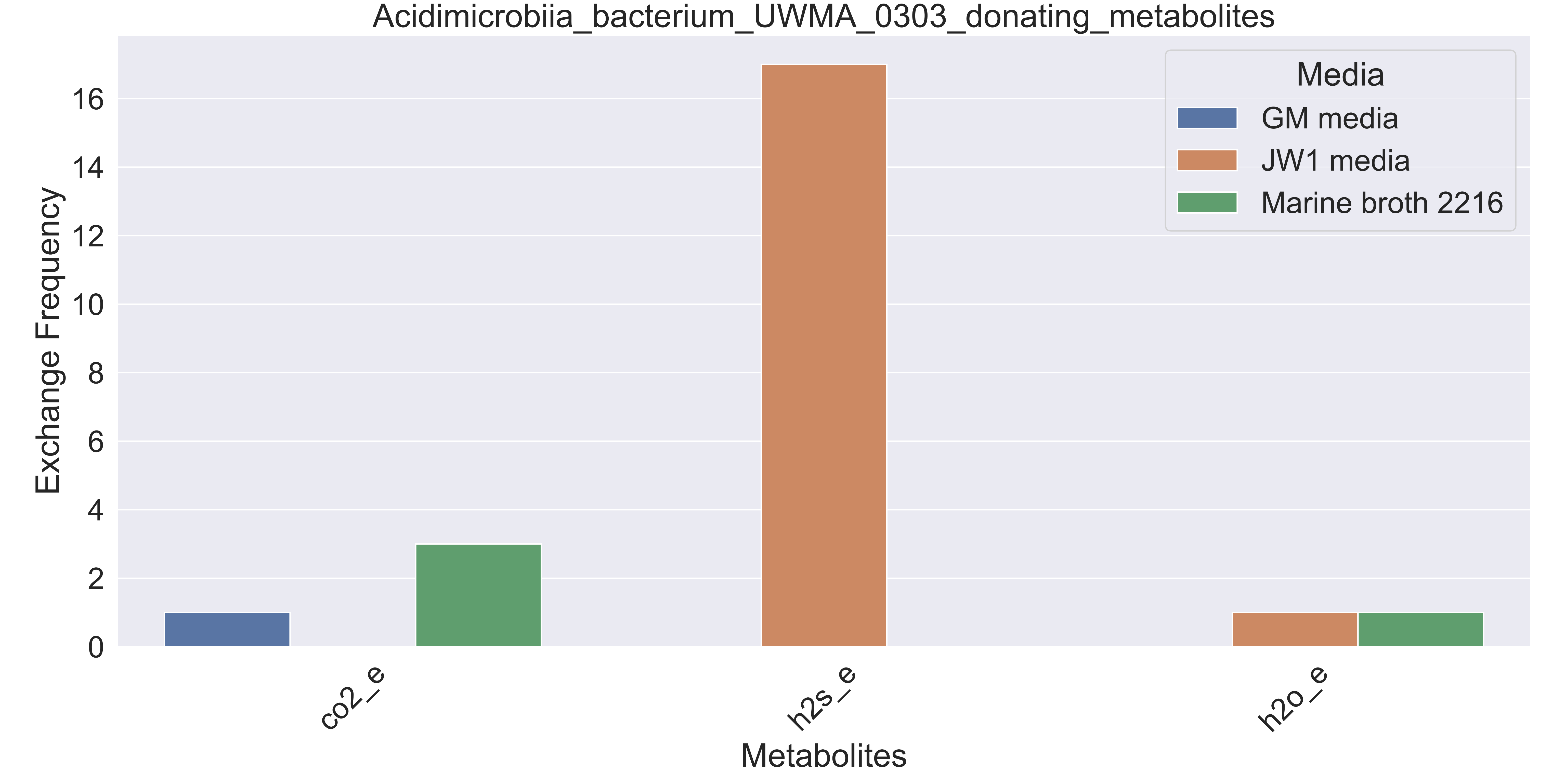

### Supplementary Figure S101 Acidimicrobiia bacterium UWMA 0319 donating metabolites.pdf

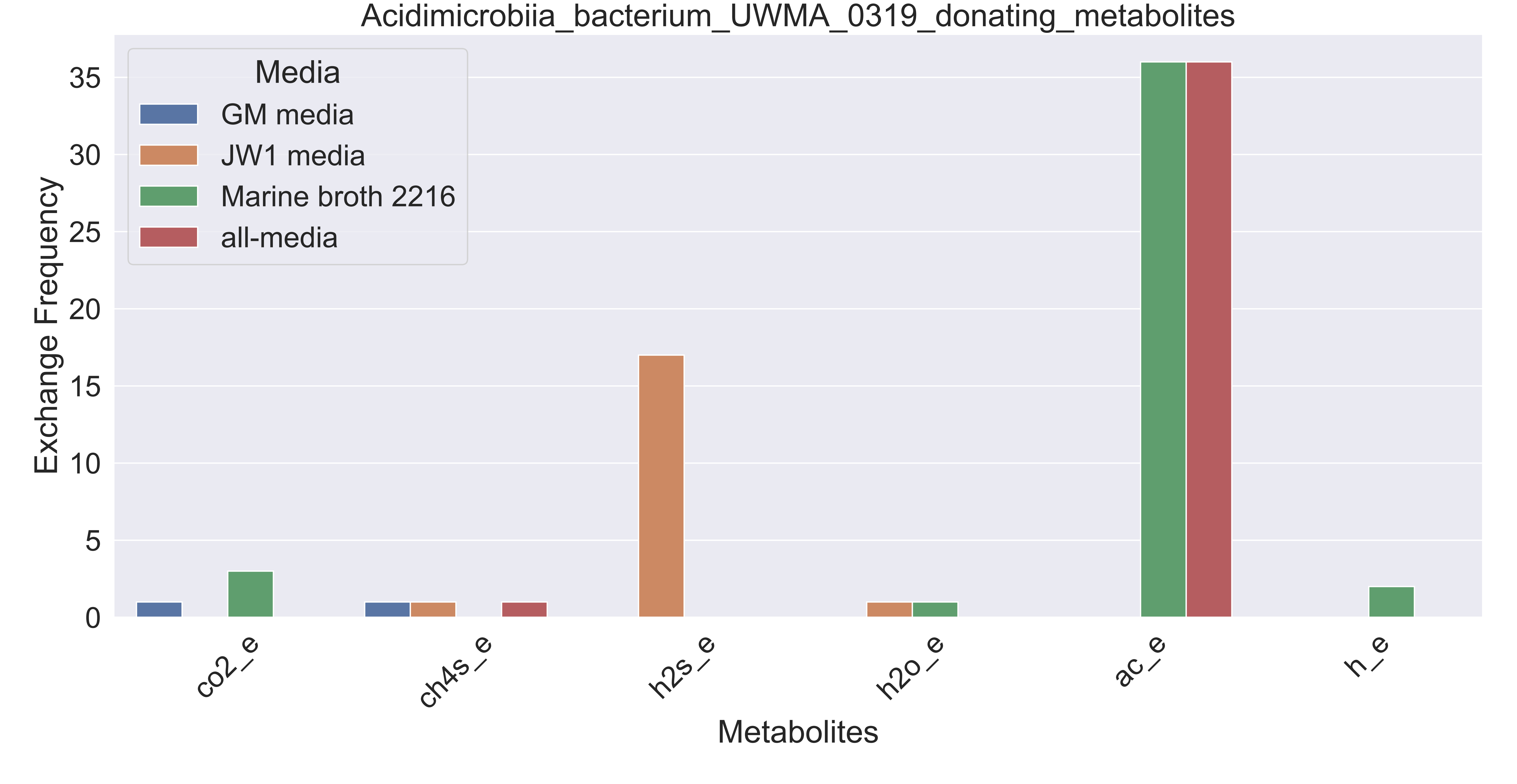
