## Supplementary figures and images for "Metagenome-based metabolic modelling predicts unique microbial interactions in deep-sea hydrothermal plume microbiomes"

### Supplementary Figure S99 Acidimicrobiia bacterium UWMA 0264 donating metabolites.pdf

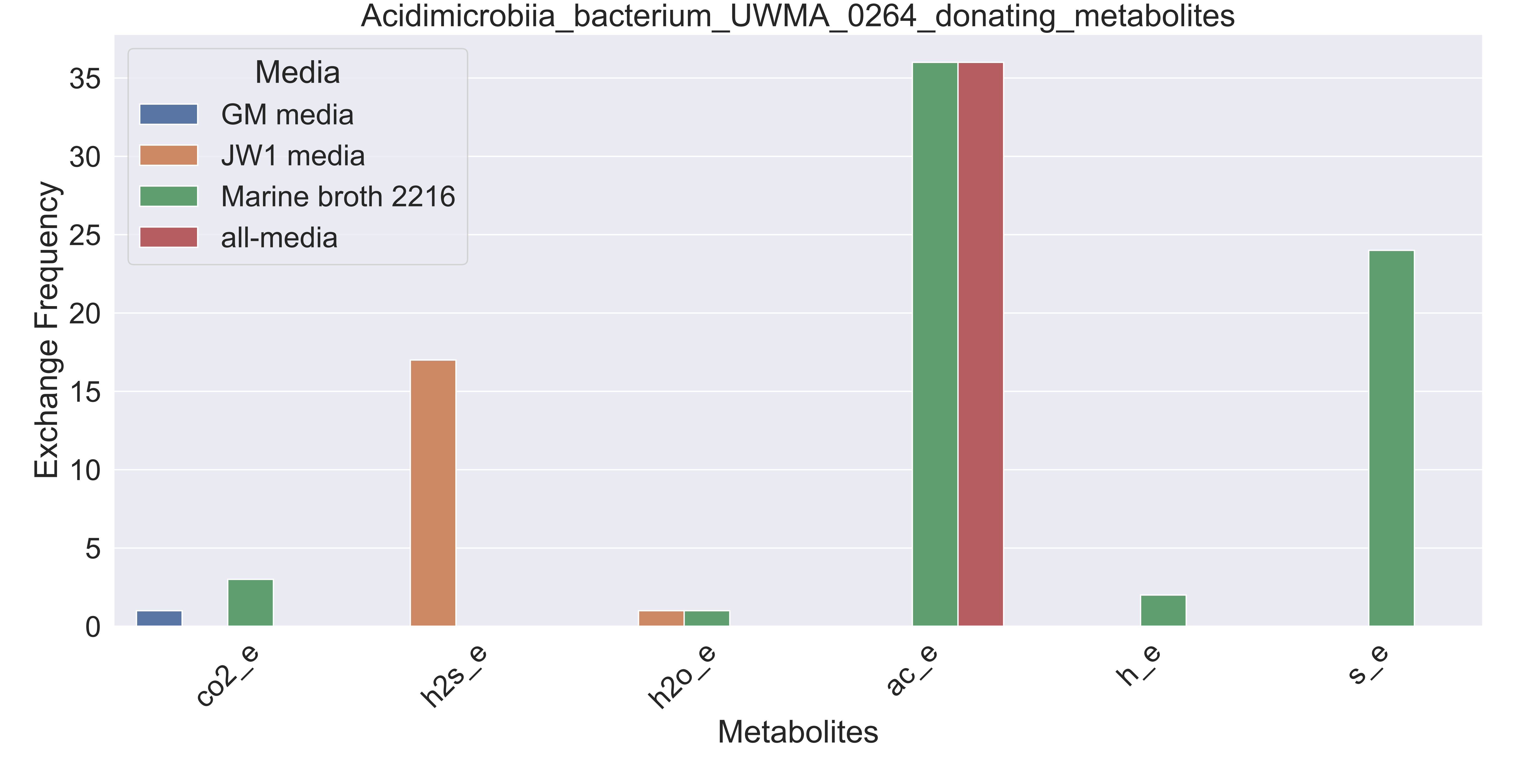

### Supplementary Figure S102 Acidobacteria bacterium UWMA 0355 donating metabolites.pdf

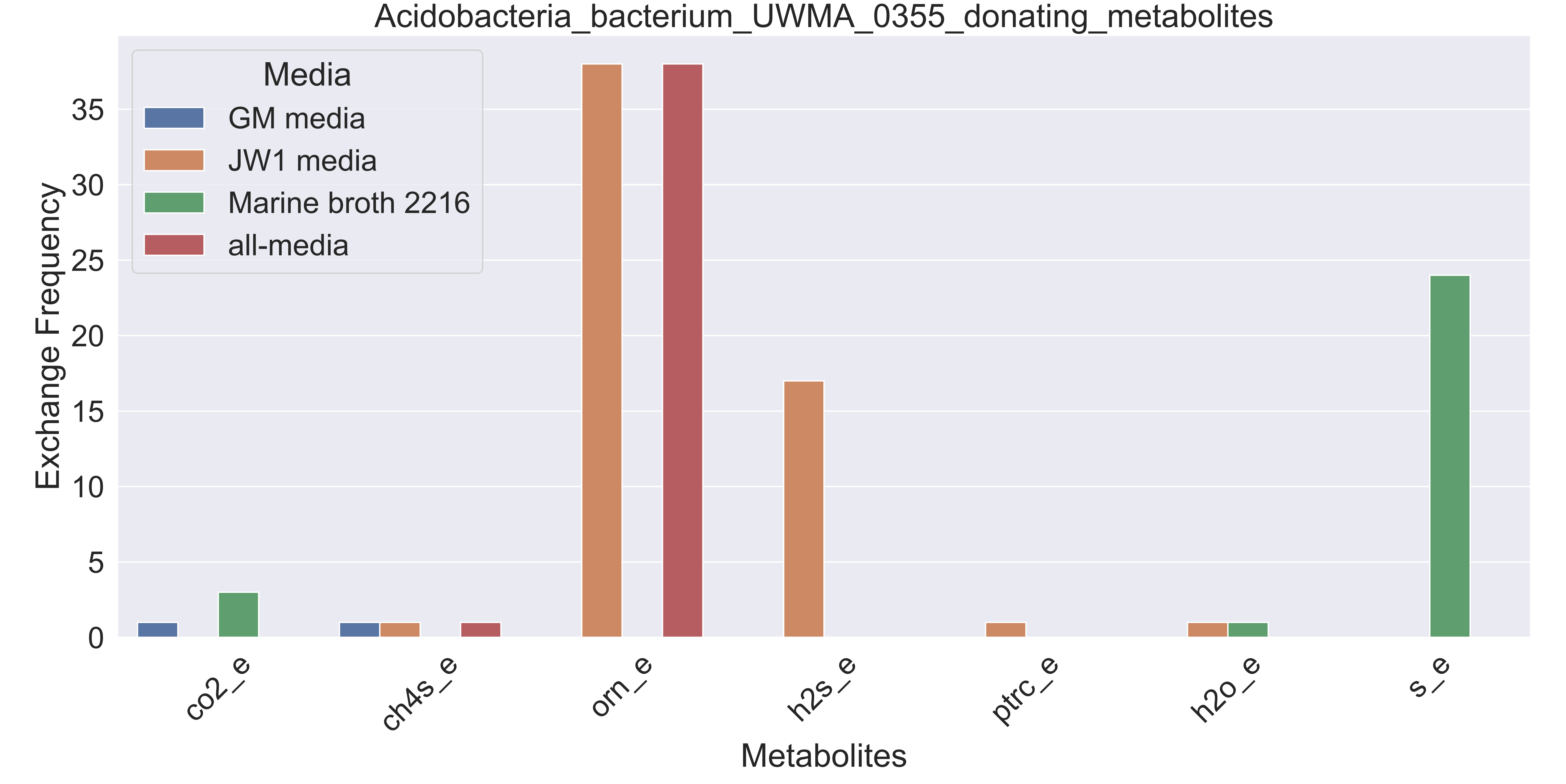

### Supplementary Figure S103 Alphaproteobacteria bacterium UWMA 0321 donating metabolites.pdf

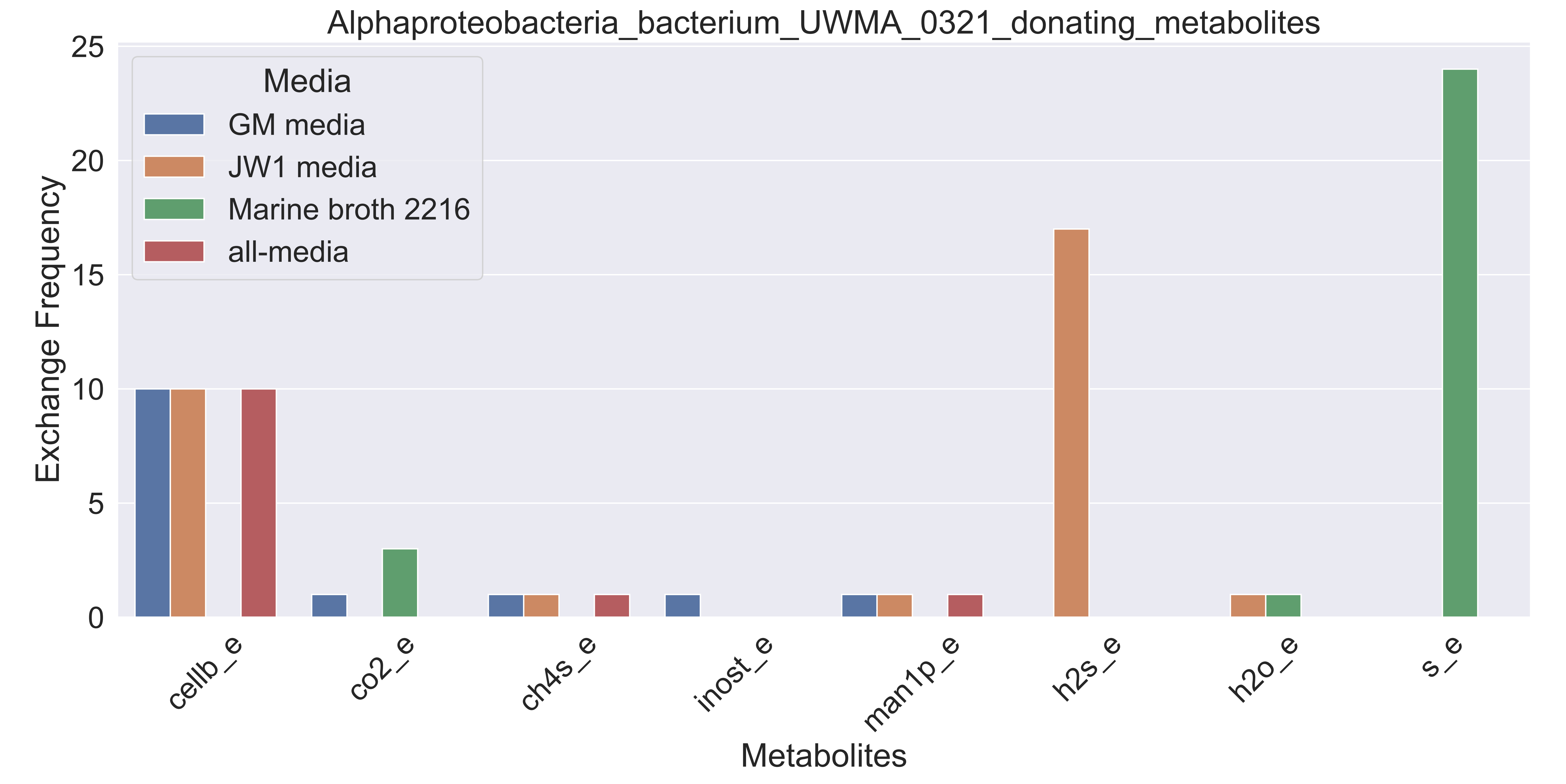

### Supplementary Figure S104 Bacteroidetes bacterium UWMA 0293 donating metabolites.pdf

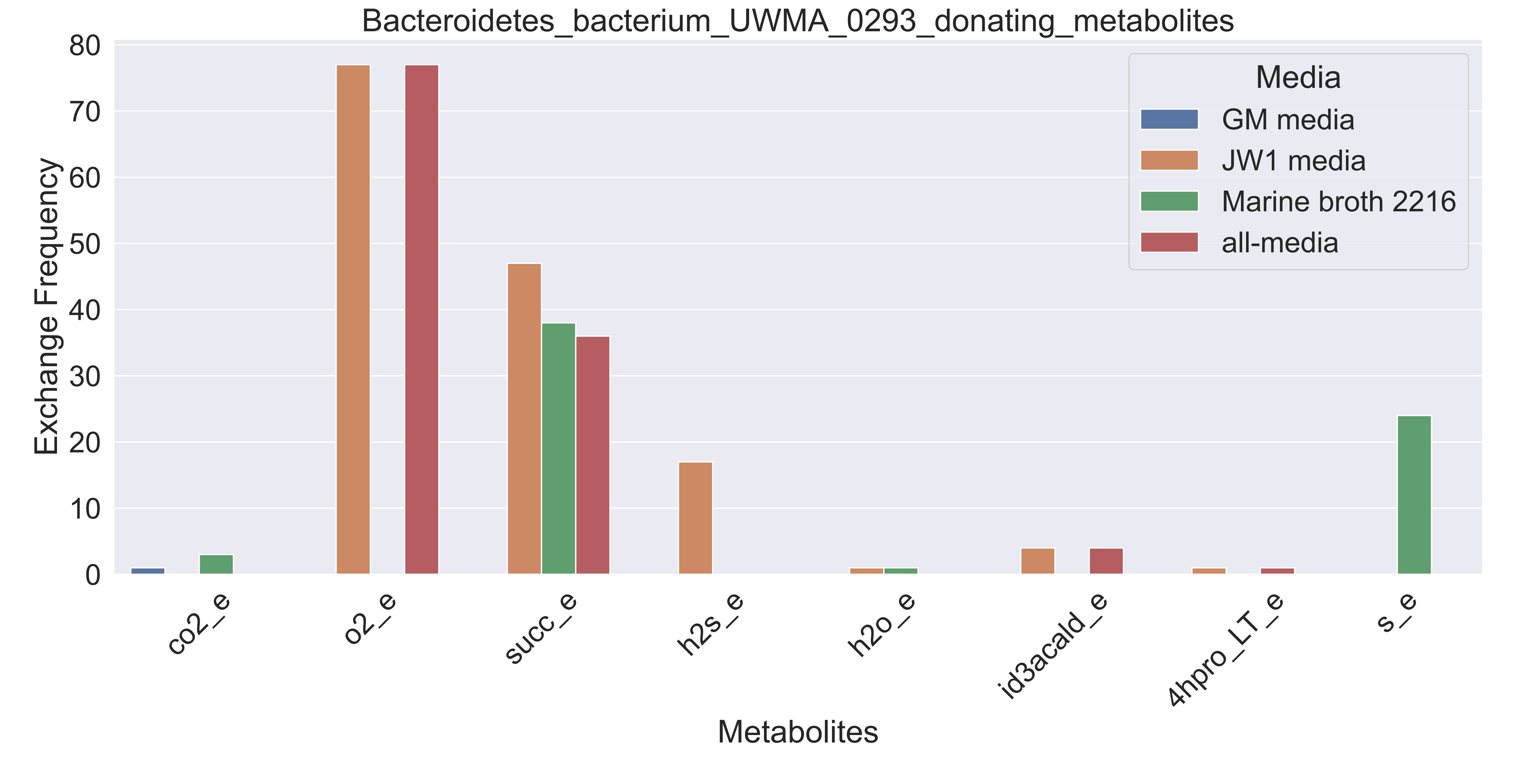

### Supplementary Figure S105 Bacteroidetes bacterium UWMA 0358 donating metabolites.pdf

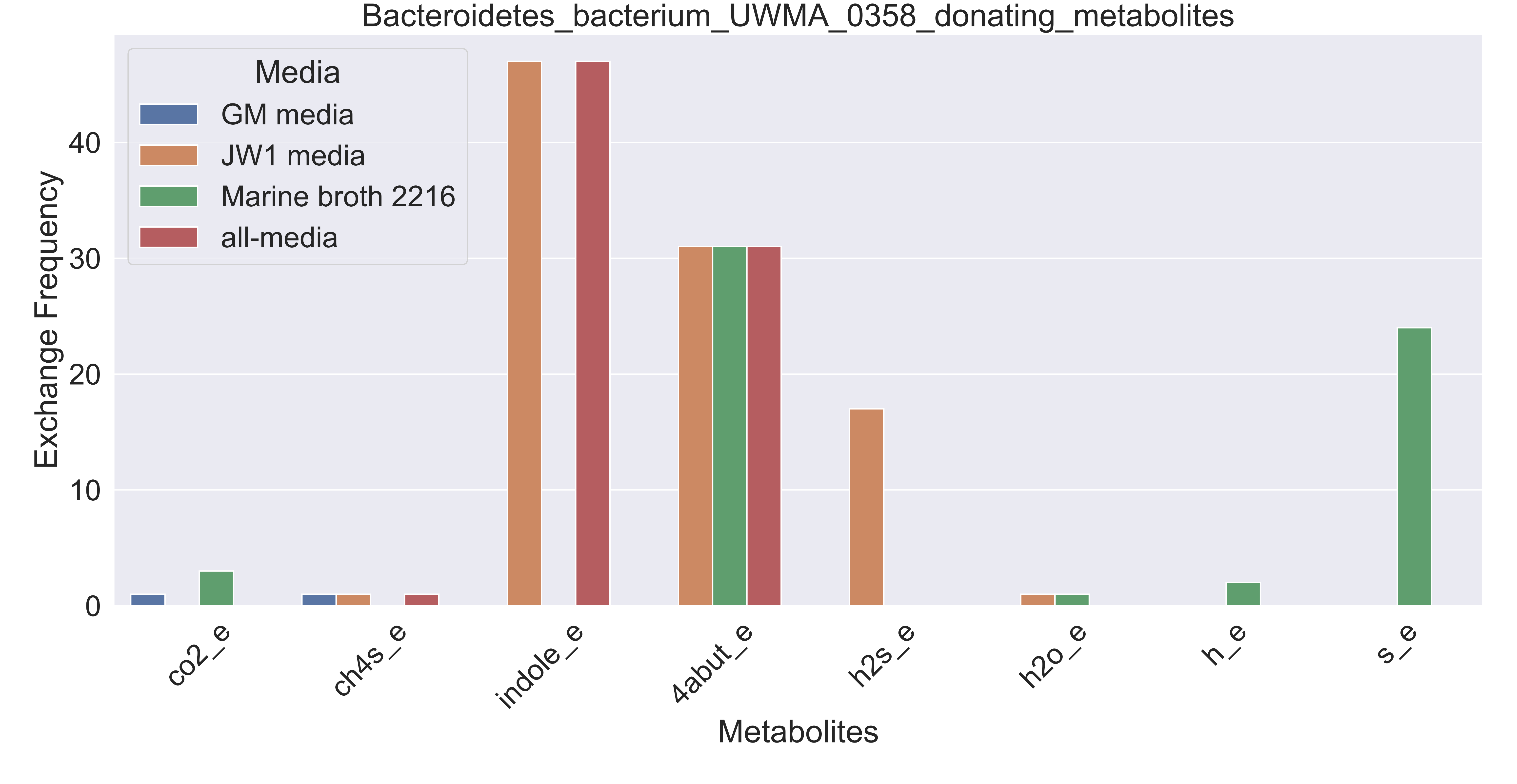

### Supplementary Figure S106 candidate division UBP10 bacterium UWMA 0304 donating metabolites.pdf

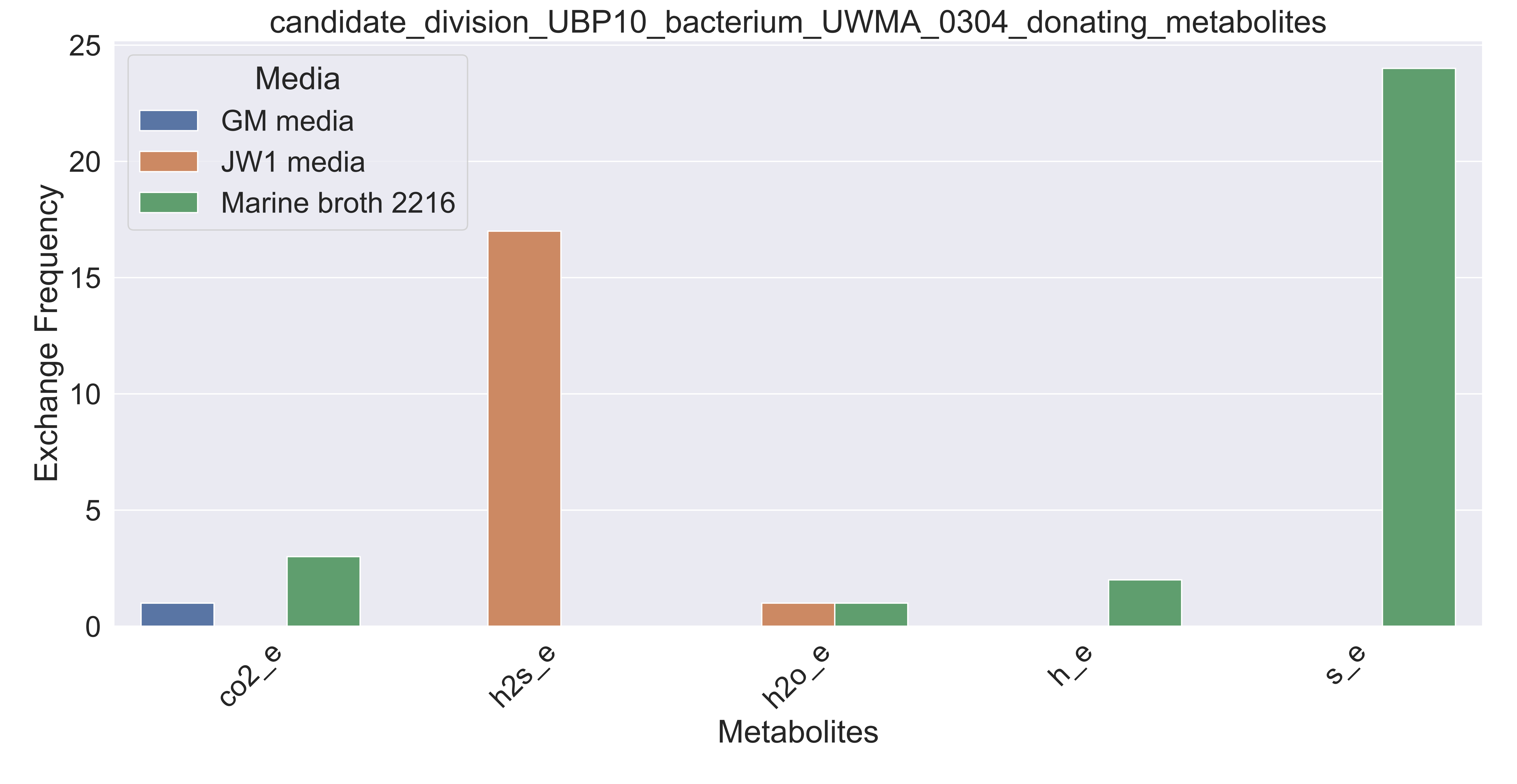

### Supplementary Figure S107 candidate division Zixibacteria bacterium UWMA 0288 donating metabolites.pdf

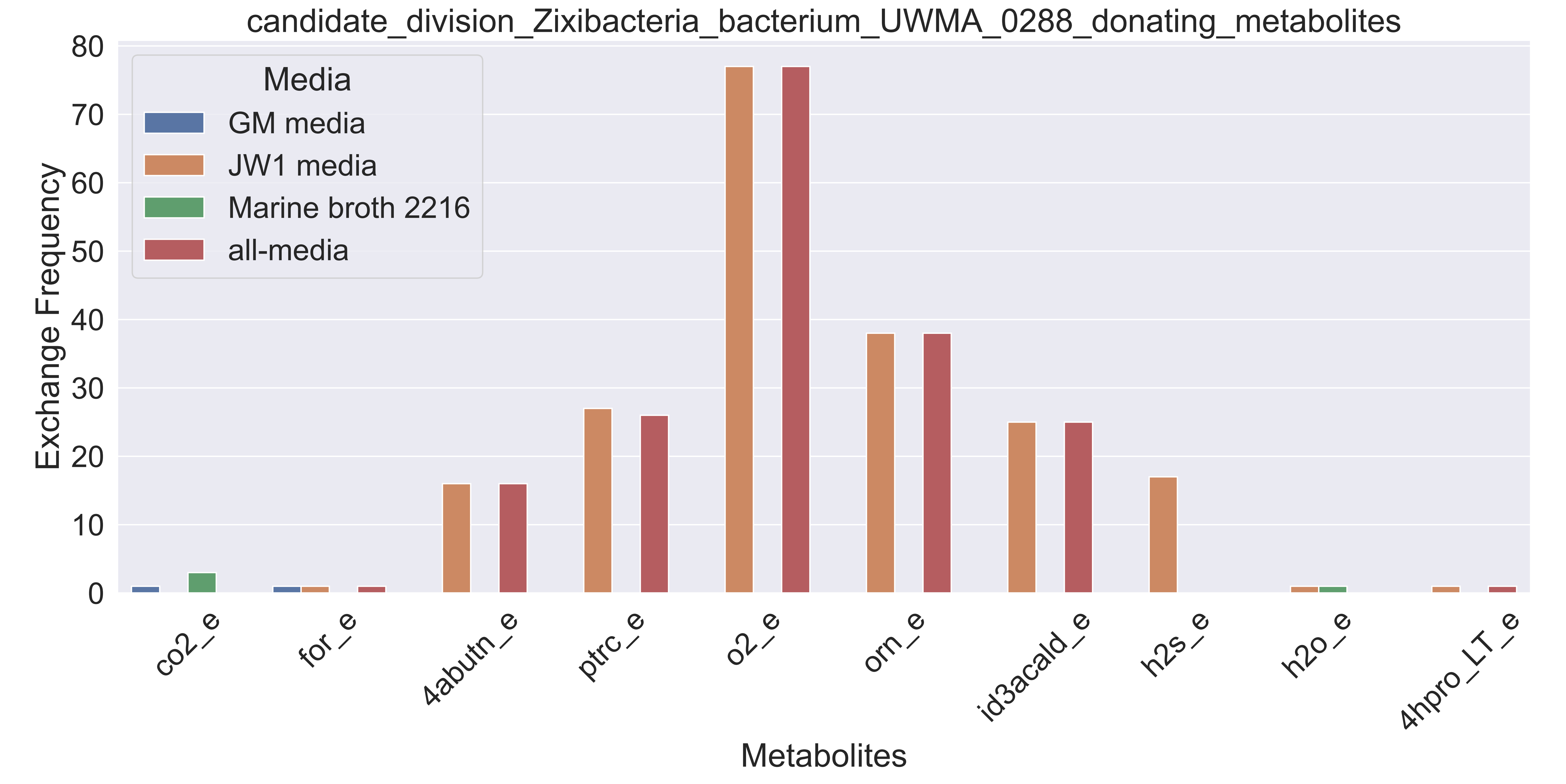

### Supplementary Figure S108 Candidatus Handelsmanbacteria bacterium UWMA 0286 donating metabolites.pdf

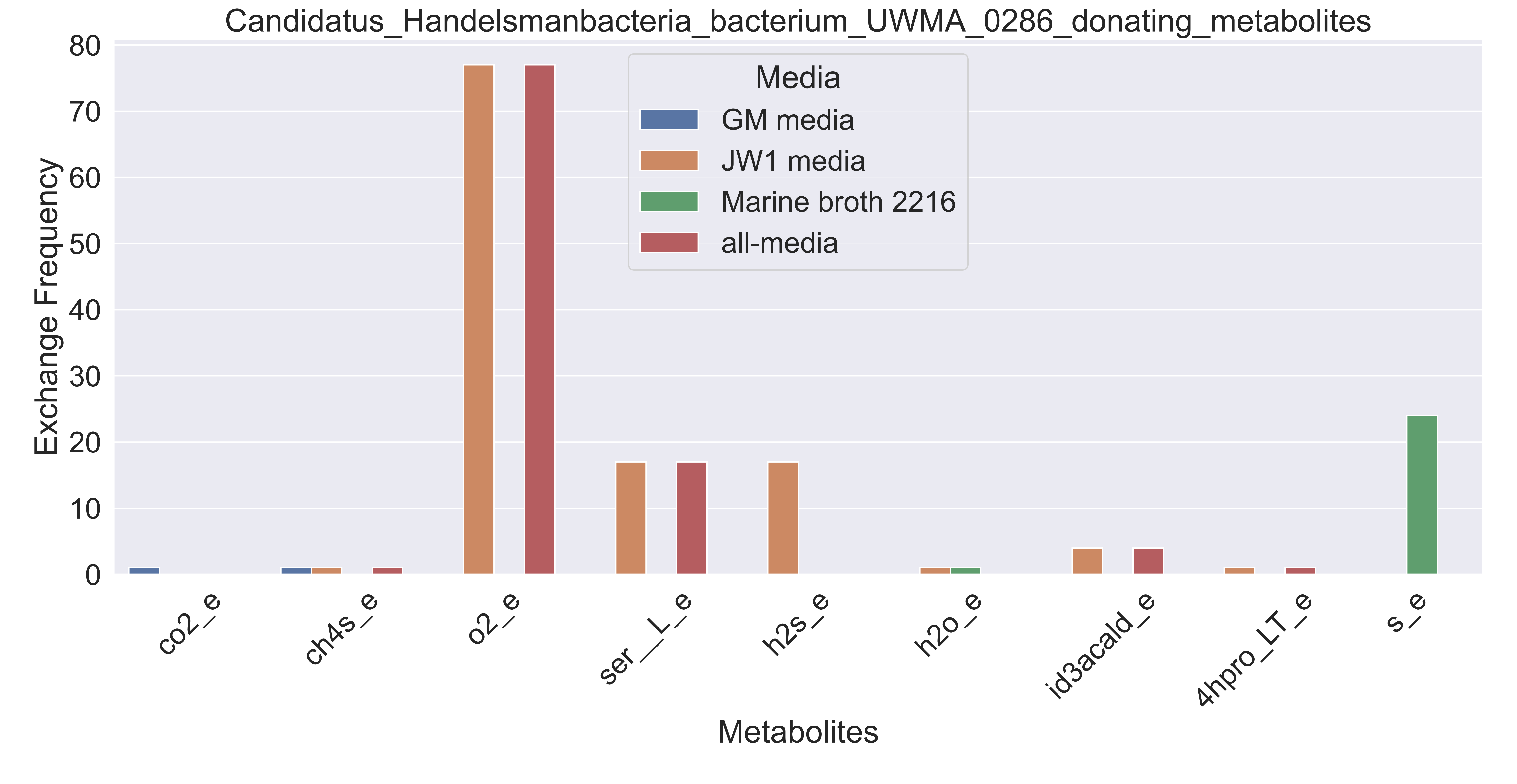

### Supplementary Figure S109 Candidatus Handelsmanbacteria bacterium UWMA 0300 donating metabolites.pdf

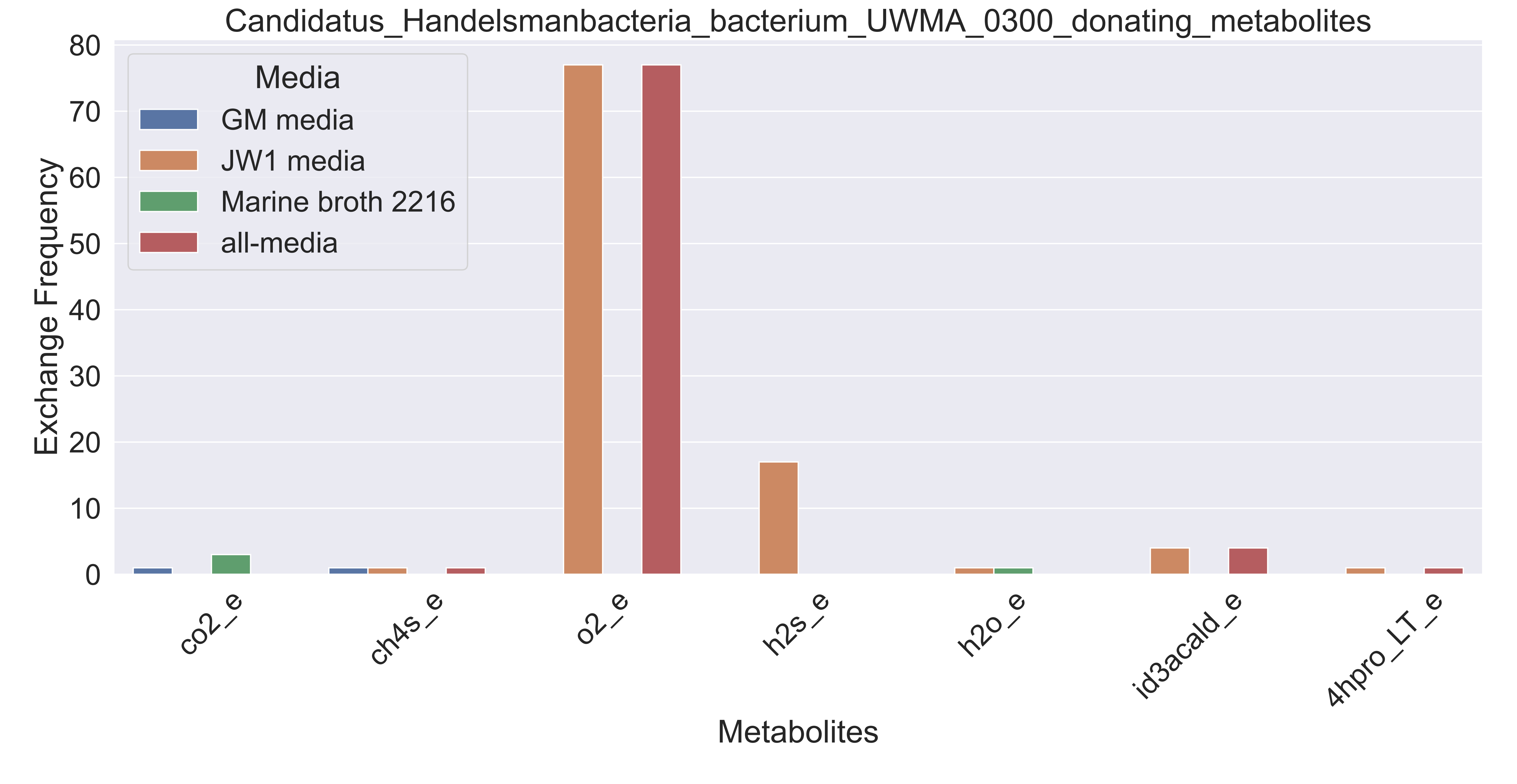

### Supplementary Figure S110 Candidatus Lambdaproteobacteria bacterium UWMA 0298 donating metabolites.pdf

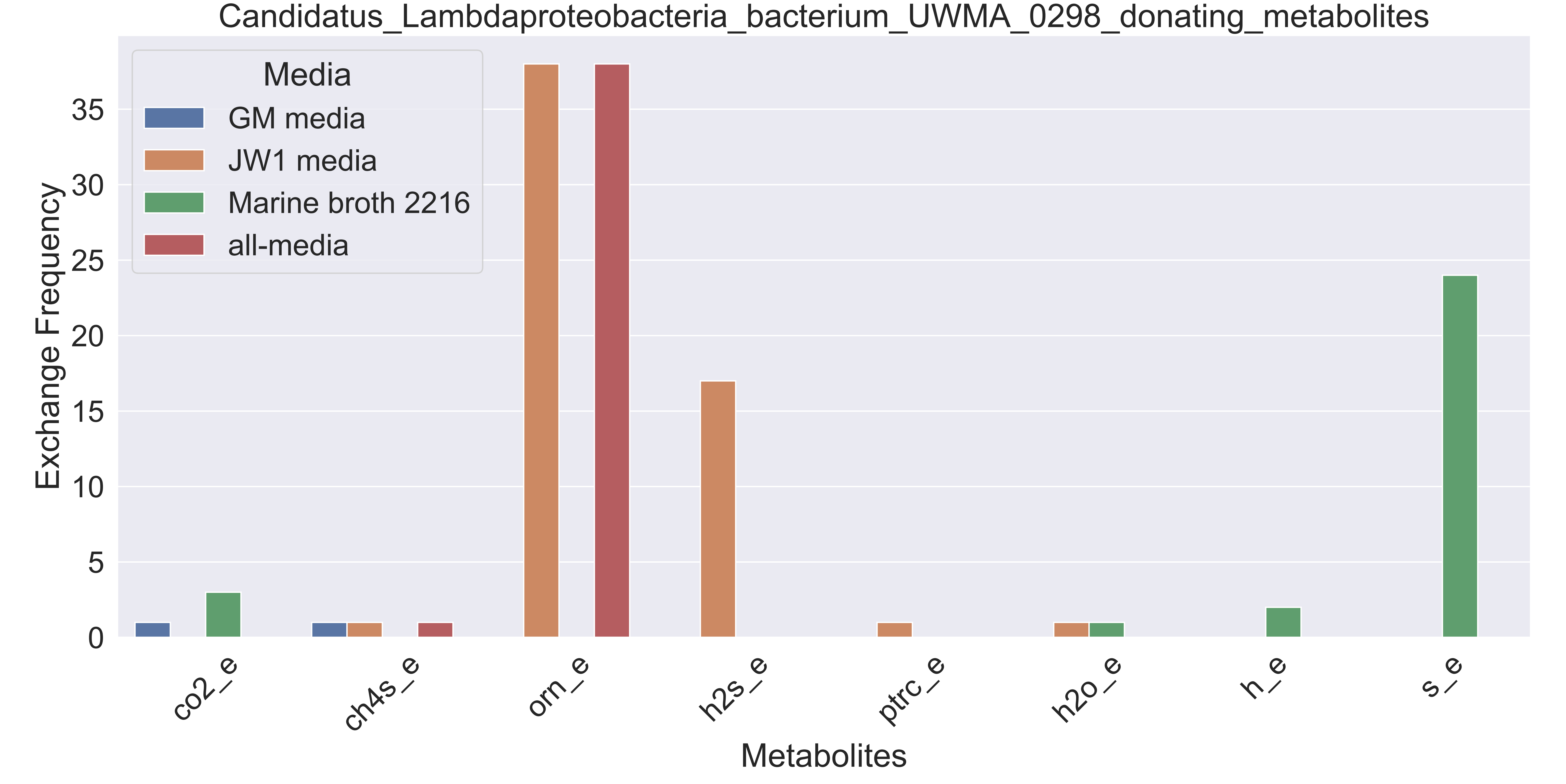

### Supplementary Figure S111 Candidatus Lambdaproteobacteria bacterium UWMA 0318 donating metabolites.pdf

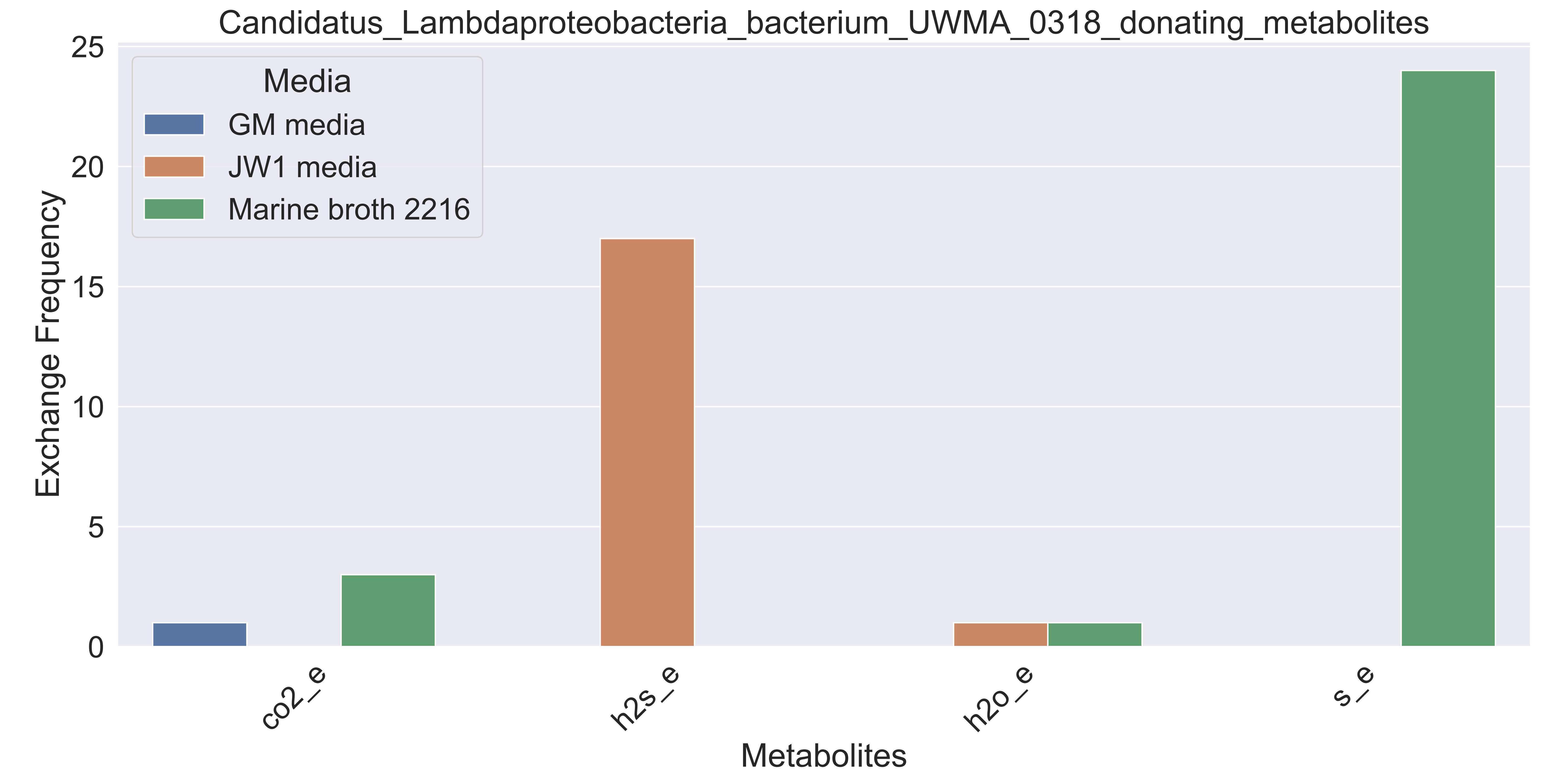

### Supplementary Figure S112 Candidatus Marinimicrobia bacterium UWMA 0285 donating metabolites.pdf

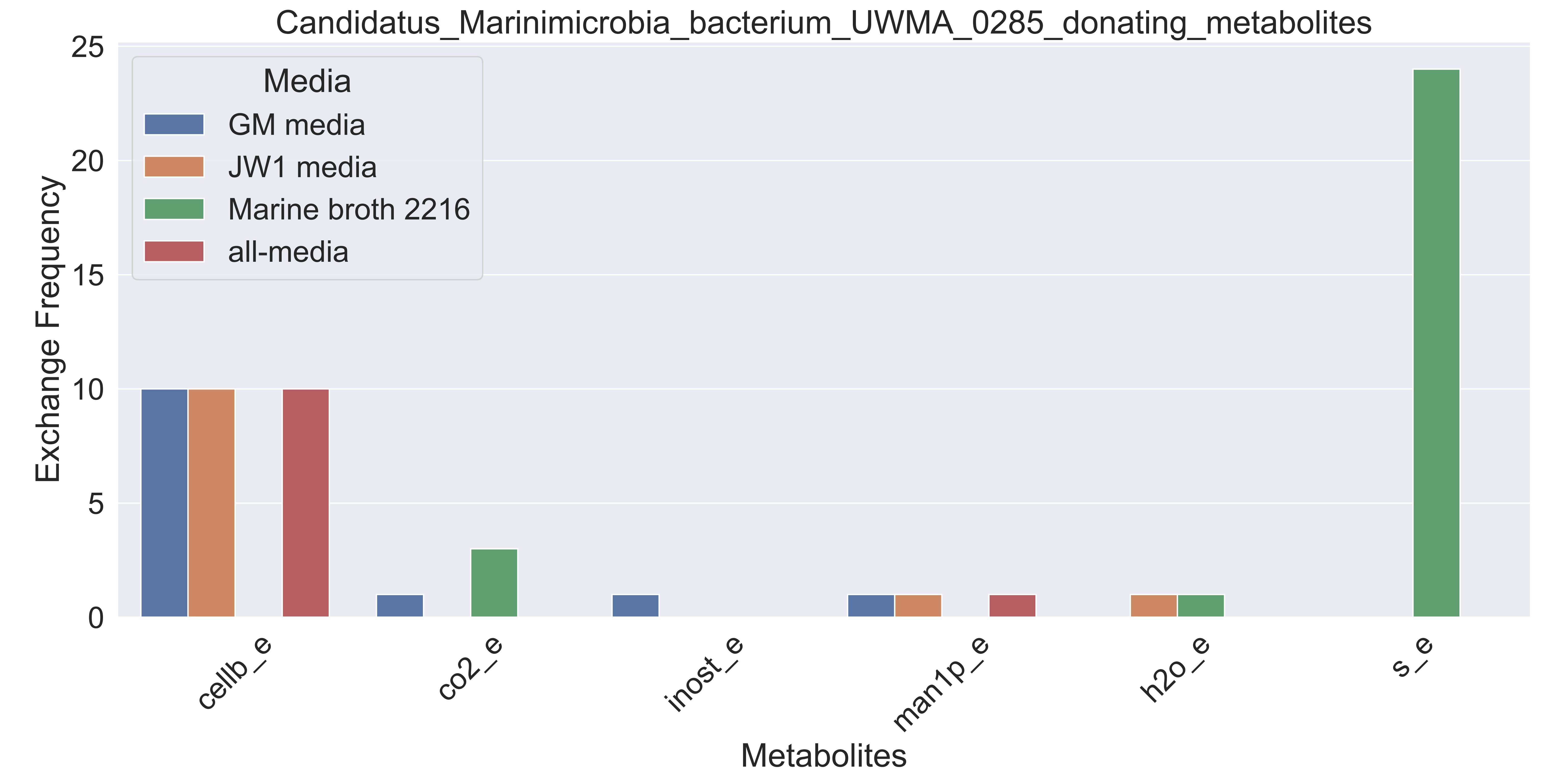

### Supplementary Figure S113 Candidatus Marinimicrobia bacterium UWMA 0306 donating metabolites.pdf

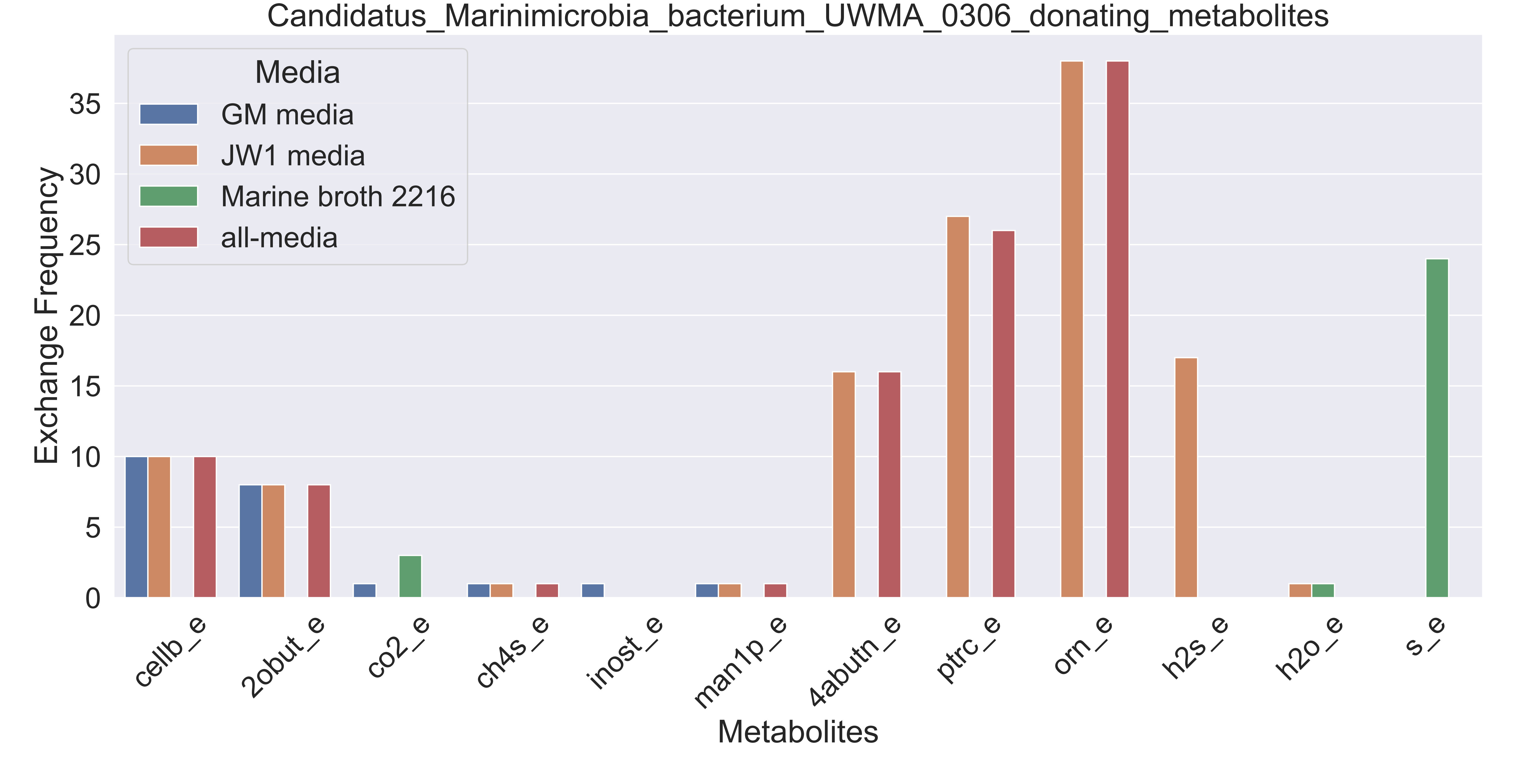

### Supplementary Figure S114 Candidatus Marinimicrobia bacterium UWMA 0309 donating metabolites.pdf

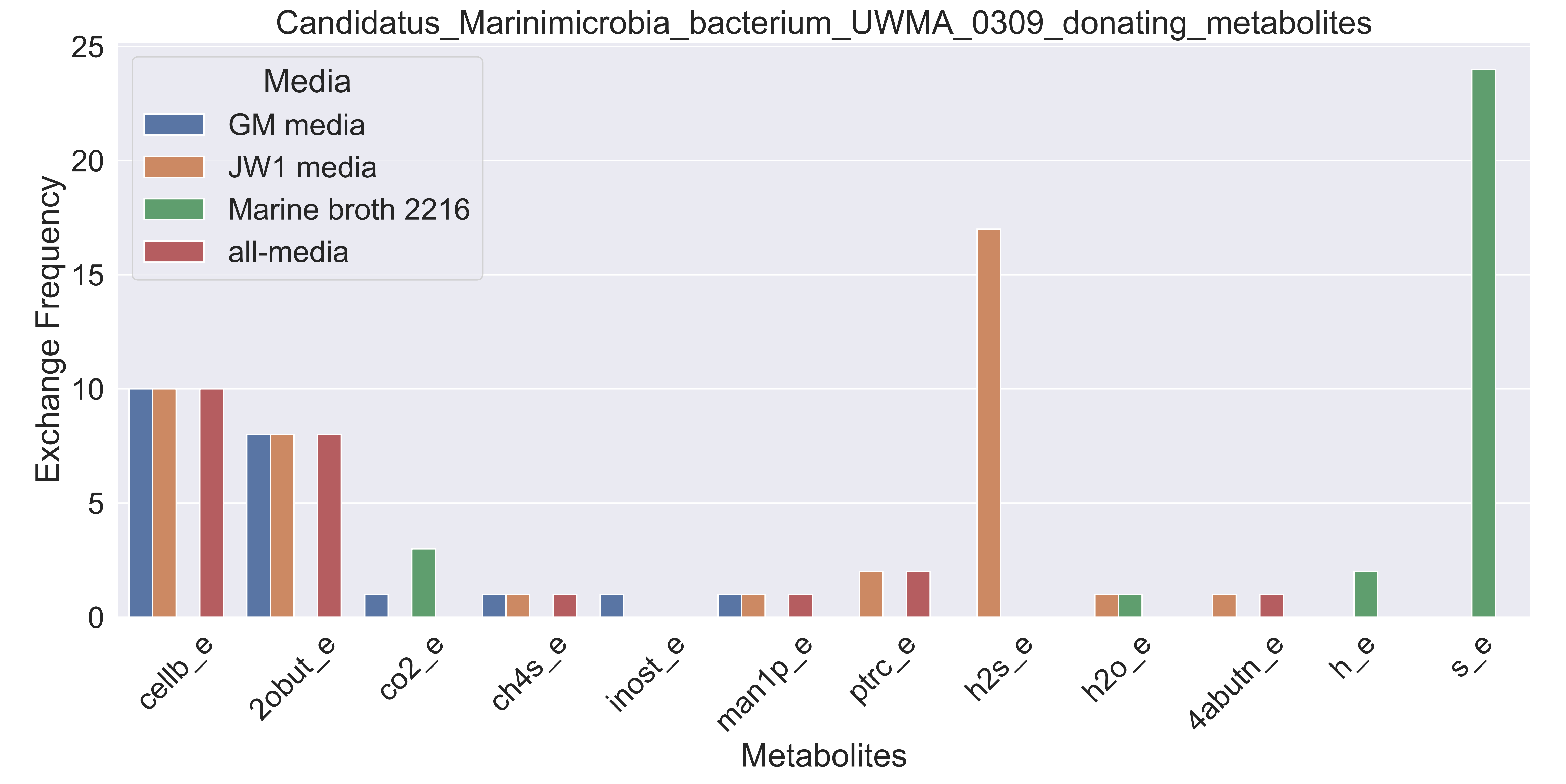

### Supplementary Figure S115 Candidatus Marinimicrobia bacterium UWMA 0347 donating metabolites.pdf

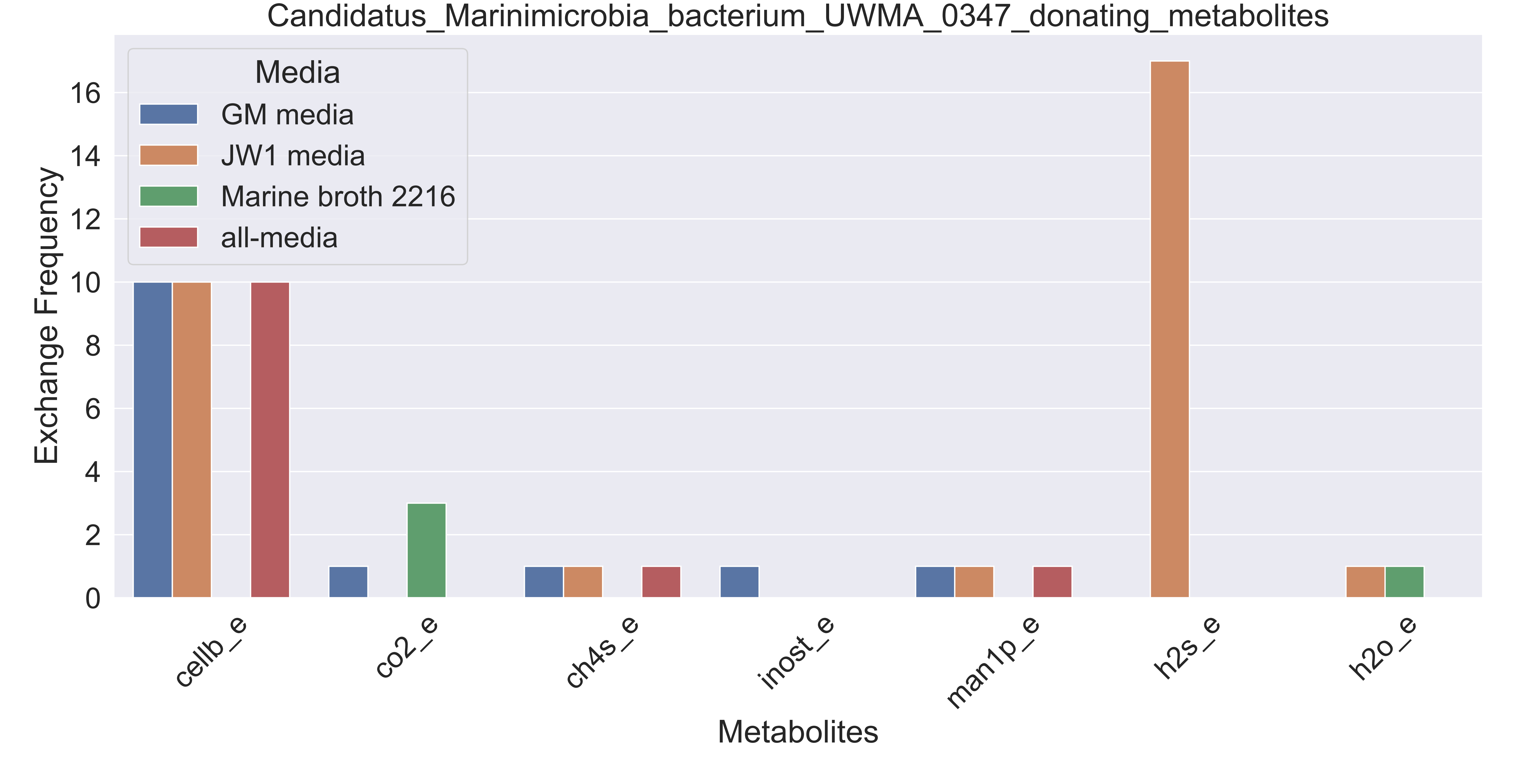

### Supplementary Figure S116 Candidatus Nitrosopelagicus sp UWMA 0359 donating metabolites.pdf

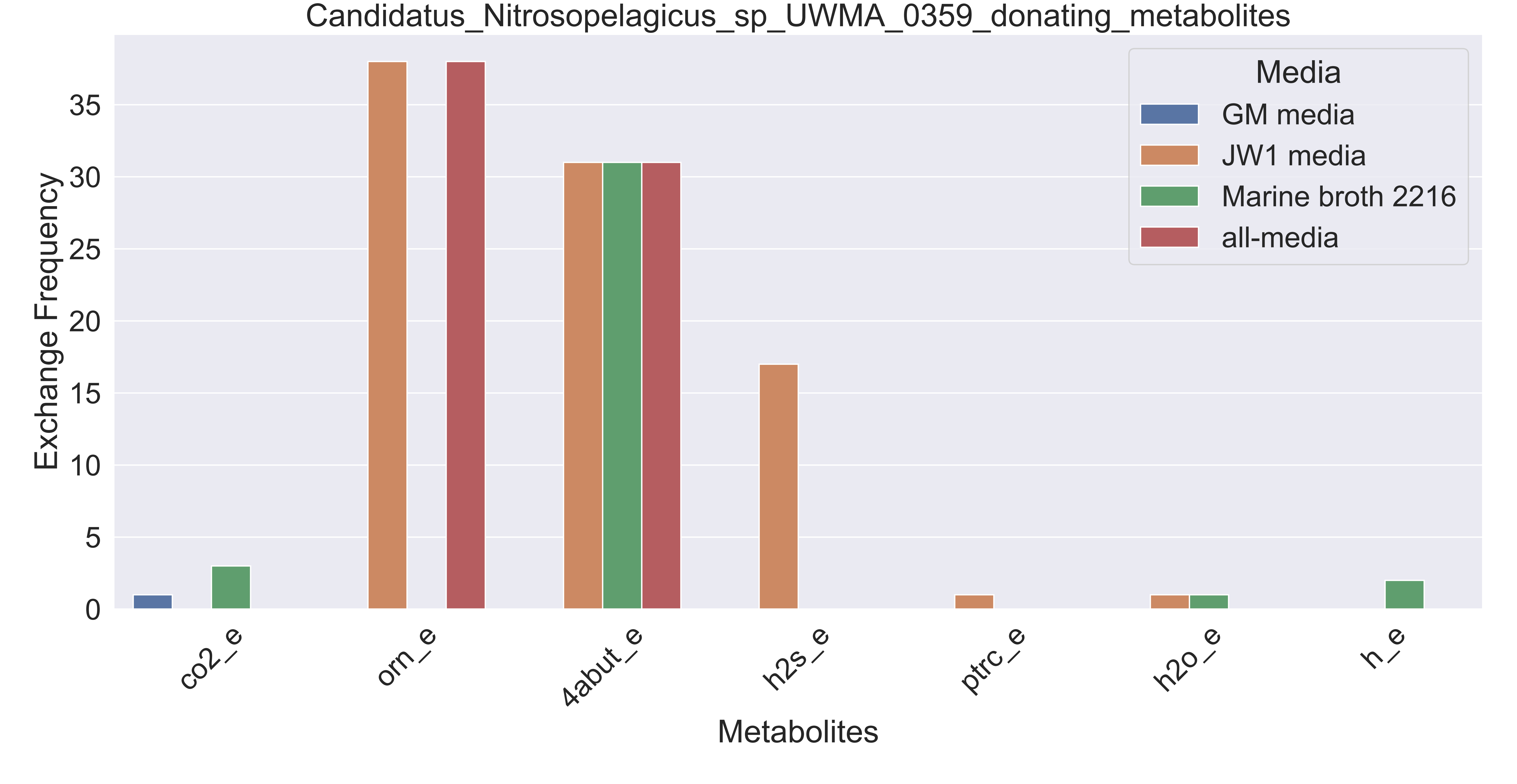

### Supplementary Figure S117 Candidatus Pacearchaeota archaeon UWMA 0287 donating metabolites.pdf

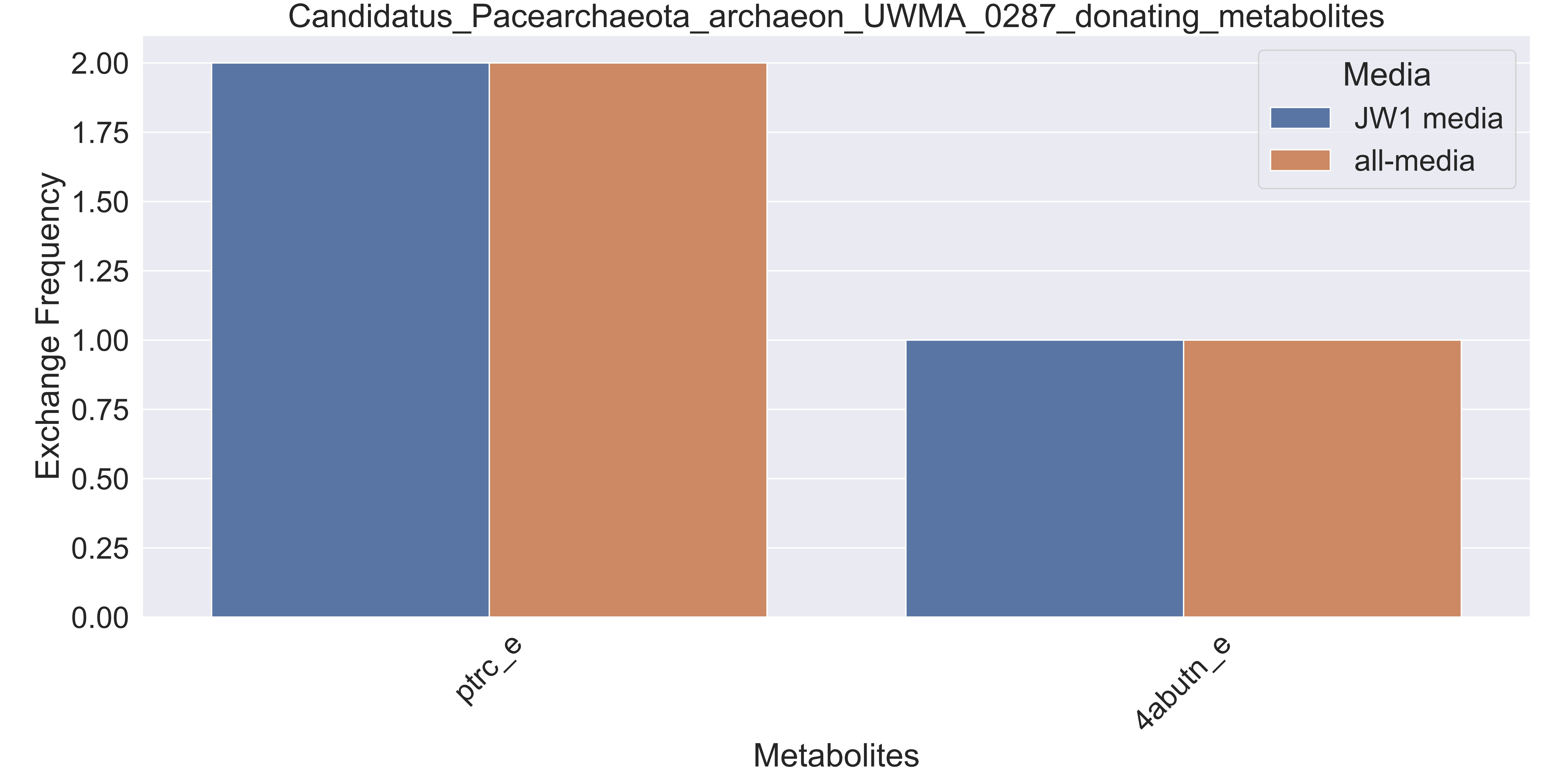

### Supplementary Figure S118 Candidatus Pelagibacter sp UWMA 0307 donating metabolites.pdf

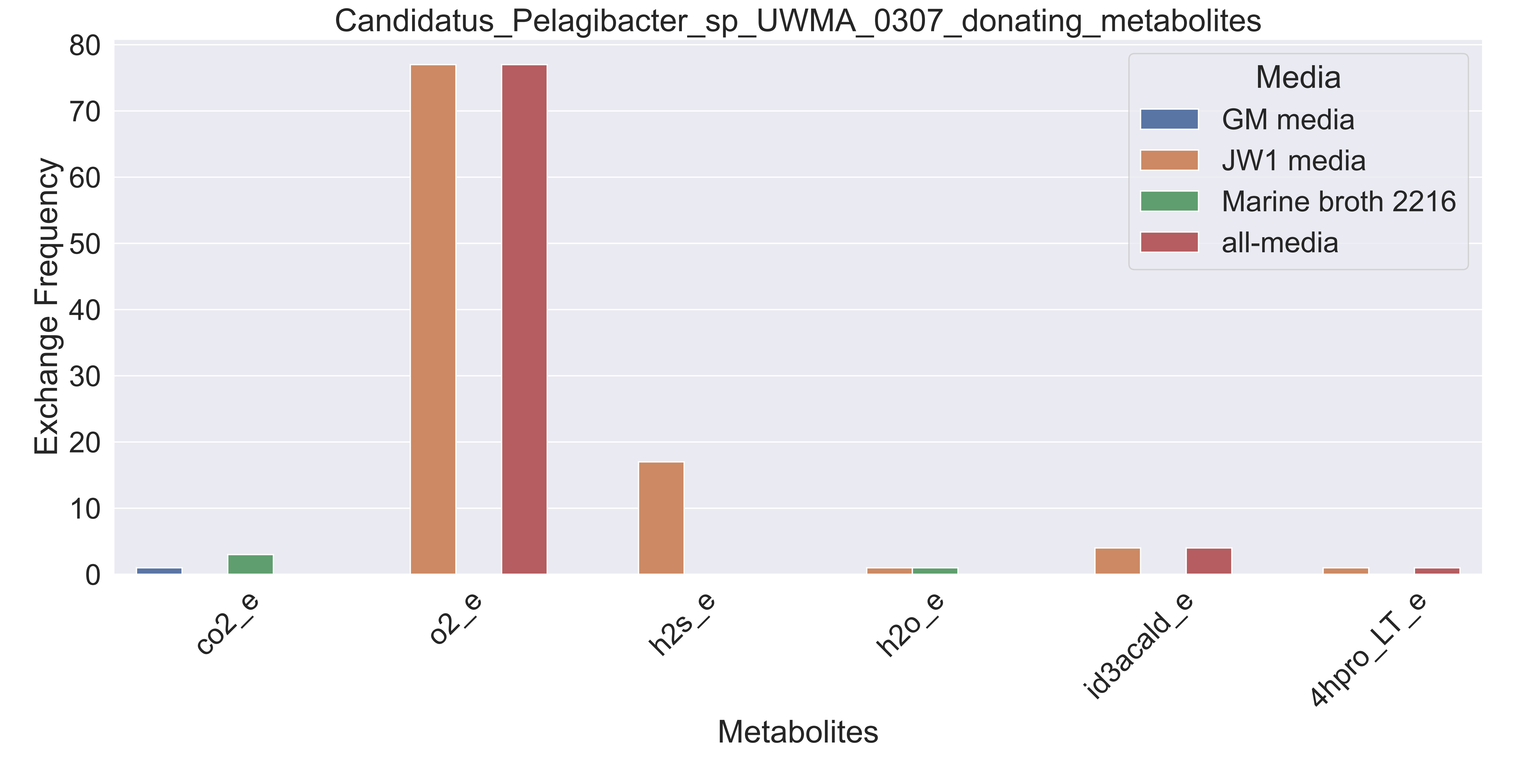

### Supplementary Figure S119 Candidatus Thioglobus sp UWMA 0259 donating metabolites.pdf

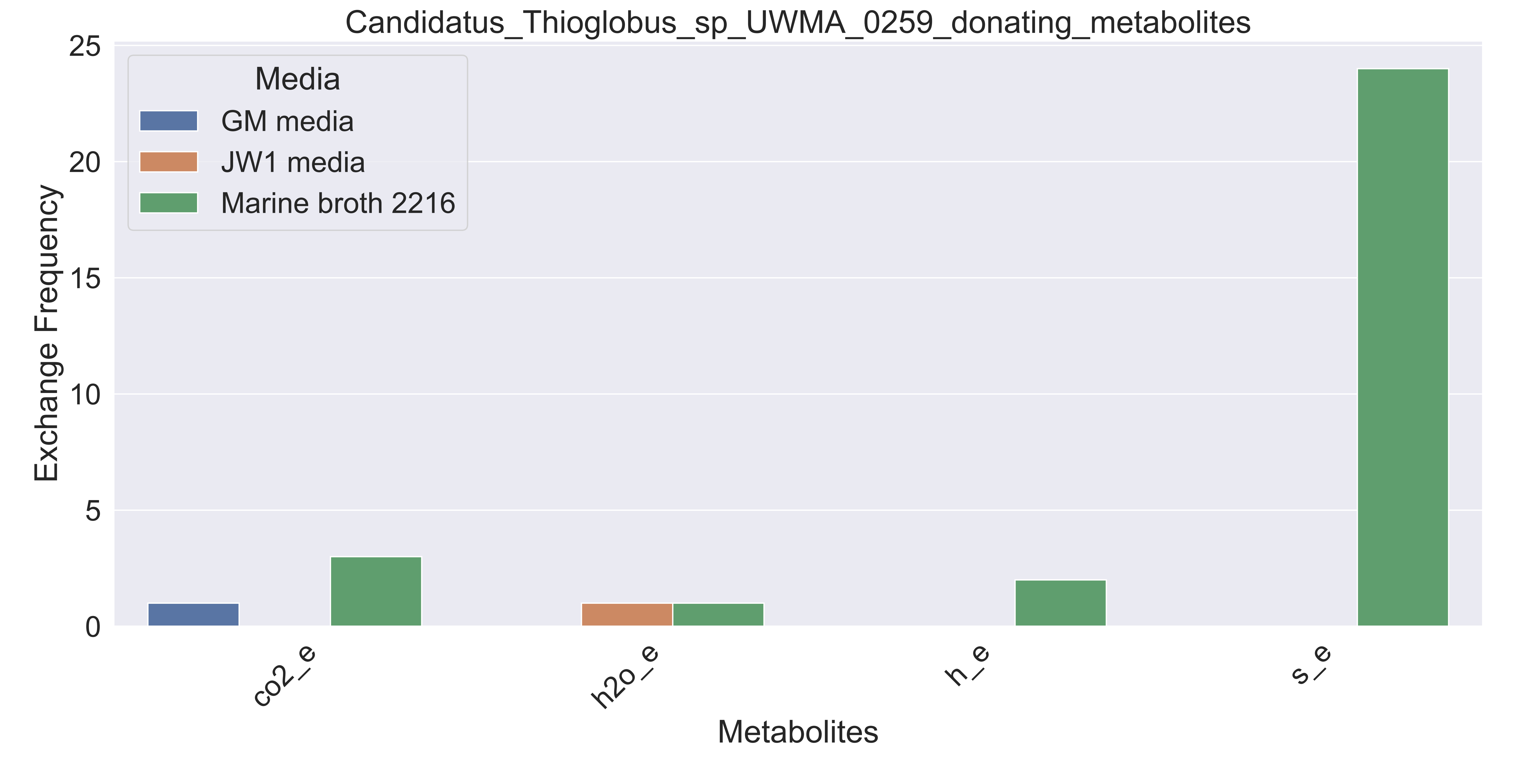

### Supplementary Figure S120 Candidatus Thioglobus sp UWMA 0272 donating metabolites.pdf

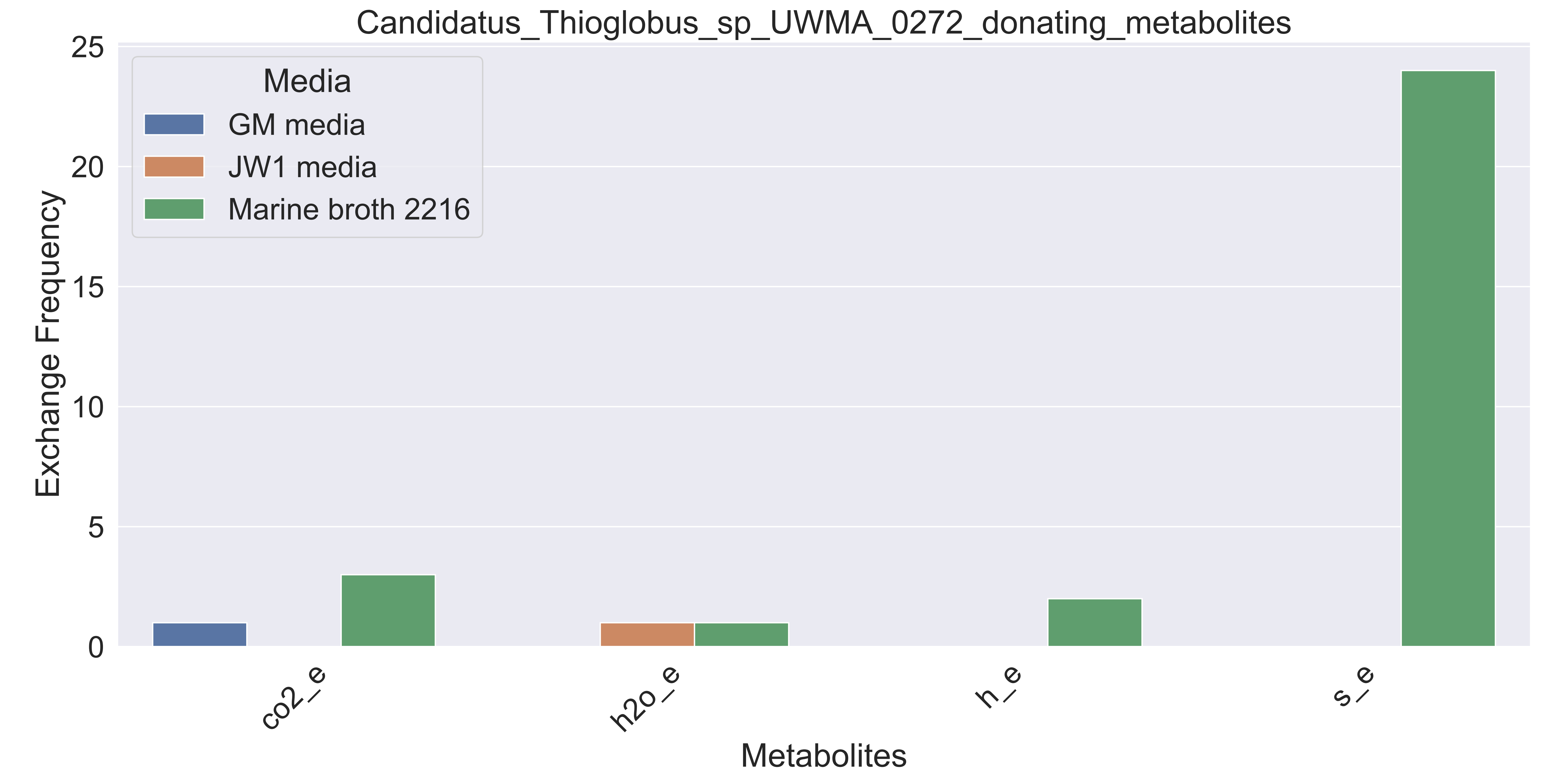

### Supplementary Figure S121 Candidatus Thioglobus sp UWMA 0322 donating metabolites.pdf

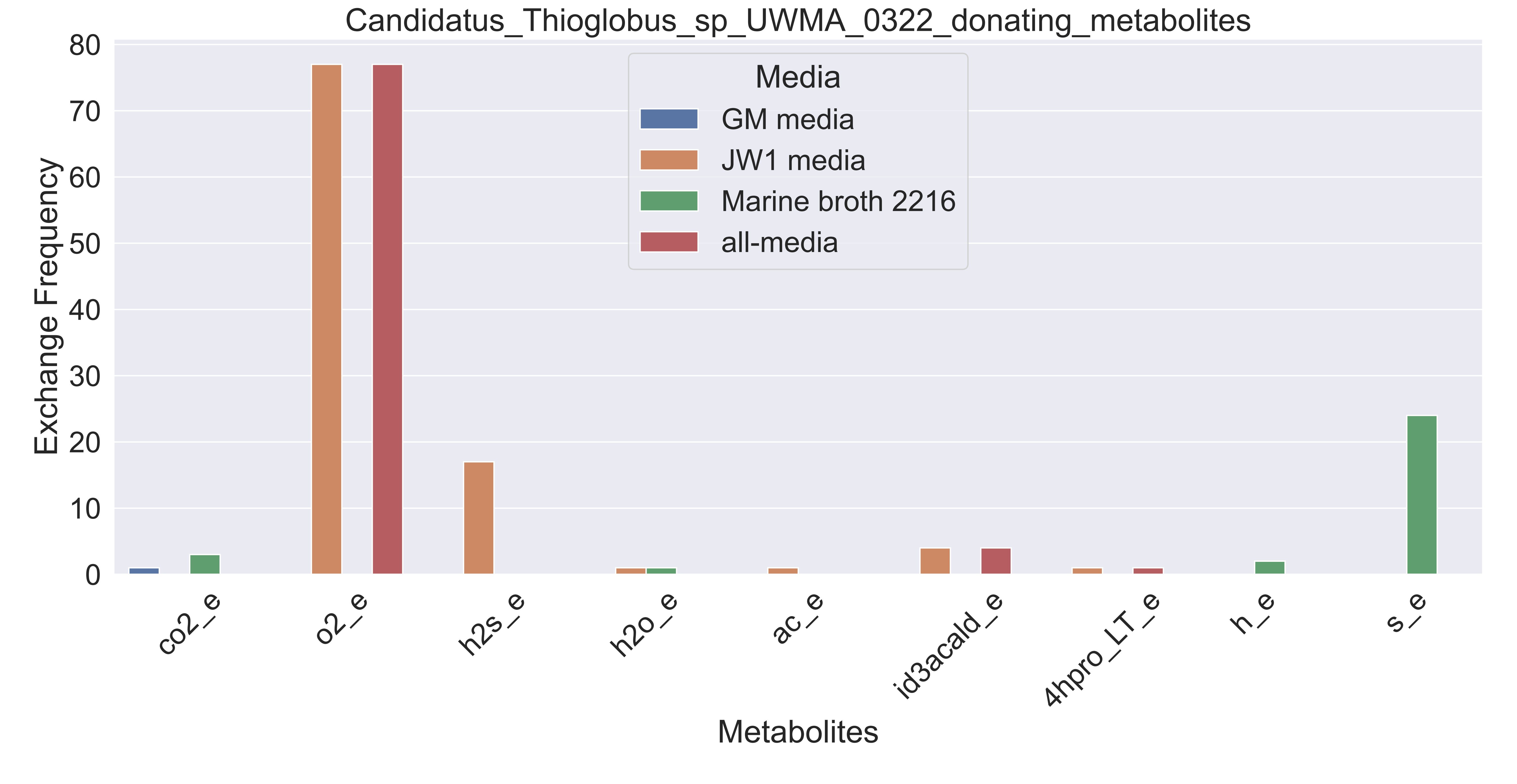

### Supplementary Figure S122 Candidatus Thioglobus sp UWMA 0342 donating metabolites.pdf

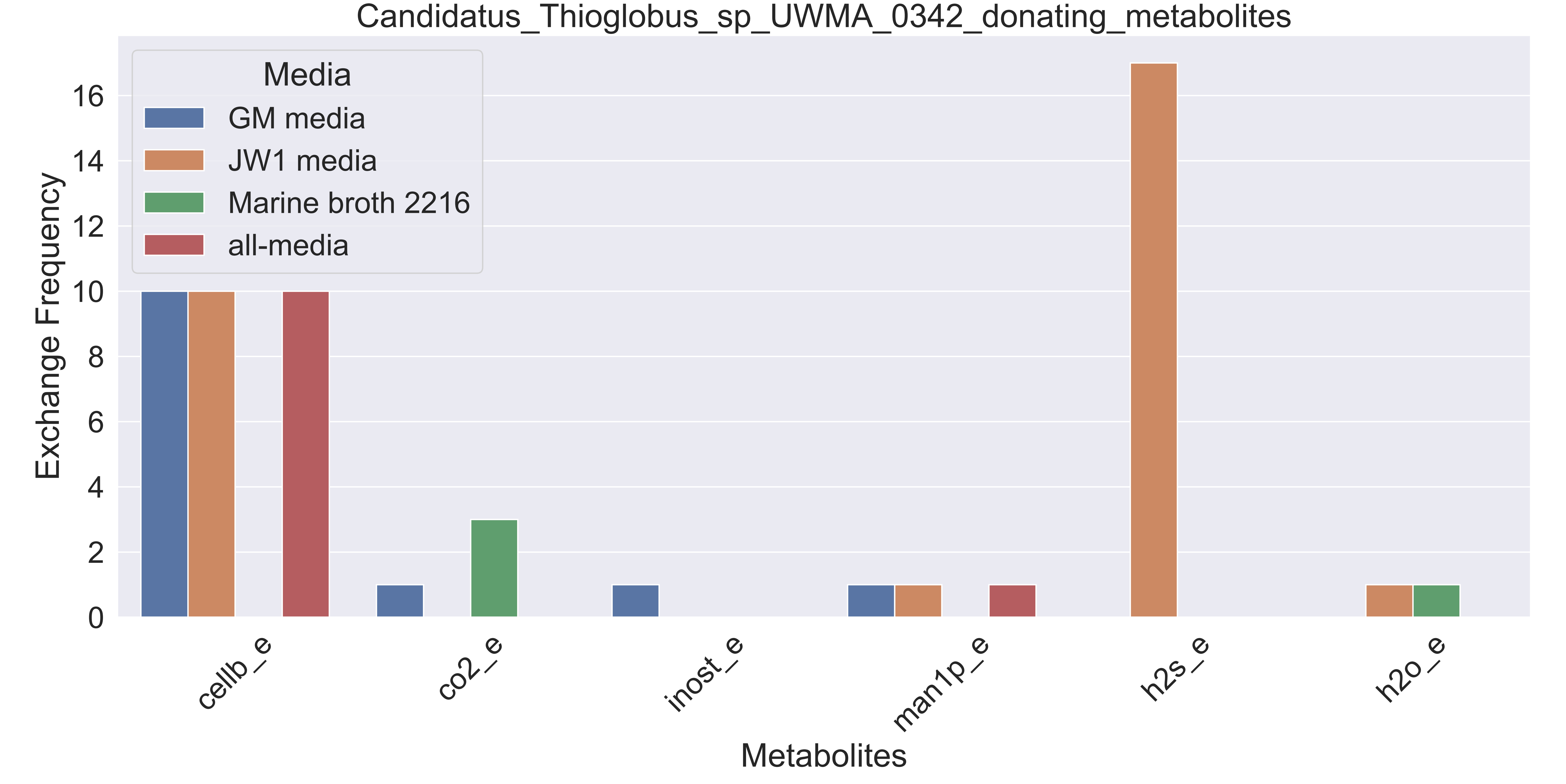

### Supplementary Figure S123 Candidatus Thioglobus sp UWMA 0360 donating metabolites.pdf

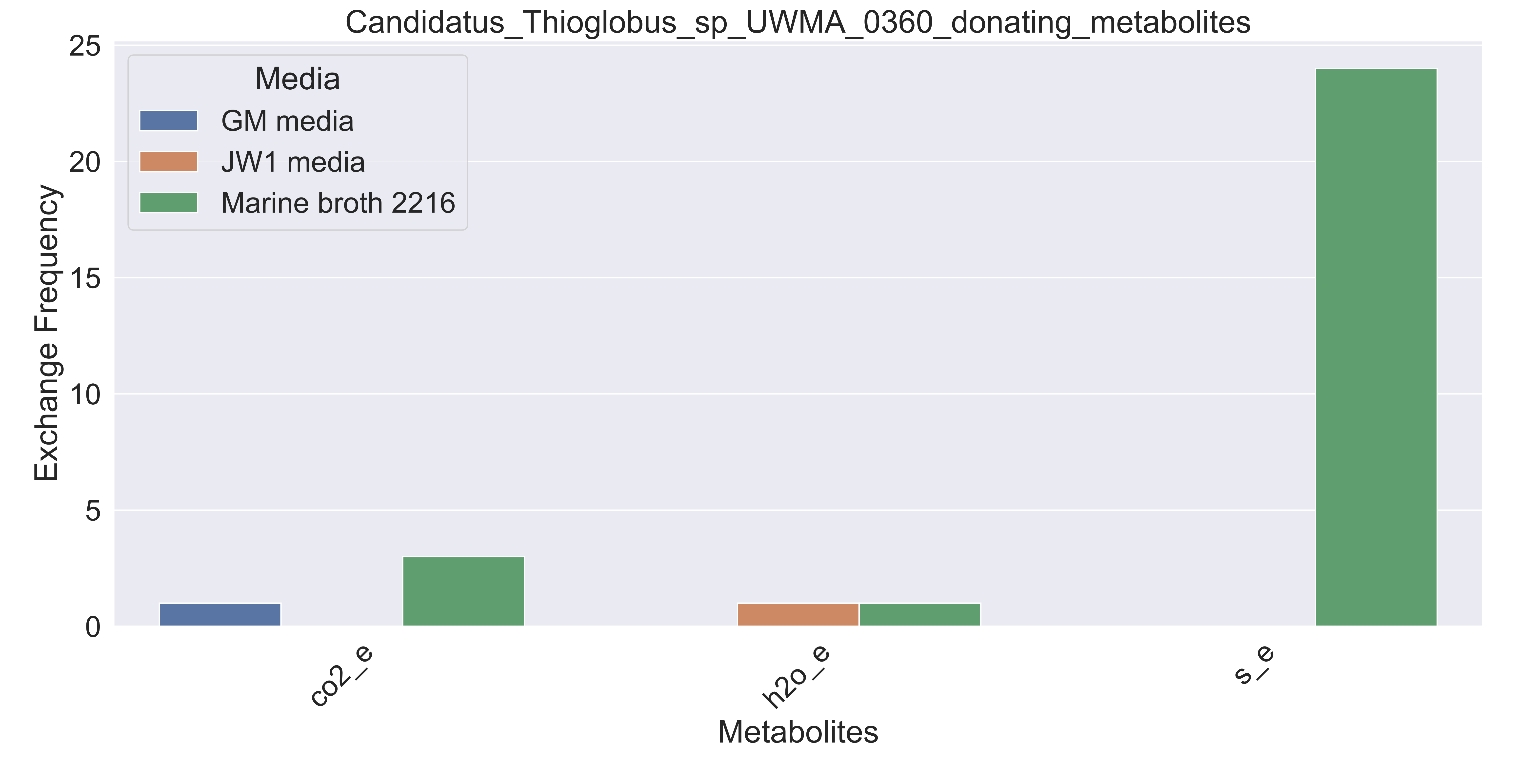

### Supplementary Figure S124 Cycloclasticus sp UWMA 0316 donating metabolites.pdf

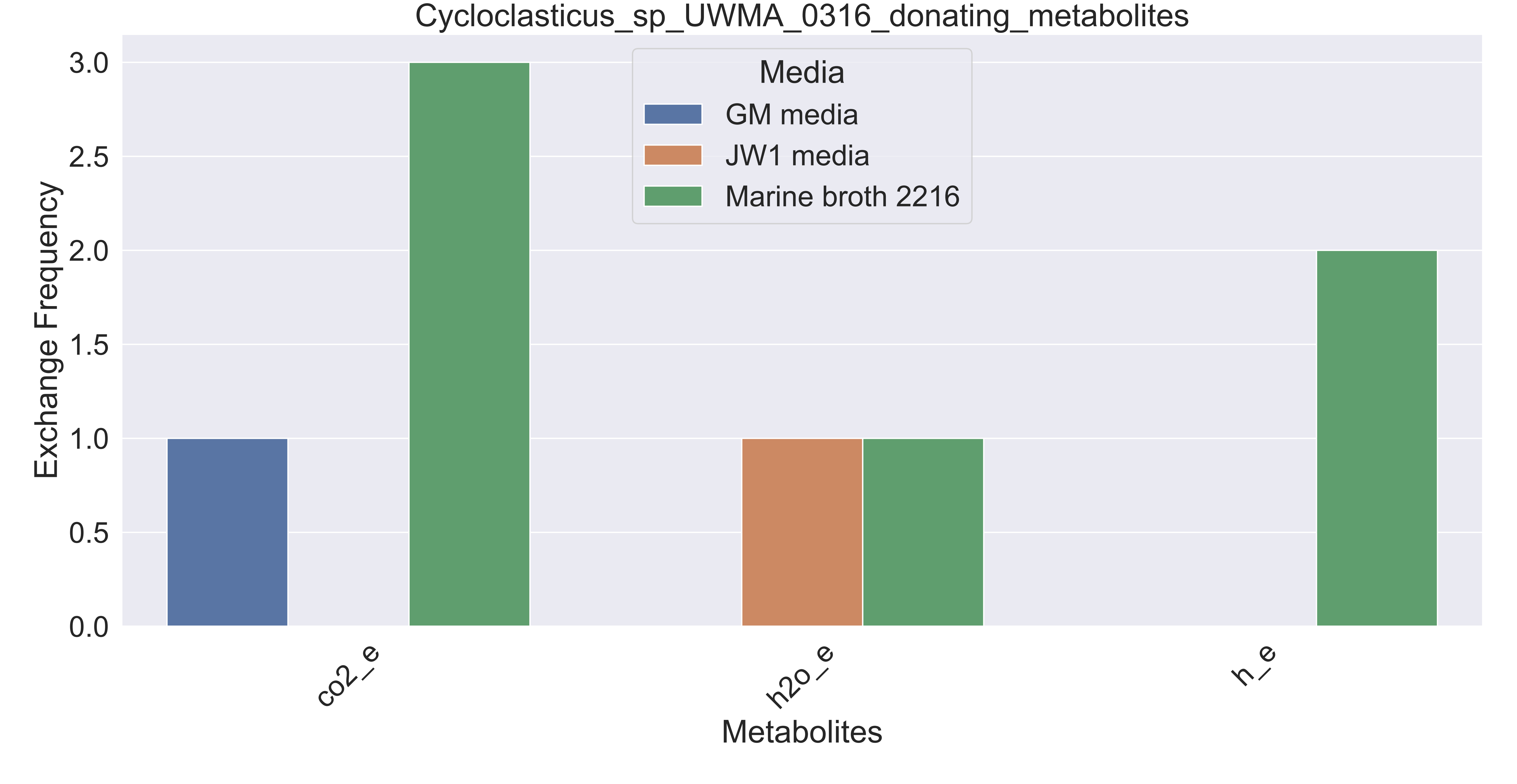

### Supplementary Figure S125 Cycloclasticus sp UWMA 0335 donating metabolites.pdf

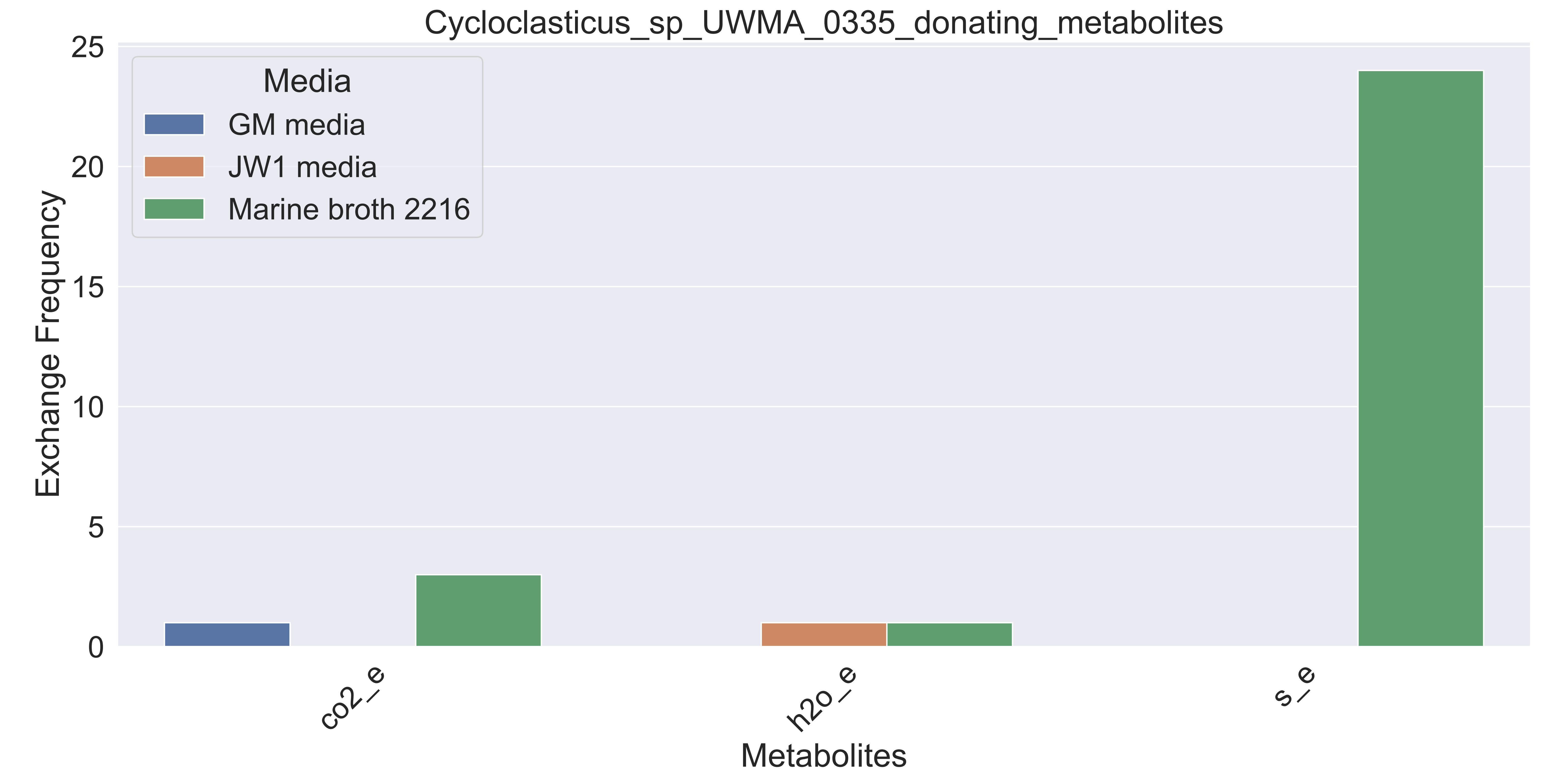

### Supplementary Figure S126 Cytophagales bacterium UWMA 0338 donating metabolites.pdf

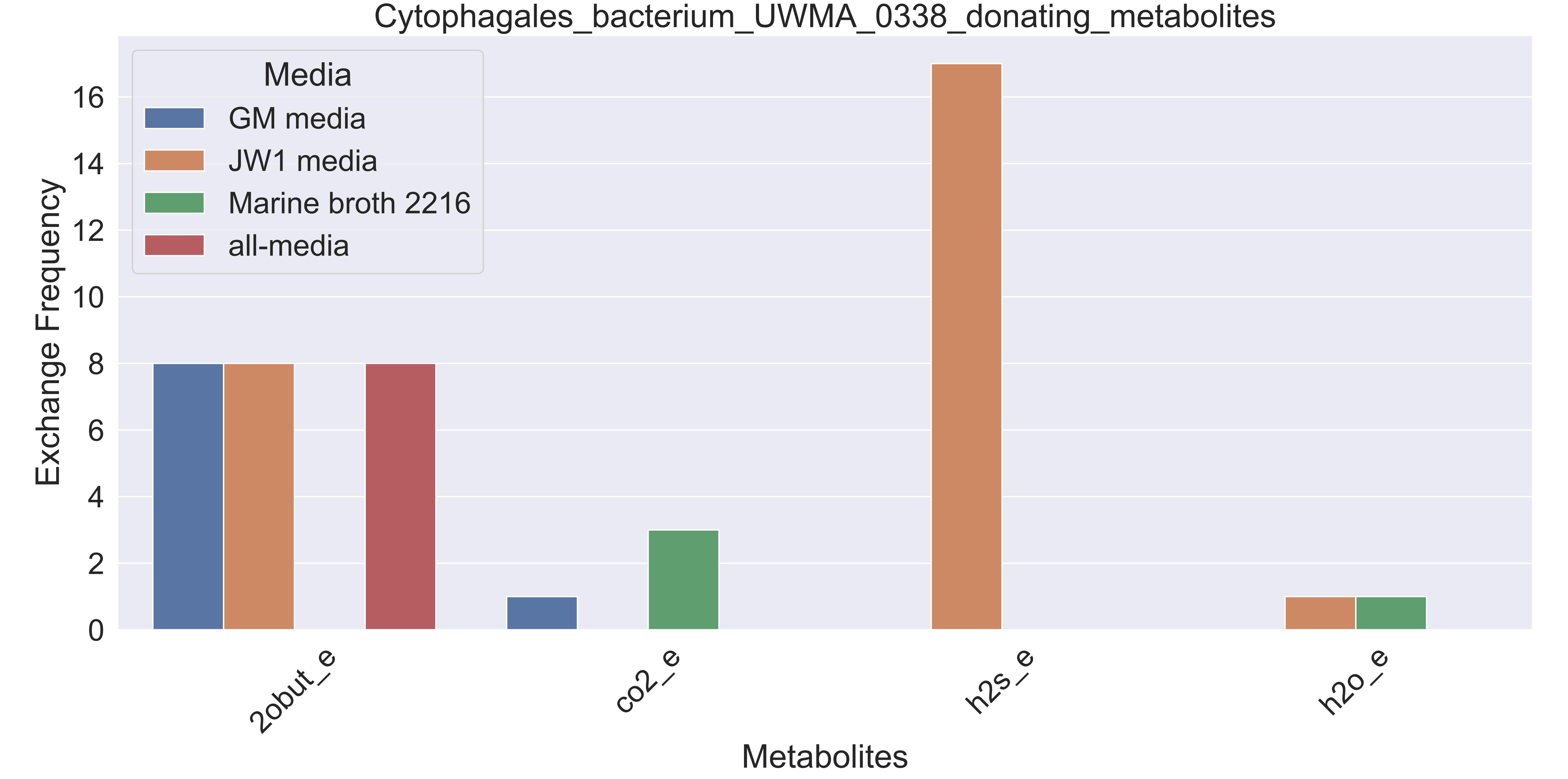

### Supplementary Figure S127 Dehalococcoidia bacterium UWMA 0267 donating metabolites.pdf

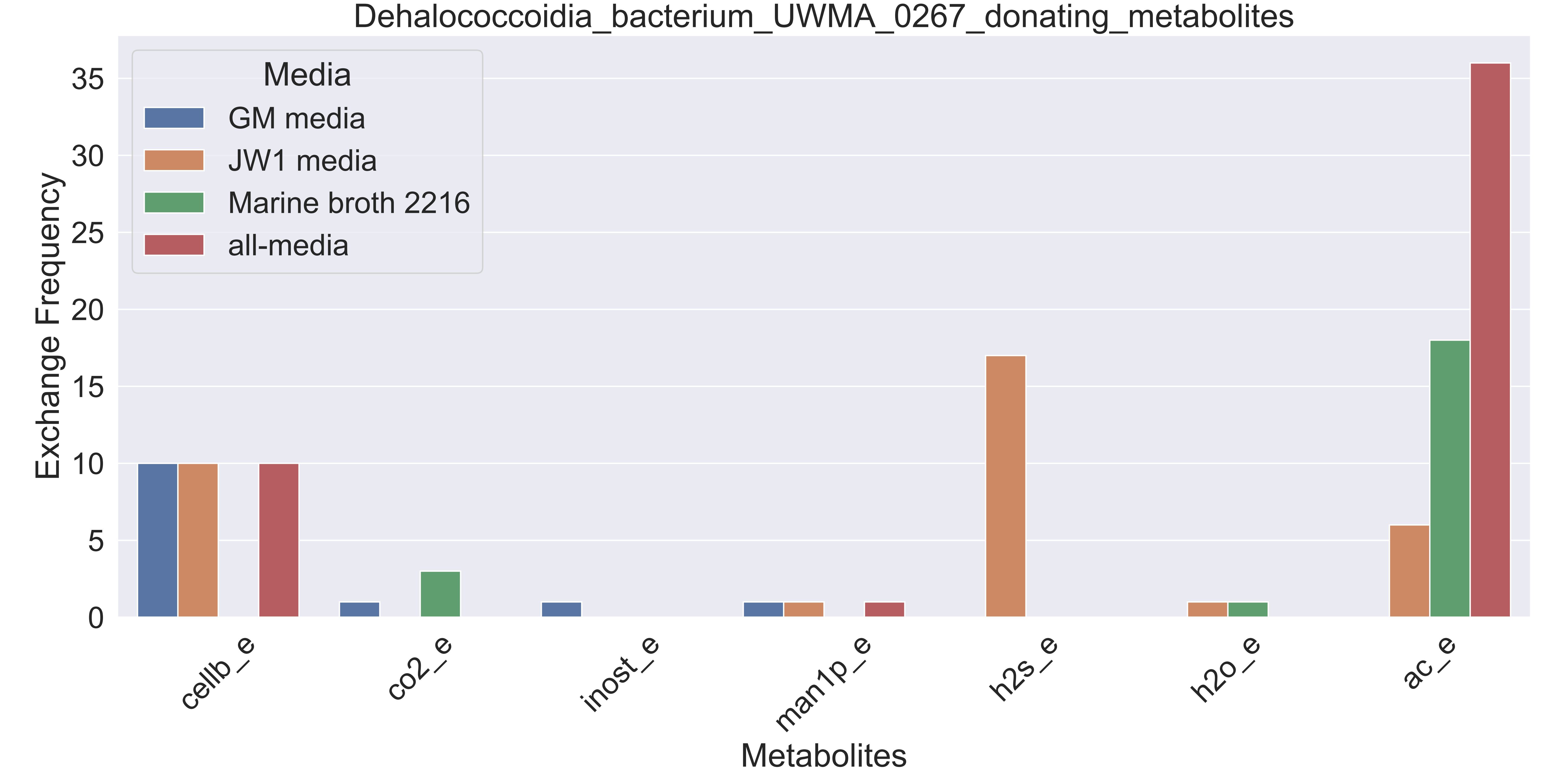

### Supplementary Figure S128 Dehalococcoidia bacterium UWMA 0331 donating metabolites.pdf

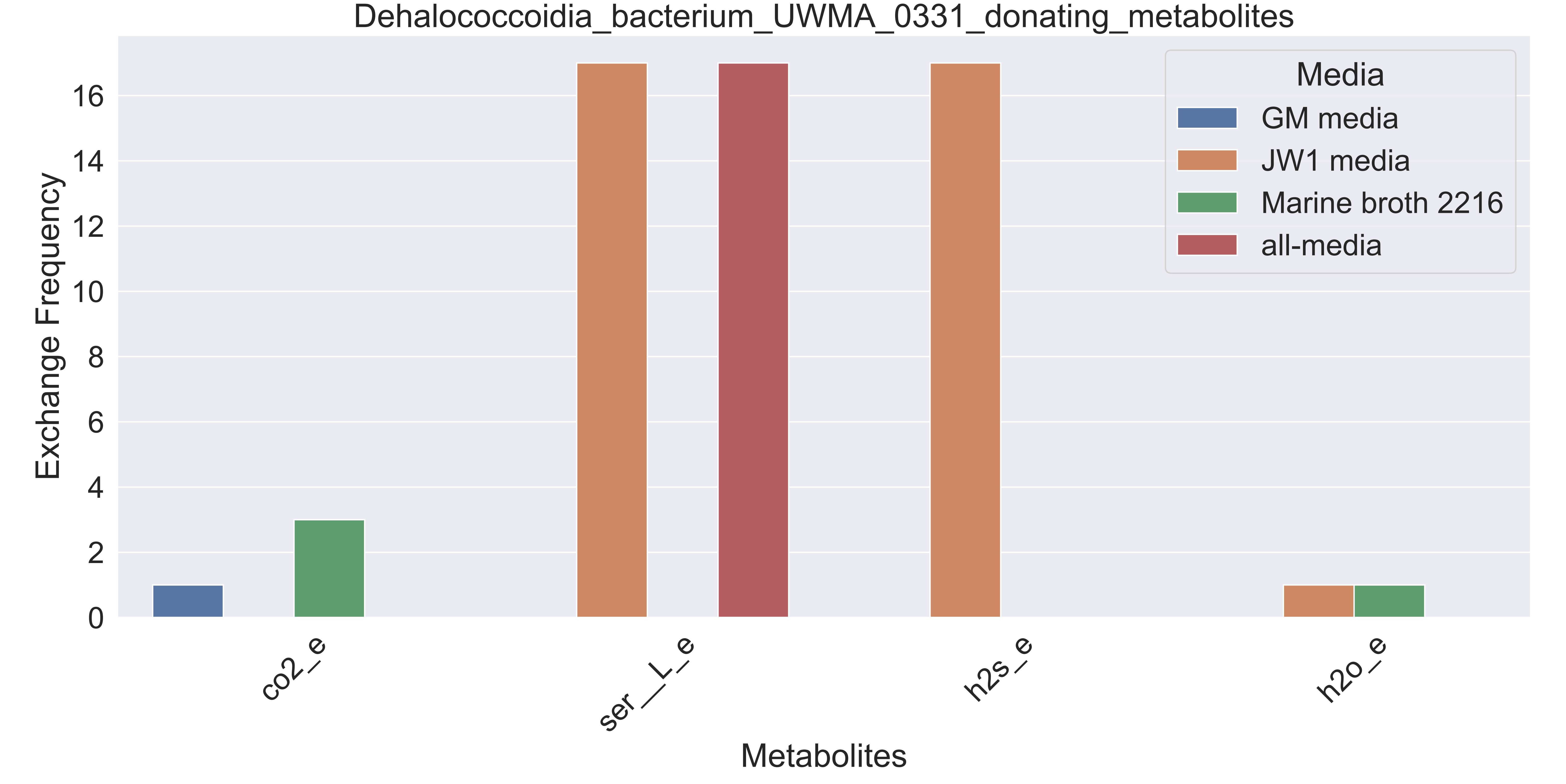

### Supplementary Figure S129 Dehalococcoidia bacterium UWMA 0337 donating metabolites.pdf

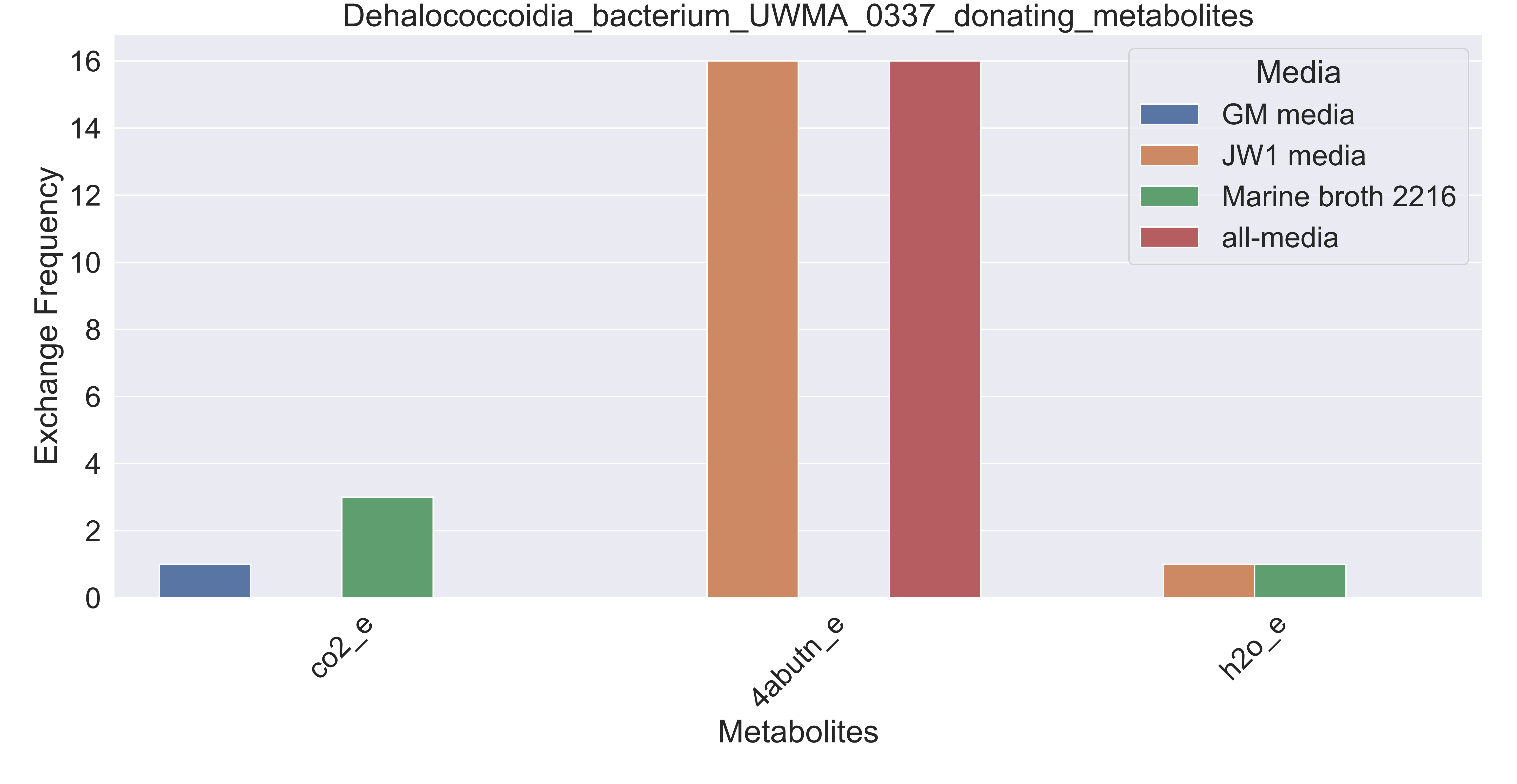

### Supplementary Figure S130 Dehalococcoidia bacterium UWMA 0354 donating metabolites.pdf

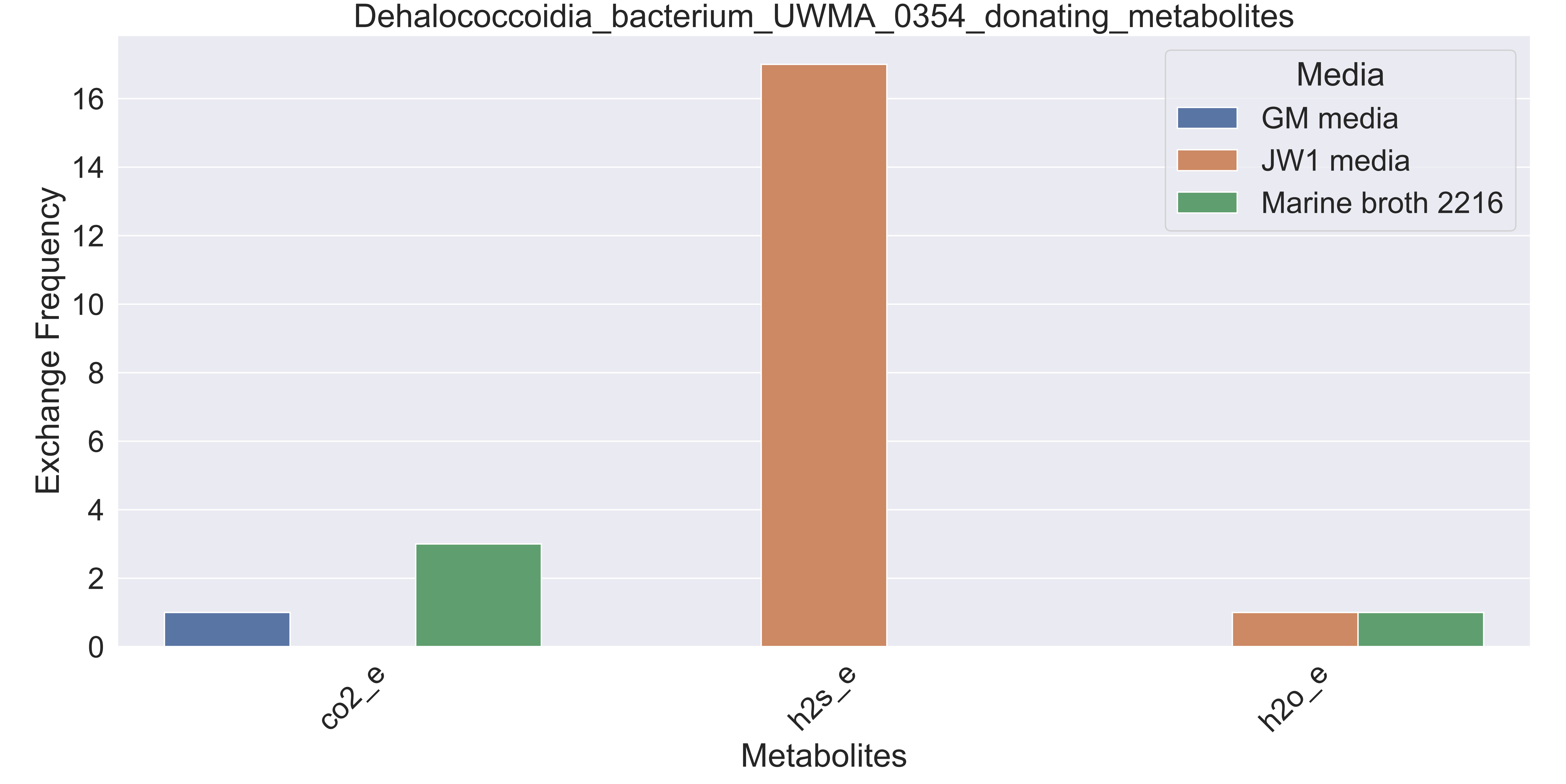

### Supplementary Figure S131 Flavobacteriaceae bacterium UWMA 0314 donating metabolites.pdf

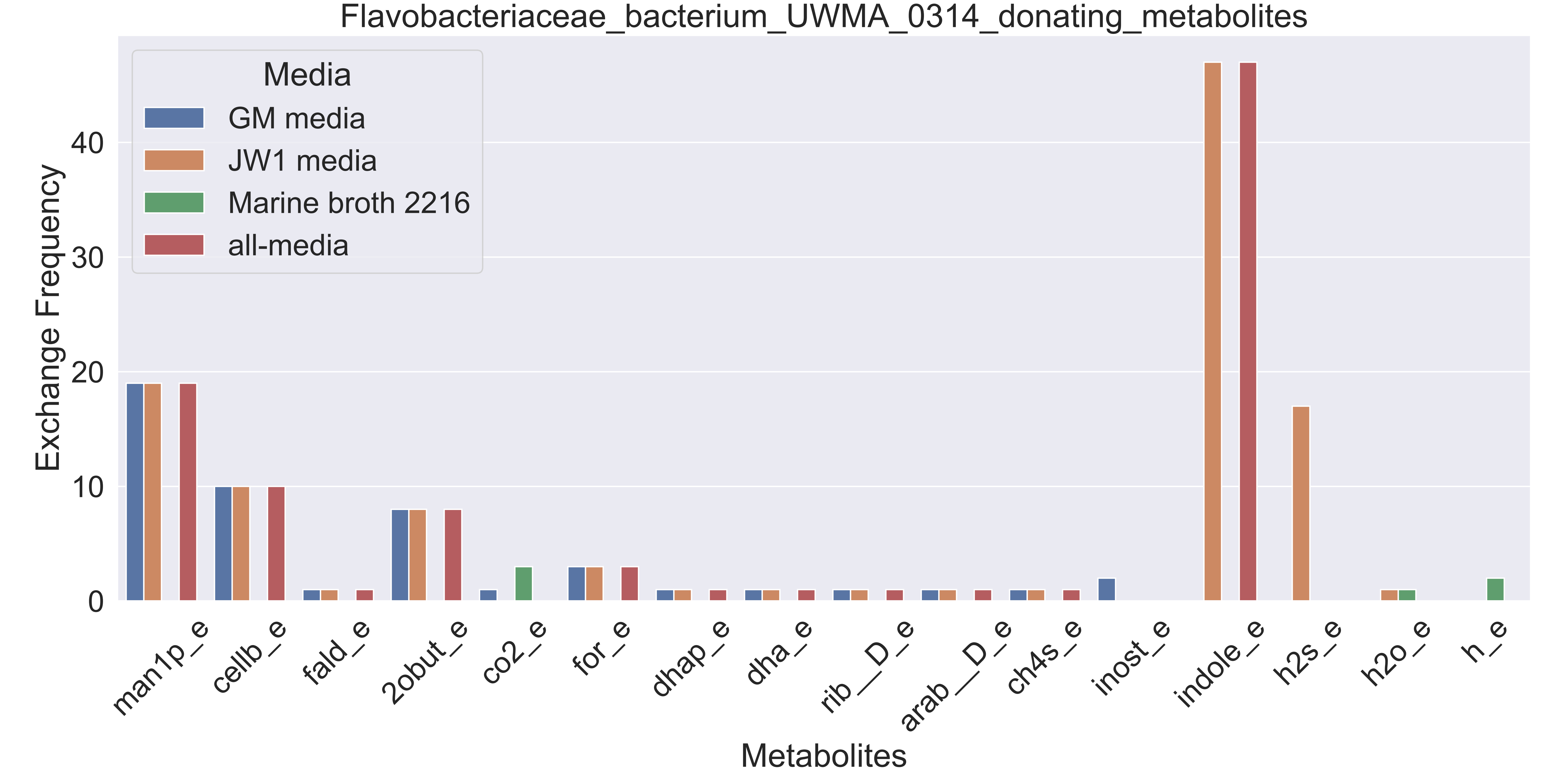

### Supplementary Figure S132 Flavobacteriaceae bacterium UWMA 0351 donating metabolites.pdf

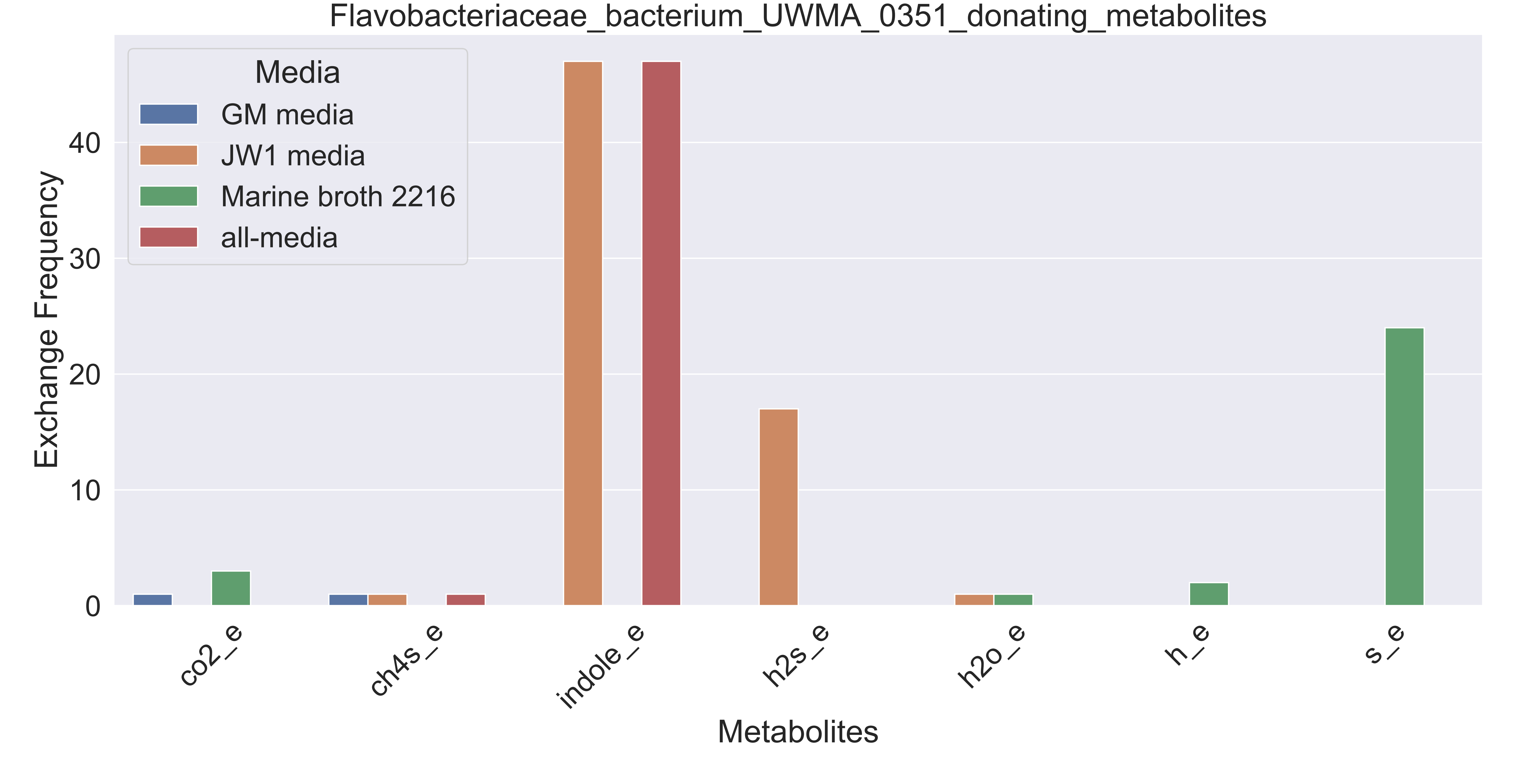

### Supplementary Figure S133 Flavobacteriales bacterium UWMA 0274 donating metabolites.pdf

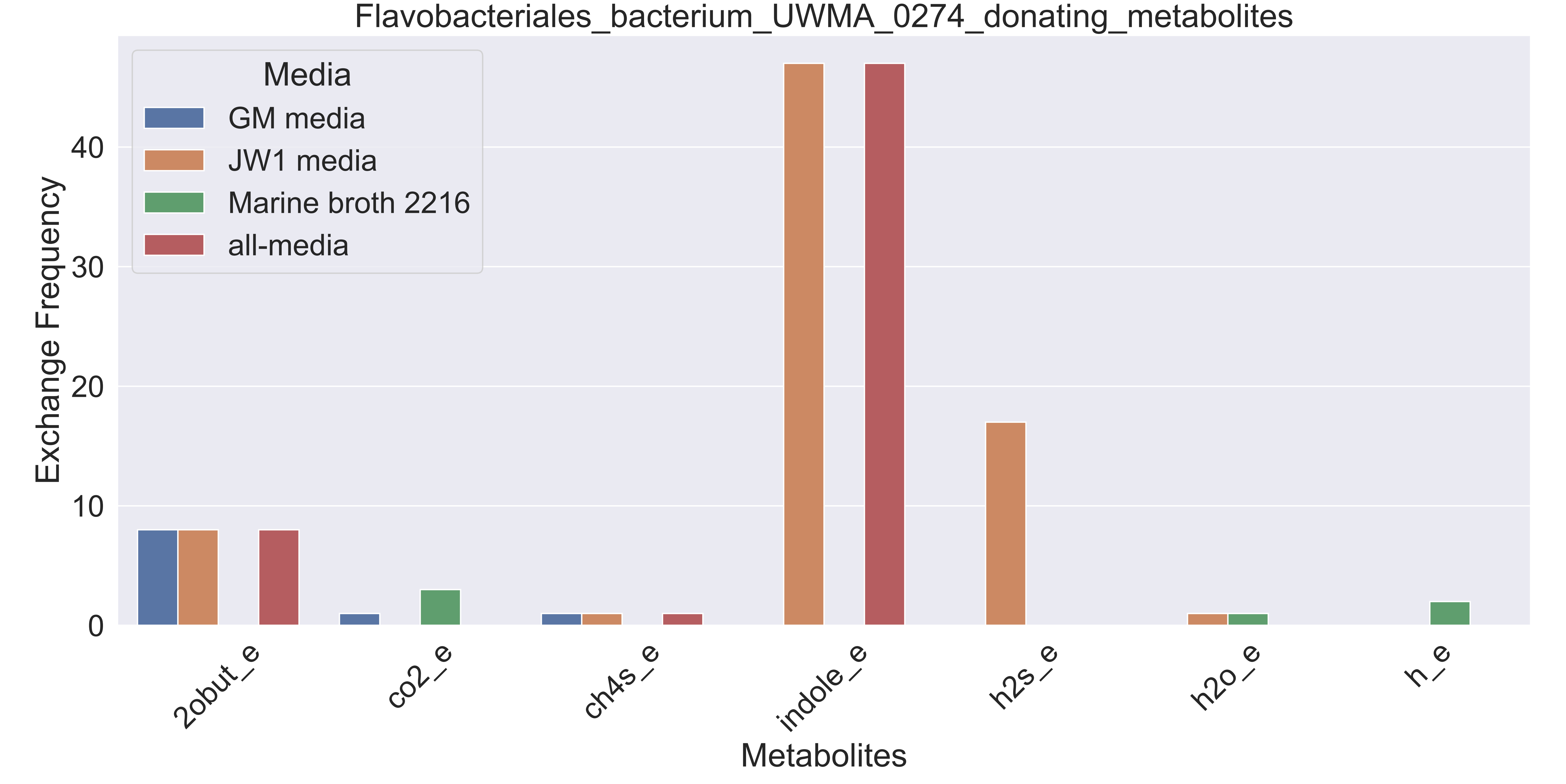

### Supplementary Figure S134 Flavobacteriales bacterium UWMA 0277 donating metabolites.pdf

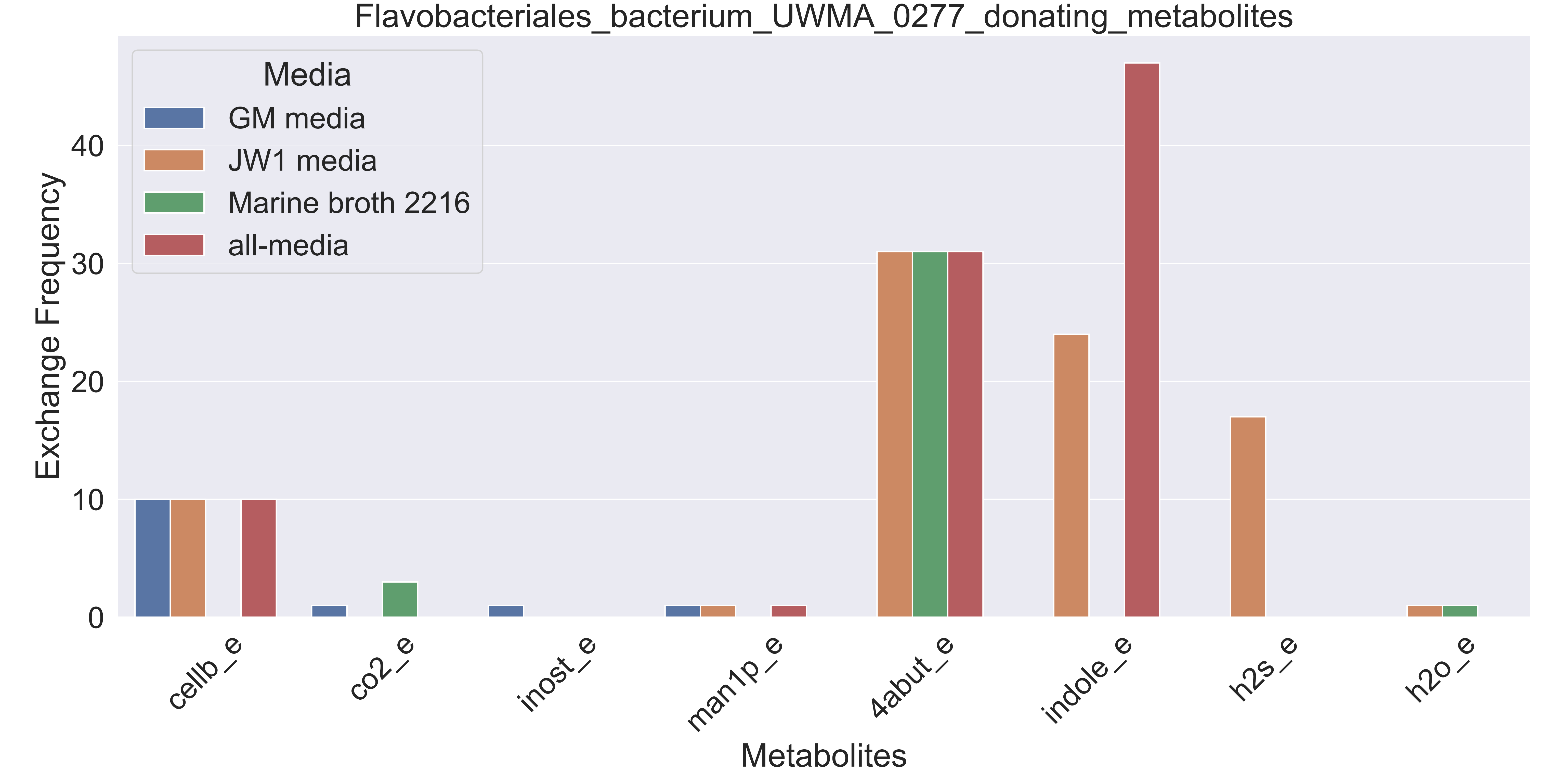

### Supplementary Figure S135 Flavobacteriales bacterium UWMA 0289 donating metabolites.pdf

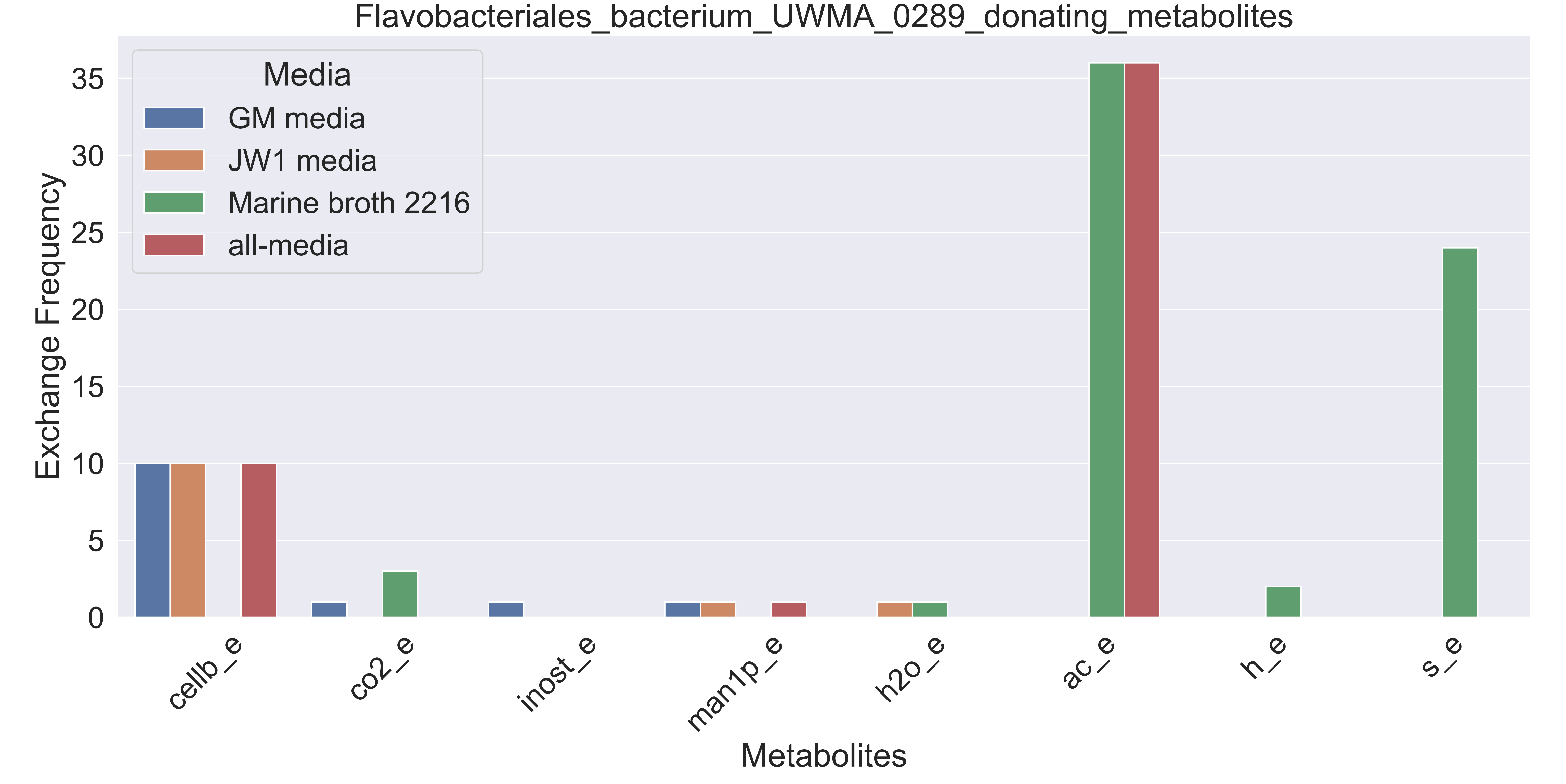

### Supplementary Figure S136 Fuerstia sp UWMA 0333 donating metabolites.pdf

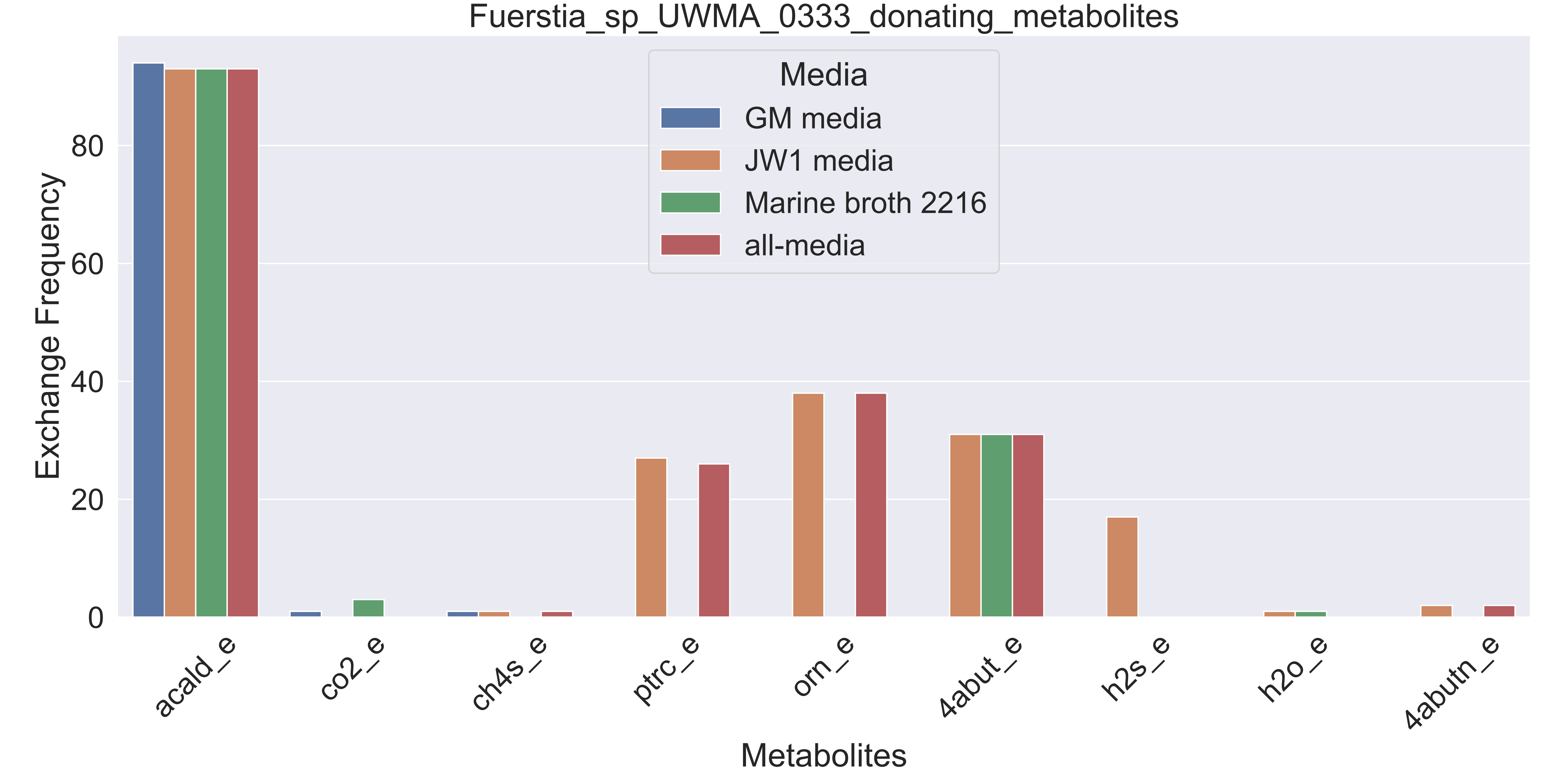

### Supplementary Figure S137 Gammaproteobacteria bacterium UWMA 0258 donating metabolites.pdf

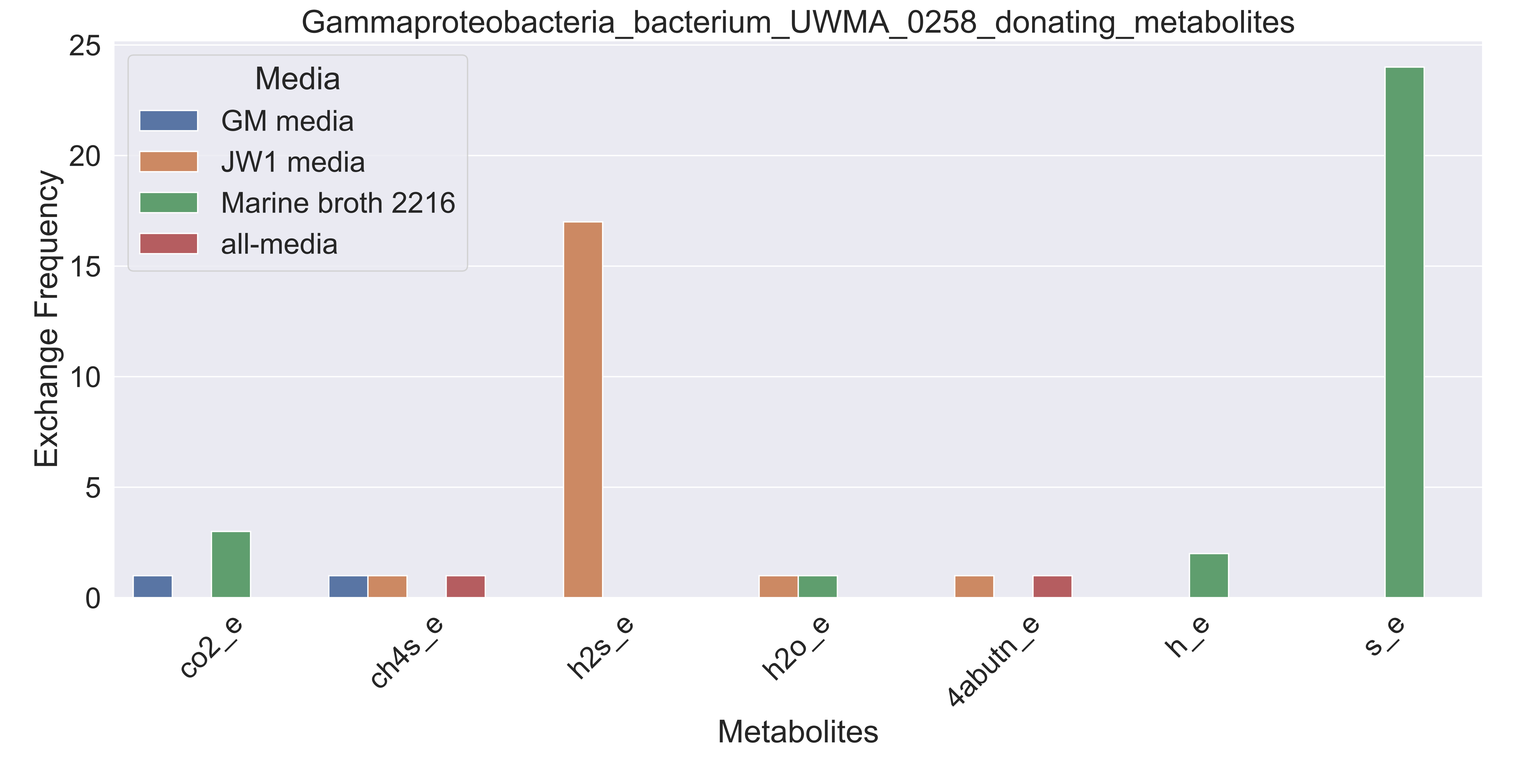

### Supplementary Figure S138 Gammaproteobacteria bacterium UWMA 0260 donating metabolites.pdf

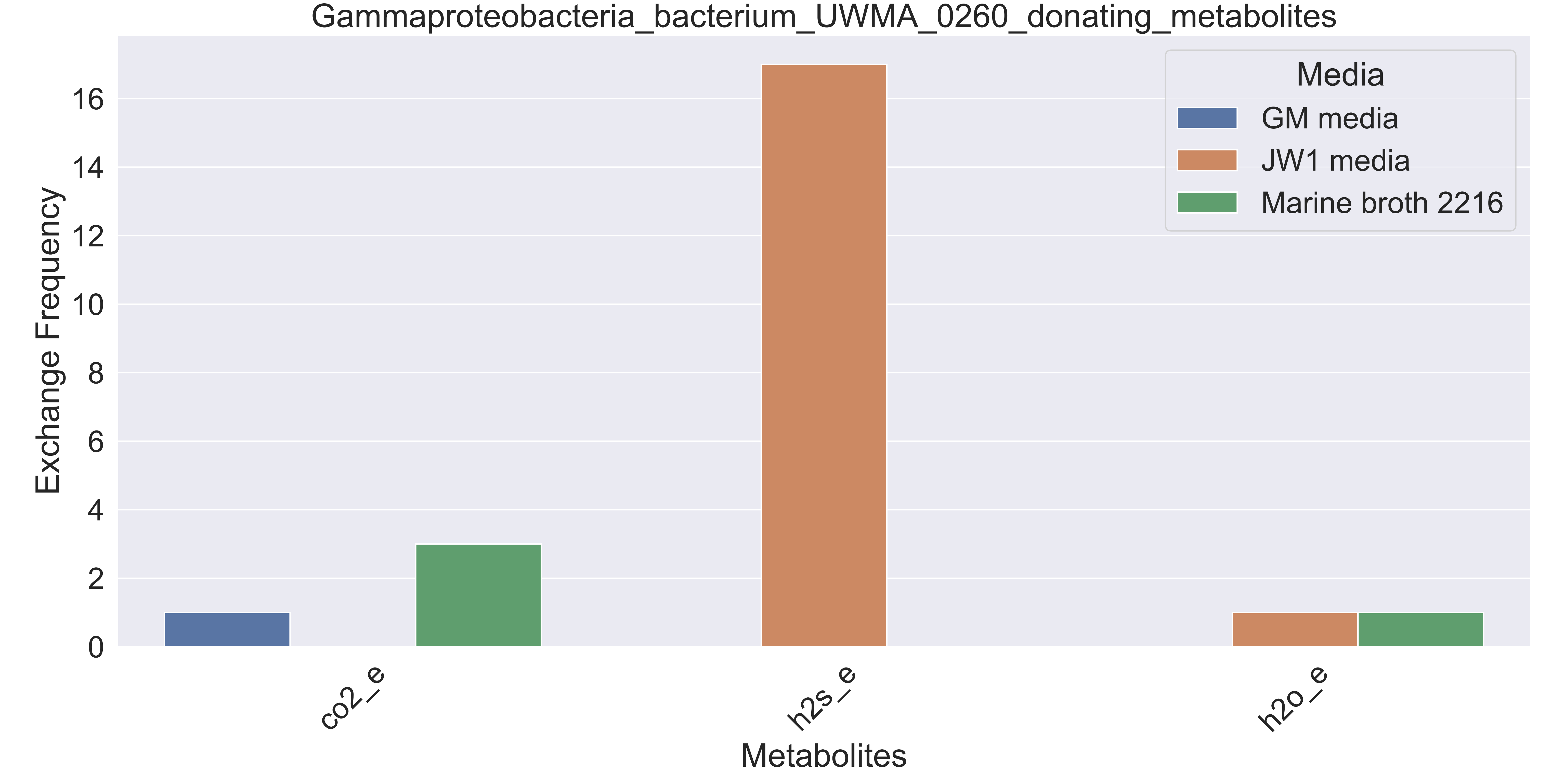

### Supplementary Figure S139 Gammaproteobacteria bacterium UWMA 0261 donating metabolites.pdf

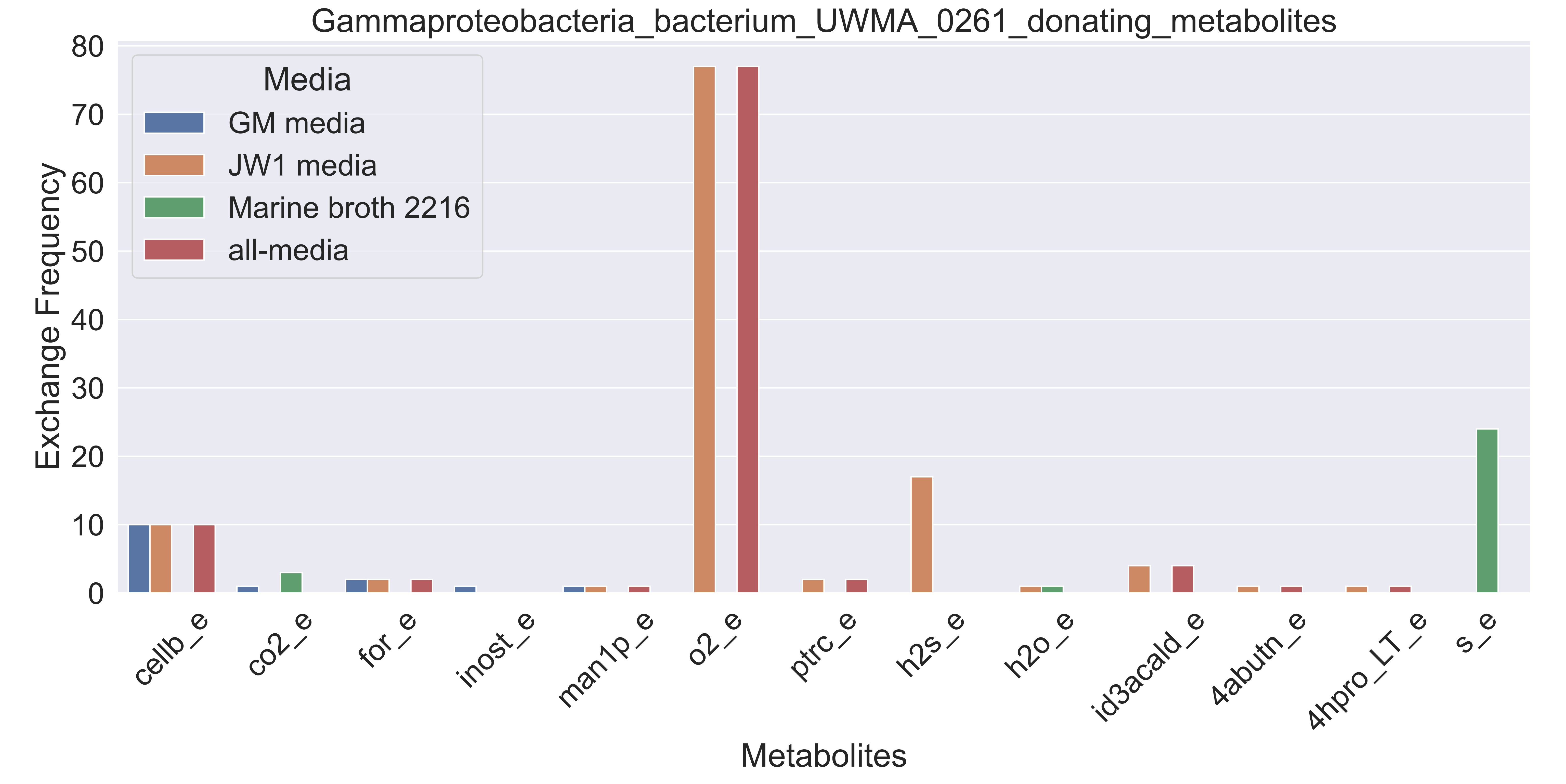

### Supplementary Figure S140 Gammaproteobacteria bacterium UWMA 0273 donating metabolites.pdf

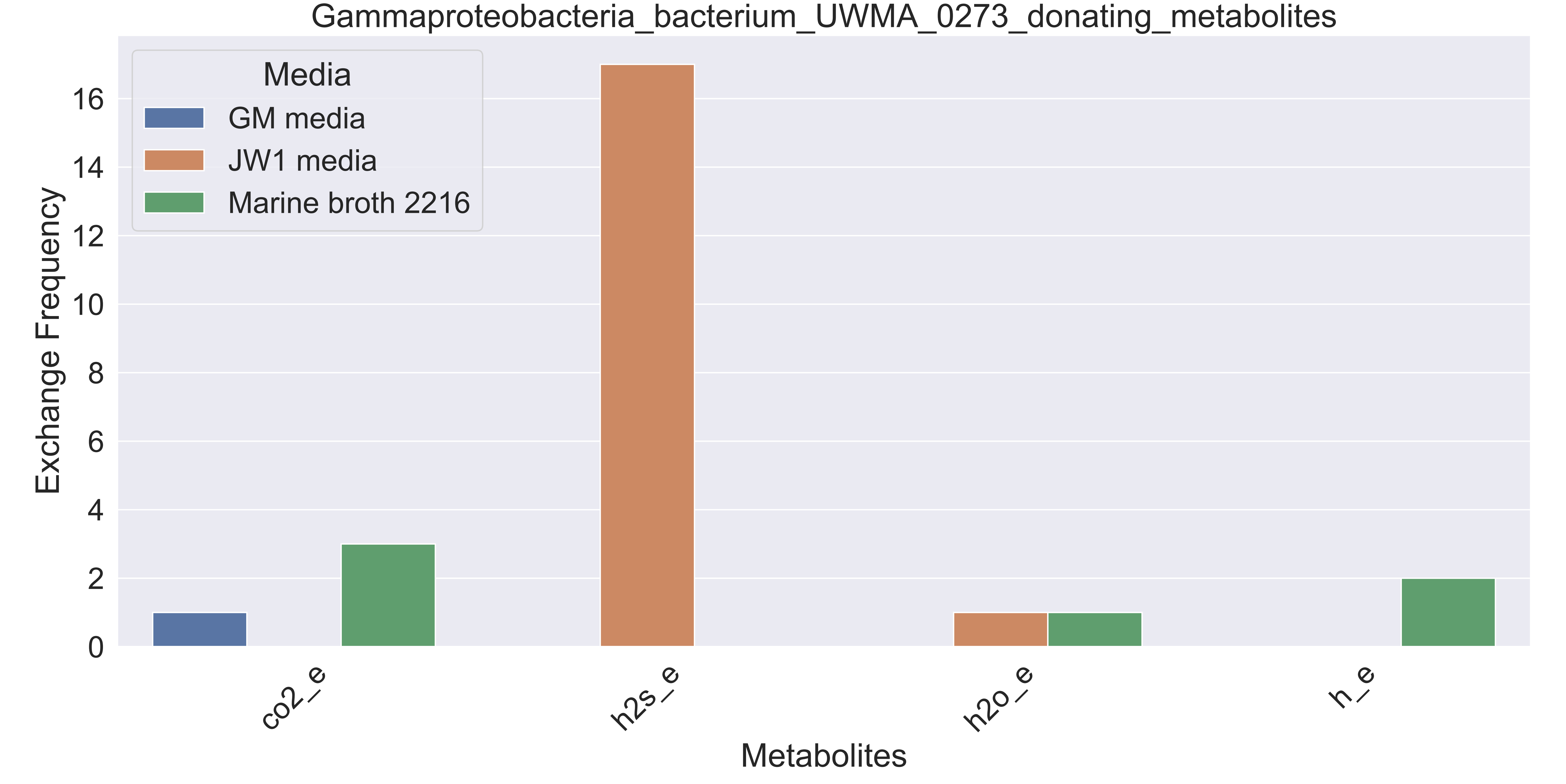

### Supplementary Figure S141 Gammaproteobacteria bacterium UWMA 0278 donating metabolites.pdf

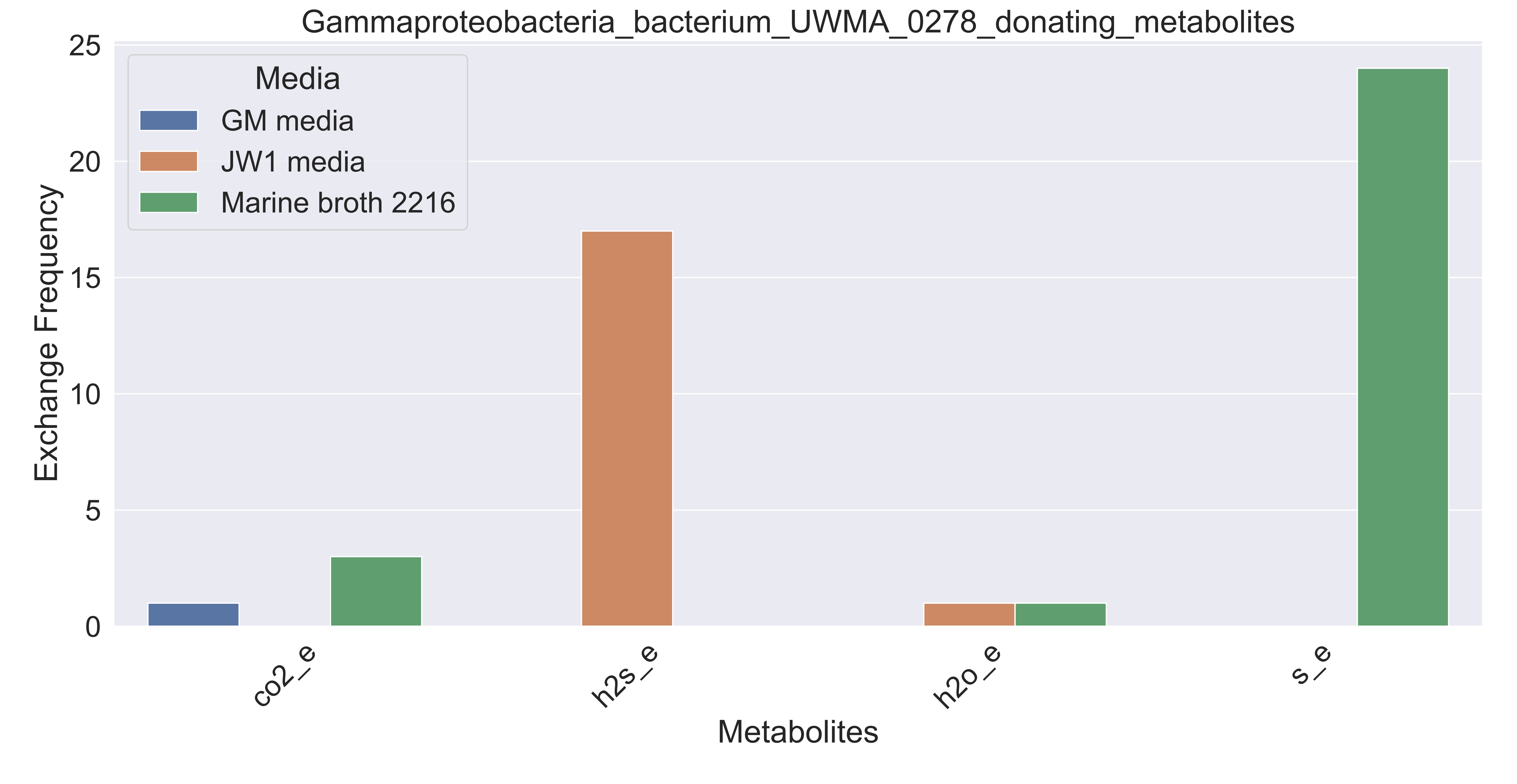

### Supplementary Figure S142 Gammaproteobacteria bacterium UWMA 0281 donating metabolites.pdf

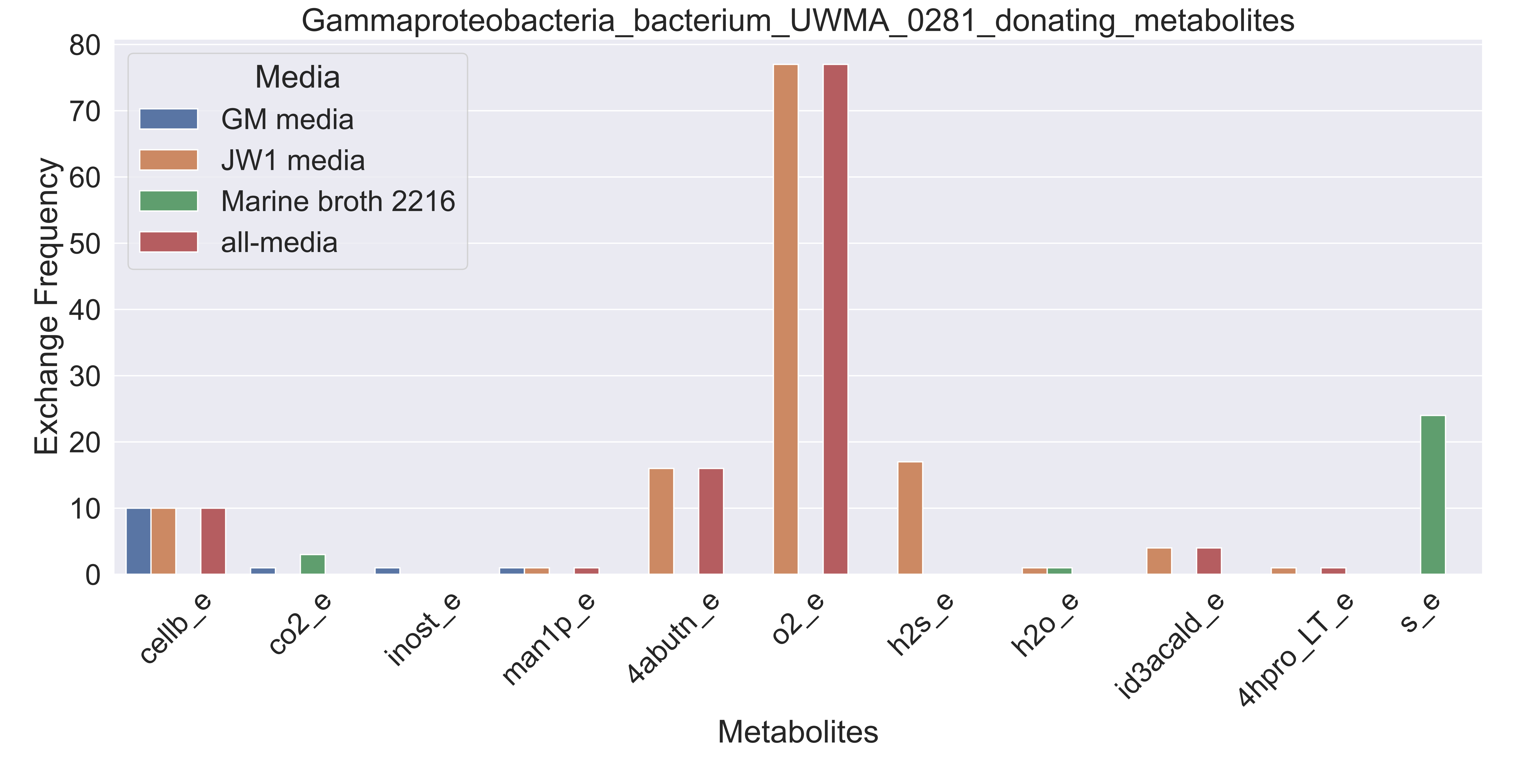

### Supplementary Figure S143 Gammaproteobacteria bacterium UWMA 0282 donating metabolites.pdf

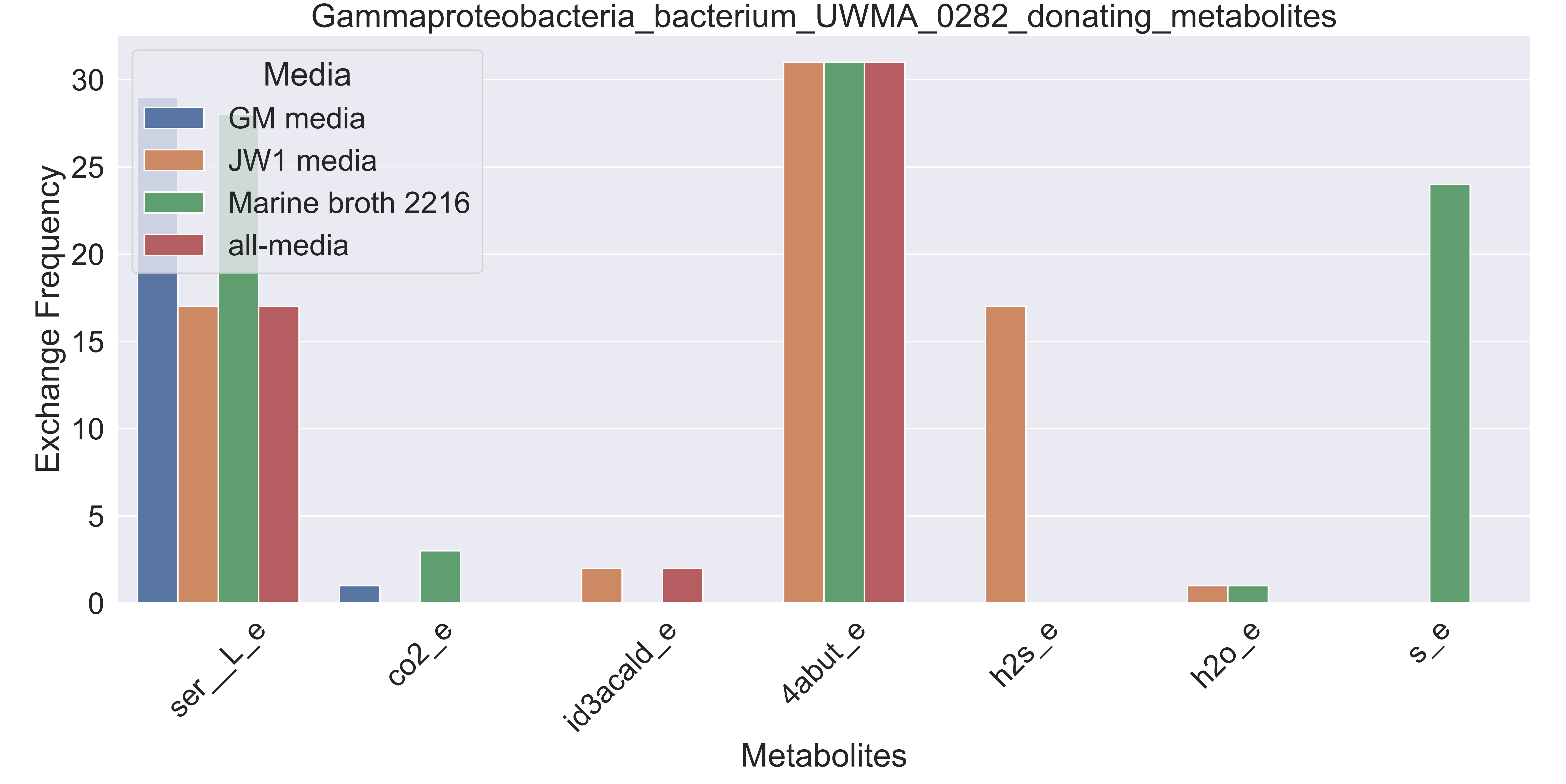

### Supplementary Figure S144 Gammaproteobacteria bacterium UWMA 0291 donating metabolites.pdf

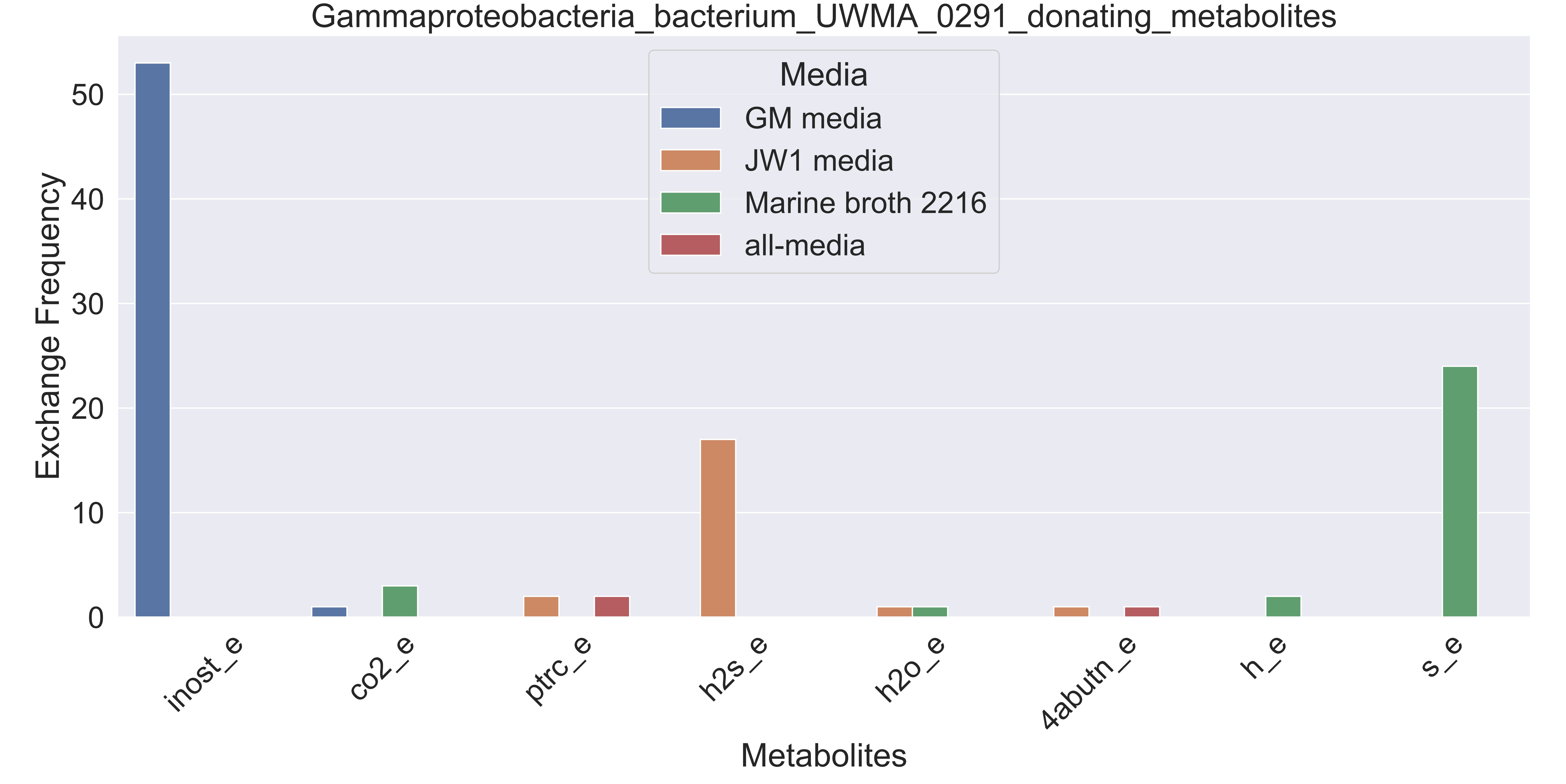

### Supplementary Figure S145 Gammaproteobacteria bacterium UWMA 0299 donating metabolites.pdf

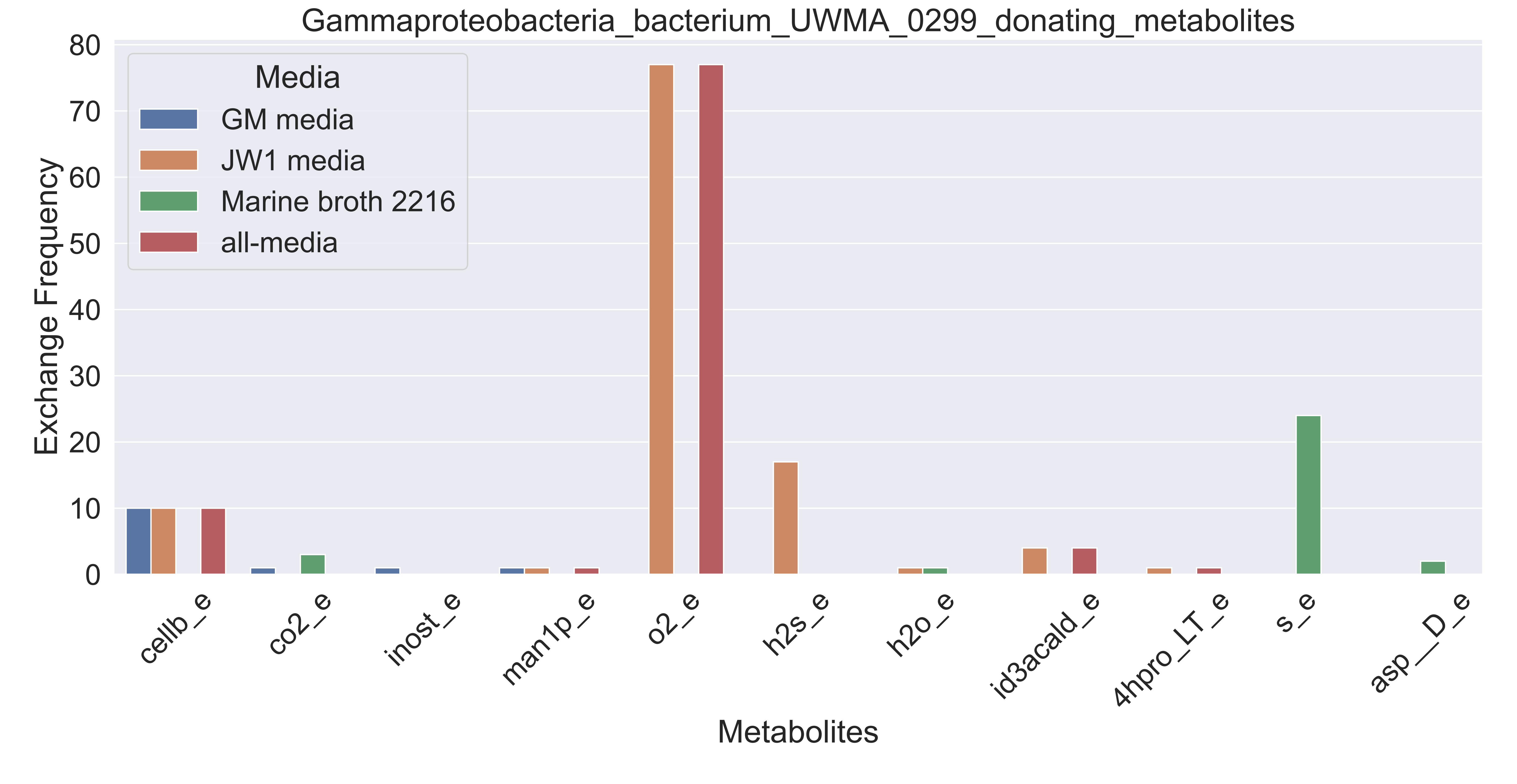

### Supplementary Figure S146 Gammaproteobacteria bacterium UWMA 0301 donating metabolites.pdf

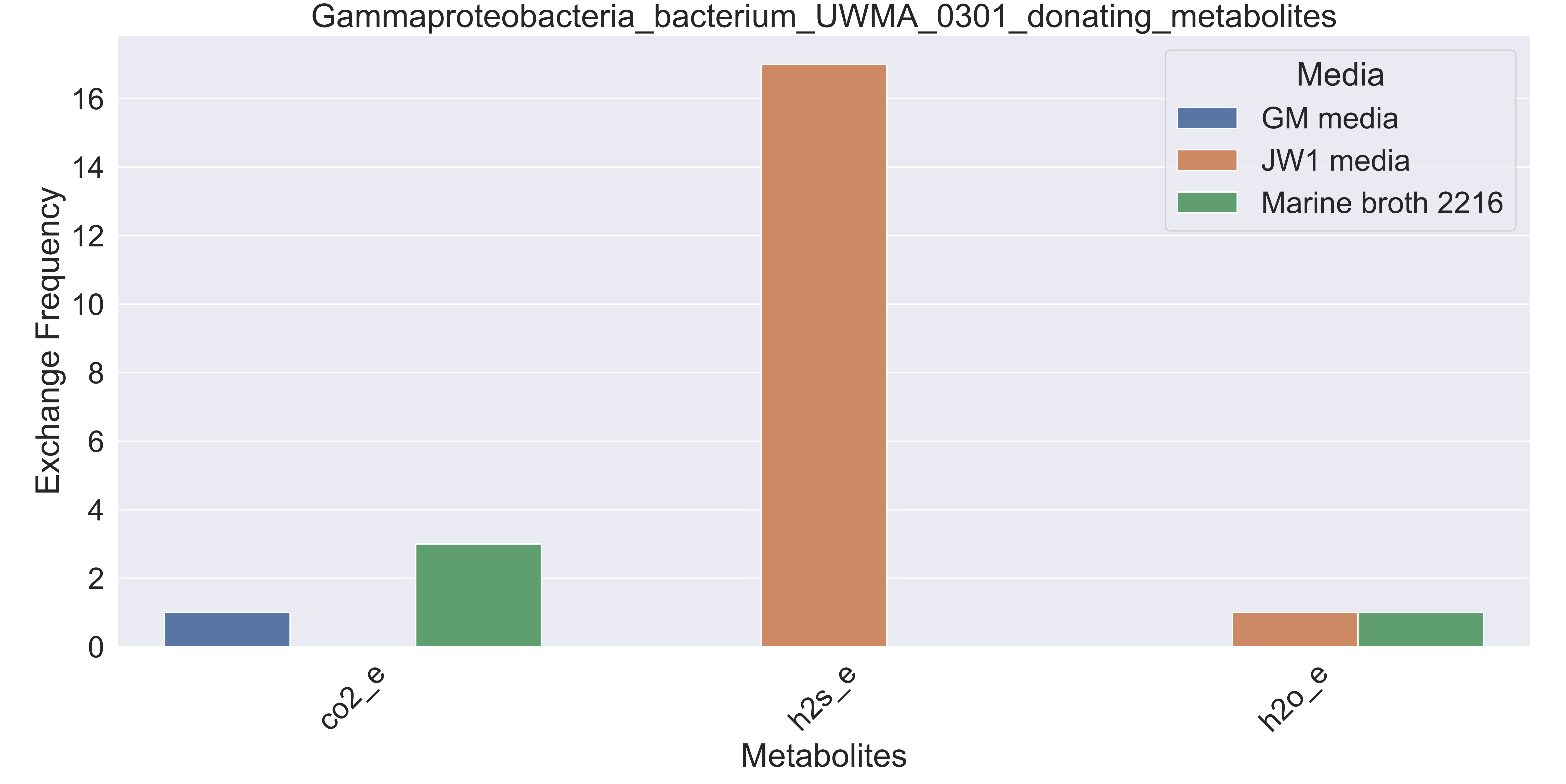

### Supplementary Figure S147 Gammaproteobacteria bacterium UWMA 0302 donating metabolites.pdf

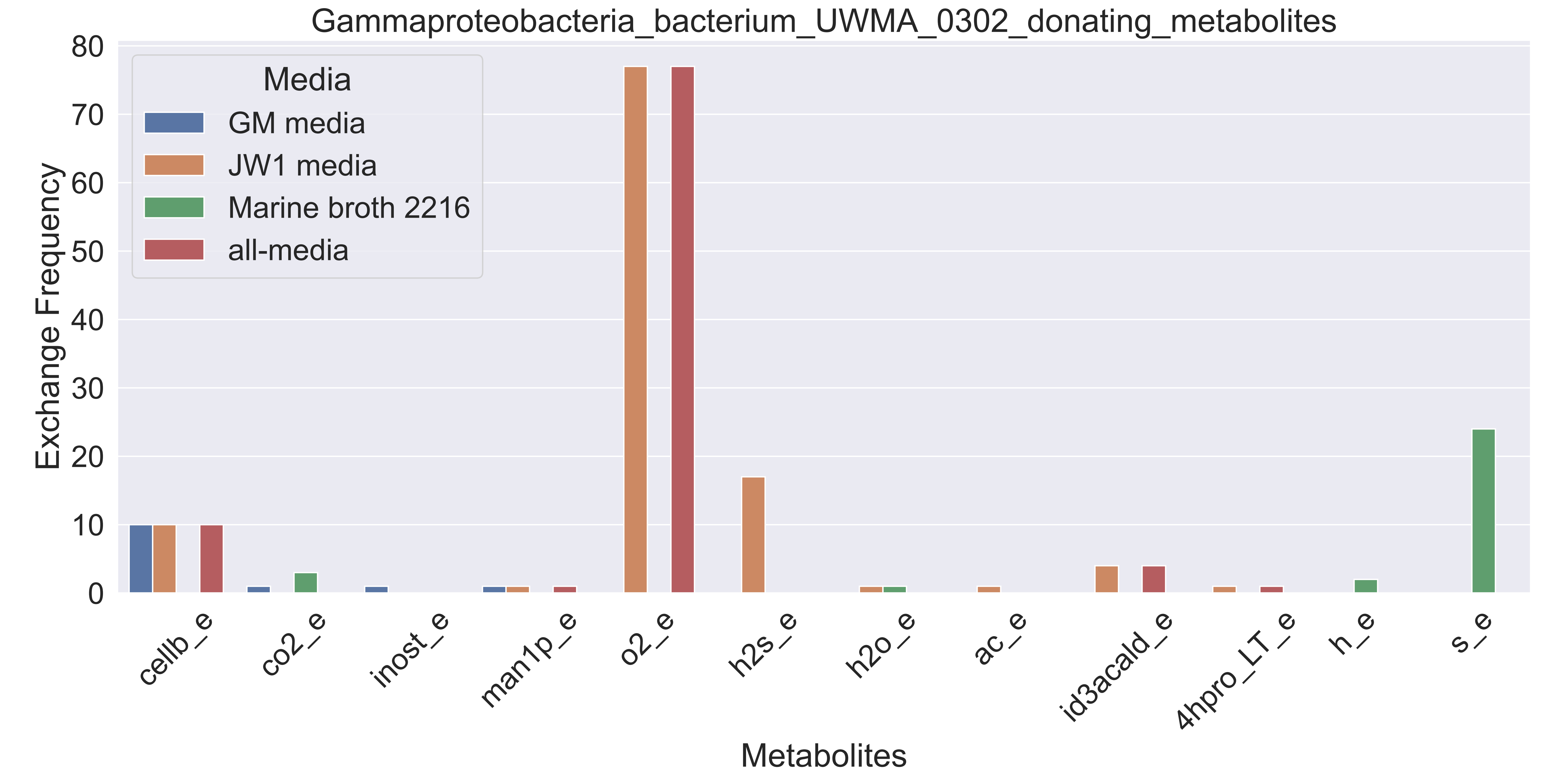

### Supplementary Figure S148 Gammaproteobacteria bacterium UWMA 0343 donating metabolites.pdf

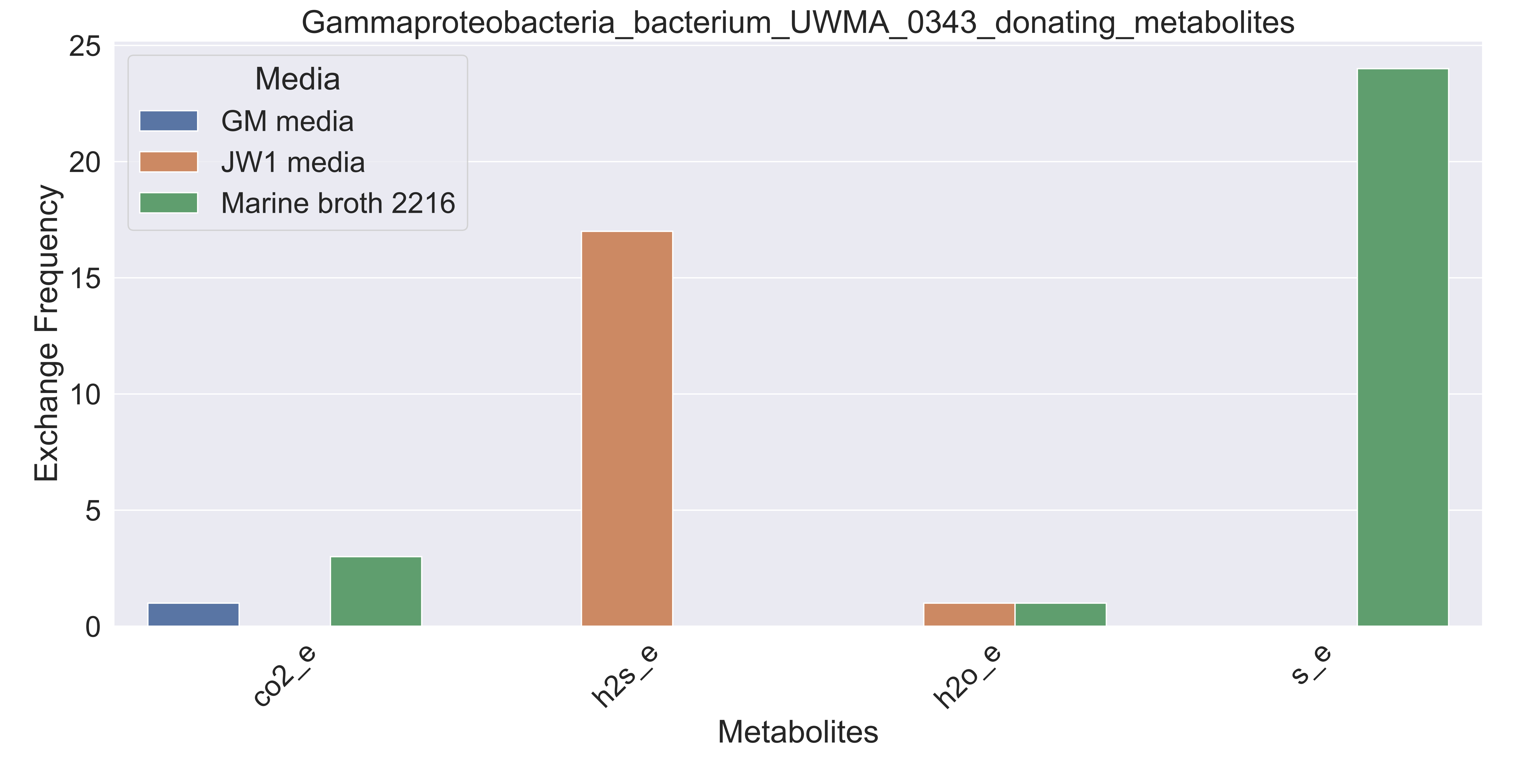

### Supplementary Figure S149 Gemmatimonadetes bacterium UWMA 0296 donating metabolites.pdf

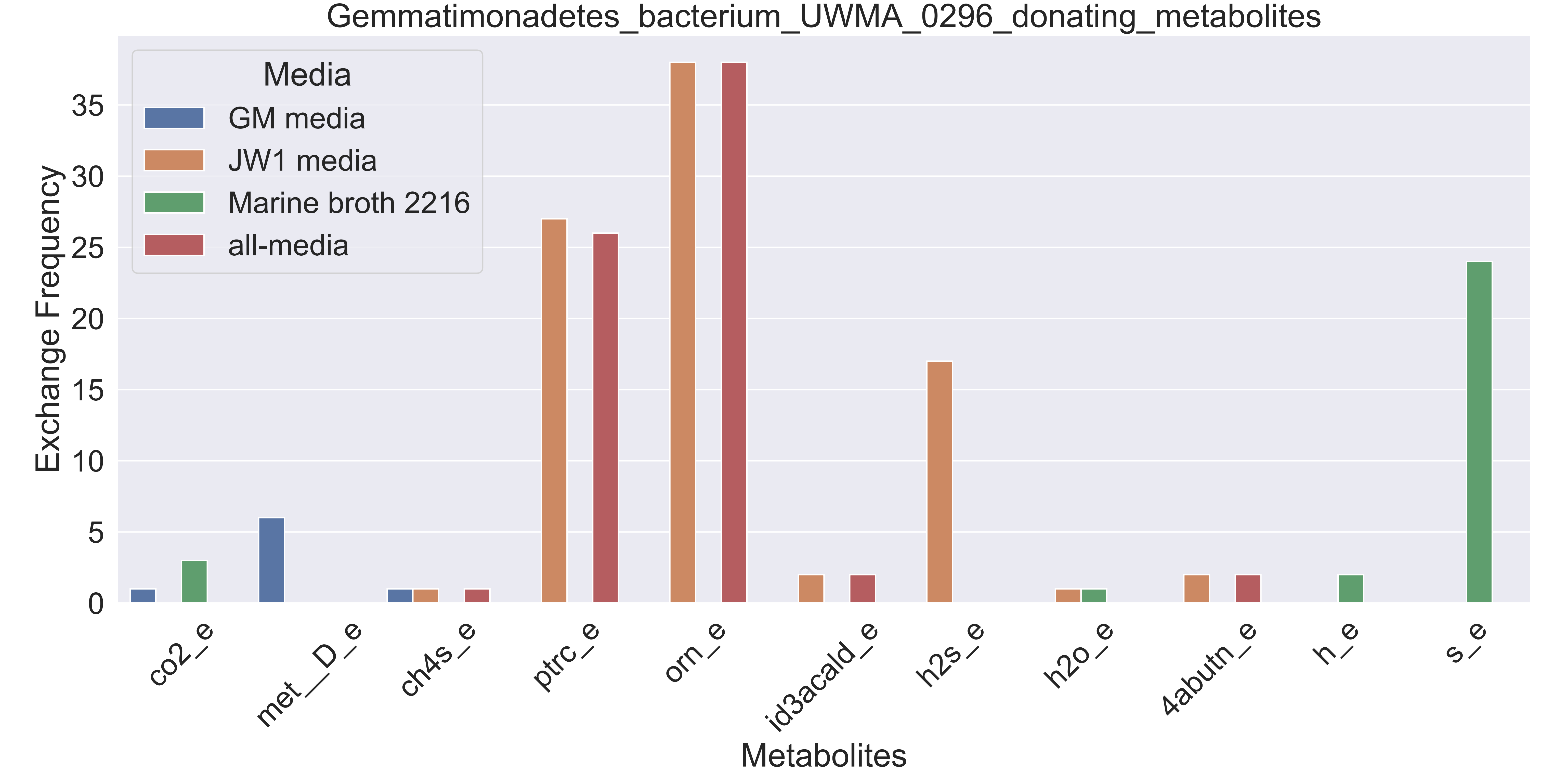

### Supplementary Figure S150 Gemmatimonadetes bacterium UWMA 0308 donating metabolites.pdf

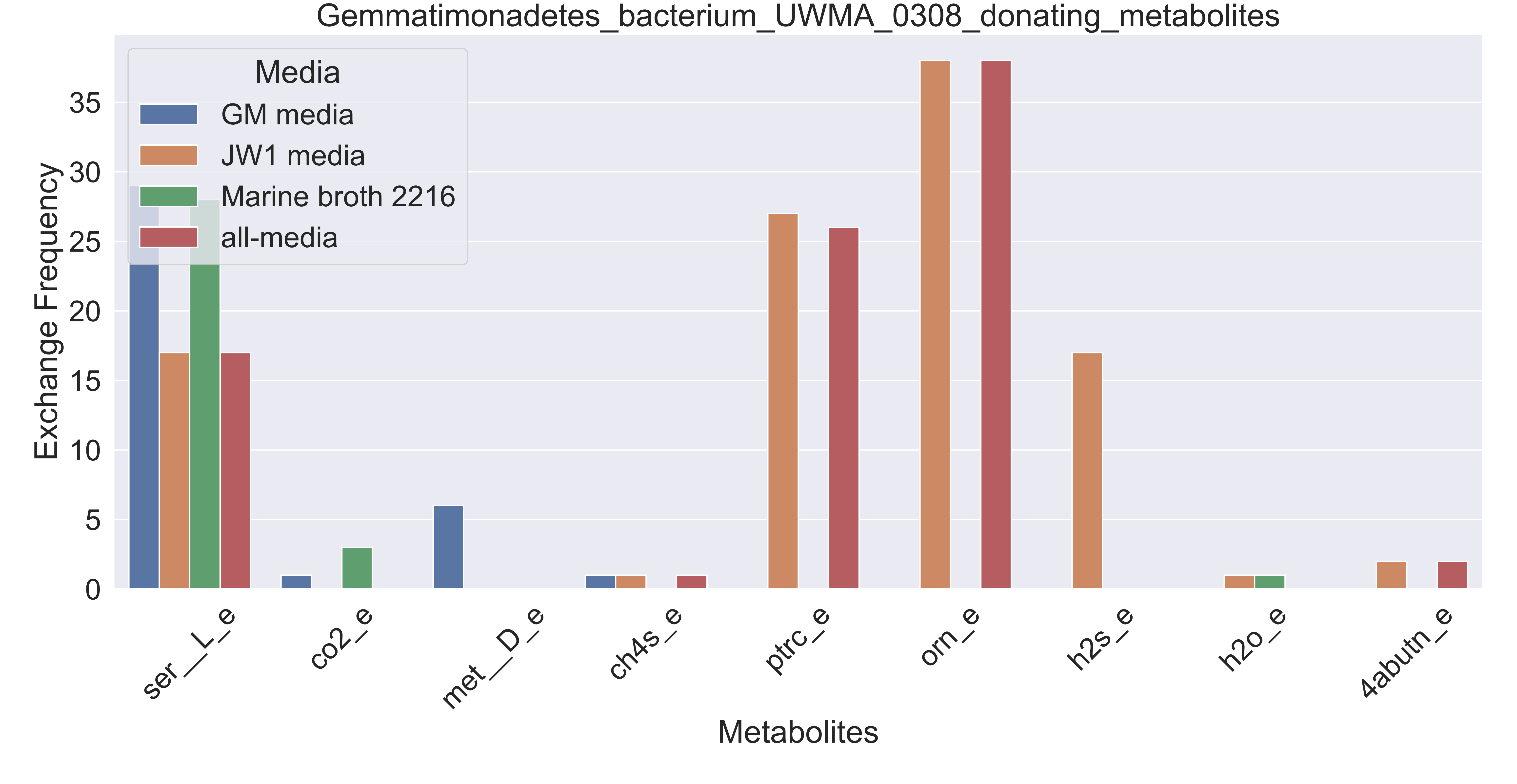

### Supplementary Figure S151 Gemmatimonadetes bacterium UWMA 0334 donating metabolites.pdf

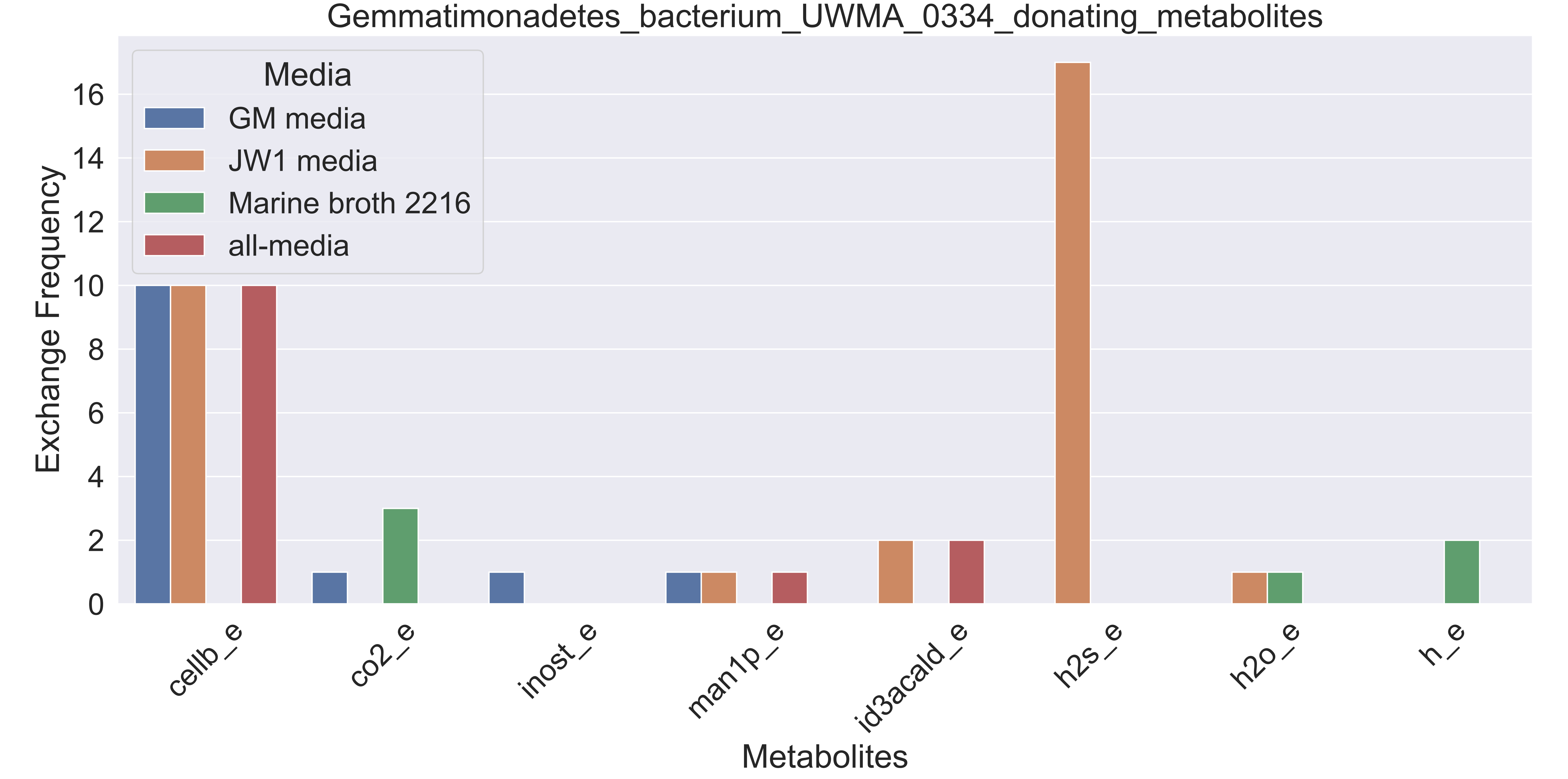

### Supplementary Figure S152 Gemmatimonadetes bacterium UWMA 0339 donating metabolites.pdf

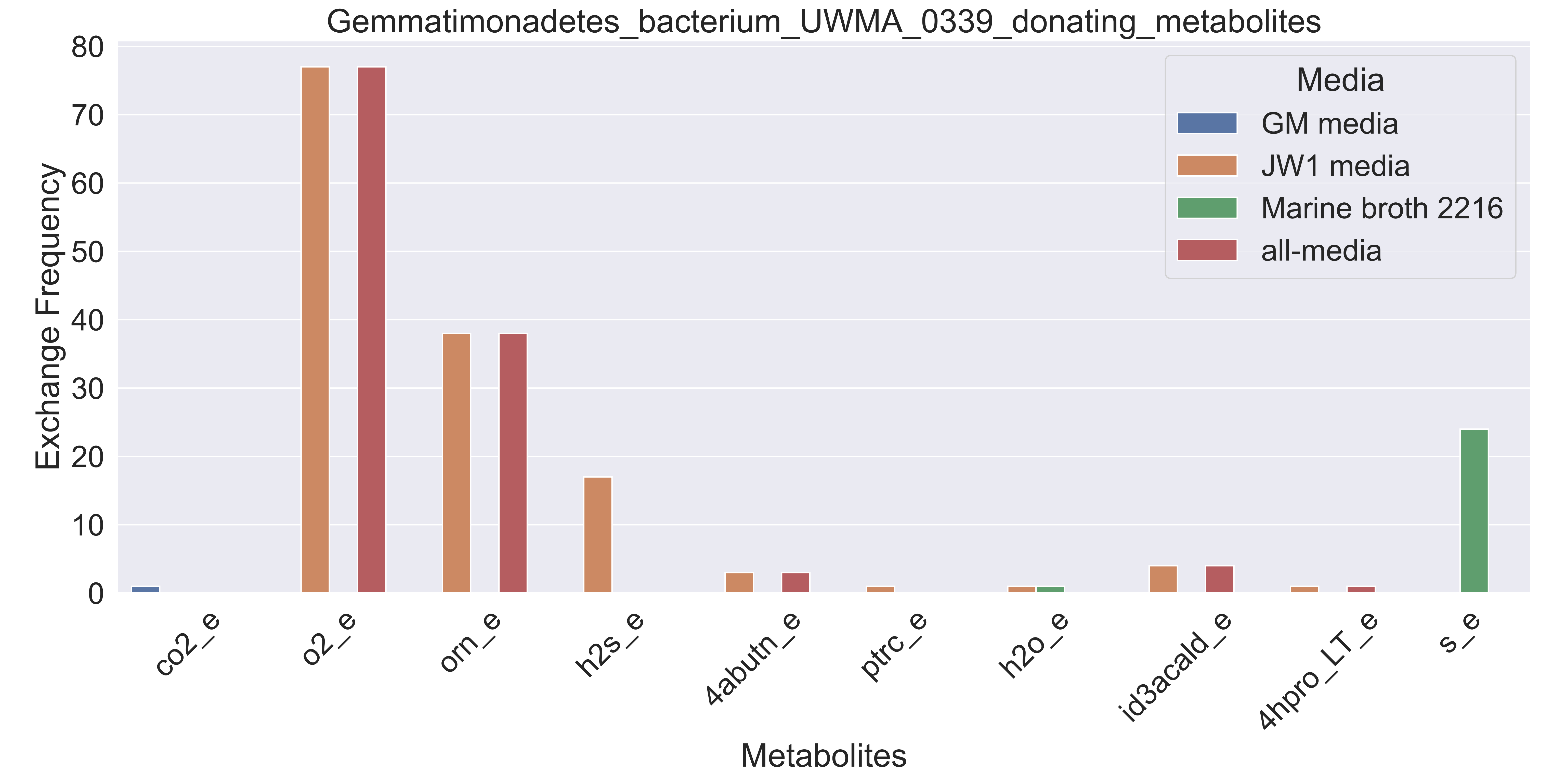

### Supplementary Figure S153 Henriciella sp UWMA 0297 donating metabolites.pdf

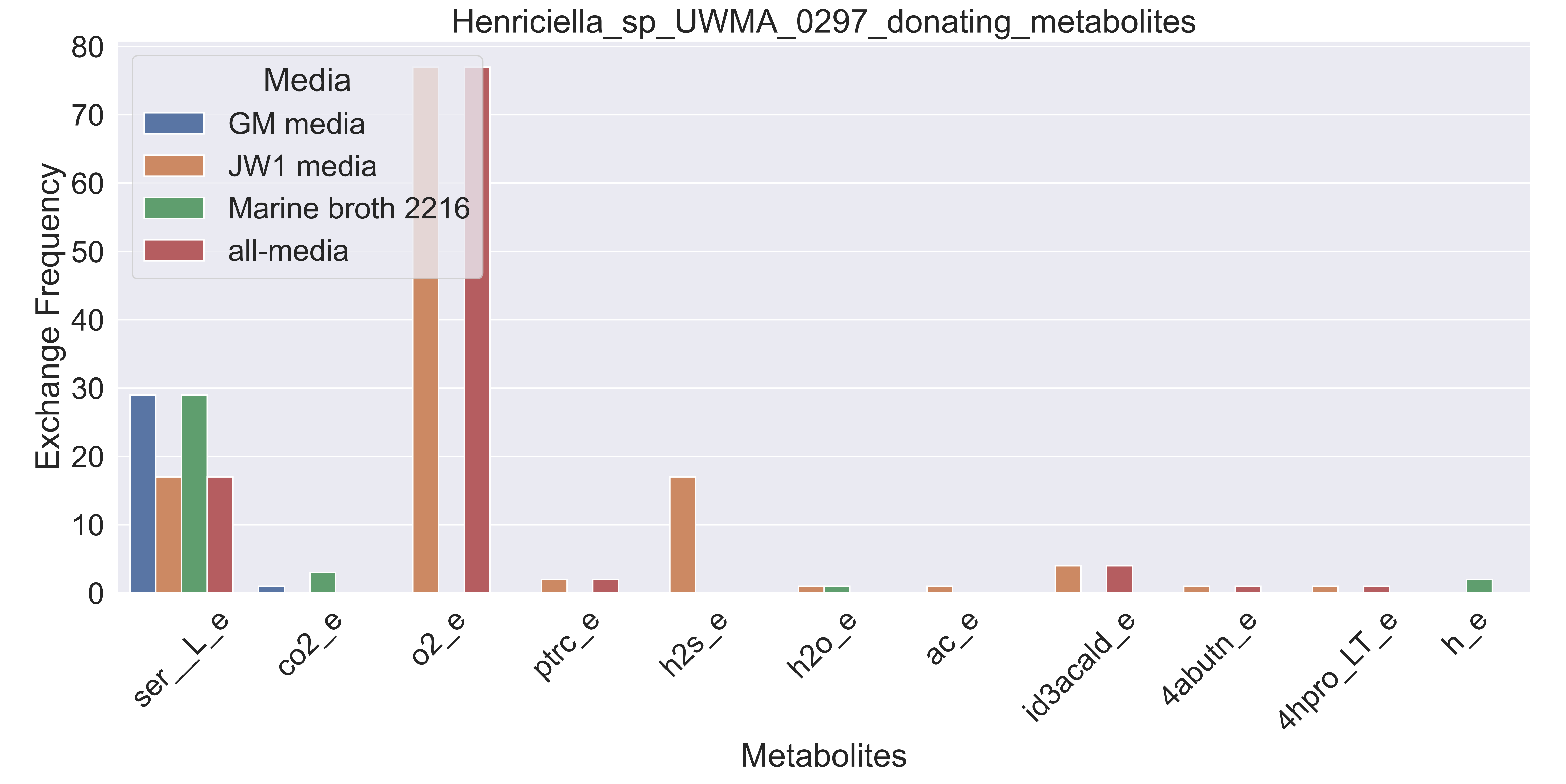

### Supplementary Figure S154 Marine Group III euryarchaeote UWMA 0284 donating metabolites.pdf

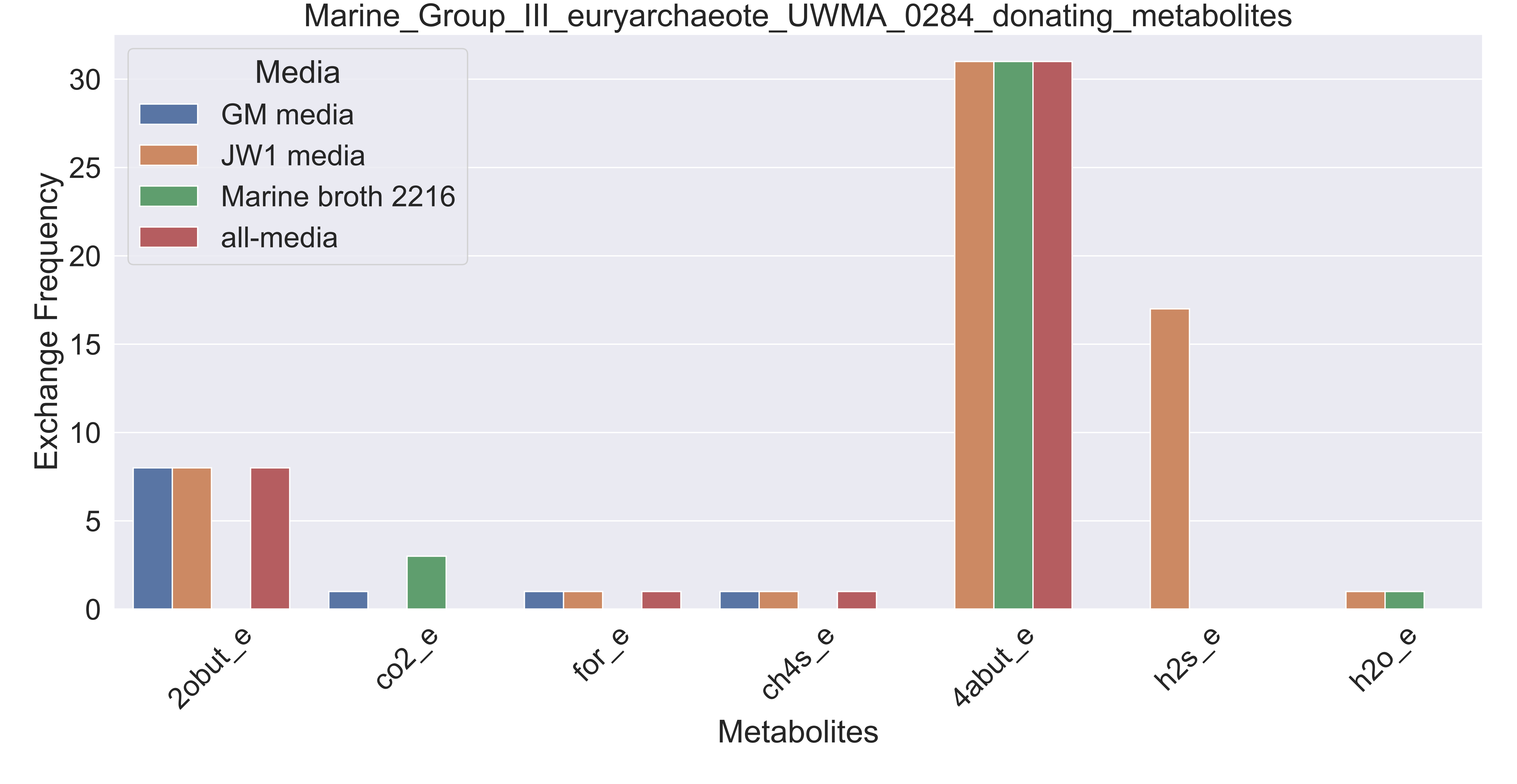

### Supplementary Figure S155 Marine Group III euryarchaeote UWMA 0340 donating metabolites.pdf

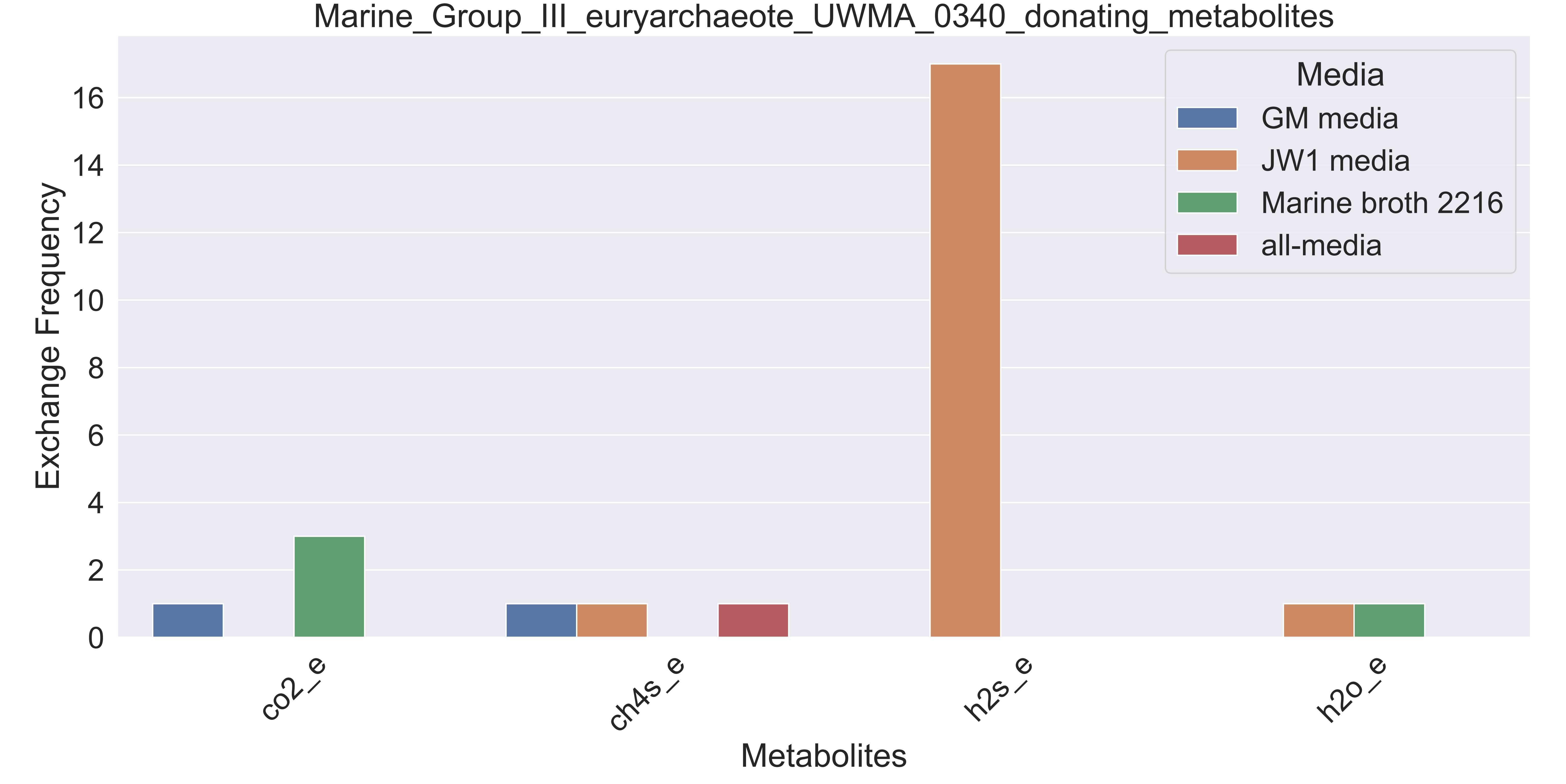

### Supplementary Figure S156 Marine Group II euryarchaeote UWMA 0266 donating metabolites.pdf

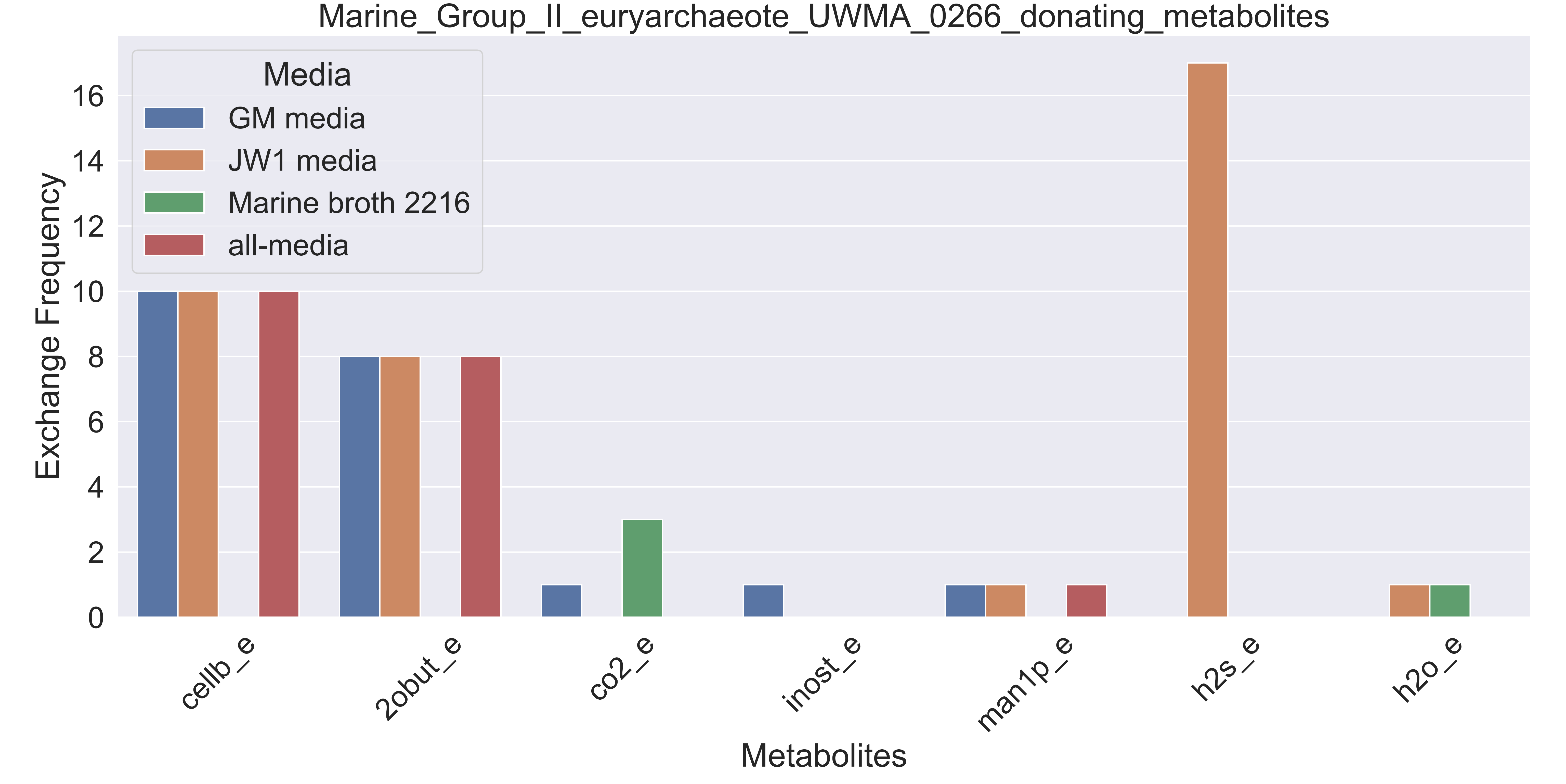

### Supplementary Figure S157 Marine Group II euryarchaeote UWMA 0275 donating metabolites.pdf

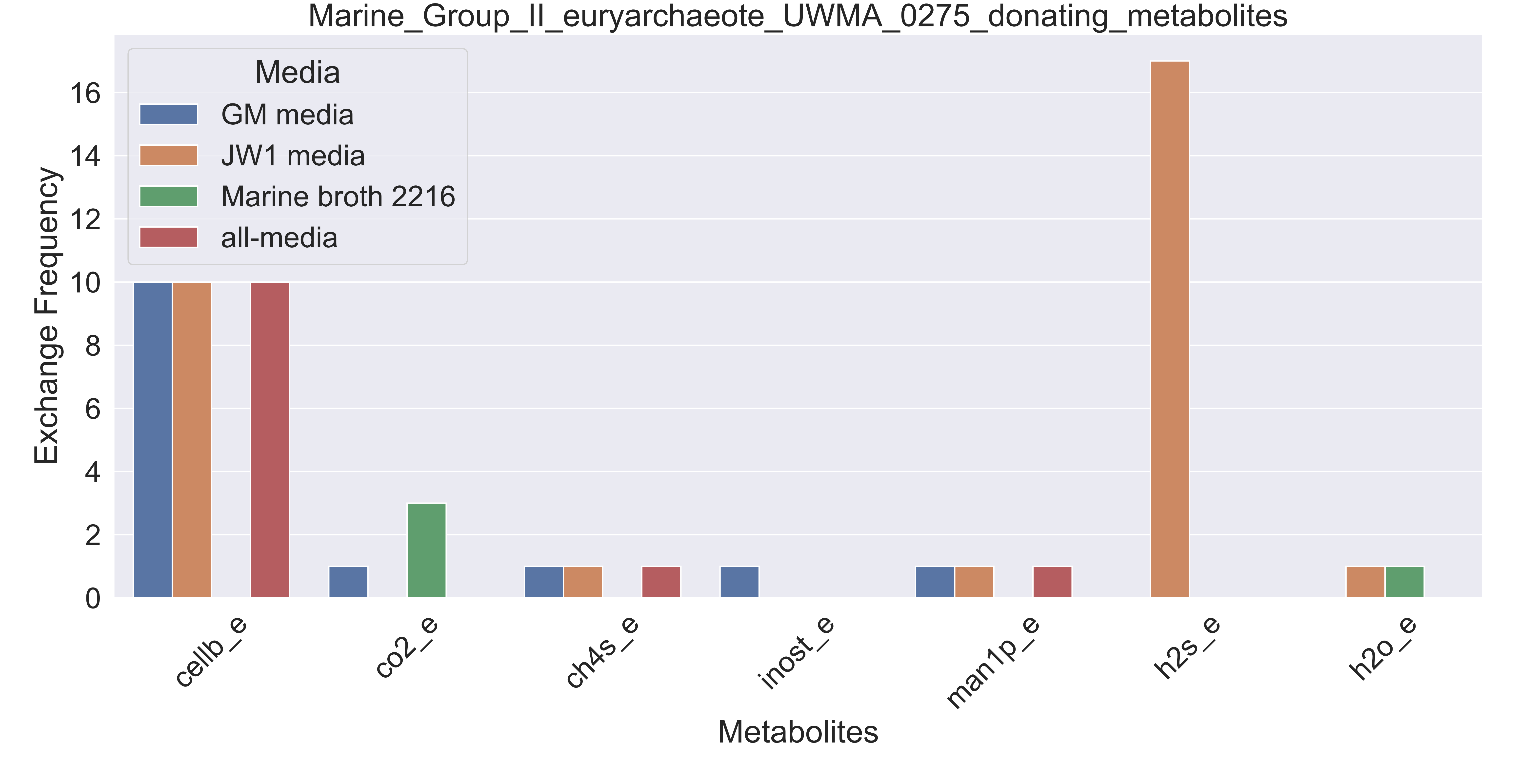

### Supplementary Figure S158 Marine Group II euryarchaeote UWMA 0279 donating metabolites.pdf

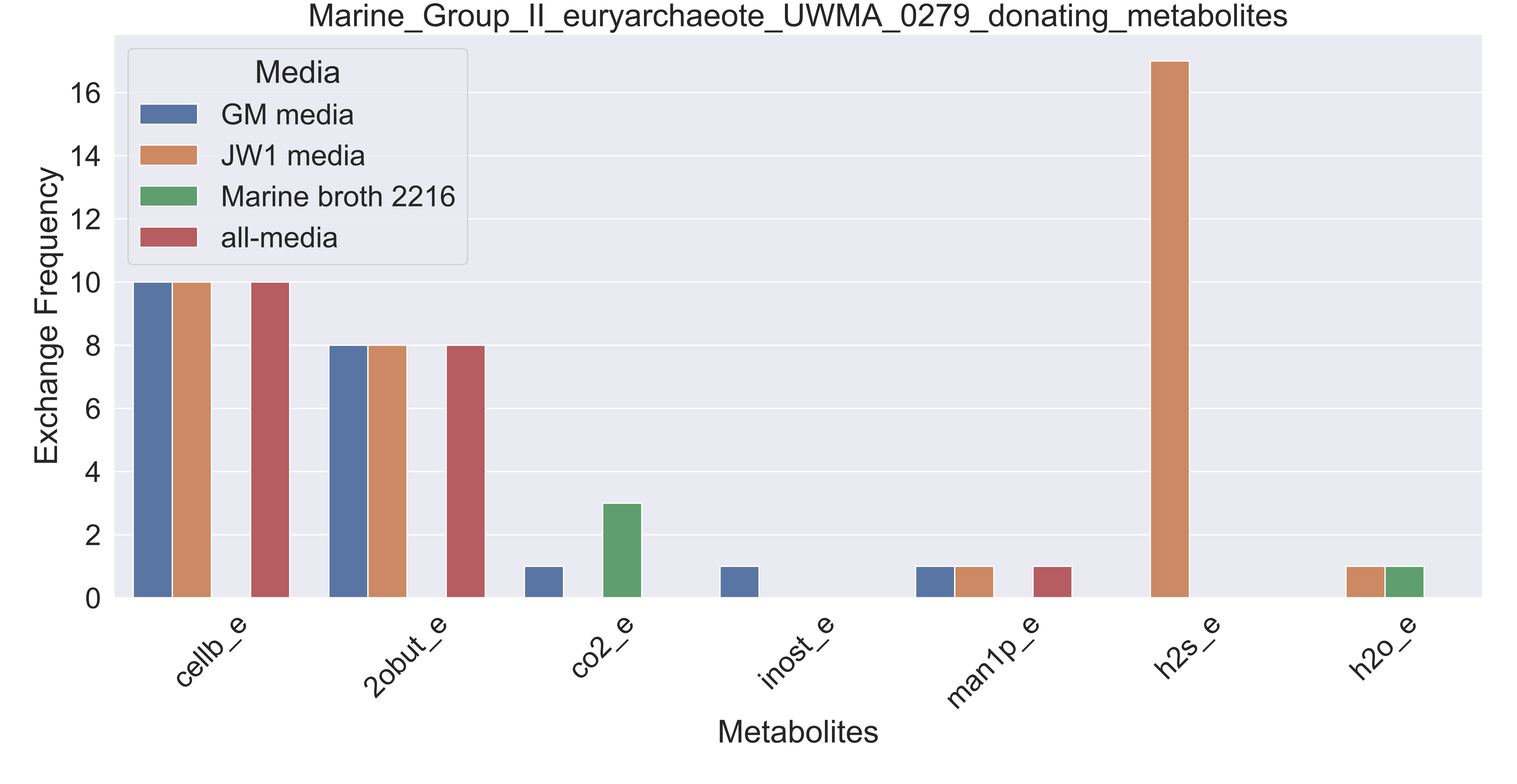

### Supplementary Figure S159 Marine Group II euryarchaeote UWMA 0283 donating metabolites.pdf

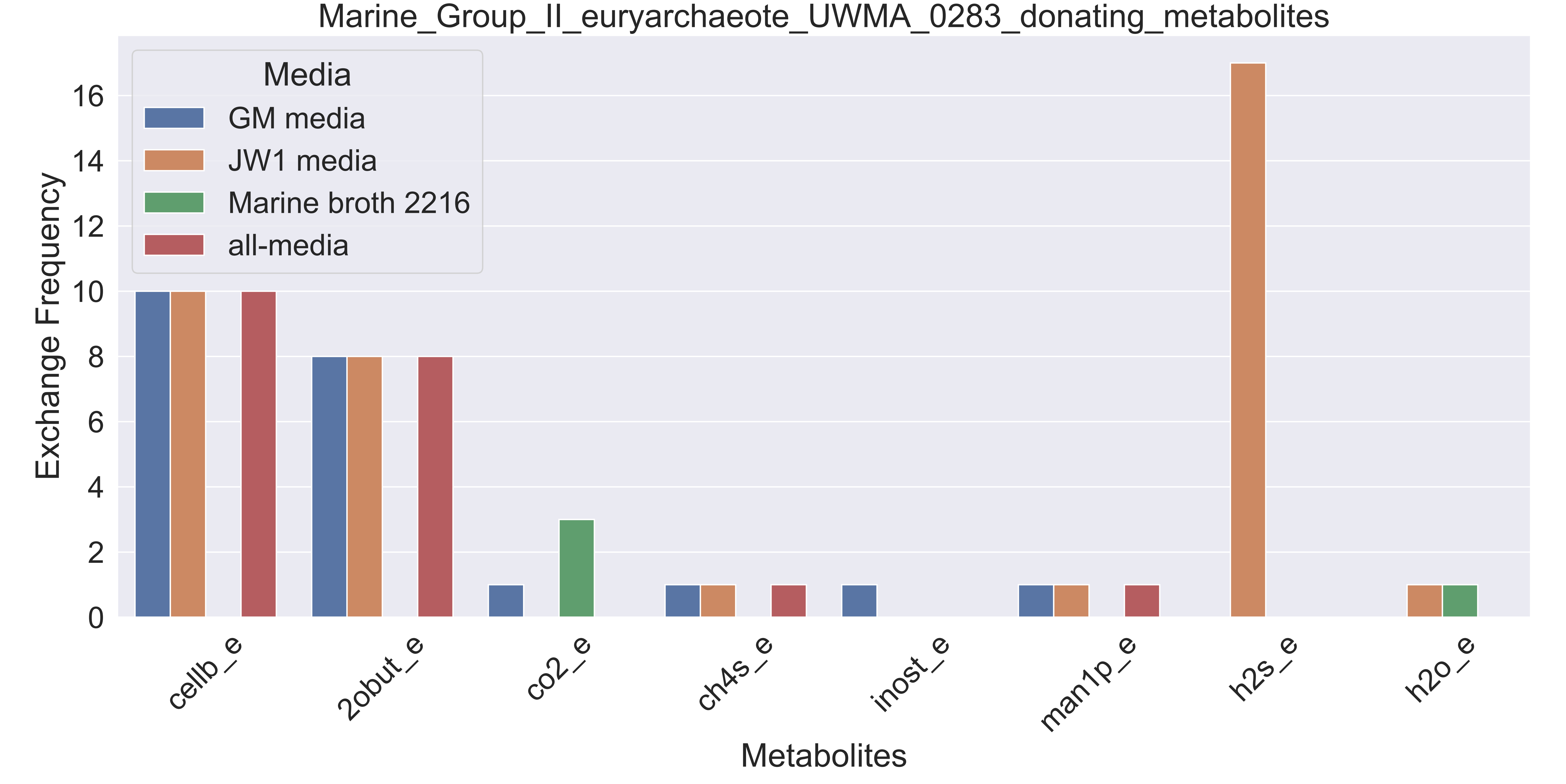

### Supplementary Figure S160 Marine Group II euryarchaeote UWMA 0323 donating metabolites.pdf

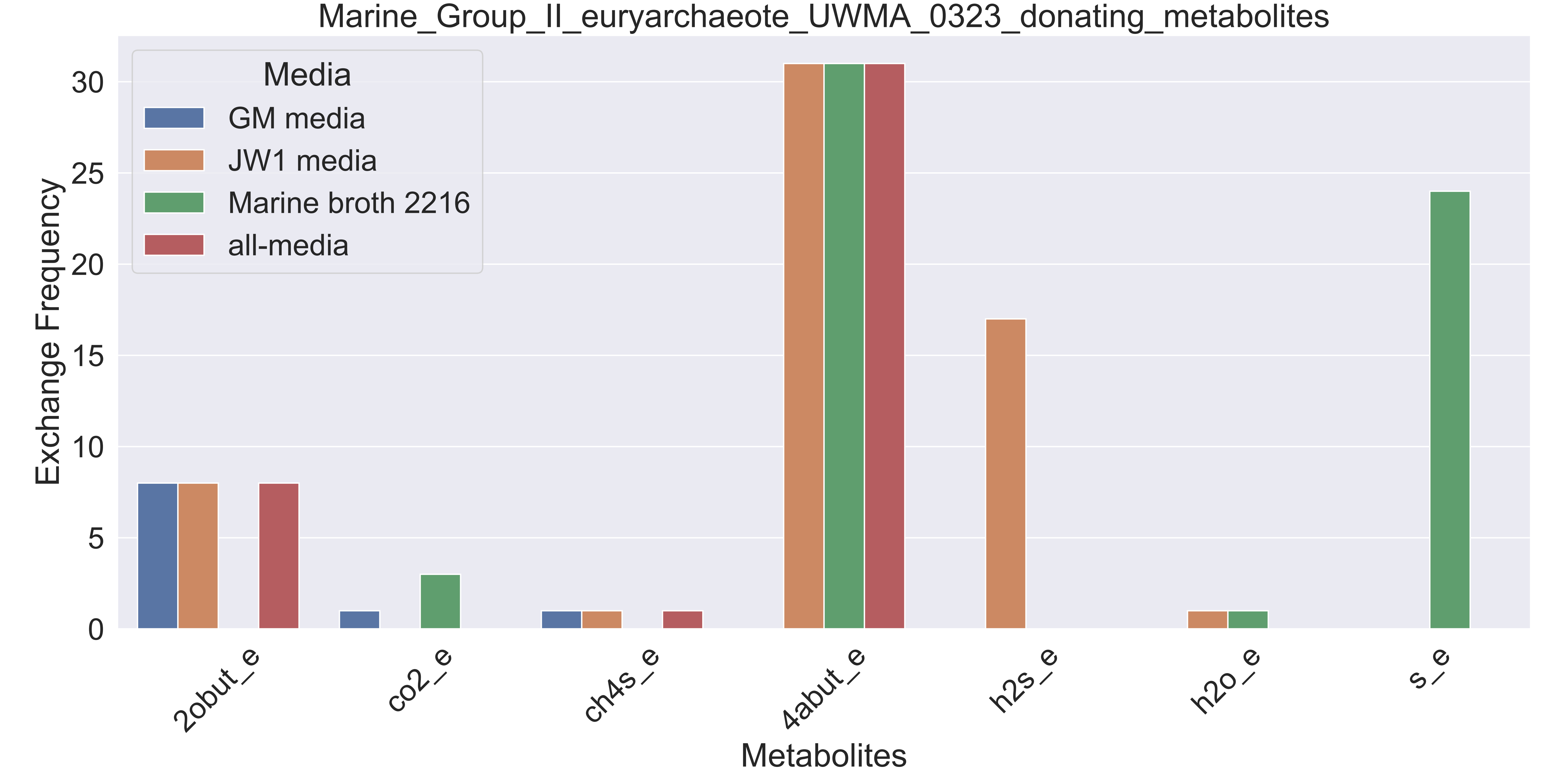

### Supplementary Figure S161 Marine Group II euryarchaeote UWMA 0328 donating metabolites.pdf

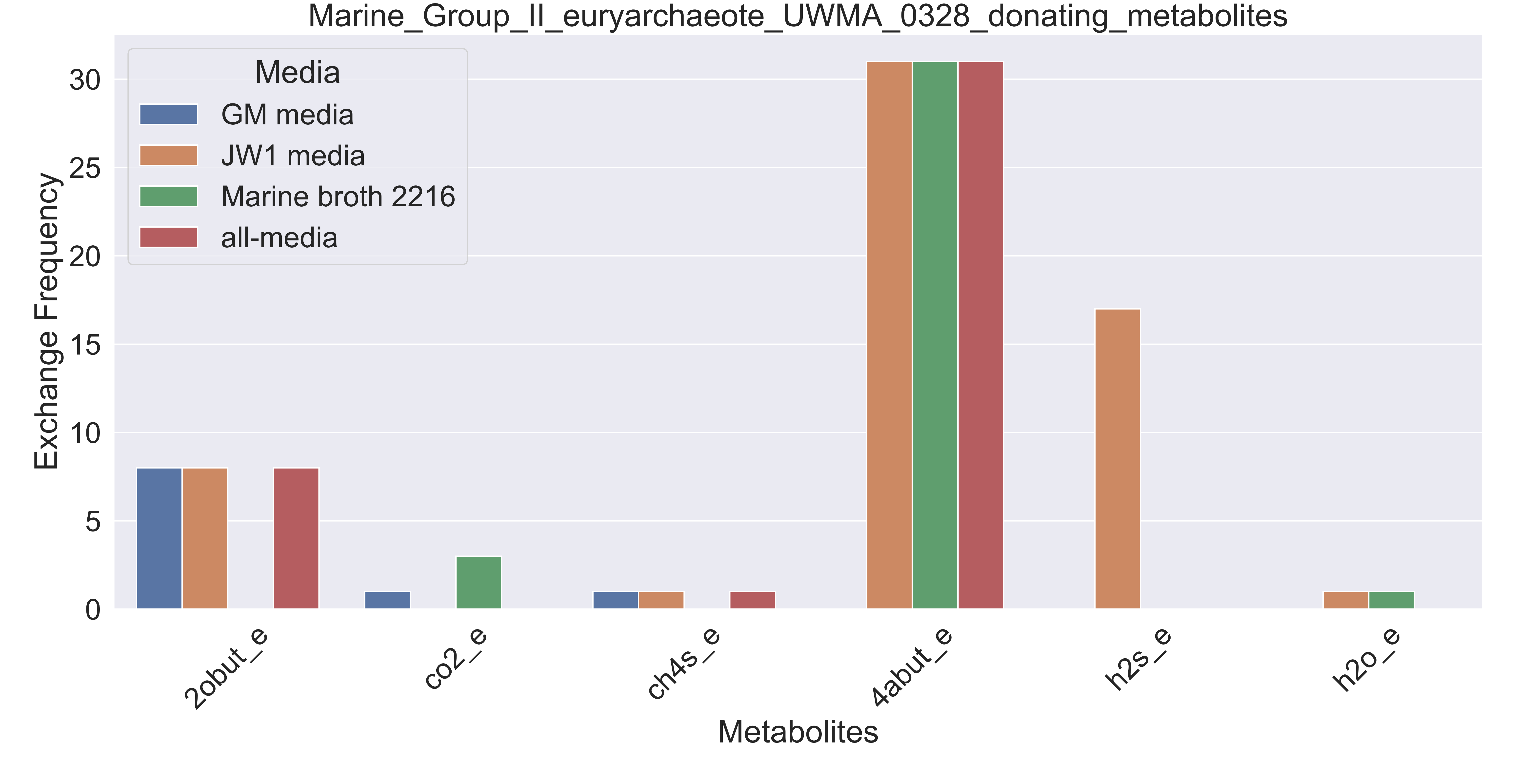

### Supplementary Figure S162 Marine Group II euryarchaeote UWMA 0344 donating metabolites.pdf

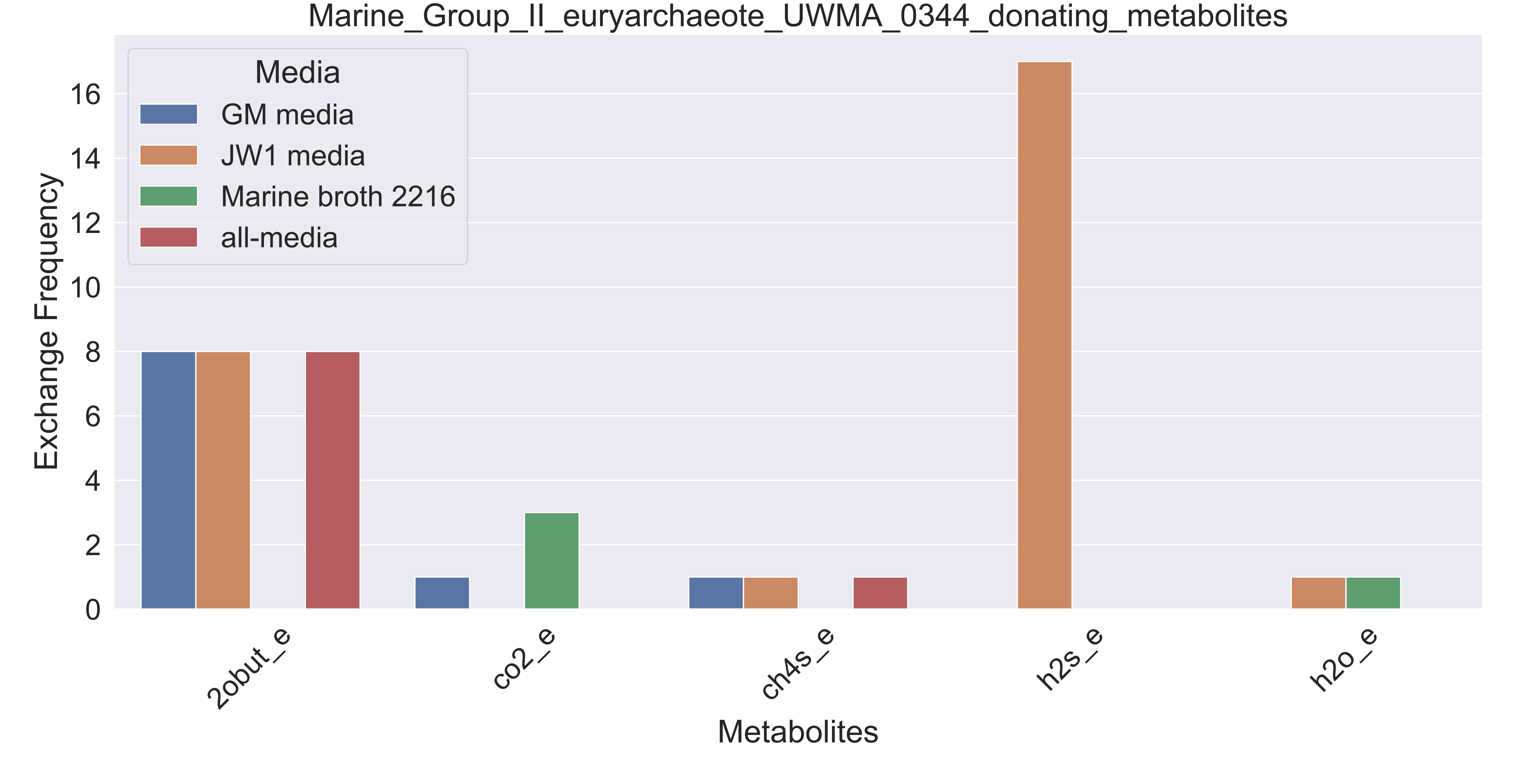

### Supplementary Figure S163 Marine Group II euryarchaeote UWMA 0350 donating metabolites.pdf

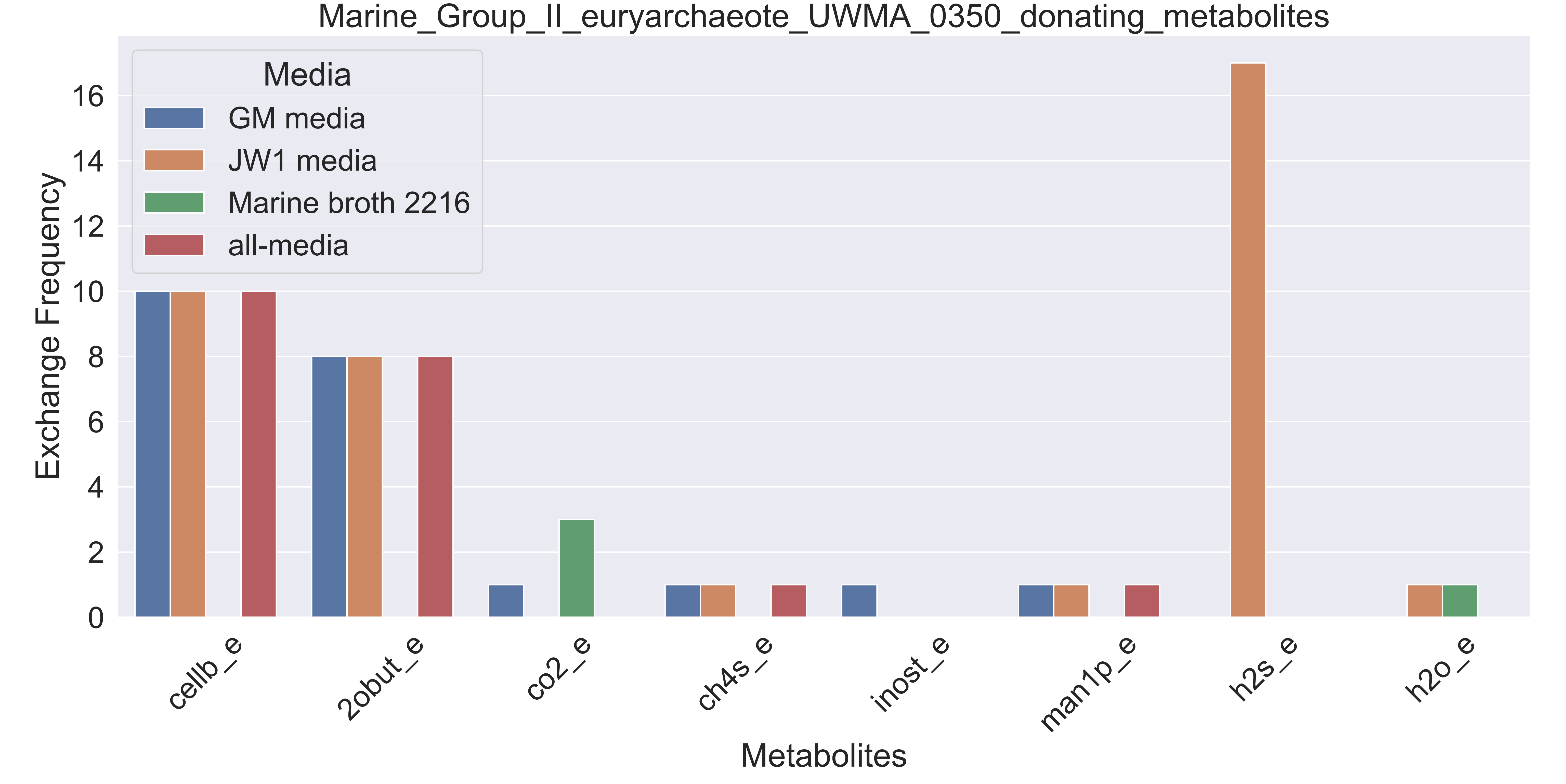

### Supplementary Figure S164 Marine Group II euryarchaeote UWMA 0352 donating metabolites.pdf

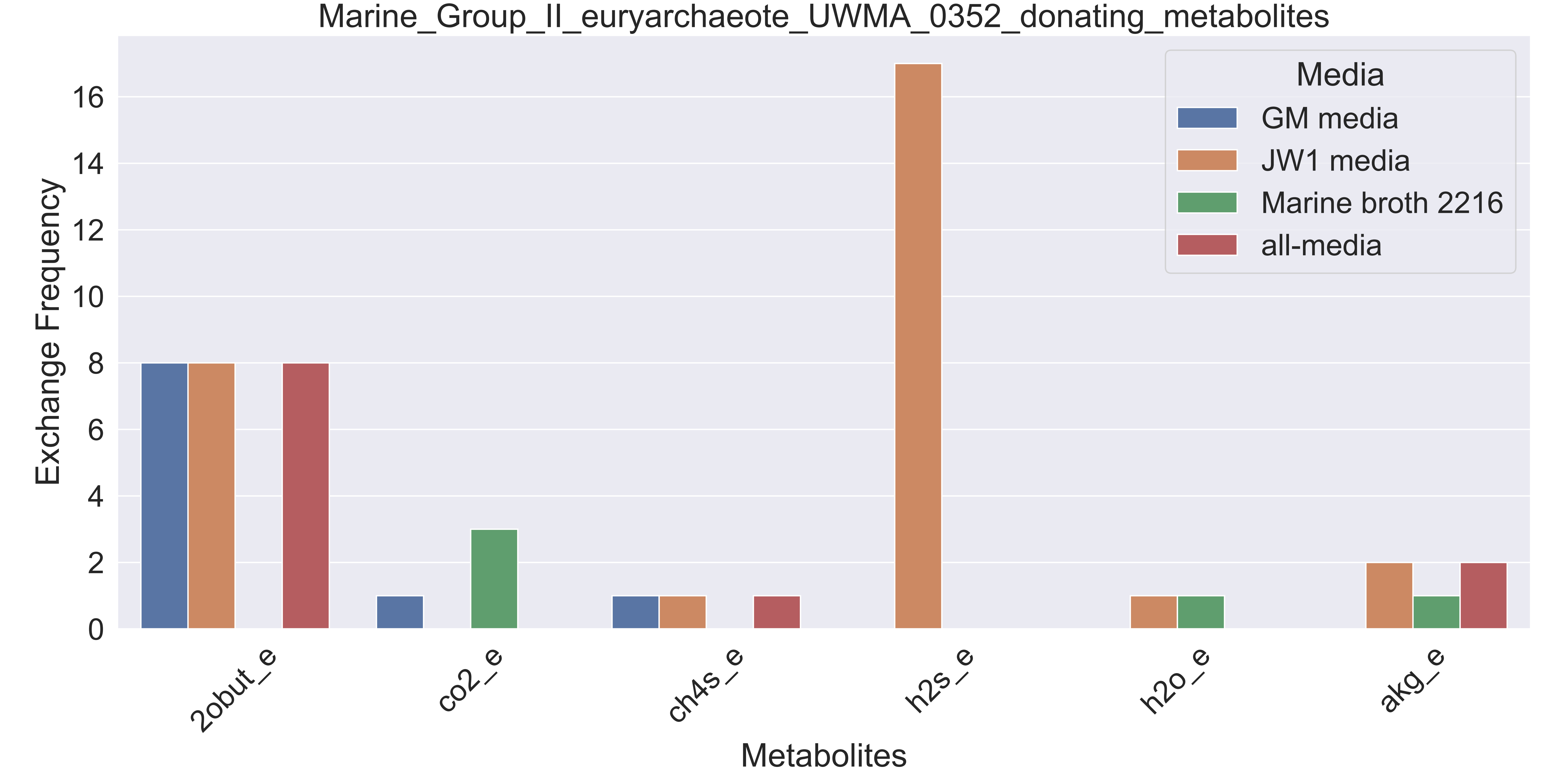

### Supplementary Figure S165 Marine Group II euryarchaeote UWMA 0357 donating metabolites.pdf

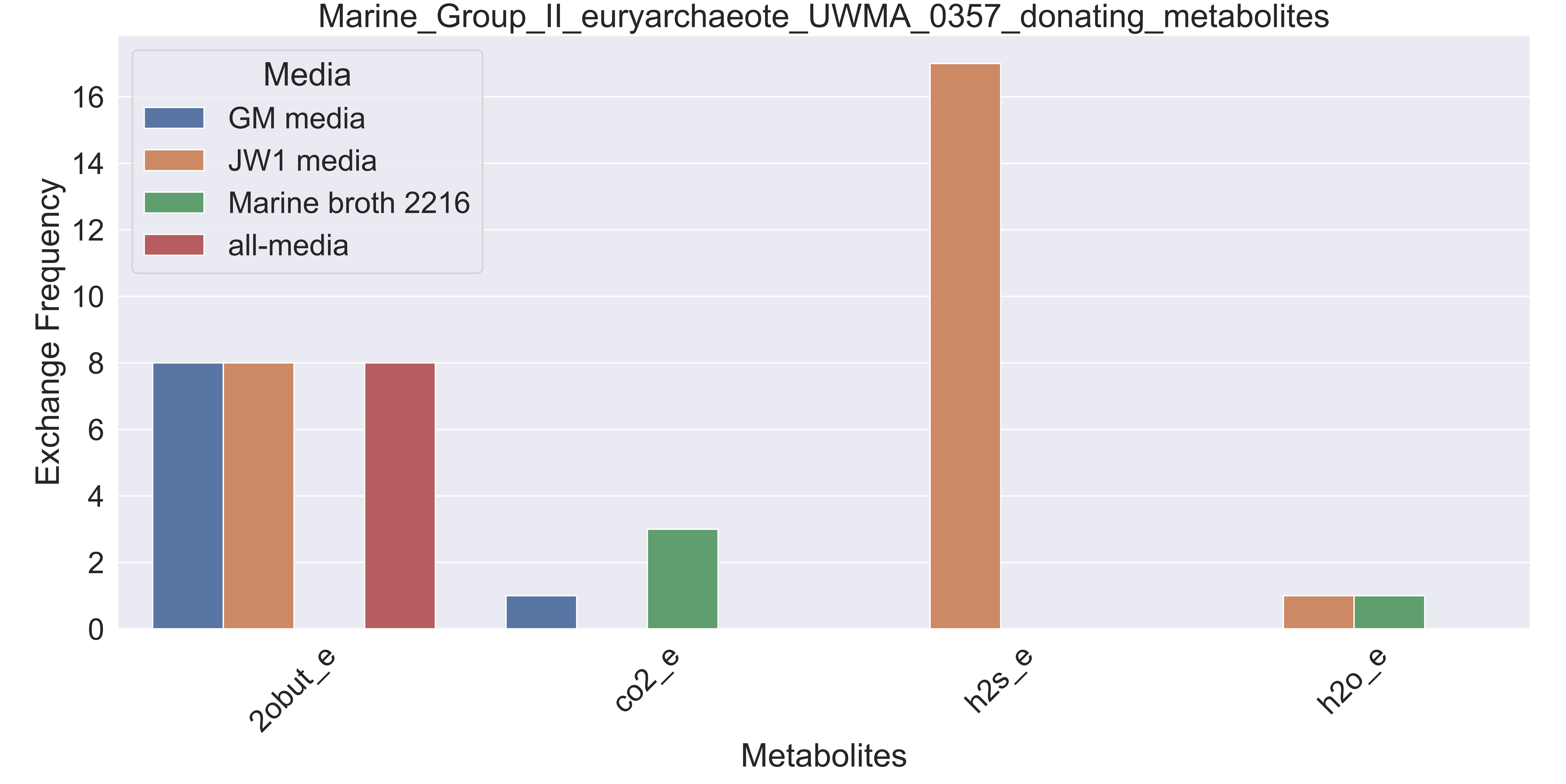

### Supplementary Figure S166 Methylococcaceae bacterium UWMA 0325 donating metabolites.pdf

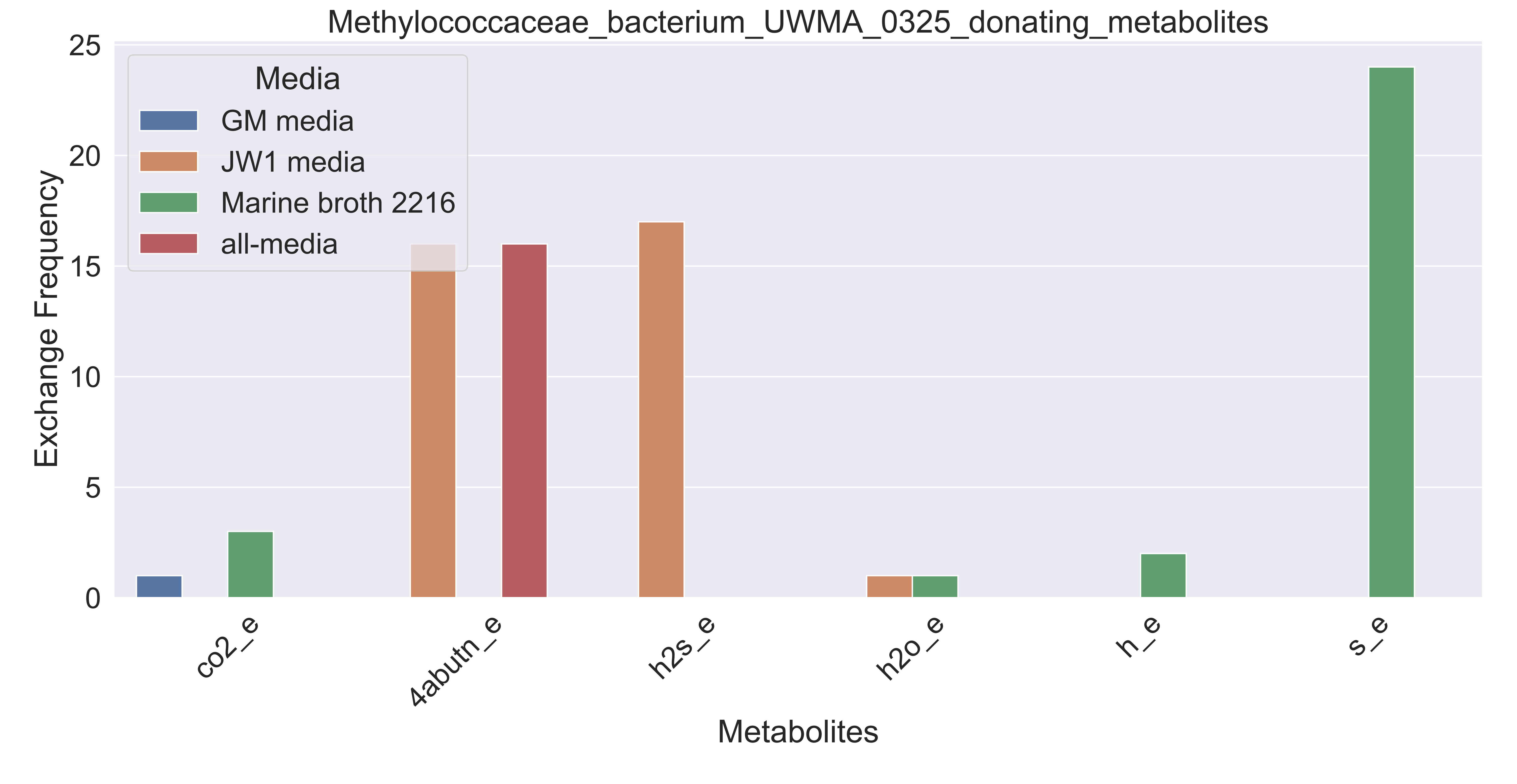

### Supplementary Figure S167 Methylococcaceae bacterium UWMA 0327 donating metabolites.pdf

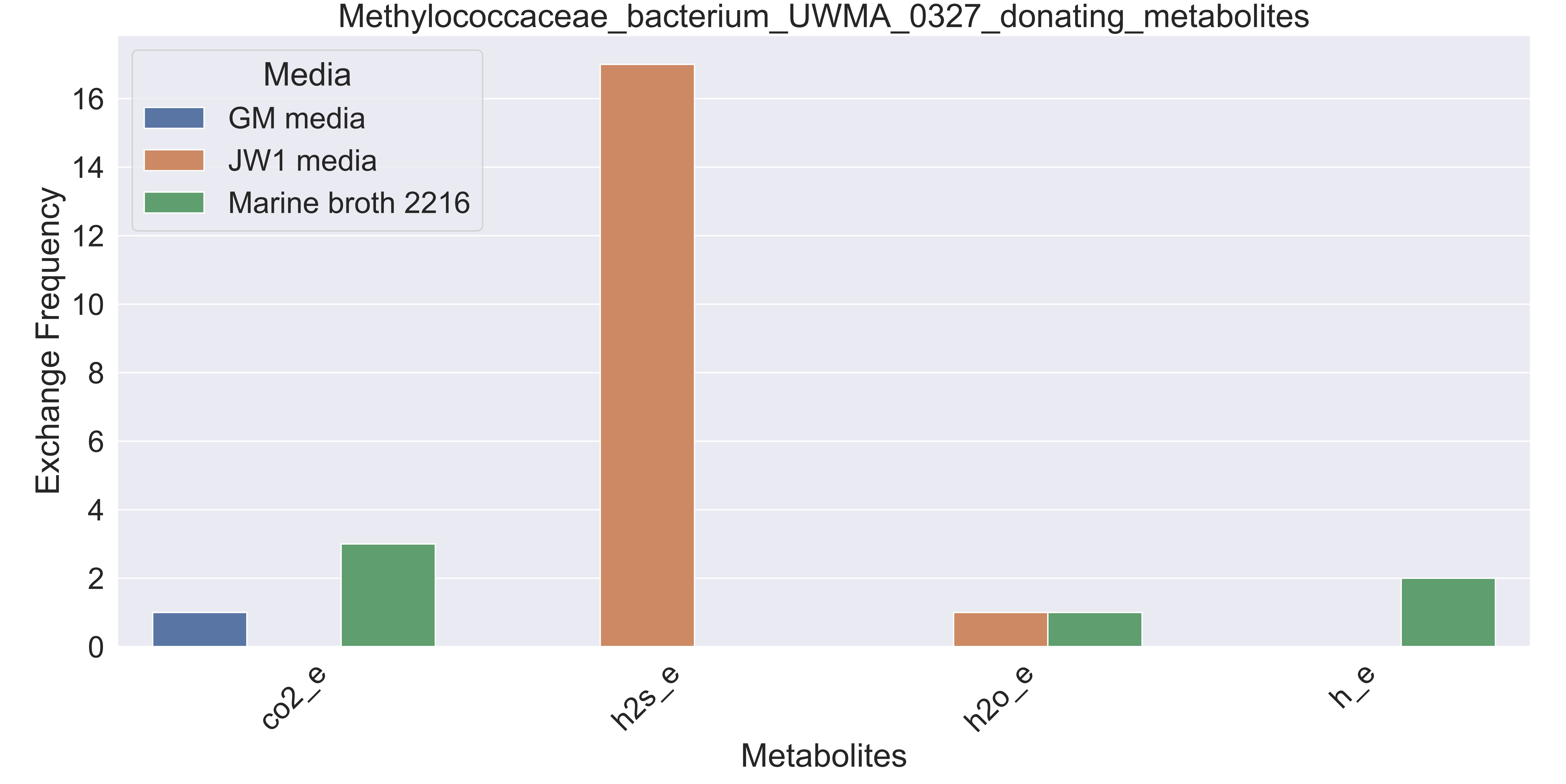

### Supplementary Figure S168 Methylococcaceae bacterium UWMA 0348 donating metabolites.pdf

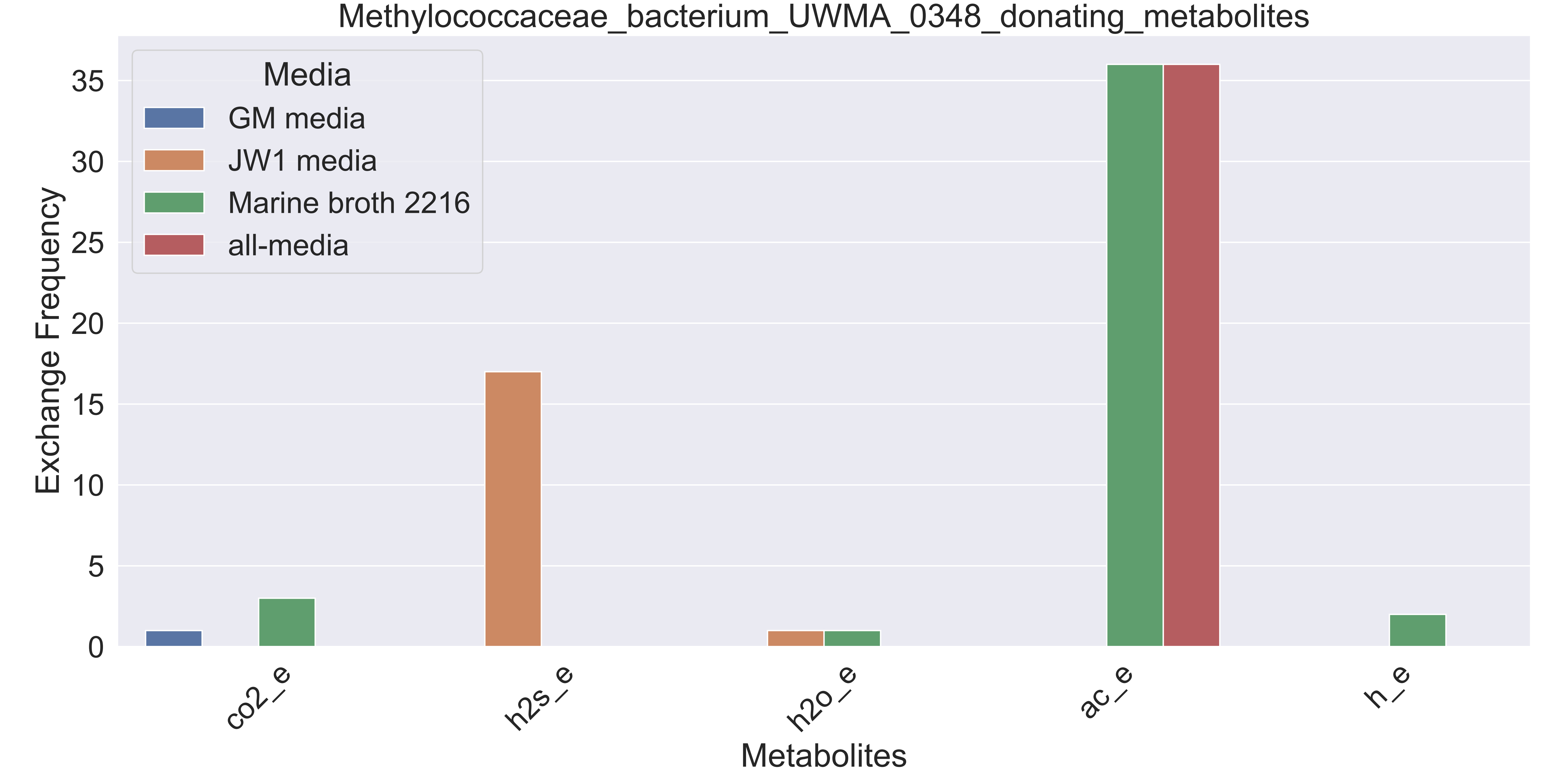

### Supplementary Figure S169 Methylophilaceae bacterium UWMA 0326 donating metabolites.pdf

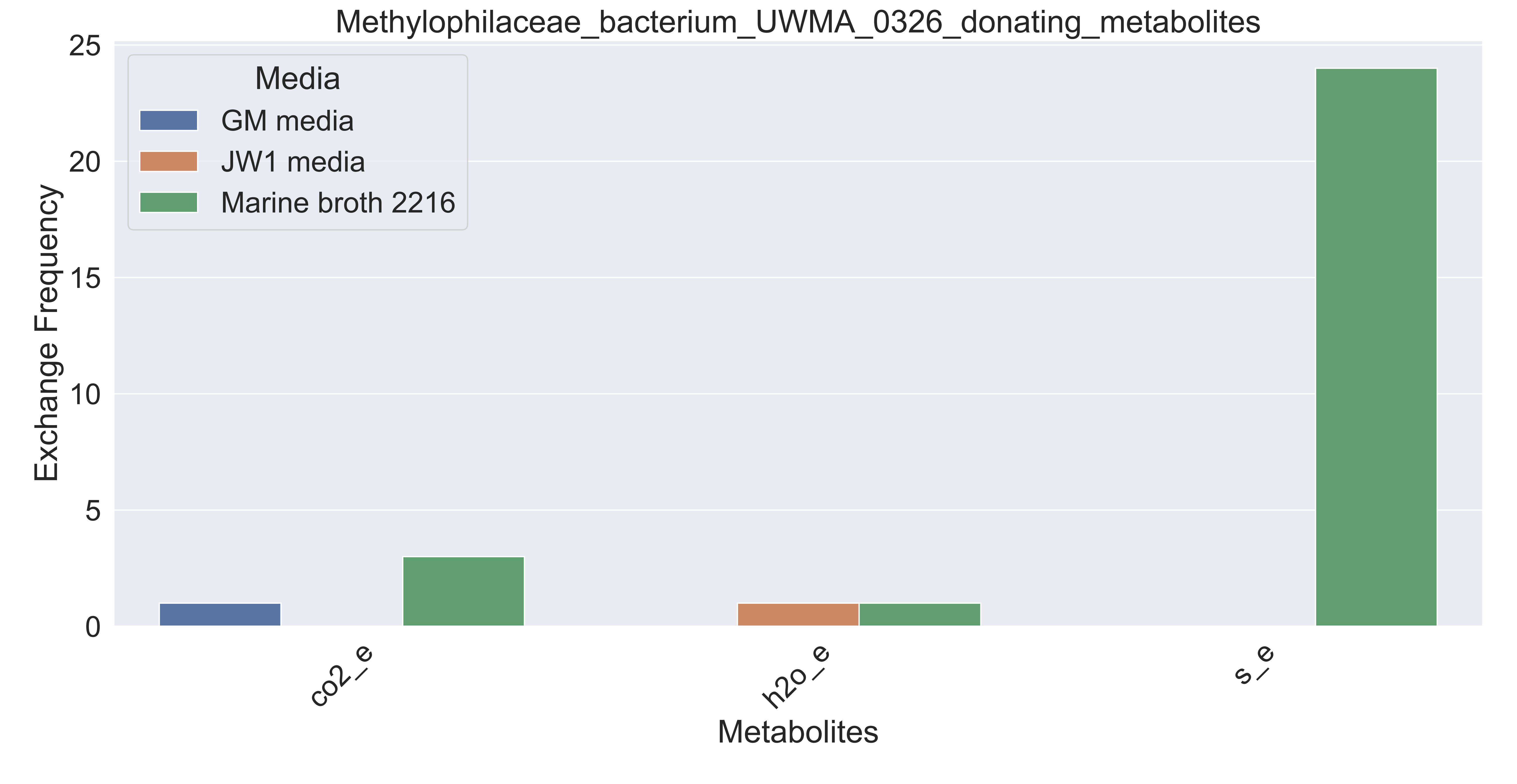

### Supplementary Figure S170 Methyloprofundus sp UWMA 0312 donating metabolites.pdf

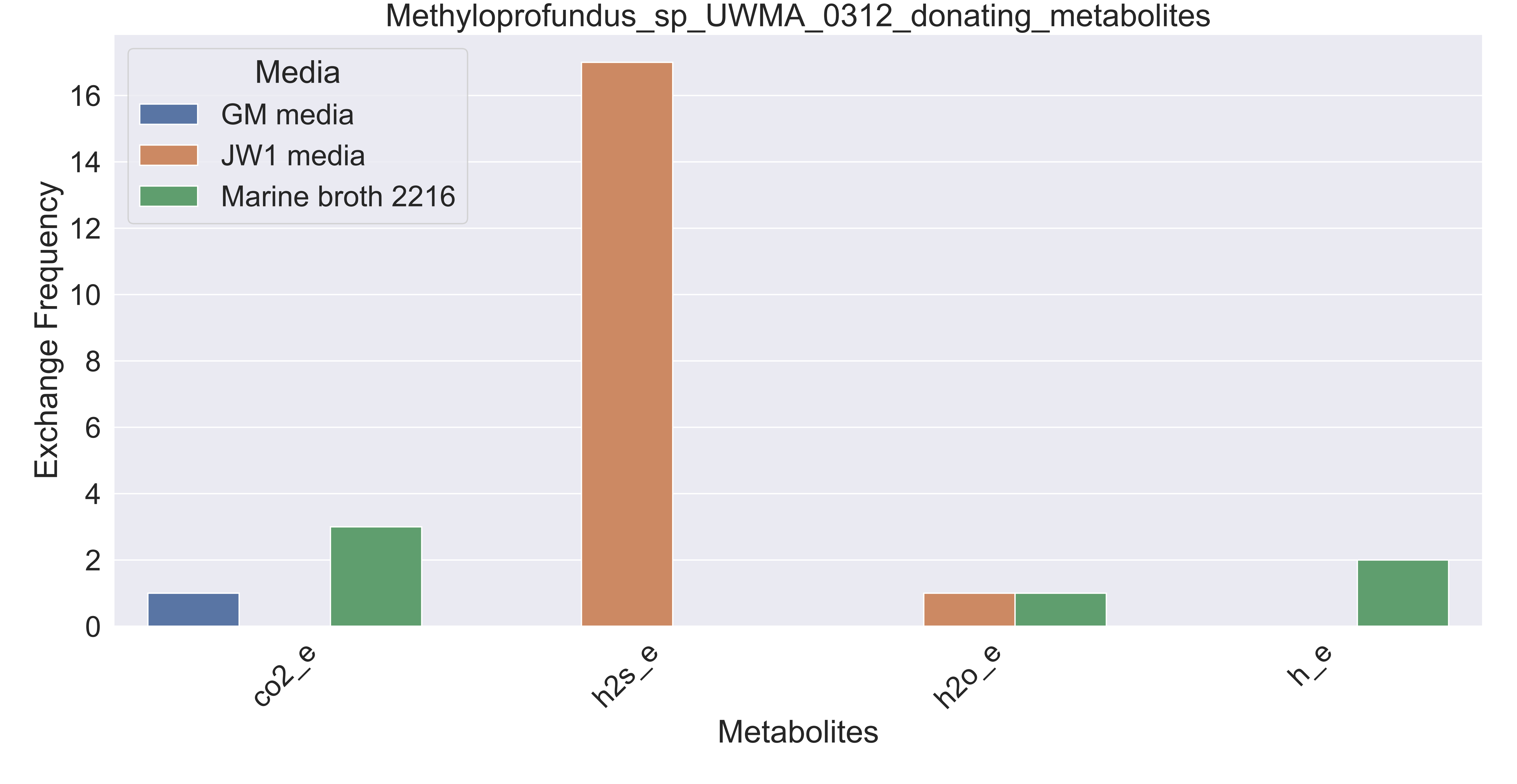

### Supplementary Figure S171 Micavibrio sp UWMA 0310 donating metabolites.pdf

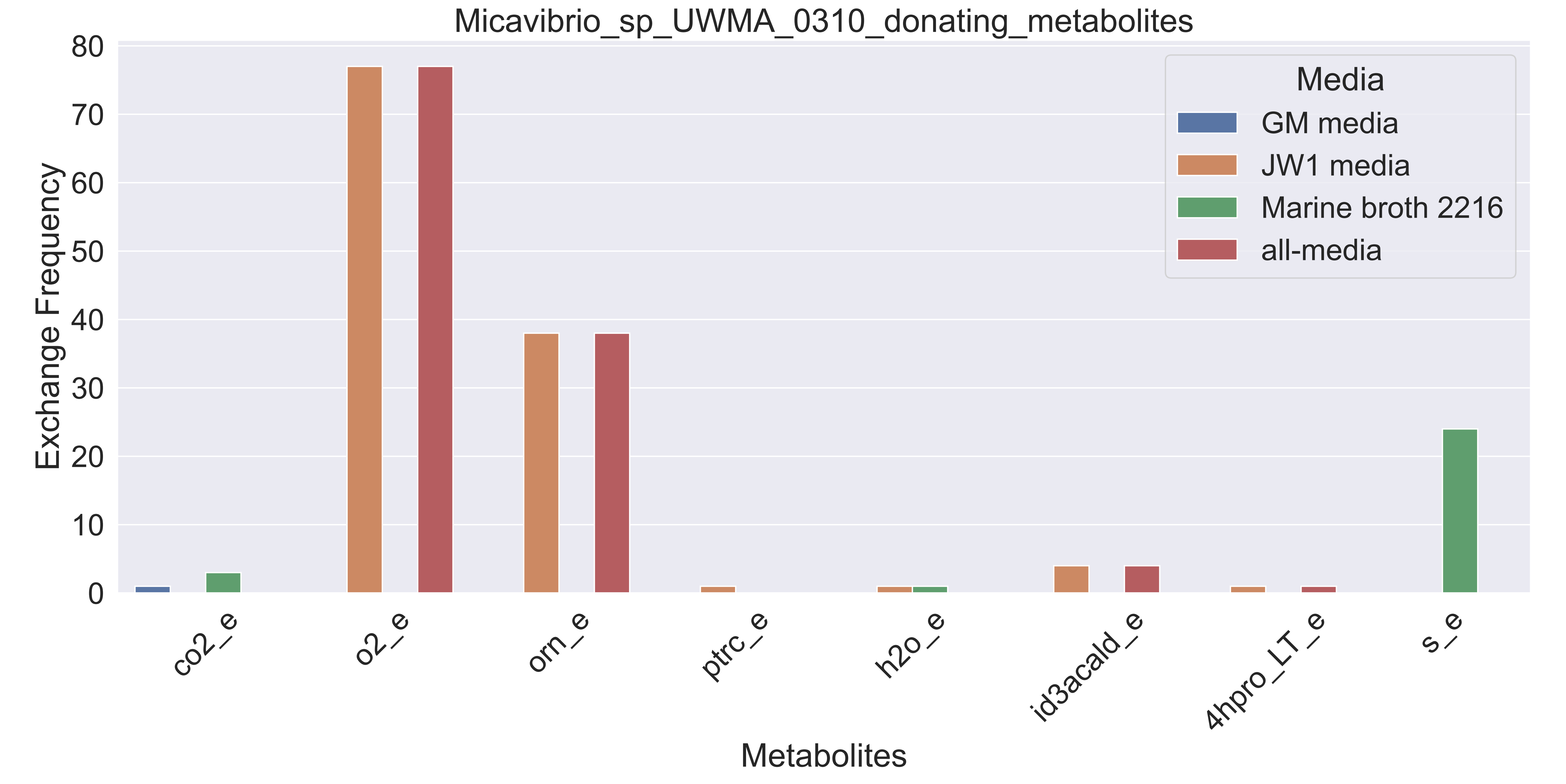

### Supplementary Figure S172 Microbacterium sp UWMA 0270 donating metabolites.pdf

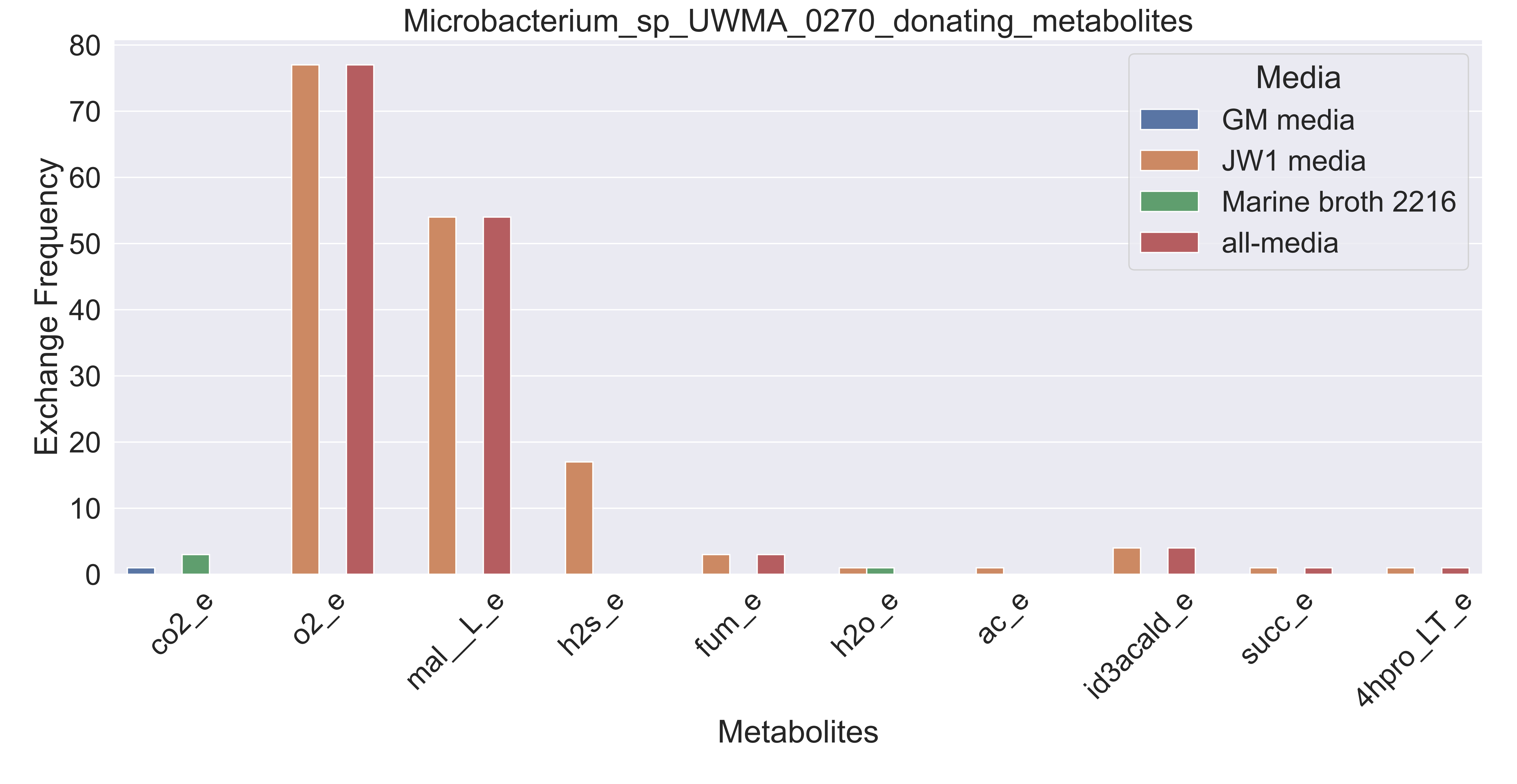

### Supplementary Figure S173 Myxococcales bacterium UWMA 0315 donating metabolites.pdf

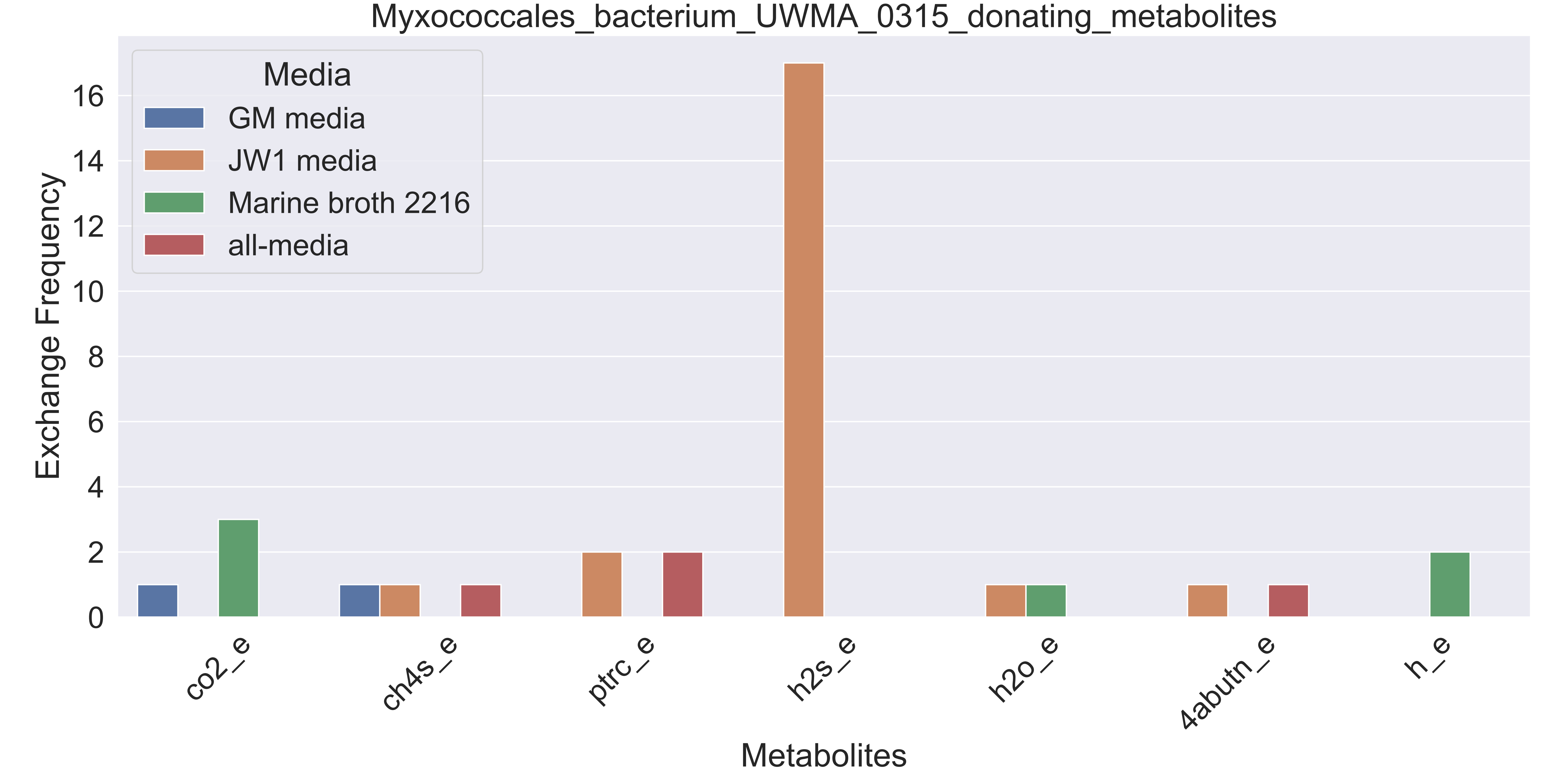

### Supplementary Figure S174 Myxococcales bacterium UWMA 0324 donating metabolites.pdf

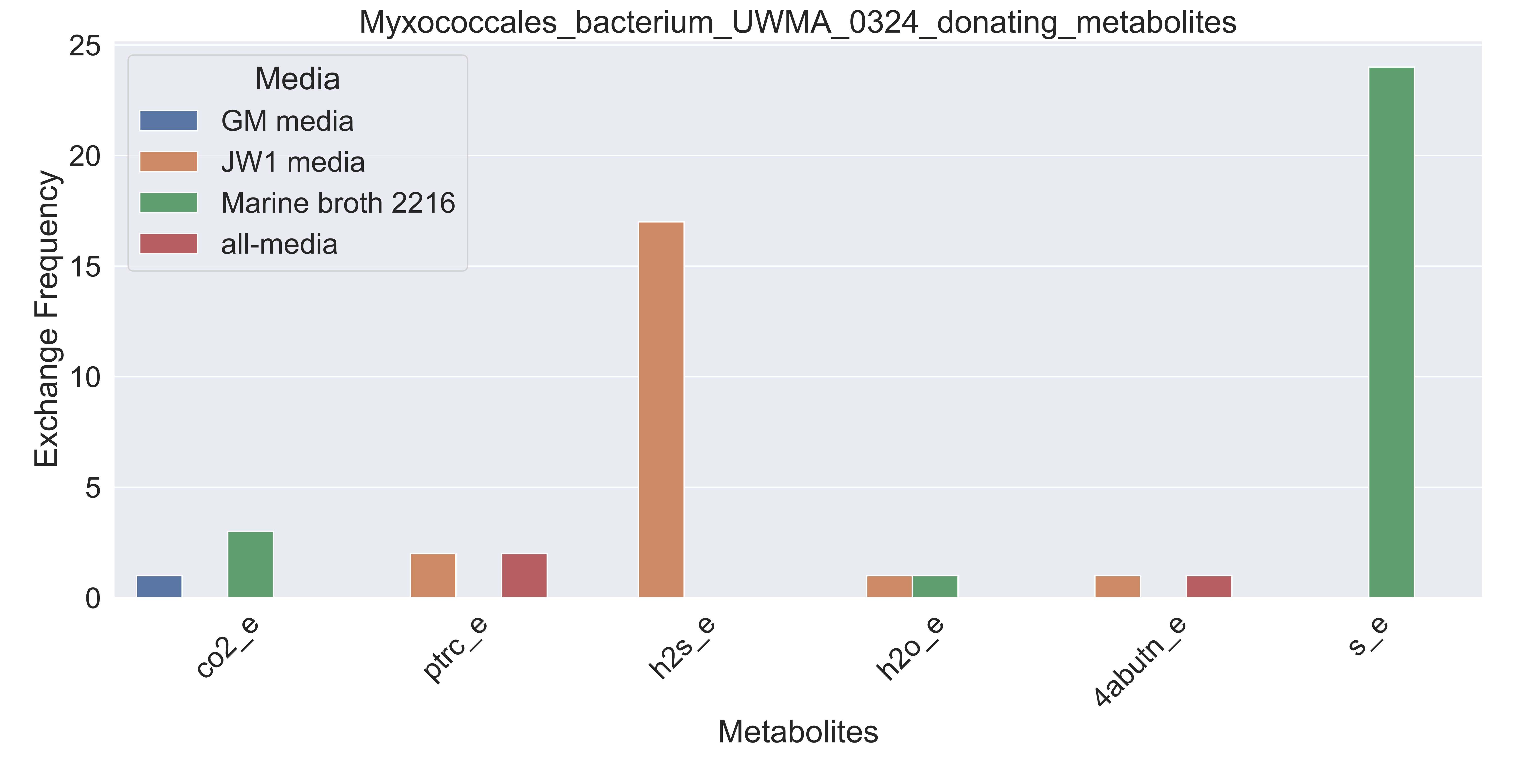

### Supplementary Figure S175 Myxococcales bacterium UWMA 0330 donating metabolites.pdf

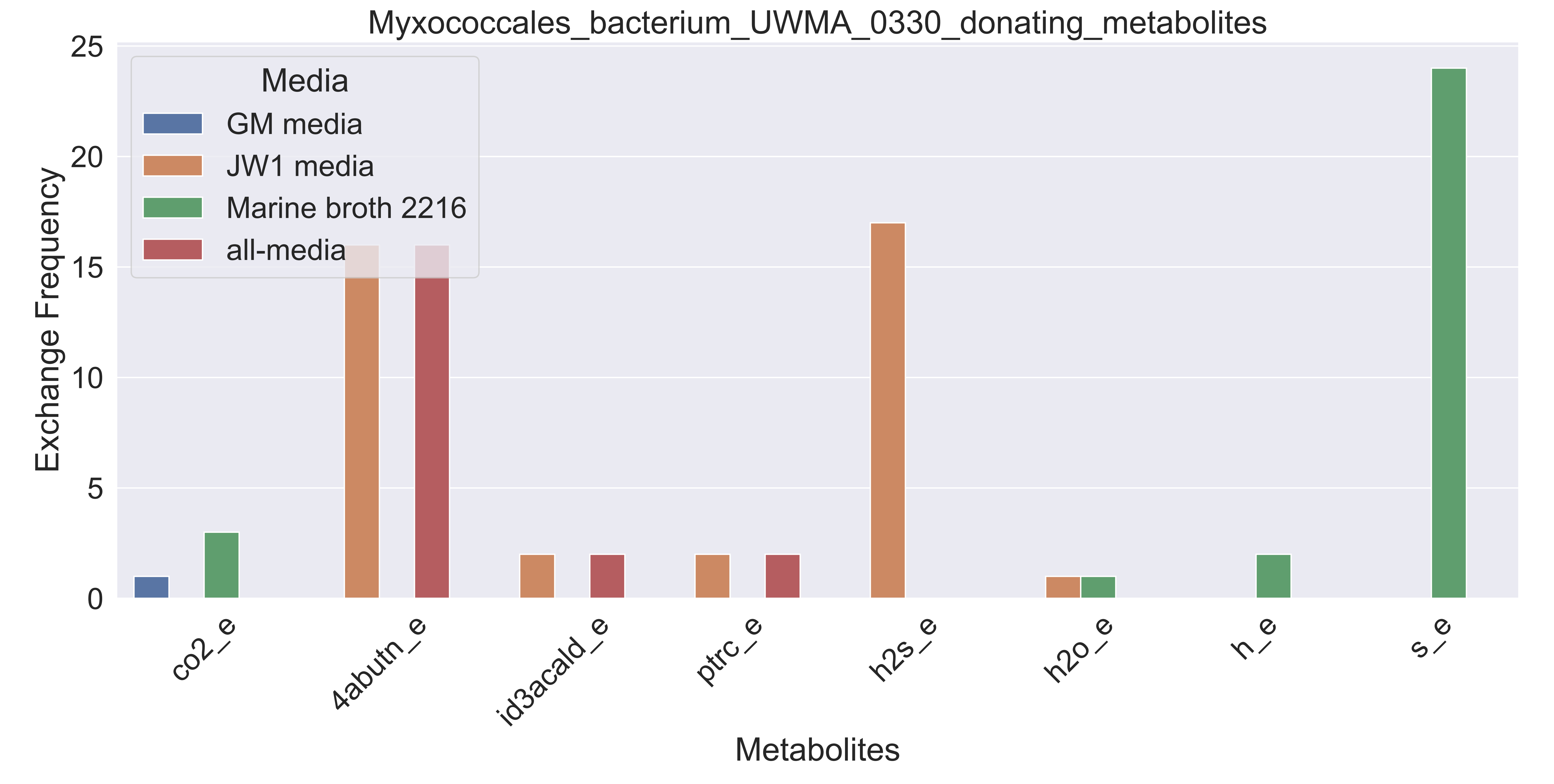

### Supplementary Figure S176 Myxococcales bacterium UWMA 0336 donating metabolites.pdf

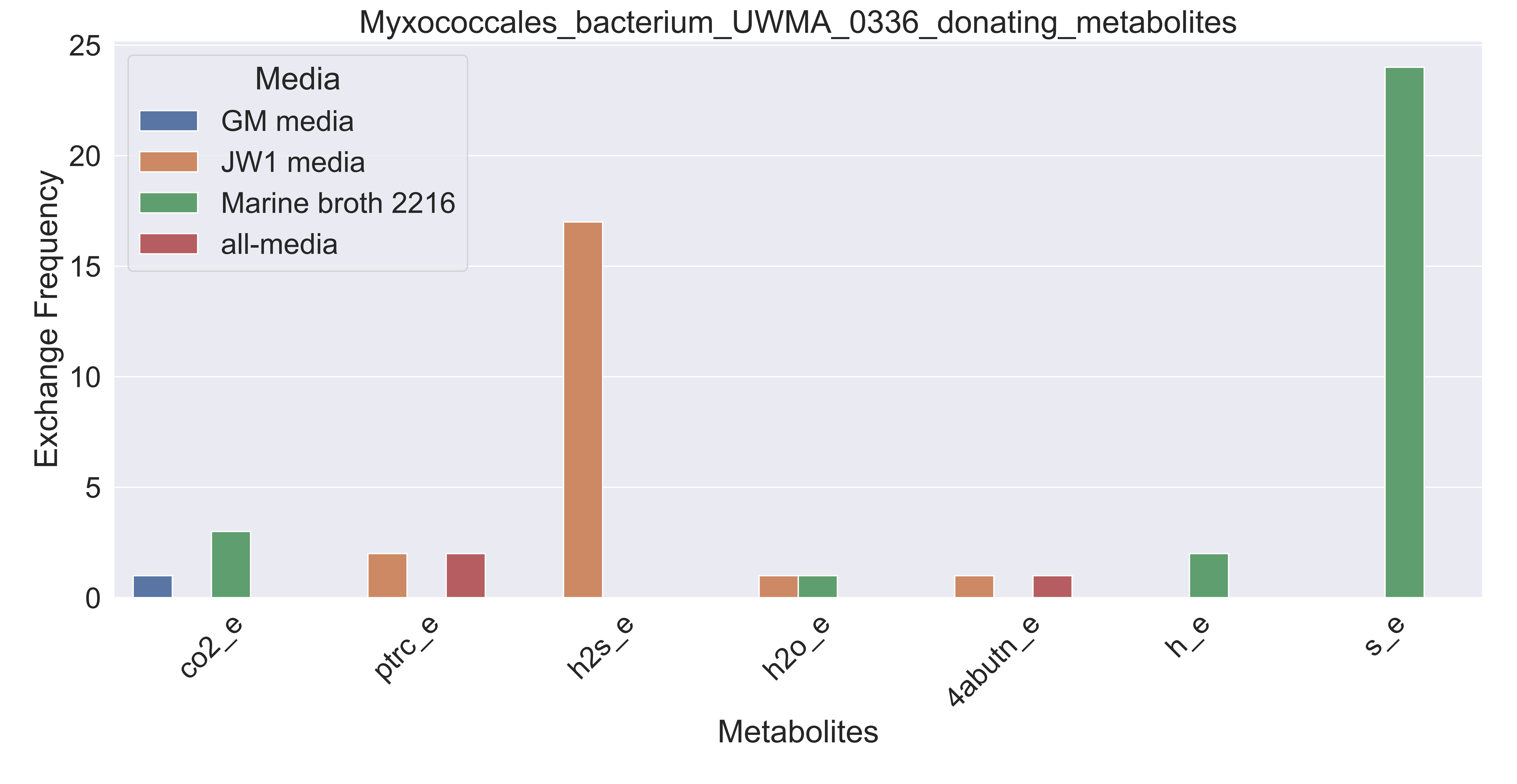

### Supplementary Figure S177 Nitrosopumilus sp UWMA 0263 donating metabolites.pdf

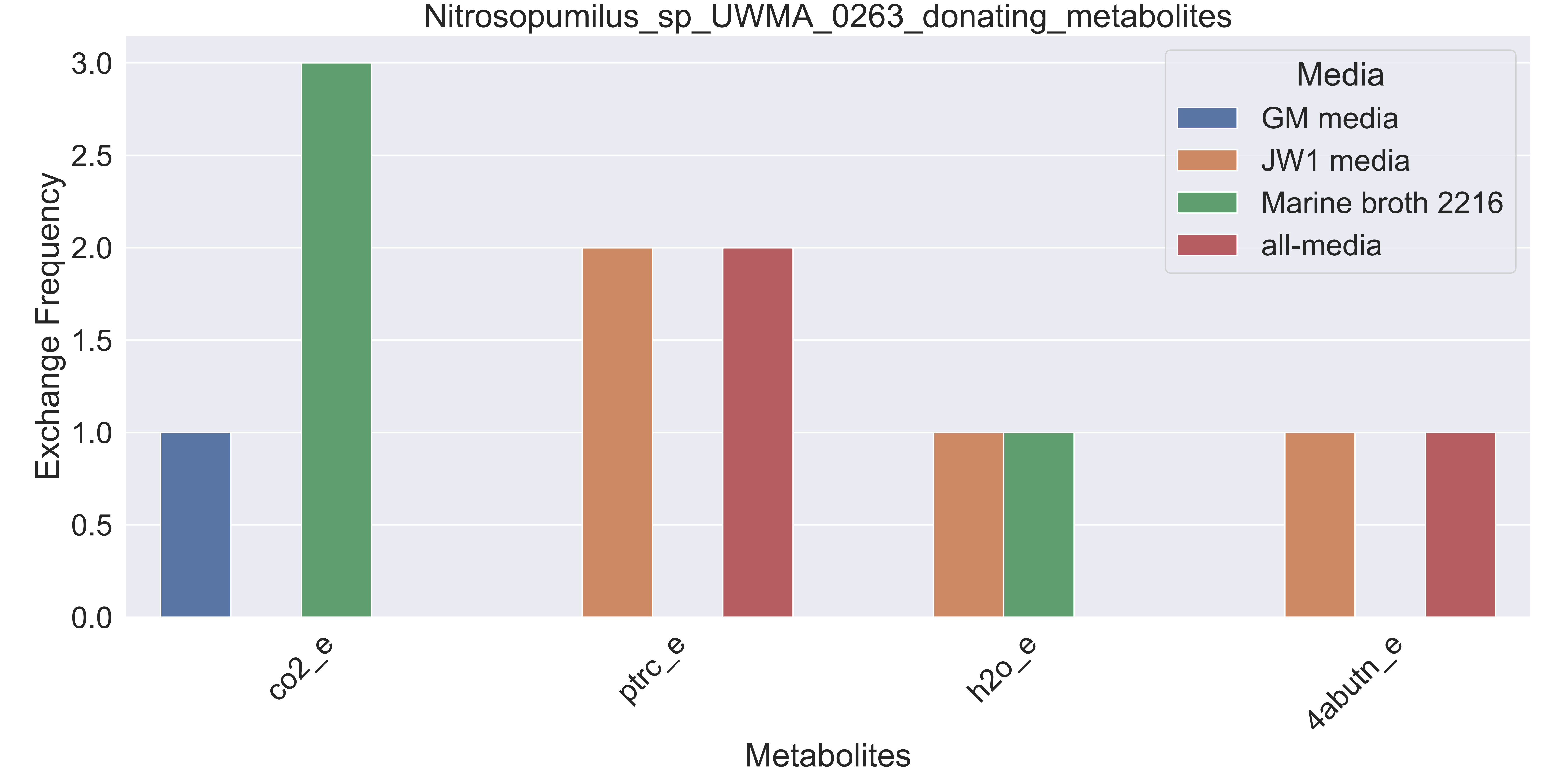

### Supplementary Figure S178 Nitrospinaceae bacterium UWMA 0262 donating metabolites.pdf

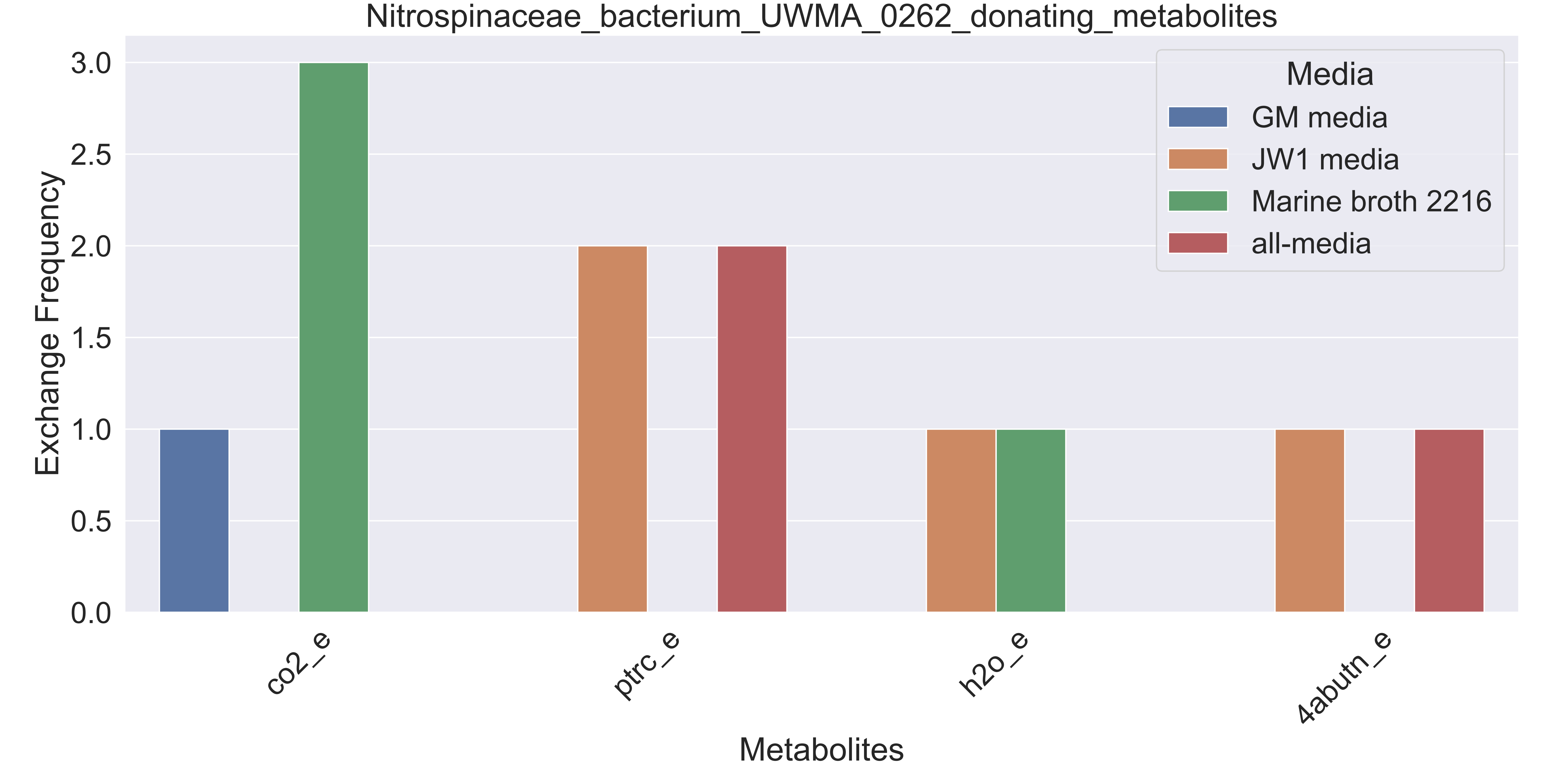

### Supplementary Figure S179 Nitrospinaceae bacterium UWMA 0271 donating metabolites.pdf

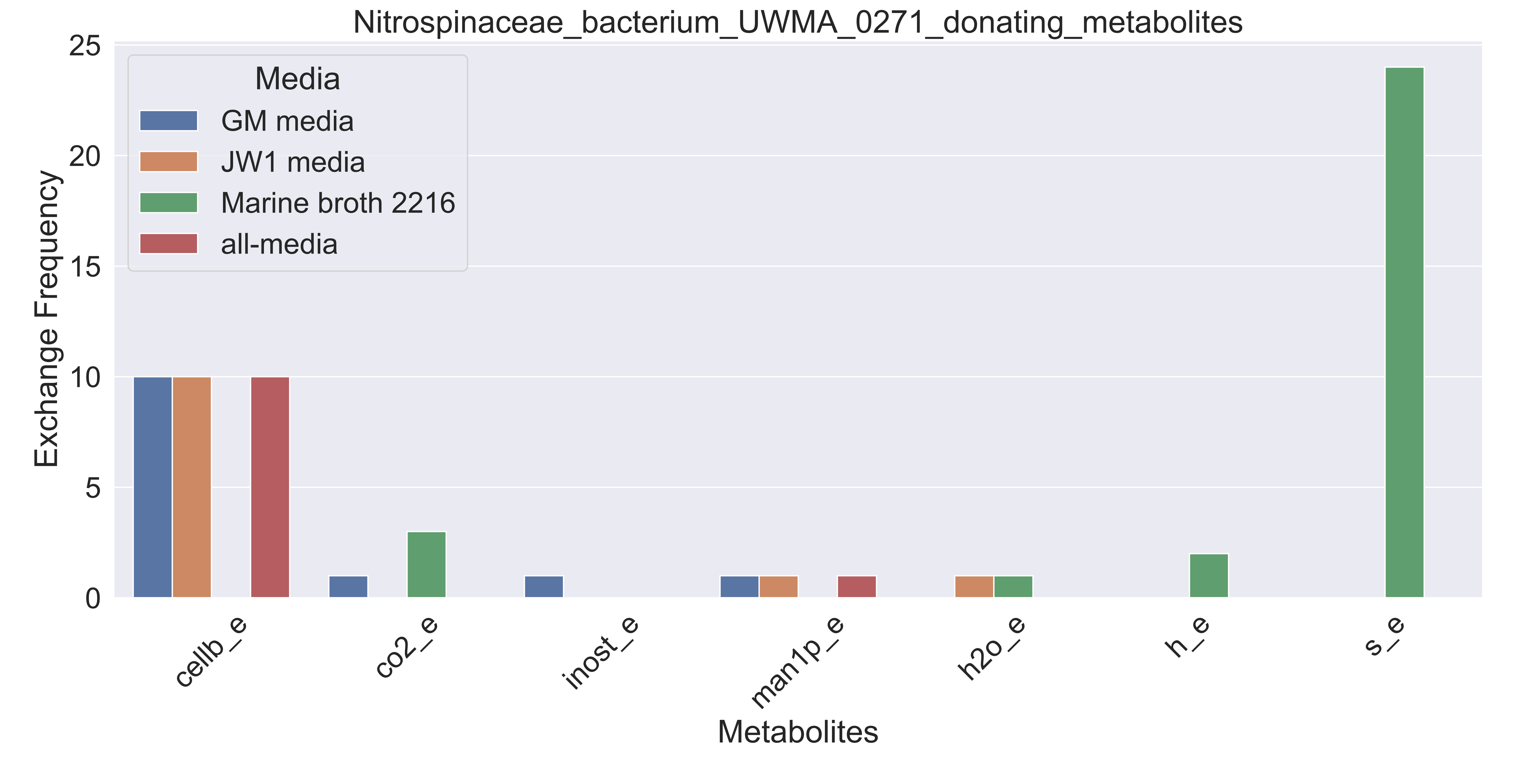

### Supplementary Figure S180 Nitrospinaceae bacterium UWMA 0356 donating metabolites.pdf

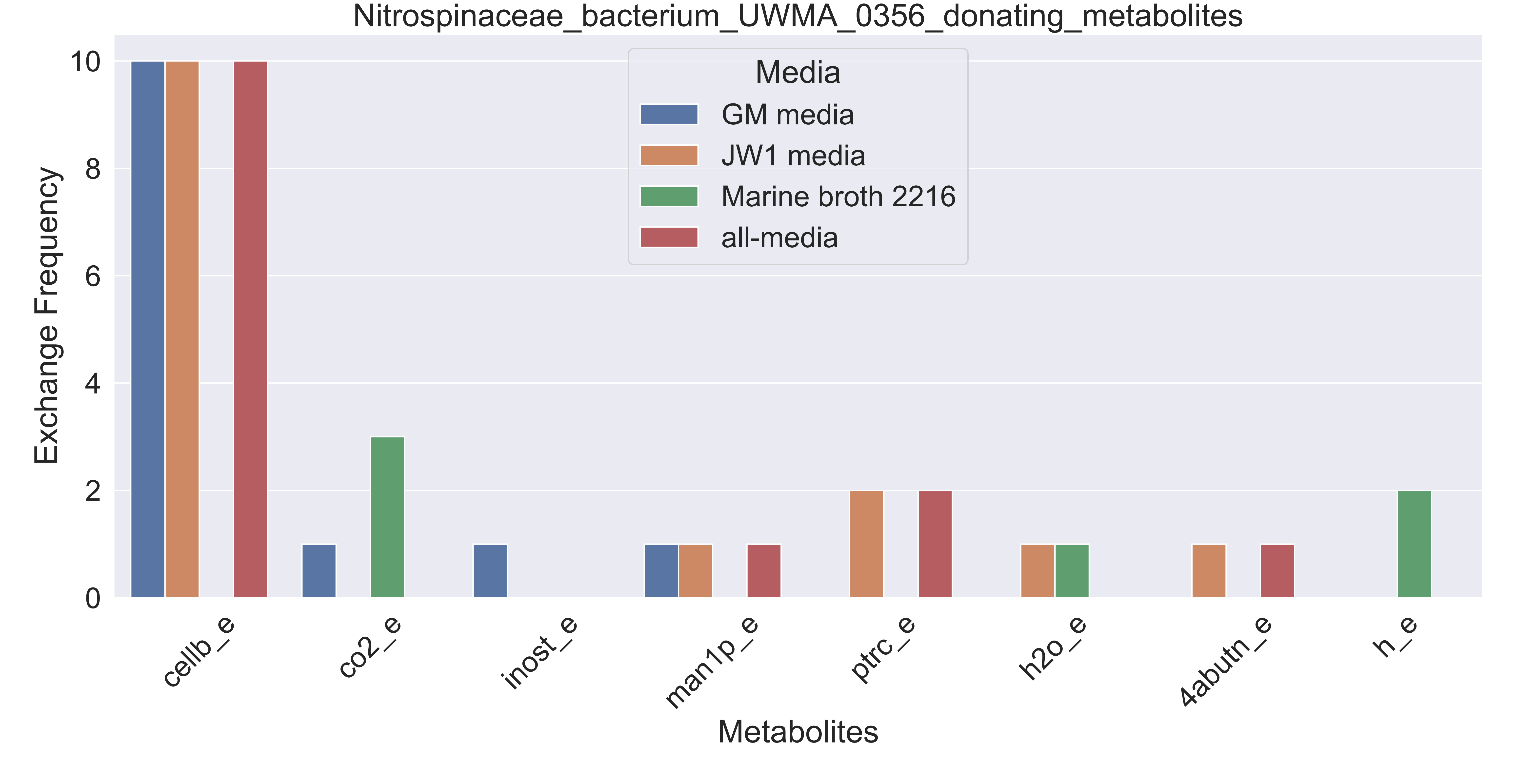

### Supplementary Figure S181 Nitrospira sp UWMA 0268 donating metabolites.pdf

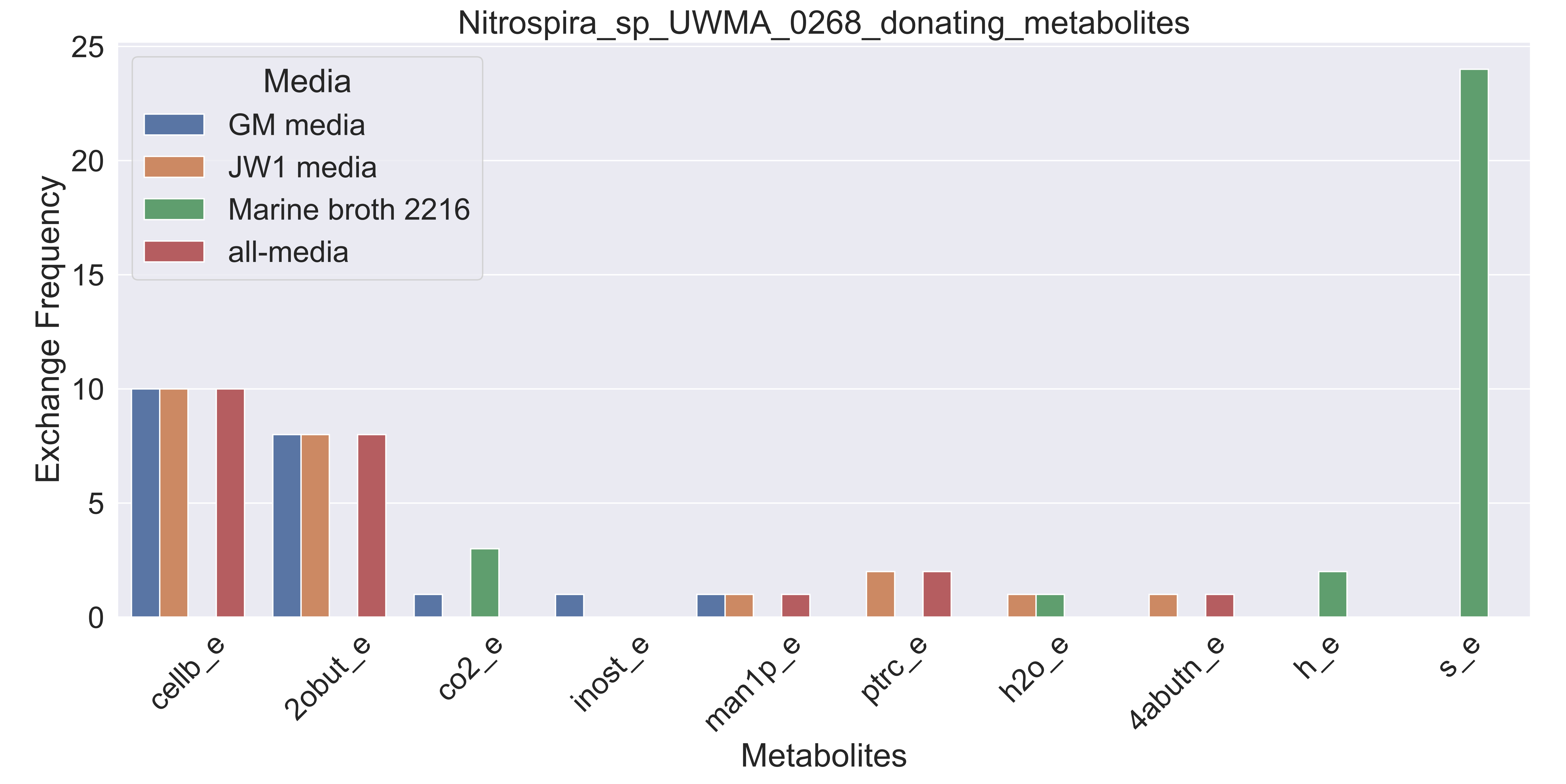

### Supplementary Figure S182 Oceanospirillaceae bacterium UWMA 0280 donating metabolites.pdf

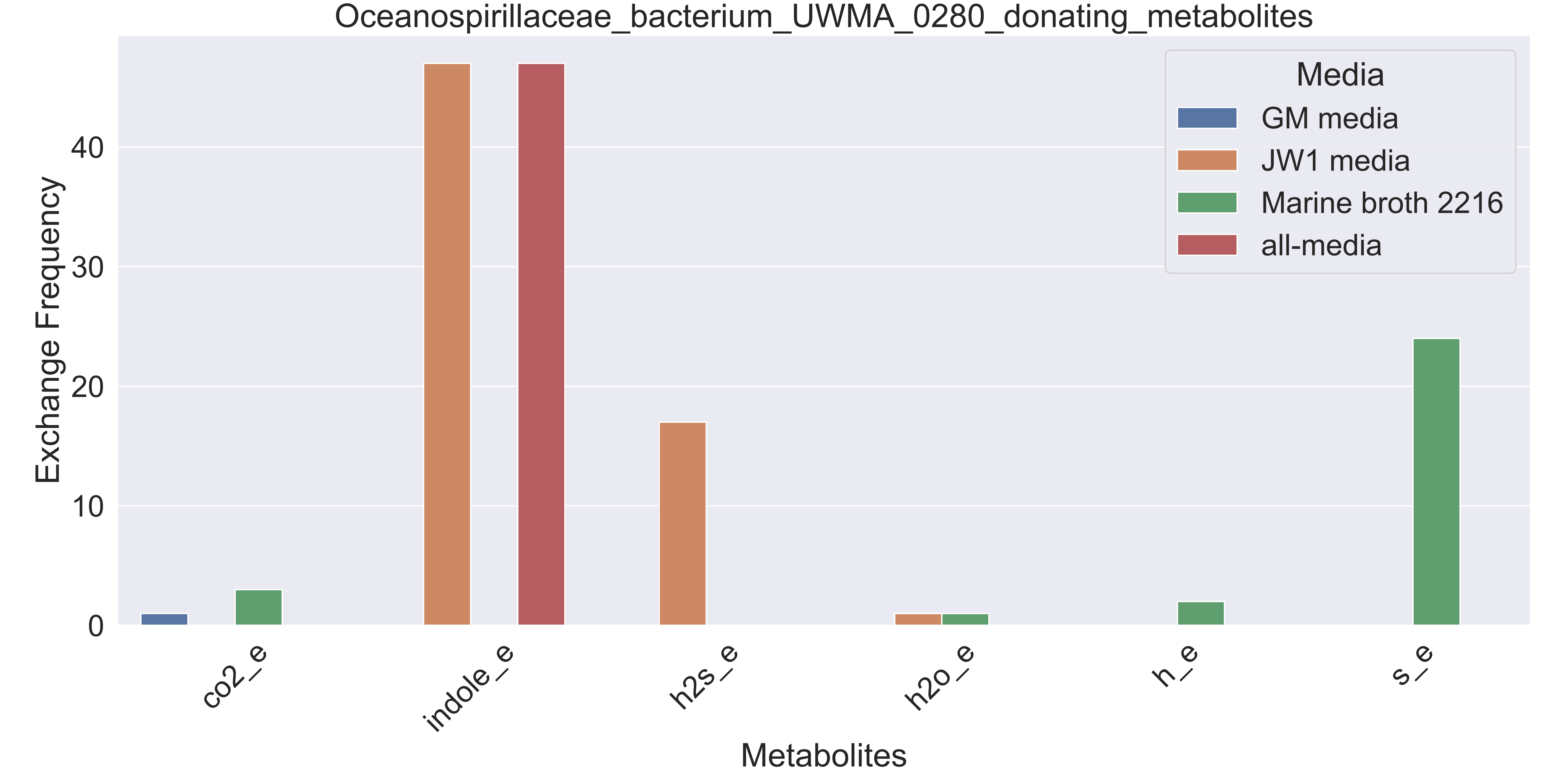

### Supplementary Figure S183 Planctomycetaceae bacterium UWMA 0346 donating metabolites.pdf

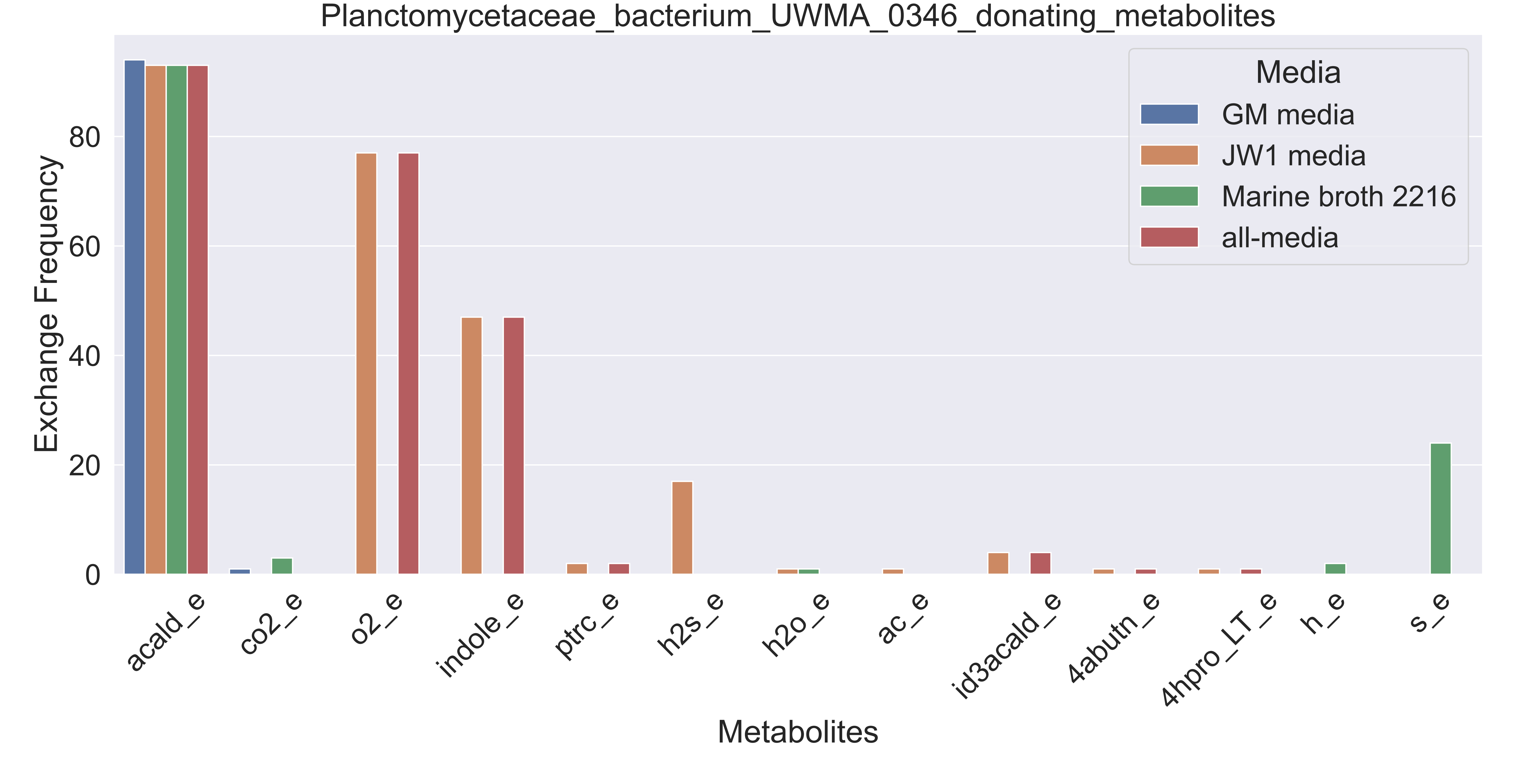

### Supplementary Figure S184 Planctomycetes bacterium UWMA 0265 donating metabolites.pdf

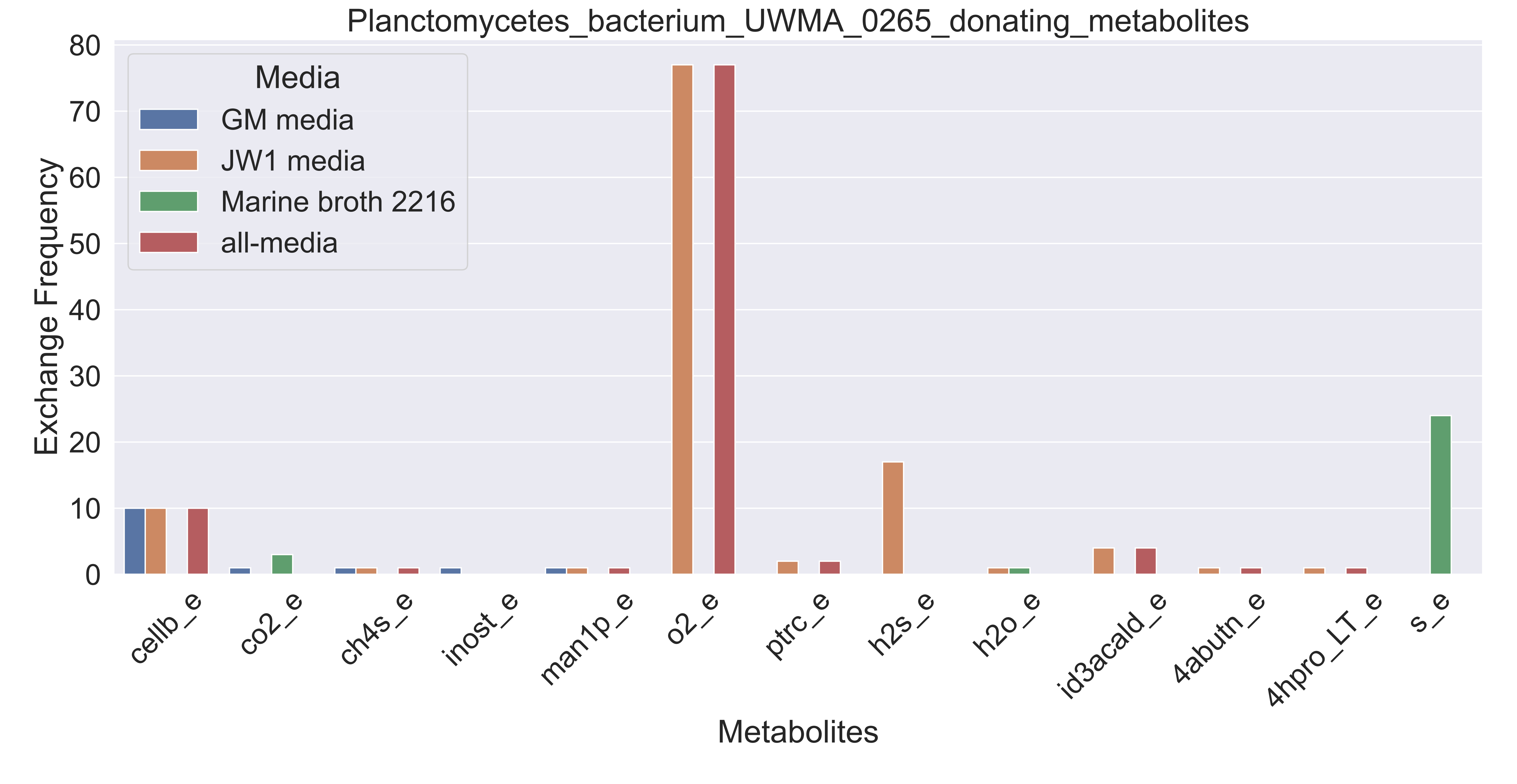

### Supplementary Figure S185 Planctomycetes bacterium UWMA 0276 donating metabolites.pdf

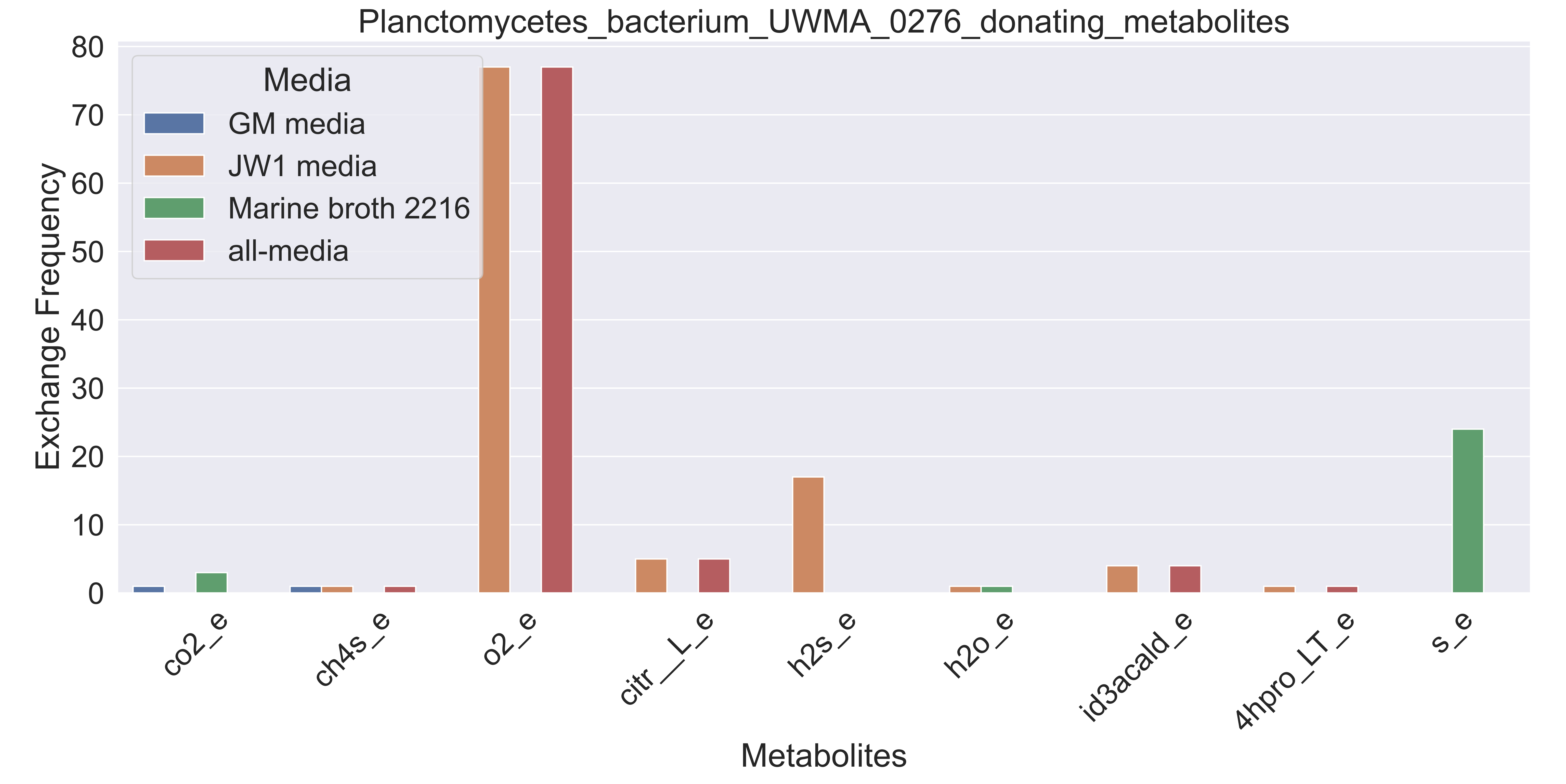

### Supplementary Figure S186 Planctomycetes bacterium UWMA 0294 donating metabolites.pdf

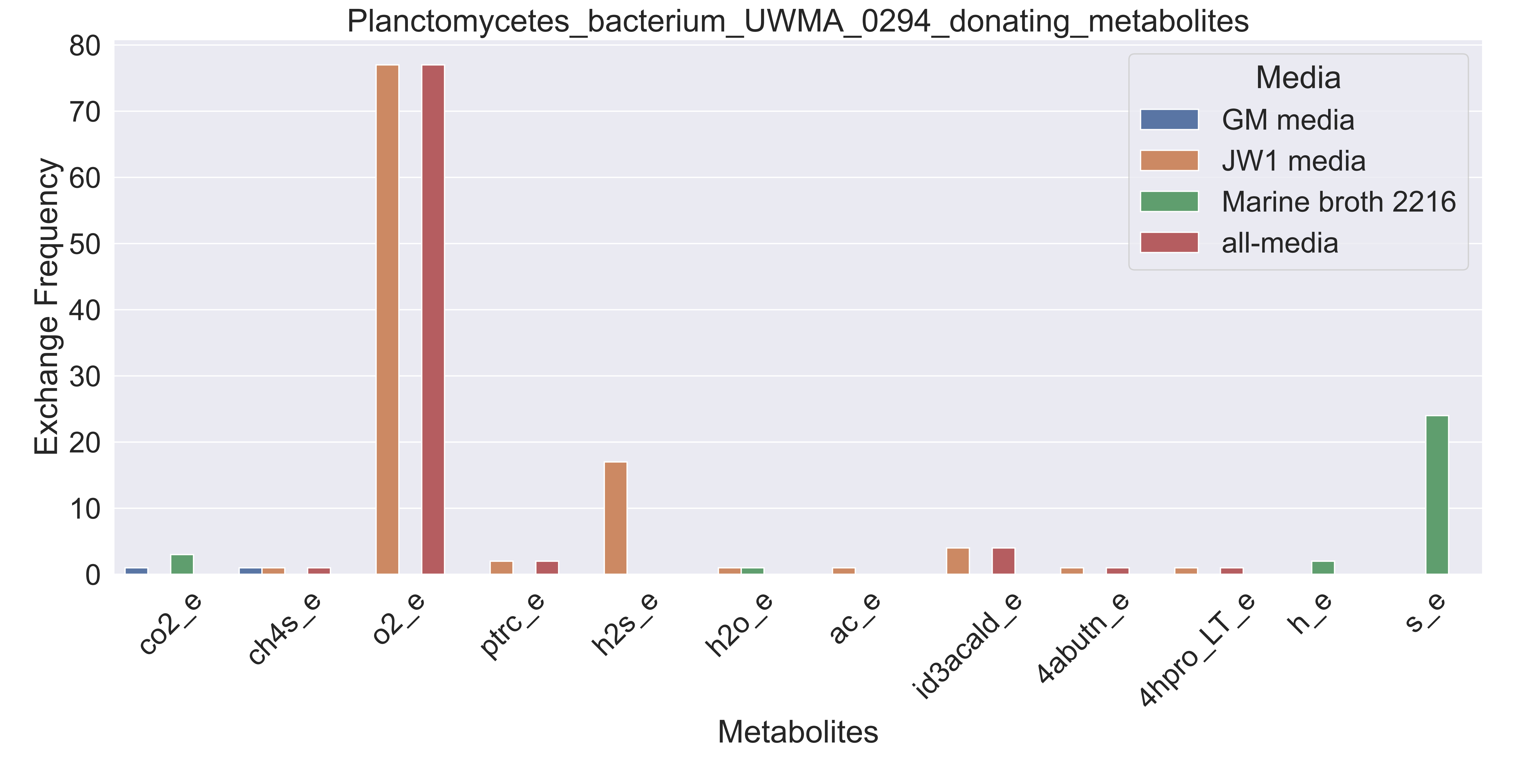

### Supplementary Figure S187 Planctomycetes bacterium UWMA 0329 donating metabolites.pdf

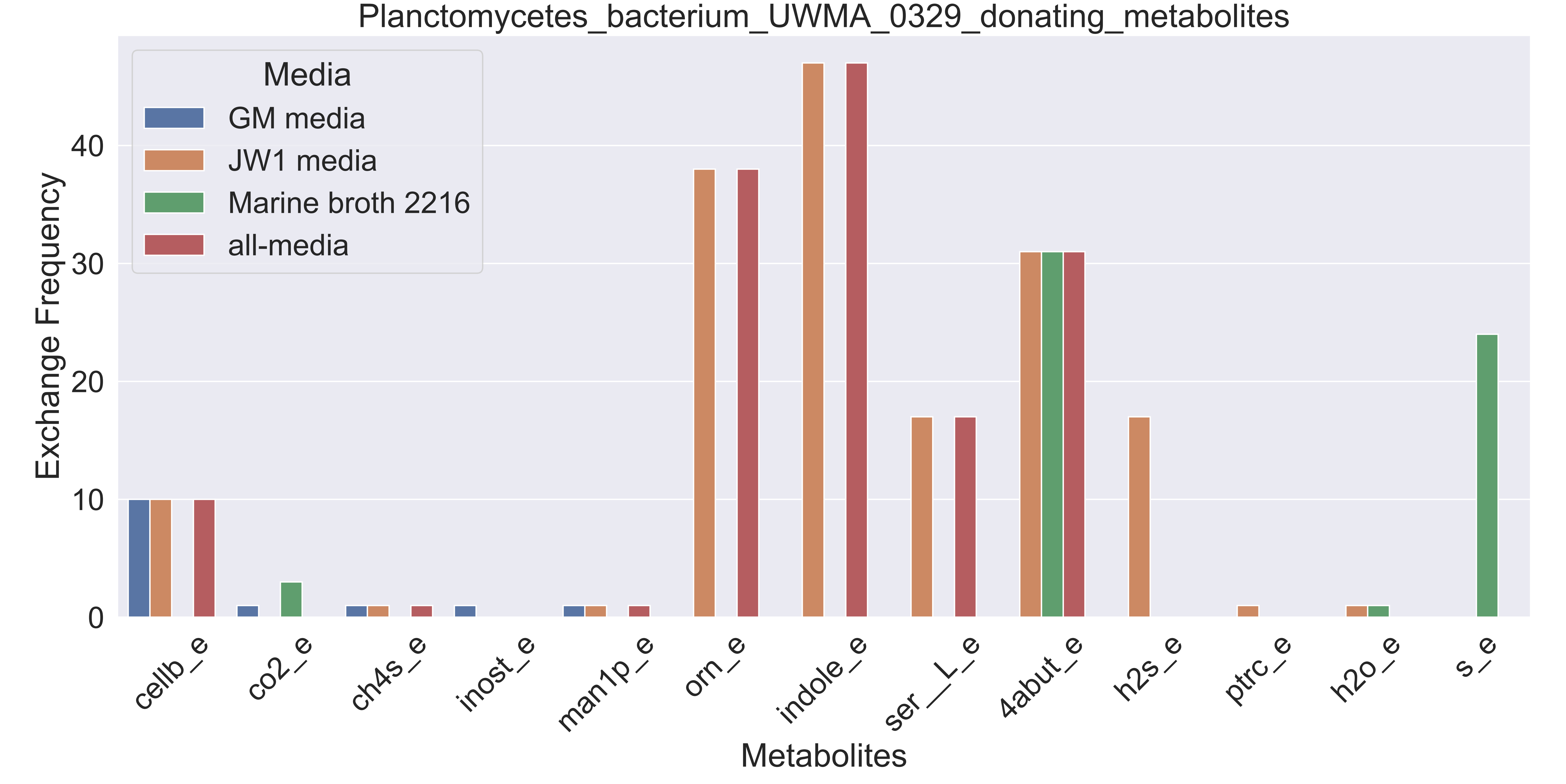

### Supplementary Figure S188 Planctomycetes bacterium UWMA 0345 donating metabolites.pdf

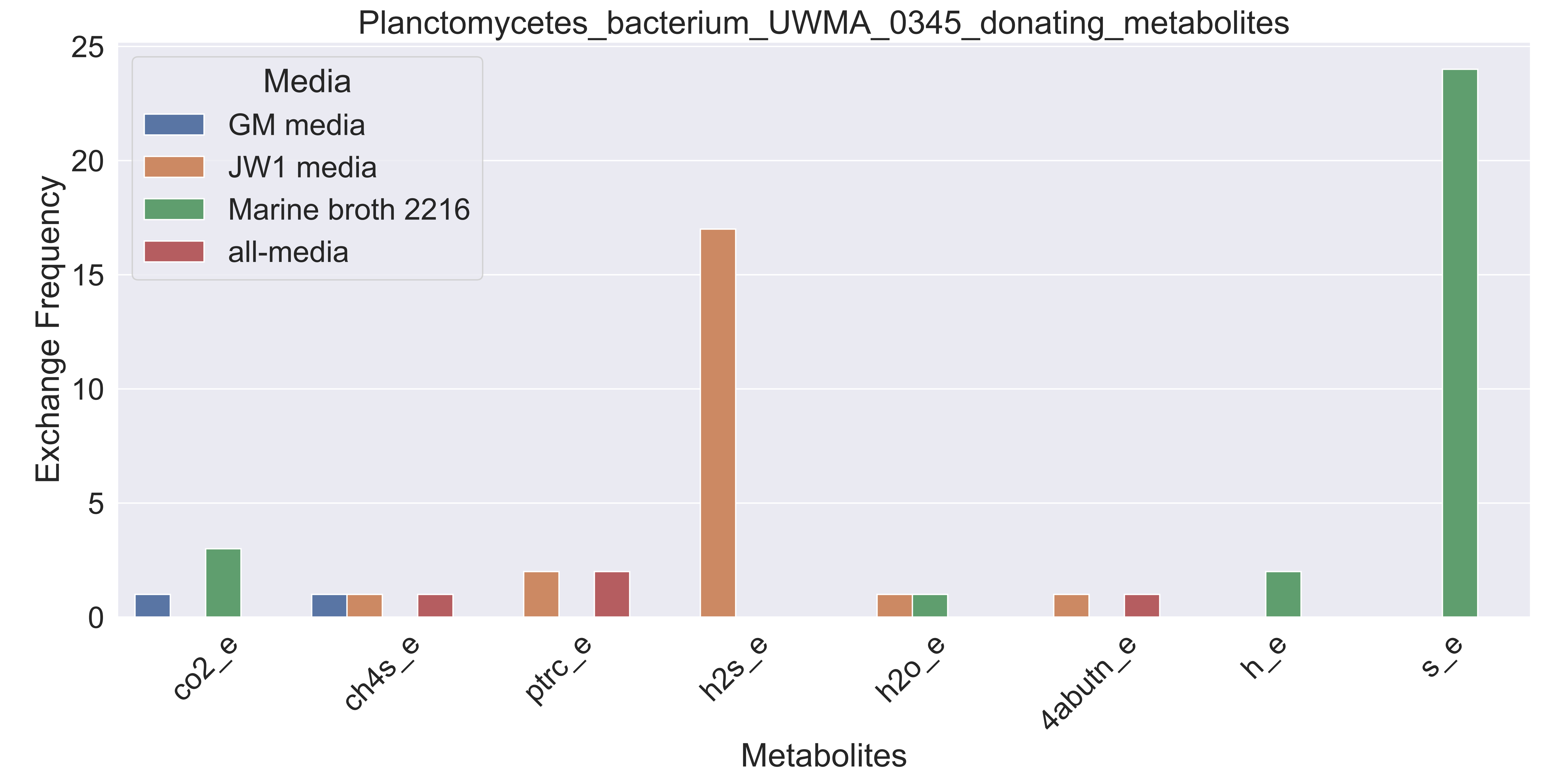

### Supplementary Figure S189 Porticoccaceae bacterium UWMA 0313 donating metabolites.pdf

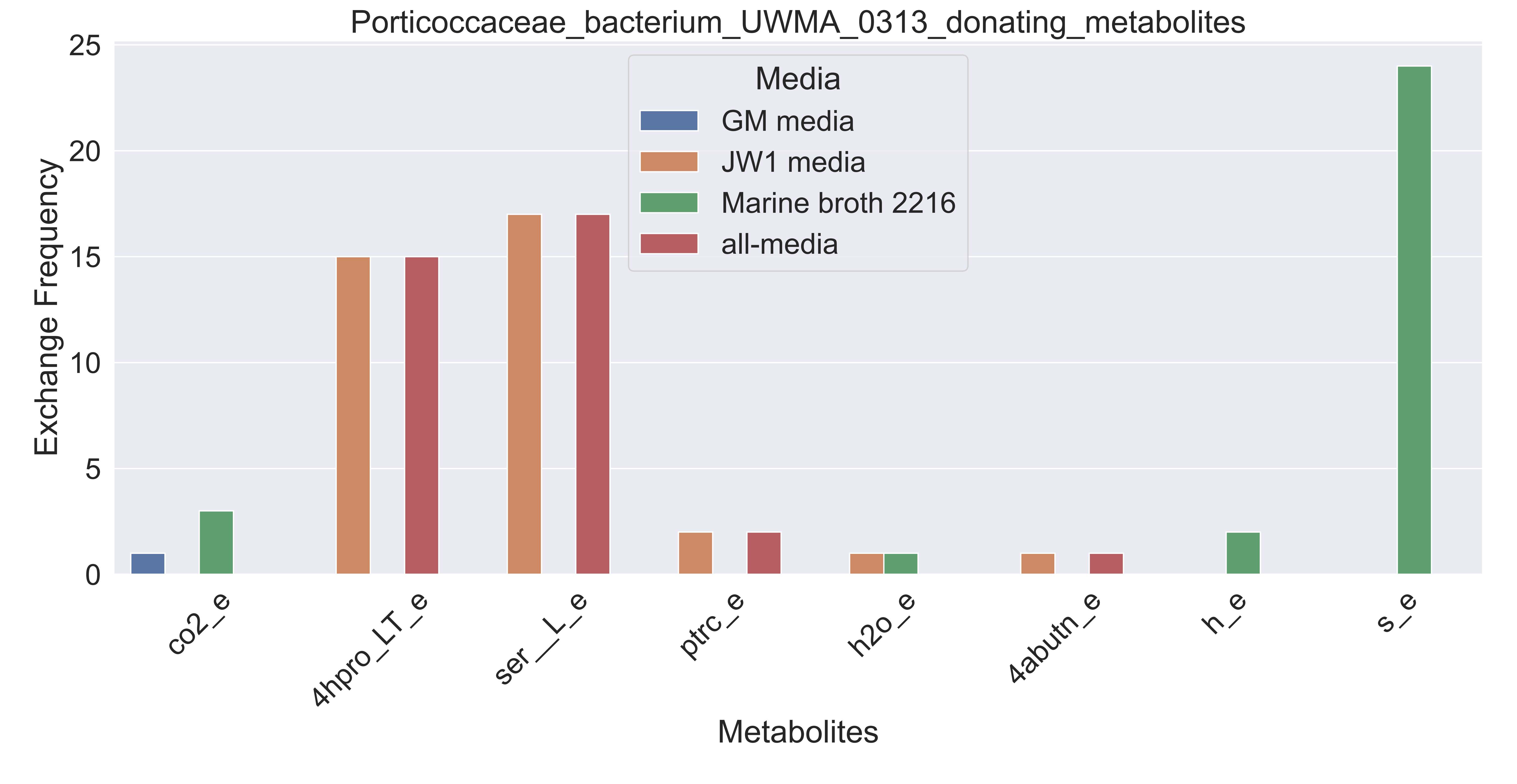

### Supplementary Figure S190 Rhodospirillales bacterium UWMA 0295 donating metabolites.pdf

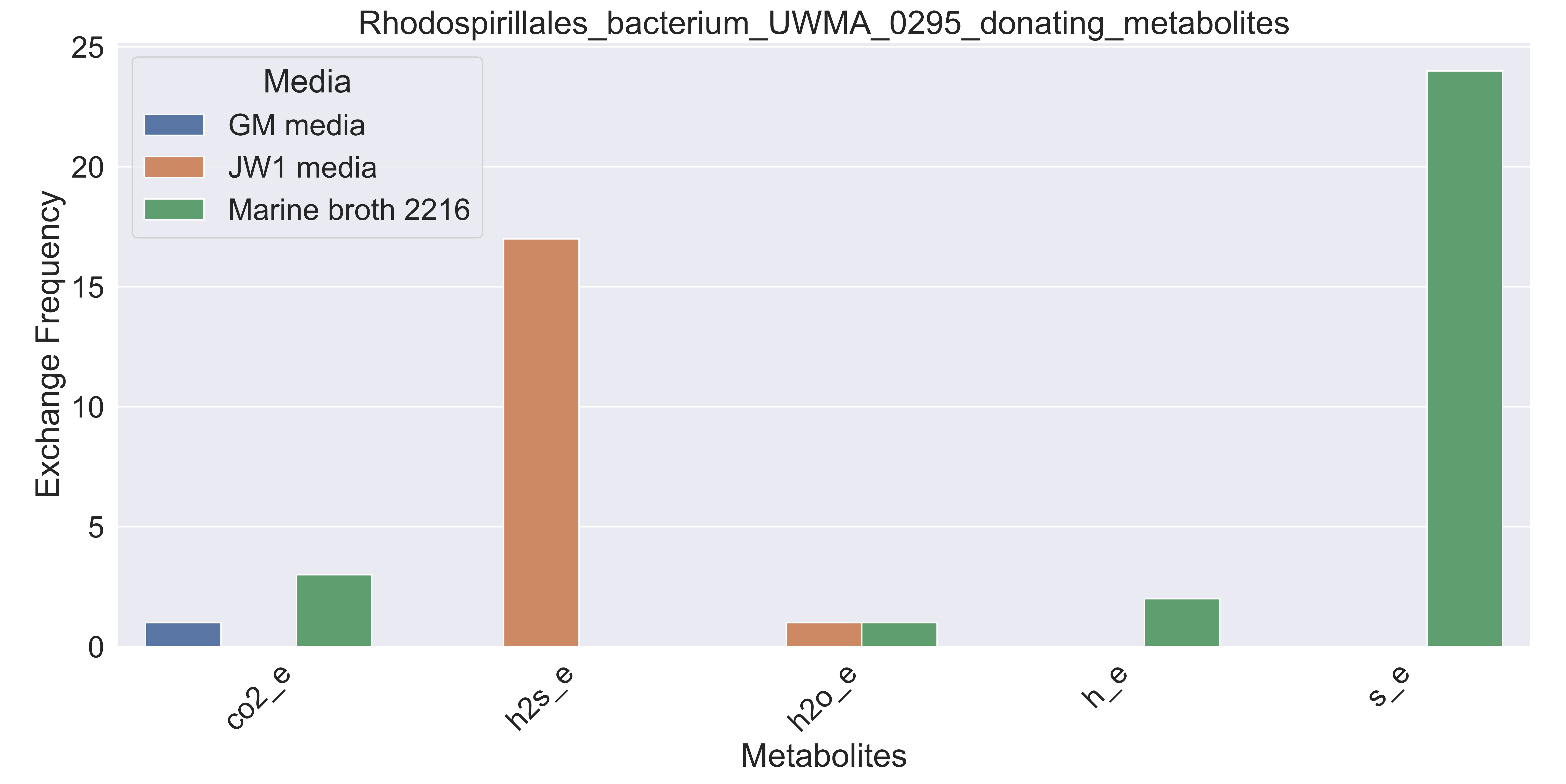

### Supplementary Figure S191 Sneathiellales bacterium UWMA 0353 donating metabolites.pdf

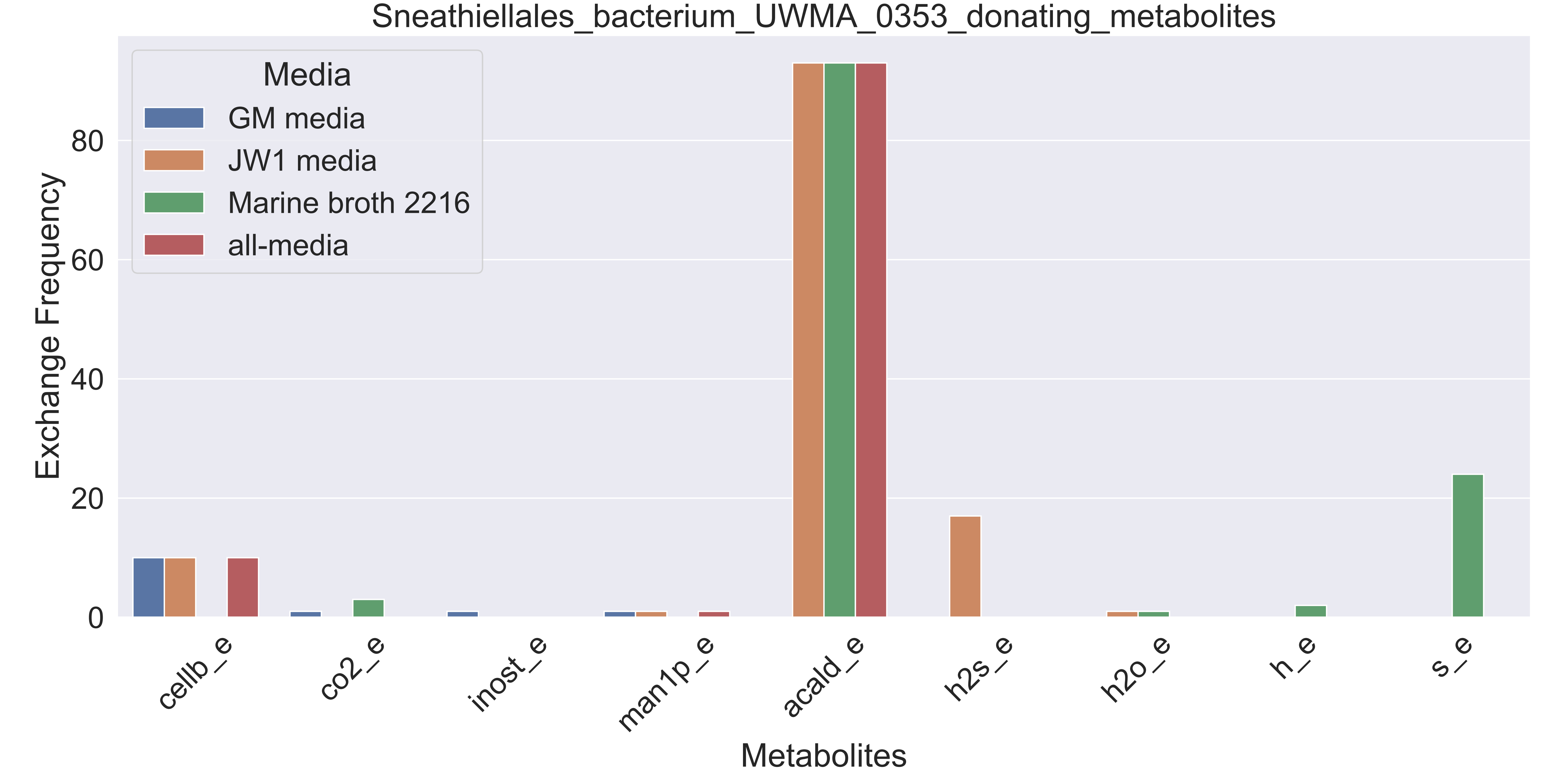

### Supplementary Figure S192 Sulfitobacter sp UWMA 0305 donating metabolites.pdf

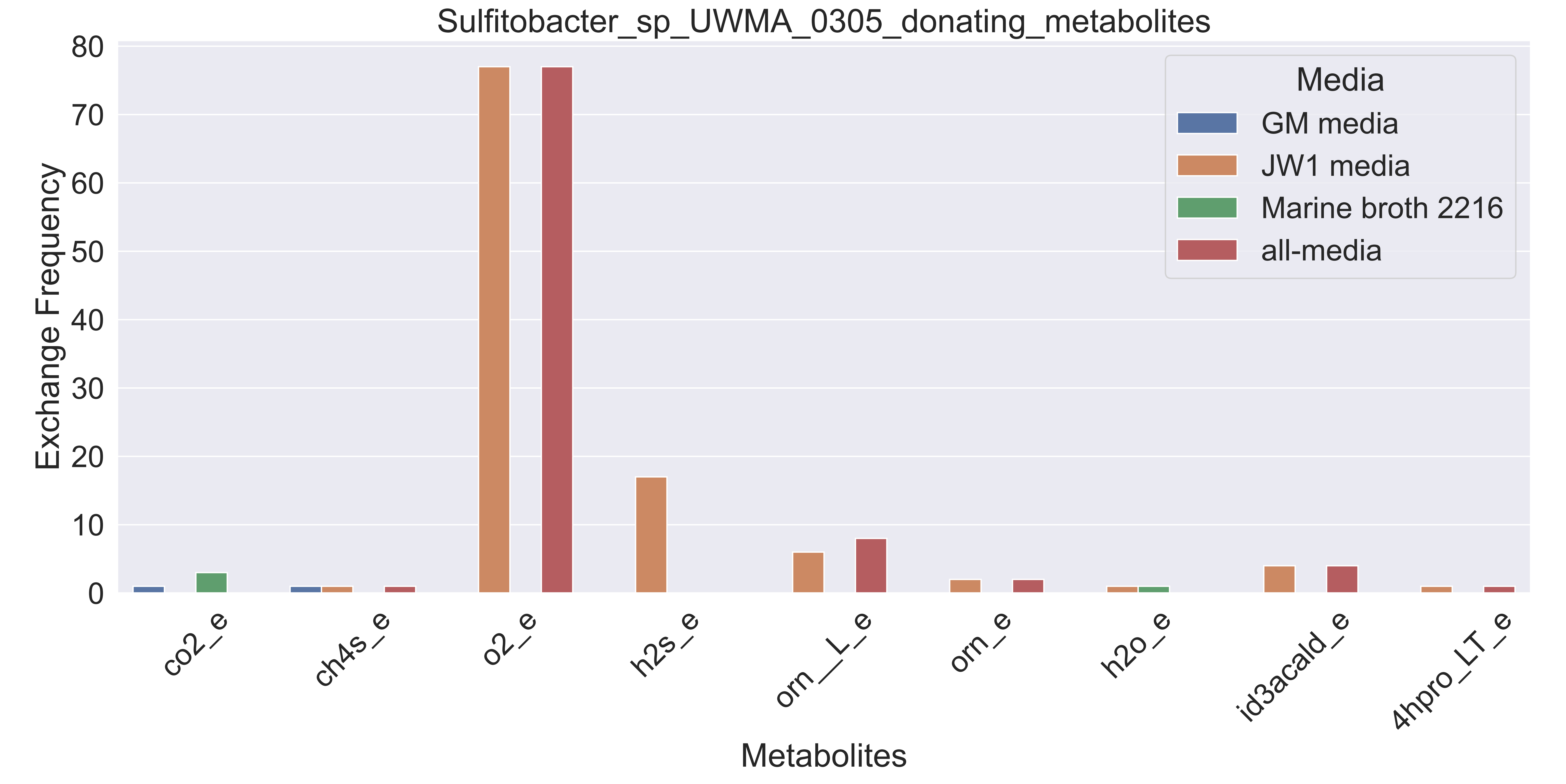

### Supplementary Figure S193 Thiotrichaceae bacterium UWMA 0311 donating metabolites.pdf

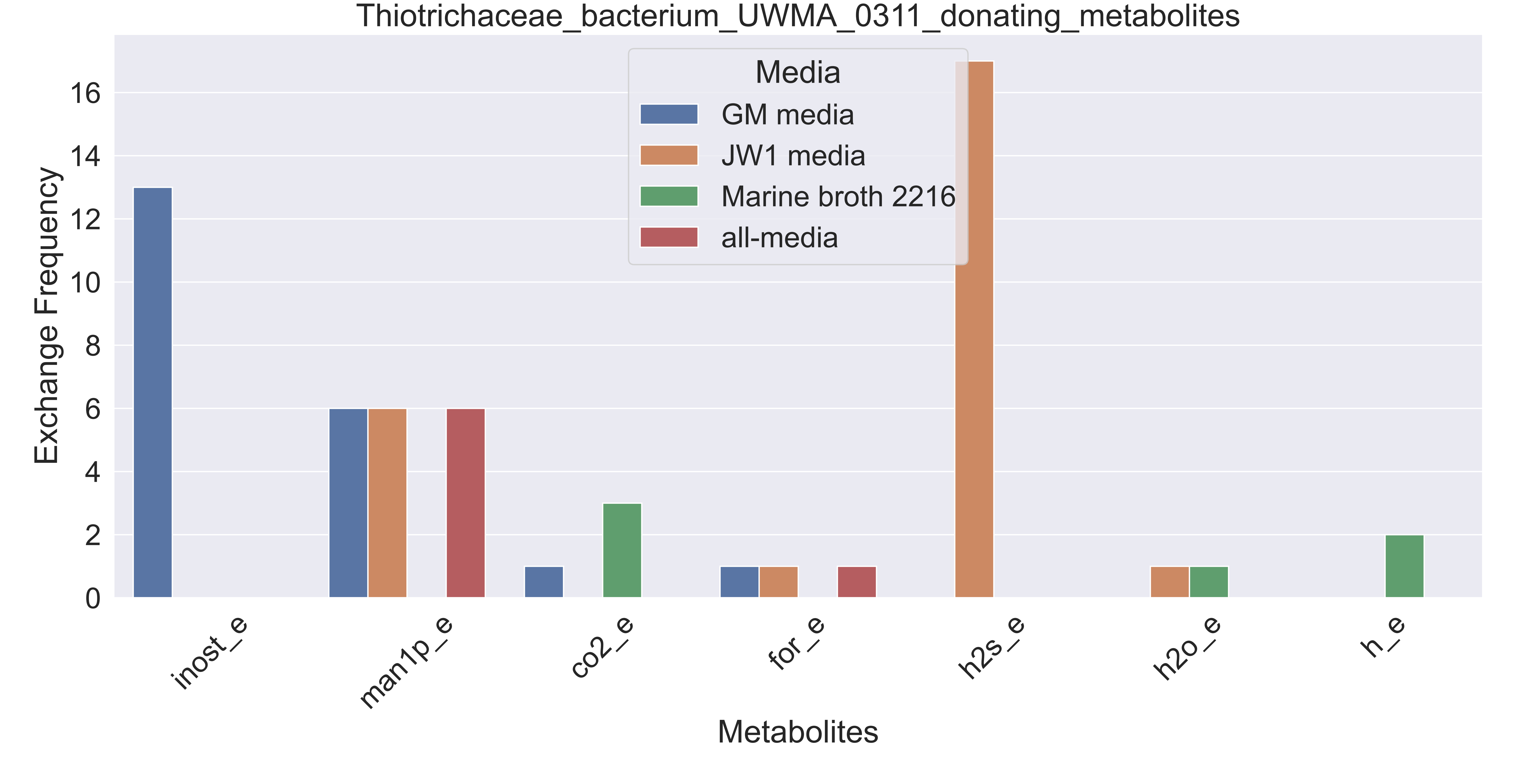

### Supplementary Figure S194 Verrucomicrobiales bacterium UWMA 0292 donating metabolites.pdf

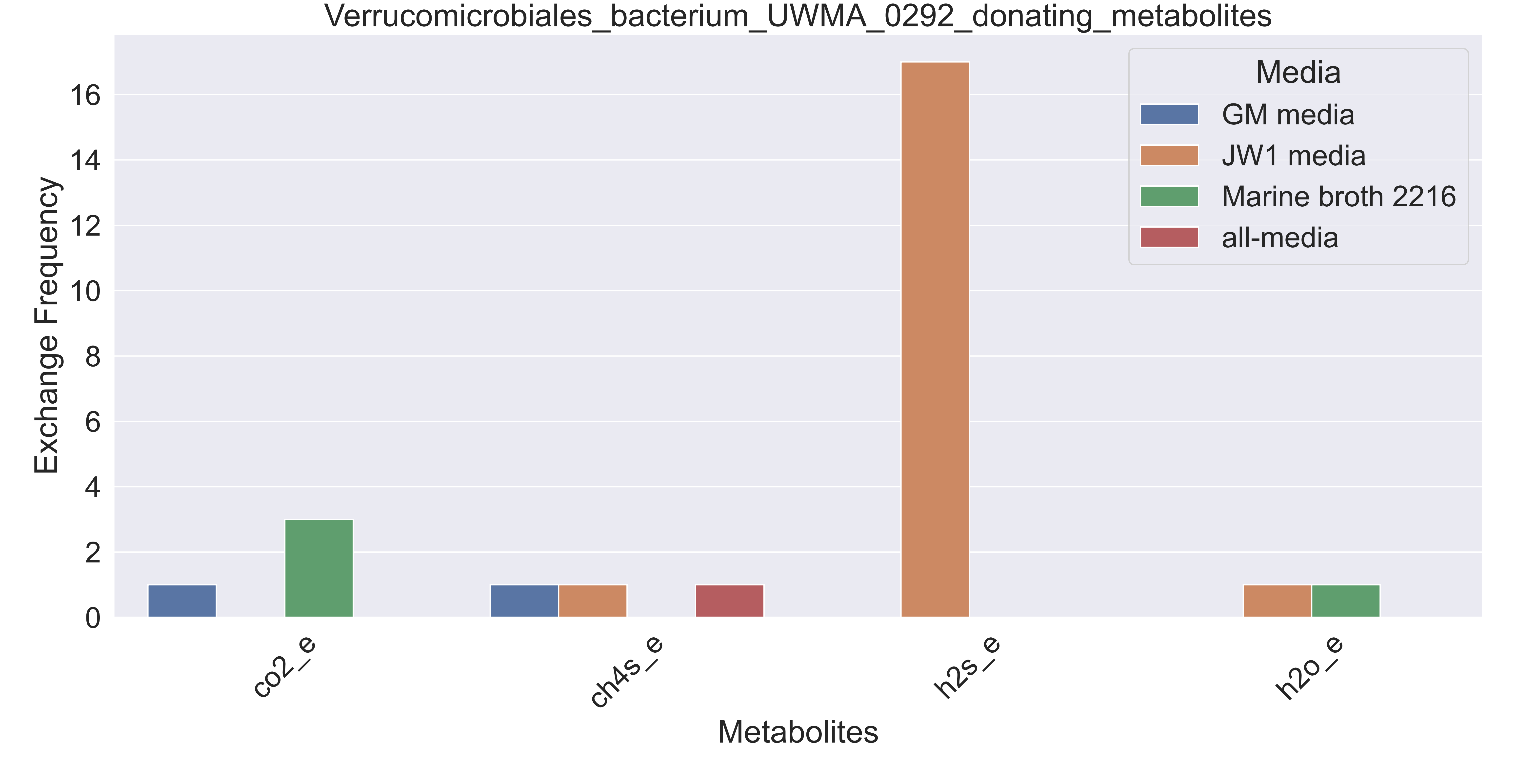

### Supplementary Figure S195 Verrucomicrobiales bacterium UWMA 0320 donating metabolites.pdf

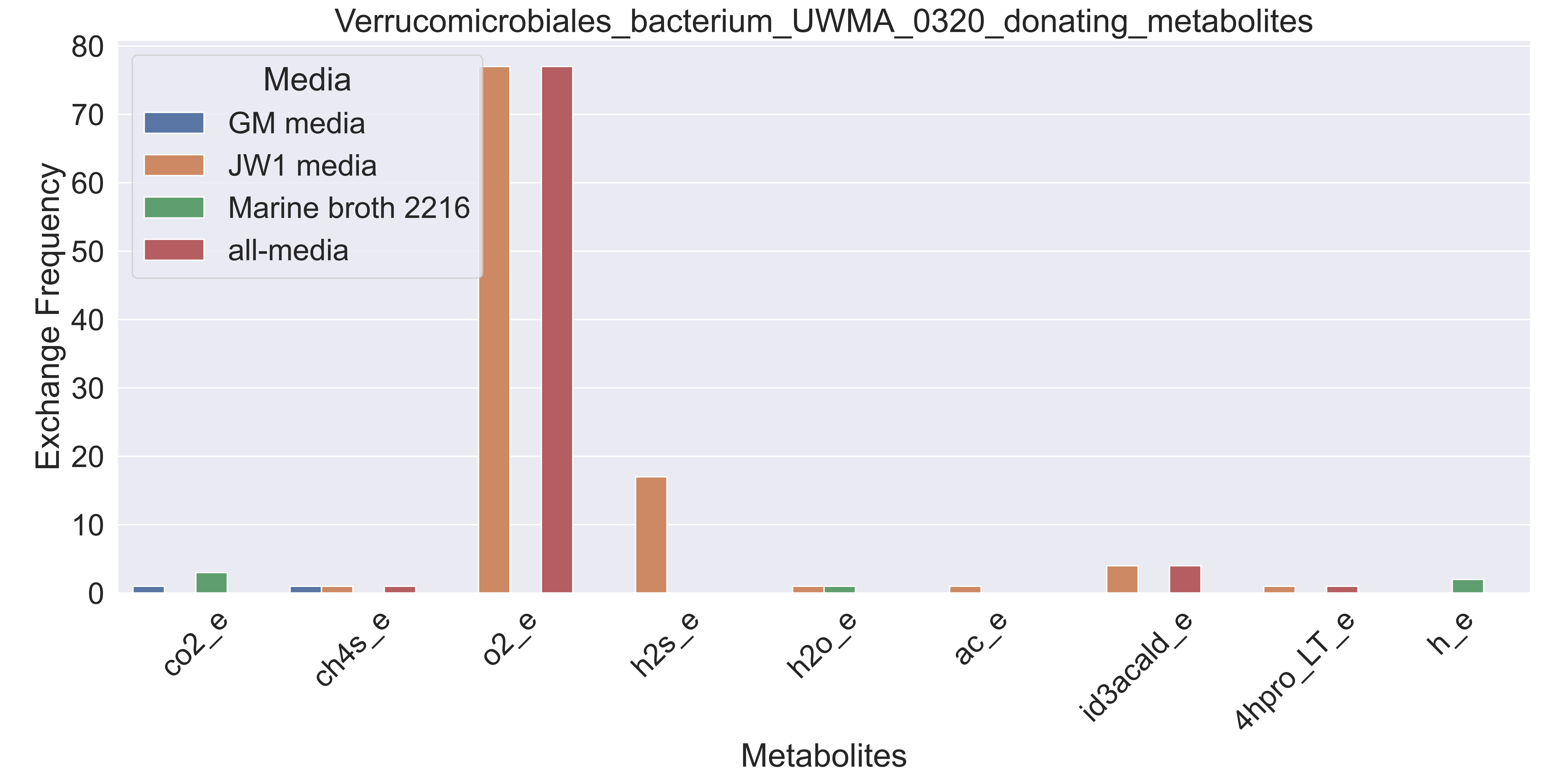

### Supplementary Figure S196 Verrucomicrobiales bacterium UWMA 0332 donating metabolites.pdf

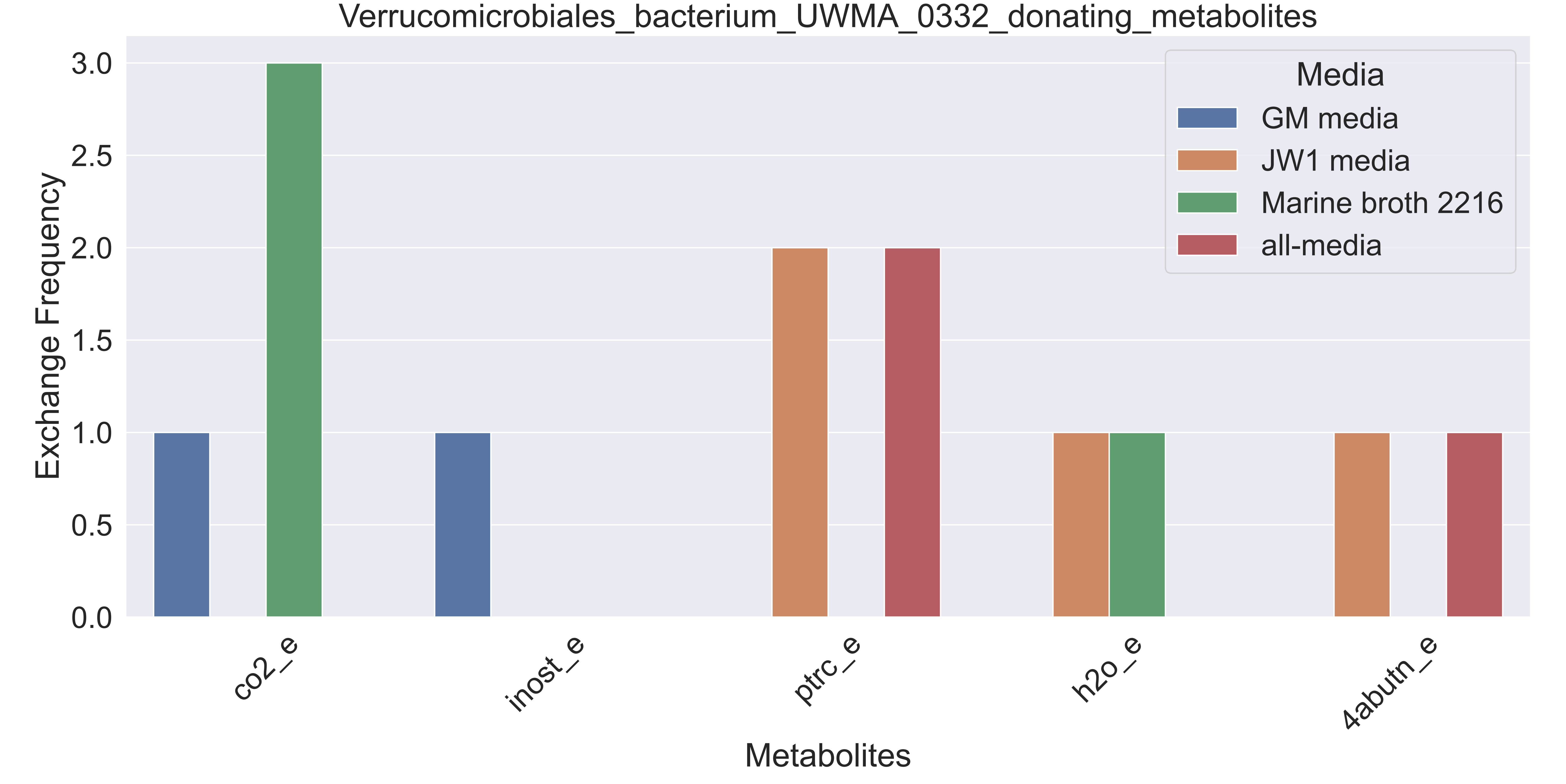
